# LAM/TREM2 ^+^ macrophages release extracellular vesicles and extracellular lipid droplets which modulate the phenotype of recipient macrophages and homeostasis of skeletal muscle cells

**DOI:** 10.1101/2025.01.26.634912

**Authors:** Stefano Tacconi, Anna Maria Giudetti, Ferdinand Blangero, Emmanuelle Meugnier, Assia El-jaafari, Serena Longo, Federica Angilé, Francesco Paolo Fanizzi, Laurence Canaple, Audrey Jalabert, Elizabeth Errazuriz-Cerda, Christel Cassin, Maurizio Zuccotti, Andrea Alfieri, Claire Crola Da Silva, Camille Brun, Benjamin Gillet, Sandrine Hughes, Jennifer Rieusset, Hubert Vidal, Luciana Dini, Sophie Rome

**Affiliations:** CarMeN Laboratory (UMR INSERM 1060/INRA 1397), Lyon-Sud Faculty of Medicine, University of Lyon, Pierre-Bénite, FRANCE; Department of Biological and Environmental Sciences and Technologies (Di.S.Te.B.A.), University of Salento, 73100 Lecce, ITALY; Ecole Normale Supérieure de Lyon, SFR BIOSCIENCES UAR3444, PBES, Lyon, FRANCE; Centre d’Imagerie Quantitative Lyon Est (CIQLE), Lyon 1 University, Lyon, FRANCE; Laboratory of Biology and Biotechnology of Reproduction, Department of Biology and Biotechnology ‘Lazzaro Spallanzani’, University of Pavia, ITALY; Centro Grandi Strumenti, University of Pavia, ITALY; Institut de Génomique Fonctionnelle de Lyon (IGFL), UCBL, ENS de Lyon, CNRS UMR 5242, Lyon, FRANCE; Department of Biology and Biotechnology “C. Darwin”, Sapienza University of Roma, Roma, ITALY; Research Center of Nanotechnologies for Engineering (CNIS), Sapienza University of Roma, Roma, ITALY

**Author notes:** Corresponding authors, &.

**Keywords:** lipid-associated macrophages, extracellular vesicles, lipid droplets, TREM2, diabetes, skeletal muscle

## Abstract

The polarization of tissue-resident macrophages is influenced by a variety of signals from the immune system and the local tissue environment, including nutrition. Although it is known that the quality and quantity of ingested lipids have a significant effect on the lipid composition of extracellular vesicles and their fate, it is unknown how the nutritional environment modifies the release and the function of macrophage-derived EVs. In this study, we used a combination of palmitate and oleate (1:2, FFA) to generate lipid-associated TREM2-expressing macrophages (LAM/TREM2^+^) *in vitro*. Using various electron microscopy techniques (TEM, SEM, CryoEM) and fluorophores, we found that FFA overload not only induces lipid storage in LAM/TREM2^+^ macrophages, but also alters their morphology and reduces the diversity and the number of the lipid-derived structures they release. In addition, LAM/TREM2^+^ macrophages accumulated lipid droplets (LDs) below the plasma membrane and we discovered for the first time that they export and disseminate full LDs into their environment, in addition to extracellular vesicles, by using a cellular pathway associated to CD81. The use of ^14^C-palmitate confirmed the presence of ^14^C-triacylglycerols in the large extracellular vesicle pellet. LAM/TREM2^+^ macrophage-derived EVs induced TREM2 and Il-10 expression in recipient M0 macrophages. These data provide potential insights into how dietary factors and metabolic perturbations can shape the functions of macrophage-derived EVs in the context of metabolic diseases such as diabetes and obesity. In addition, LAM/TREM2^+^ macrophage-derived EVs modulated insulin-sensitivity, mitochondrial oxidative capacity, lipid profiles and the expressions of genes encoding extracellular matrix components in recipient skeletal muscle cells. Although previously postulated but never demonstrated, these data also highlight the LAM/TREM2+ macrophage-derived EVs as important players in SkM tissue renewal and metabolic homeostasis.

## INTRODUCTION

Macrophages are versatile immune cells playing critical roles in tissue homeostasis and repair. Being professional phagocytes, they clear tissues from cellular debris, and apoptotic cells during tissue remodelling and development. They also produce both pro- and anti-inflammatory cytokines during the initiation, maintenance, and resolution of inflammation. In addition, macrophages also release extracellular vesicles (EVs) (**1**). EVs are lipid-derived nanovesicles generated either from the plasma membrane (PM) budding (*i.e*; Microvesicles, 100-400nm) or from the endolysosomal system in which intraluminal vesicles progressively accumulate inside acid and bis(monoacylglycerol)phosphate-enriched endosomes to form multivesicular bodies (MVBs). By fusion with PM, some MVBs release their intraluminal vesicles into the microenvironment (*i.e*.; exosomes, 50-110nm) (**2**). Until now, it has been difficult to obtain pure preparations of exosomes or microvesicles, and the International Society for Extracellular Vesicles (ISEV) recommends the terminology of large EVs (lEVs, >200nm) to consider EVs mainly derived from PM, and small EVs (sEVs, <200nm) for EVs enriched in exosomes or small PM budding (**3, 4**). All EVs contain cargoes of lipids, proteins, DNA and RNA. Upon release, EVs are taken up by neighbouring cells where they modulate gene expression and thus participate in the regulation of tissue homeostasis (**5**). Because EV cargoes simultaneously affect multiple signalling pathways, they have a more potent and complex influence on the homeostasis of recipient cells than individual lipids, hormones, or cytokines. EVs also have a trophic role supplying recipient cells with lipids, proteins and micronutrients (**6, 7**).

The polarization of tissue-resident macrophages is influenced by a variety of signals from the immune system and the local tissue environment, including nutrition. Diets rich in unsaturated fats and low-gIycaemic index carbohydrates support the M2-phenotype. In contrast, saturated fats and refined carbohydrates promote the pro-inflammatory M1-phenotype (**8**). However the balance between M1 and M2 macrophages is altered during the development of obesity. Pro-inflammatory immune cells predominate in early obesity, whereas non-resident anti-inflammatory macrophages predominate in chronic obesity (**9**),(**10**). How this phenotypic shift affects the biological activity of macrophage-derived EVs is poorly understood. Recently, we have showed that an hyper glucose environment polarized macrophages into glycolytic macrophage M1 which released EVs able to polarize activated macrophages to M2 (**11**). Similarly, in a mouse model of endometriosis, M1-derived EVs could repolarize macrophages into M2, thereby reducing the inflammation associated with the pathology (**12**). These two studies demonstrated for the first time that the dialog between macrophages via the EV pathway is an integral part of the immune response and that M1-derived EVs can mitigate inflammation. In addition, it was shown that M2-derived EVs reduced inflammatory responses in ulcerative colitis in mice (**13**), but worsened the progression of non-small cell lung cancer by promoting cell growth in a hypoxic microenvironment (**14**). Taken together these data suggest that M2-derived EVs, may favor an anti-inflammatory ‘M2-like environment’.

However, this conclusion is partial as there is no data on the role of M2-derived EVs on macrophage polarization. Furthermore, although it is known that the diet, and in particular the quality and quantity of ingested lipids, has a significant effect on the lipid composition of EVs and their fate (**6**), it is unknown how the nutritional environment modifies the release and the function of macrophage derived-EVs. In this context, a subset of macrophages has recently been identified, the lipid-associated TREM2-expressing macrophages (LAM/TREM2^+^) which arise from circulating monocytes under obesity conditions and are located around enlarged adipocytes (**6, 9**). Through active phagocytosis they remove free-fatty acids (FFA), lipoproteins and cholesterol which are stored as lipid droplets (LDs) to reduce the effects of adipose tissue lipotoxicity. TREM2 macrophages are also detected in the liver under high-fat diet conditions (**10**) and show similarity to brain DAM/TREM2^+^ macrophages (**15**)

In this study, we have investigated how an FFA-enriched environment modulates the metabolism and phenotype of macrophages, and consequently the properties and composition of their released EVs. In addition to the crosstalk between macrophages through the EV exchange, we have determined whether FFA-EVs could affect insulin-sensitivity and lipid composition of skeletal muscle (SkM) recipient cells. The present study reveals unexpected new mechanisms to explain how LAM/TREM2^+^ release lipids in their environment to recruit new LAM/TREM2^+^ macrophages and to modulate the metabolic homeostasis of SkM cells.

## MATERIAL AND METHODS

### Cell cultures and treatments

THP-1 monocytes (ATCC TIB-202) were grown in RPMI-1640 and C2C12 myoblasts (CRL-1772™) in DMEM 4.5g/L glucose, both supplemented with 10% FBS, 2mM L-glutamine, 100IU/mL penicillin/streptomycin (Sigma-Aldrich) (37°C, 5% CO_2_). THP-1 differentiation into M0 macrophages was induced with 100ng/ml Phorbol 12-Myristate 13-Acetate (72 hours). Confluent C2C12 cells were differentiated into myotubes in DMEM with 2% horse serum and used 7 days post-differentiation. To mimic a hyperlipidemic conditions, M0 were treated for 24h either with oleate+palmitate (2:1 in BSA-free fatty acids, 500 µM final) (FFA-condition) or with BSA (100 µM) (control-condition). Myotubes or macrophages were treated with 2µg/mL of macrophage-EVs. After 24h of treatment, myotubes were starved for 4h to quantify insulin-stimulated AKT phosphorylation or GLUT4 translocation (10min, 100nM insulin, Sigma-Aldrich).

### Bone marrow-derived macrophages

Blood samples were from the Lyon Blood Bank (France), following institutionally approved guidelines. Monocytes were harvested from healthy human peripheral blood by density gradient centrifugation (Ficoll-Histopaque, Sigma-Aldrich). Monocytes were stored in liquid nitrogen prior to use.

### MTT assay

Macrophages were incubated with 1 mg/mL of MTT in RPMI-1640 for 2h and were washed three times in PBS (0.2 M, pH 7.4) and the reduced MTT formazan crystals were solubilized with DMSO (Carlo Erba, IT). The absorbance was read at 570 nm.

### EVs isolation and characterization

EVs were purified from the conditioned medium (CM) of mycoplasma-free macrophages grown in EV-depleted medium by differential centrifugation. Briefly, CM were centrifuged successively at 500g (10min, RT), 800g (10min, RT), and 2,000g (20min, RT), to remove dead cells, apoptotic bodies and aggregates. The supernatant was centrifuged at 20,000g (30min, 4°C). The lEV-enriched pellet was rinsed and resuspended in calcium/magnesium-free PBS and the supernatant was filtered (0.22 µm polyethersulfone filter units, Thermo Fisher Scientific). The filtered supernatant was centrifuged at 100,000g (70min, 4°C) (Beckman Coulter Ultracentrifuge Optima XE, fixed angle rotor) to collect the sEV pellet which was rinsed and resuspended in calcium/magnesium-free PBS. EV proteins were quantified by Bradford protein assay (Biorad) and EV size distributions by Dynamic Light Scattering (DLS, Malvern). Refractive index and viscosity of dispersant were 1.332 and 1.029 cP, respectively, at 20°C (**16**).

### Electron microscopy

Macrophages were fixed in 2.5% glutaraldehyde, 0.1 mol/L cacodylate (pH 7.4) for 1h and post-fixed in 1% OsO_4_ in the same buffer for 2h. Cells were dehydrated with ethanol (25%, 50%, 70%, 90%, 100%) and embedded in Spurr resin. 60nm sections were examined by transmission electron microscopy (TEM). EVs were fixed in 0,1% paraformaldehyde for 30 min, RT. Fixed EVs were stained with 2% uranyl acetate (7min, RT), and loaded onto 200 mesh carbon-coated grids. Both EVs and macrophages were visualized with a Zeiss Auriga (Zeiss, DE) equipped with STEM module at 20 kV.

For scanning electron microscopy (SEM) macrophages grown on glass coverslips were fixed as above and dehydrated with acetone (25%, 50%, 70%, 90% and 100%) followed by Critical Point Dryer CPD EMITECH K850 (Quorum Technologies Ltd, UK). Stub-mounted specimens were prepared with a Balzers Union SCD 040 (Balzers Union) and examined under a Zeiss EVO HD 15 scanning microscope. For cyoelectron microscopy (CryoEM), Ssamples were prepared in a Vitrobot instrument (Thermo Fisher Scientific) as follows: after glow discharging the carbon/copper grids (Q C-Flat 1.2/1.3 Cu 300 or Lacey carbon TH Cu 200), 4 µL of EV sample (1:10 dilution in filtered PBS) was blotted once for 3 to 6 sec at 4 °C and 95% relative humidity. Loaded grids were immediately plunged into liquid ethane. Images were acquired under low dose conditions (30 e-/Å2) at several different magnifications (73000 to 120000x) and −3.0 μM defocus value with the software EPU (Thermo Fisher Scientific, USA) on a 200kv Glacios cryo-transmission electron microscope (Thermo Fisher Scientific, USA) equipped with a Falcon 3EC direct electron detector (Thermo Fisher Scientific, USA).

### Western and Dot Blot analyses

Cells were lysed in NaCl 150mM, Tris-HCl 50mM pH=8, MgCl2 2mM, 0,1% SDS, 0,5% deoxycholic acid, 1% NP40, containing PMSF and protease inhibitors (ThermoFisher Scientific) and sonicated for 5min. Insoluble material was pelleted at 13,000g, 4°C. Proteins from the supernatant were separated by SDS-PAGE and transferred onto nitrocellulose membranes. For Dot Blot analyses, proteins were directly spotted onto nitrocellulose membranes and completely dried for 30min at RT. Membranes were saturated in 5% milk or 3% BSA in Tris-buffered saline with 0.1% Tween20 (TTBS) (1h, RT), and were incubated overnight at 4°C with each primary antibody (**Table S1-A**). After washing in TTBS, membranes were incubated for 1h at RT with horseradish peroxidase-conjugated antibodies. Bands were detected by chemiluminescence (ECL reagent; Merck Millipore) and quantified by densitometry. All original blots are provided in **Fig. S1**.

### RNA sequencing and qRT-PCR

Total RNA was extracted with Trizol (Invitrogen, CA). Genomic DNA was removed (DNA-free DNA removal kit, ThermoFisher Scientific). PolyA RNAs purified from 1µg of total RNA (NEBNext Poly(A) mRNA Magnetic Isolation Module (NEB)) were used to build barcoded libraries (CORALL mRNA-Seq Library Kit (Lexogen)). Libraries were pooled in equimolar ratios and sequenced on Illumina NextSeq 500 sequencer in single end (84 bp, High Output run) (IGFL-ENS Lyon-France sequencing platform). At least 24 million reads per sample were obtained (GEO #GSE199222). For qRT-PCR, 1.5 µg of RNA was reverse transcribed into cDNA using the single-step SuperScriptTM IV kit (Invitrogen, CA). PCR was performed by using the CFX ConnectTM Real-Time PCR Detection System (Biorad, CA) (primers are in **Table S1-B**).

### Bioinformatic analyses

RNAseq data were analysed with the pipeline of Lexogen for the CORALL kit on the BlueBee Genomics platform. After removing reading artifacts, sequences were aligned with STAR Aligner to eliminate PCR duplicates. Clean sequences were re-aligned to retrieve transcript number of counts by using Mix2. After removal of all non-expressed genes, analyses were performed on the count tables by using Rstudio and DESEQ2 (V_1.30.1). Statistical analyses were carried out using a linear regression model including conditions to detect differences between 2 conditions and using contrast to extract comparisons of interest. The *p* values were adjusted by using the Benjamini and Hochberg procedure. PCA plots were performed by using ClustVis 2.0 (https://biit.cs.ut.ee/clustvis/). Significantly enriched Gene Ontology pathways were identified by gprofiler (http://biit.cs.ut.ee/gprofiler/gost) considering only the annotated genes. Heatmaps were performed with MORPHEUS (https://software.broadinstitute.org/morpheus/). Cellular pathways and protein networks were determined with STRING 12.0 (https://string-db.org/).

### Intracellular neutral lipid staining with Oil Red-O

Macrophages differentiated onto 11mm glass coverslips were washed in PBS, fixed in 10% formalin (5min), washed with 60% isopropanol (5min) and left at RT until completely dry. Cells were treated with 0.5% Oil Red-O/isopropyl alcohol (20min), and washed in PBS. Nuclei were stained with 1M DAPI (20min). Intracellular fluorescence was visualized with a Leica DMi8 Thunder Imager 3D cell culture microscope and analysed with Fiji 2.15.1.

### Thin Layer Chromatography (TLC)

Lipids were extracted by the Bligh and Dyer procedure. Lipids loaded on silica gel plates were separated with hexane/ethyl ether/acetic acid (70/30/1; v/v/v) for neutral lipid, with chloroform/methanol/water (65/25/4; v/v/v) for polar lipids or toluene/methanol (70/30; v/v/v) for sphingolipids. After separation, plates were sprayed with 10% cupric sulfate in 8% aqueous phosphoric acid, dry (10min, RT), and heated at 145 °C (10min). Identification of lipid species was made by migrating and developing specific standards (11). Spot intensity was measured by densitometry.

### 1H-NMR spectroscopy analysis

All measurements were performed on a Bruker Avance III 600 Ascend NMR spectrometer (Bruker, Biospin Milan) operating at 600.13 MHz for ^1^H observation, equipped with a TCI cryoprobe incorporating a z-axis gradient coil and automatic tuning-matching. Experiments were acquired at 300K in automation mode after loading individual samples on a Bruker Automatic Sample Changer, interfaced with the software IconNMR (Bruker). Lipid extraction and spectra analyses (2D ^1^H Jres, ^1^H COSY, ^1^H-^13^C HSQC) were performed as previously (11).

### GLUT4 Immunostaining

C2C12 were fixed in 4% formalin (30min, RT), incubated in 3% (w/v) BSA (30min, RT) and then with anti-GLUT4 (4°C, ON). Following exposure to FITC-conjugated secondary antibodies (1:800 in 3% BSA), cells were counterstained with TexasRED-phalloidin (#T7471, Thermo Fisher Scientific) for F-actin staining and DAPI for nuclear staining. Samples were observed under an Axio Observer fluorescence microscope (Zeiss) equipped with Apotome 3.

### Radiolabelled-EV isolation

To produce radiolabelled-EVs, 6×10^6^ THP-1 were treated with FFA supplemented with ^14^C-palmitate for 4h in RPMI (0.005 μCi/ μmol of FFA mix, [1-^14^C]Palmitic acid #NEC075H050UC, Revvity). After validation that macrophage intracellular lipids were radiolabelled (Fig. S2A), they were washed thoroughly to remove exogenous radioactivity and were incubated with a fresh medium for 24h. EVs were purified from CM (as described above). Of note: to avoid radioactivity contamination of the incubator, the lids of the flasks were carefully closed which may have induced a state of hypoxia, but without altering TAG accumulation inside THP-1 (Fig. S2A). After TLC separation of cell lipids, radioactivity was counted in each lipid fraction with a Beckman Coulter LS 6500.

### Flow Cytometry

Cells were washed in PBS and surface antigens were labeled with antibodies for 30 min at 4°C (**Table S1-A**). After washing cells were resuspended in FACS staining buffer. Analyses were performed with a LSR II cytofluorometer (BD Bioscience). Gating was based on forward and side scatter properties while excluding debris. Events within gates were then analyzed for specific expression of surface antigens and compared to the corresponding isotypic controls. Markers were placed to identify the positive cells. Data were analyzed using the FCS Express software (Fig. S4A).

### Bioenergetic measurements using the Seahorse analyzer

Oxygen consumption rate (OCR, pmoles/min) was measured using a Seahorse XFe 24 metabolic flux analyzer (Agilent Technologies, CA) as previously described (**11**).

### Statistical analysis

Results are expressed as means ± SD. Data normality, comparisons between groups and visualization of the data were performed with GraphPad Prism 9 software (Graphpad Software Inc, San Diego, CA). Differences between groups were considered statistically significant for *p* < 0.05. The statistical test used for each analysis is described on the figure legends.

## RESULTS

### FFA-overloaded macrophages switch to a LAM/TREM2^+^ phenotype

To understand how FFA affects EV release and lipid composition, we first determined the consequences of FFA overload on macrophage metabolism and phenotype. TEM images (Fig. 1A, Fig. S3B-C) and fluorescence labelling of neutral lipids (Fig. 1B) showed a strong accumulation of lipid droplets (LDs) inside FFA-treated THP-1 and human bone marrow-derived macrophages *vs* CTR. Lipid profiles confirmed the strong enrichment in triacylglycerols (TAG), diacylglycerols (DAG) and FFA, in FFA-treated macrophages *vs* untreated (Fig. 1C). In parallel, the expressions of DGAT1 and DGAT2 which are involved in TAG synthesis were increased (Fig. 1D-E). LD areas ranged from 0.01 to 2.5 µm^2^ (Fig. S2B) and their accumulation did not affect macrophage viability (Fig. S2C). Since LDs and mitochondria work closely together to balance lipid storage and energy production, we hypothesized that macrophages might accumulate LDs due to impaired fatty acid oxidation. Consistent with this, no difference in the basal respiration rate was observed in FFA-treated macrophages *vs* CTR (Fig.1 F-H). The small increase in mitochondrial oxygen consumption (Fig.1G) and in spare respiratory capacity (Fig. I) observed in FFA-treated macrophages indicated that they tend to accumulate lipids rather than oxidize them efficiently in contrast to tumor-associated macrophages which also accumulate lipids (**17**). Taken together, these data indicated that FFA-treated macrophages had characteristics of lipid-associated macrophages (LAM) (9). Indeed, they had increased expression of TREM2 (Fig. J-K), expressed CD9 (Fig. 1L), and increased expression of FABP4, CD36, PLIN2, and ABCA1 (Fig. 1M-P), a transcriptional signature identified in LAM from adipose tissue (10). Furthermore, FFA-treated macrophages exhibited an immunosuppressive phenotype, *i.e*.; increased release of the cytokine IL-10 (Fig. S2-D), without modulating the pro-inflammatory cytokine IL-1β (Fig. S2-E), a phenotype observed in chronic inflammation involving TREM2^+^ macrophages.

**Figure 1:**
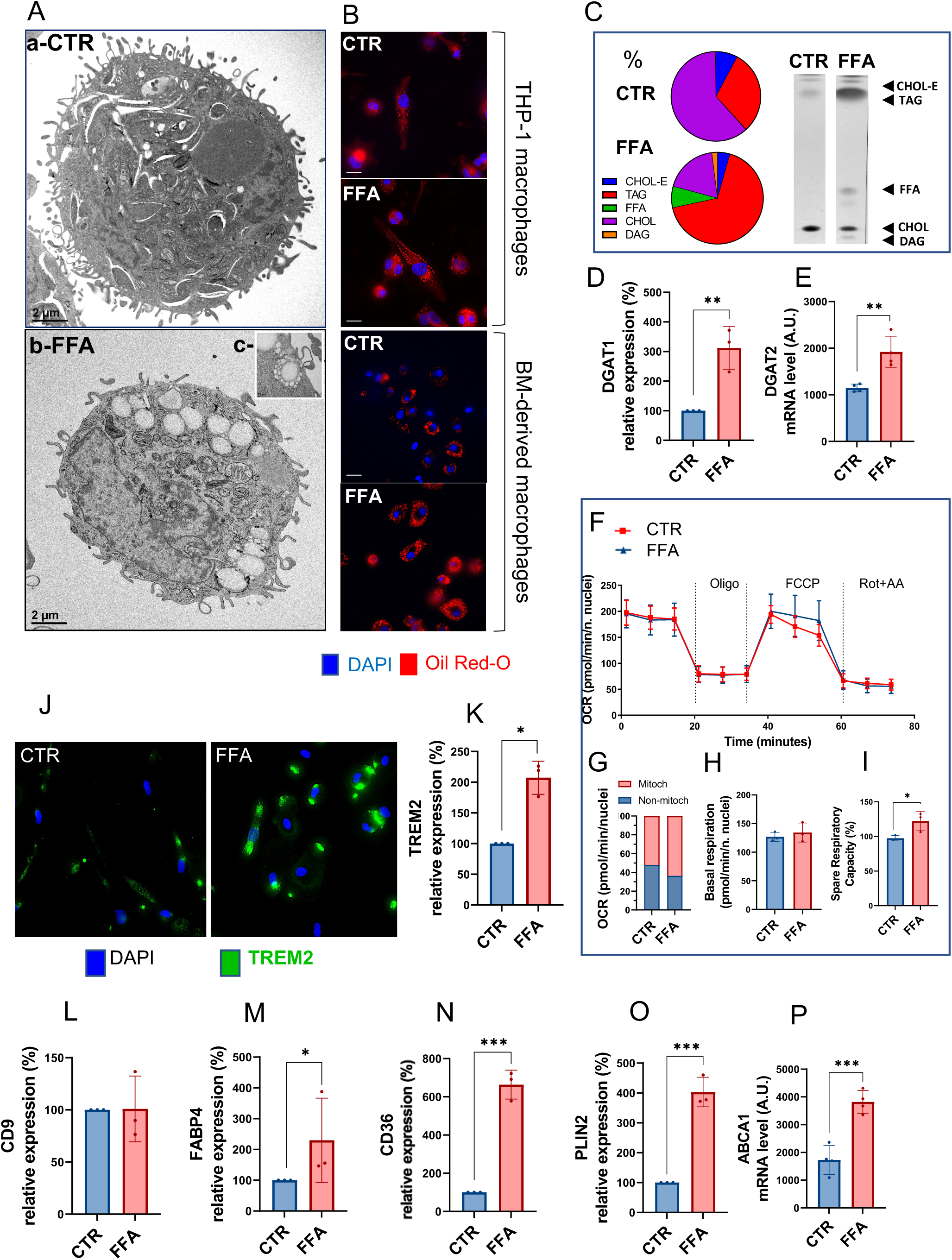
THP-1 and Bone marrow-derived macrophages have a LAM/TREM2^+^ phenotype in response to FFA. **A**- BSA-treated (**a**) and FFA-treated (**b**) THP-1 macrophages, visualized by TEM; (**c**) magnification showing the fusion of small LDs to generate large LD (original image in S3-B). **B**- THP-1 macrophages and human bone marrow (BM)-derived macrophages treated for 24h either with BSA or FFA. After fixation, neutral lipid accumulation was detected by Oil Red O (red, 557-576nm) and nucleus with DAPI (blue, 350-465nm). **C**- lipid profiles of THP1 macrophages analysed by TLC. Data are expressed as percentage of the total (TAG: triacylglycerol; FFA: free fatty acid; CHOL: cholesterol; CHOL-E: cholesterol ester, DAG: diacylglycerol). Right, representative TLC profile. **D**- DGAT1 protein level quantified by WB, expressed as % of CTR. **E**- DGAT2 mRNA levels determined by qRT-PCR (Arbitrary Unit). **F-I**, Bioenergetic measurements of THP-1 macrophages in response to FFA *vs* CTR using the Seahorse analyzer. Measurements were normalized by the number of nuclei/well. **F-** Oxygen consumption rate (OCR) profiles (Oligo = Oligomycin; Rot = Rotenone; AA = Antimycin A), **G-** Mitochondrial and non-mitochondrial oxygen consumption, **H-** Basal respiration, and **I**- Spare respiratory capacity. OCR and spare respiratory capacities are expressed as % of CTR. **J**- TREM2 Immuno-detection of non-permeabilized THP-1 macrophages. **K-O**- TREM2, CD9, FABP4, CD36, PLIN2 protein level quantified by WB, expressed as % of CTR. **P**- ABCA1 mRNA levels determined by qRT-PCR (Arbitrary unit). Values are means ± SD (n=3); *p* values are from student *t*-test (FFA *vs* CTR), (*) p<0.05, (**) p<0.01, (***) p<0.001.

### LAM/TREM2^+^ macrophages export a heterogenous population of EVs containing lipid droplets (LDs)

Since modulation of lipid metabolism affects EV biogenesis and function (**6**), we hypothesized that the phenotypic switch to LAM/TREM2^+^ macrophages in response to FFA might also affect EV release. SEM images revealed that altered lipid metabolism in response to FFA was associated with modifications of the macrophage surface (less ‘hairy’) and a suppression of the production of retractosomes involved in cell migration (**18**), *vs* untreated macrophages (Fig. 2A). Overall, macrophages appeared to produce and release fewer lipid-derived structures to the external environment in response to FFA. Different types of EVs were identified by TEM, after differential centrifugation of the macrophage-conditioned medium (Fig. 2B, a-b). Larger vesicles surrounded by a single membrane, around 1μm, were also observed in the 20K pellet (Fig. 2B, c). EVs in the 100K pellet (*i.e*.; sEVs) had a mean size of 190nm, whereas those in the 20K pellet (*i.e*.; lEVs) had a mean size of 250nm (Fig. 2C-D). Both populations showed multilamellar structures (Fig. S3A). After the FFA treatment, both sEV-FFA and lEV-FFA populations had reduced sizes (Fig. 2C-D). Interestingly, DLS analyses identified a second peak within the lEV-FFA population, not observed in the lEV-CTR, with a mean size of 800nm (Fig. 2D) close to the size of the large single-membrane vesicles observed in the lEV-FFA pellet (Fig. 2B, c). SEM images showed numerous blisters on the surface of FFA-macrophages, not detectable on the “hairy” CTR-macrophages (Fig. 2E a-b). We suspected that these extracellularly sorted blisters (Fig. E c) might correspond to LDs, as TAG and DAG were identified in the lEV-FFA lipid profile (Fig. 2F). To confirm this, macrophages were treated with ^14^C palmitate and the lipid profile of the released ^14^C-EVs was determined. Radiolabelled TAG and DAG were found in lEV-FFA but not in sEV-FFA (Fig. 2G) confirming the presence of LDs within the lEV-FFA pellet. Using SEM, we observed LDs in the vincity of FFA-treated macrophages, but not in the vincity of the control macrophages (Fig. 2E d-e). The presence of LDs in response to FFA was also observed by using human BM-derived macrophages treated with FFA (Fig. 2F-f, Fig. 3A), indicating that the release of LDs is not related to the cancerous origin of the THP-1. Finally, the size distribution of the released LDs (Fig. 3B-C) corresponded to the size of the second peak of lEV-FFA (Fig. 2D). TEM images showed that LDs pushed the PM out of the cells (Fig 3D-a), and PM ruptures near LDs were observed (Fig. 3D-b), suggesting that LDs are exported by a different mechanism than PM-derived EVs, such as microvesicles (**19**). To validate this, we performed fluorescent co-labelling of CD81 (predicted to be expressed at the PM) and PLIN2 (marker of LDs) on non-permeabilized THP-1. Surprisingly, CD81 was not uniformly expressed on the surface of macrophages (Fig. 3E a-b, Fig. S3C), and co-localized with PLIN2 at the PM sites of LD release (Fig. 3E c-d). 3D reconstruction revealed a corona of CD81 proteins surrounding LD release sites (Fig. 3F). TEM images of FFA-treated macrophages showed that CD81 was more frequently expressed at or near LDs (Fig. 3G) and also near the ER, the site of LD formation (Fig. S3D-E), than at the PM.

**Figure 2:**
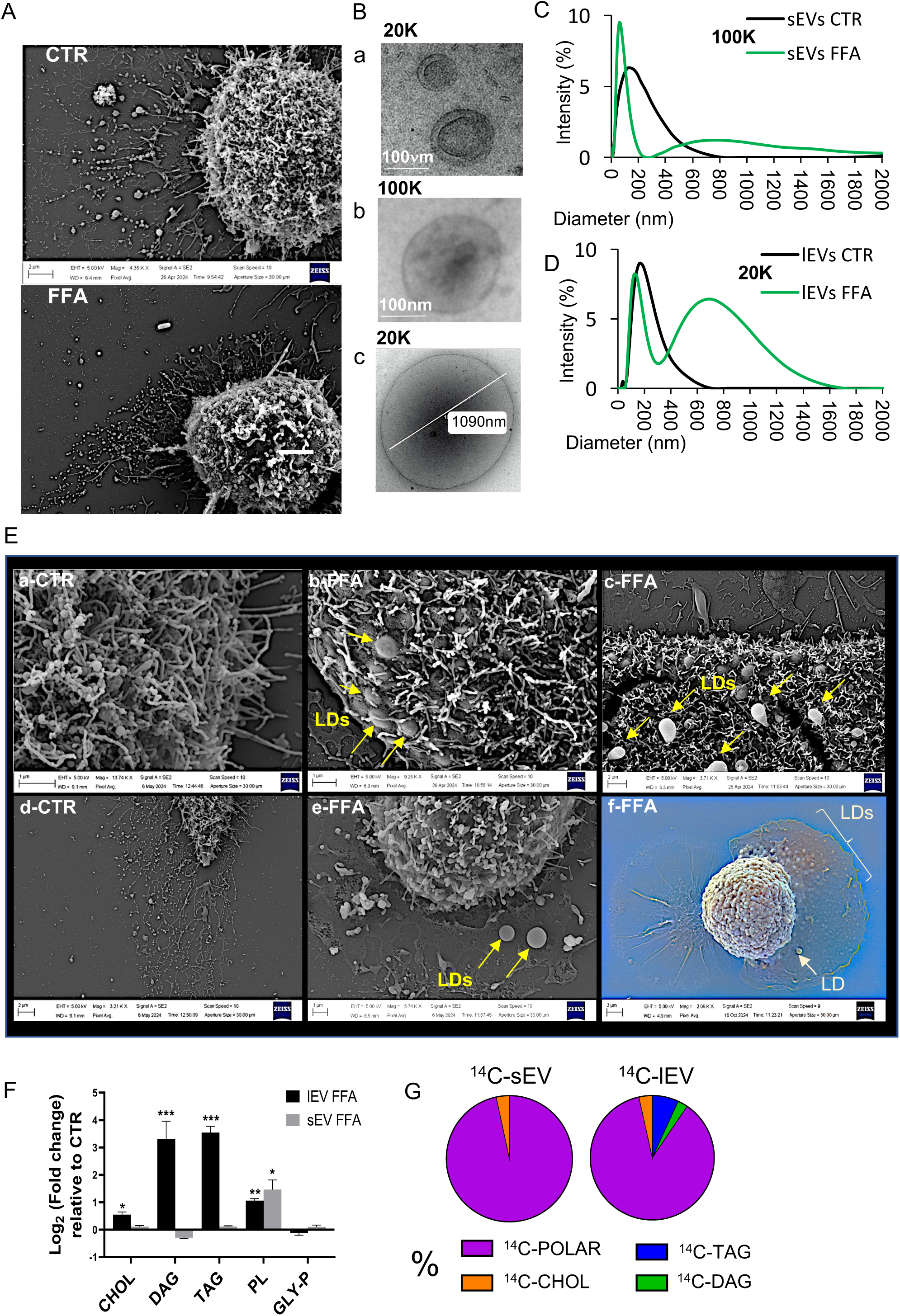
FFA-overloaded LAM/TREM2^+^ macrophages released extracellular vesicles containing TAG. **A**- SEM images showing that the diversity of the extracellular vesicles released from THP-1 macrophages (up) is reduced after the FFA treatment (down). **B**- TEM images showing EVs purified by differential centrifugations of THP-1 conditioned medium. **C**-**D** Extracellular vesicle size distribution from the 20k or 100k pellet, respectively, determined by dynamic light scattering. **E**- Lipid species from lEV-FFA and sEV-FFA detected by ^1^H-NMR spectroscopy analyses. Data are expressed as Log_2_ fold changes of lEV-CTR and sEV-CTR, respectively. F-TLC was used to quantify the radioactivity in each lipid species from ^14^C-sEVs and ^14^C-lEVs. Data are expressed as percentage of the total radioactivity. **G**- SEM images showing the release of LDs from FFA-treated macrophages (**b,c,e**) *vs* CTR (**a, d**). yellow arrays: LDs. (**f**) EVs and LDs released from a bone marrow (BM)-derived macrophages treated with FFA.

**Figure 3:**
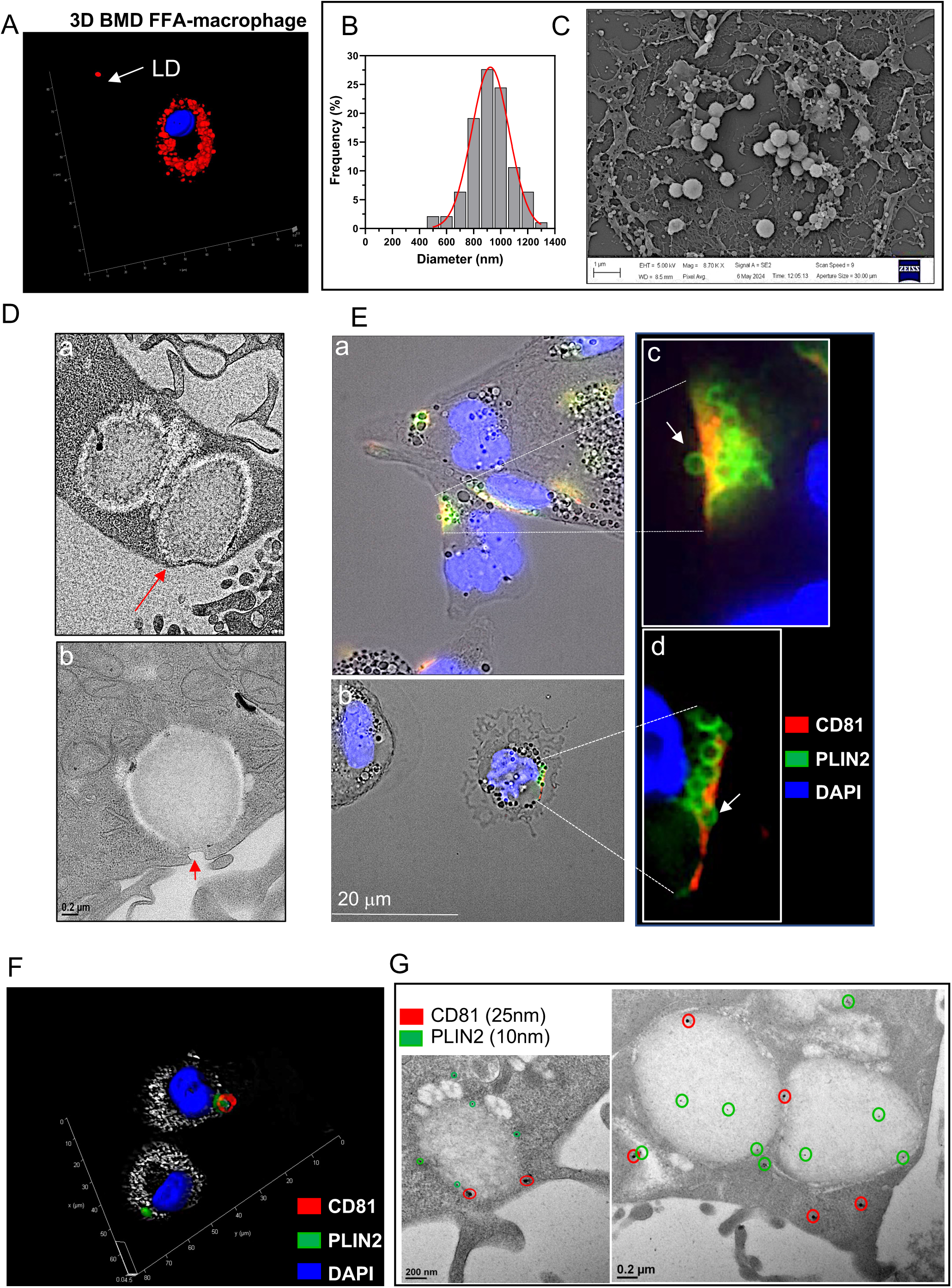
FFA-overloaded LAM/TREM2^+^ macrophages released extracellular lipid droplets. **A**- 3-D reconstruction from Z-stack images of BM-derived macrophages treated with FFA for 24h (x63), processed with the Leica LAS X software 5.2.2. Neutral lipids are visualized (red dot) after staining with Oil red O. Array = extracellular lipid droplet. **B**- Size distribution of the LDs observed in the vicinity of FFA-macrophages. **C**- Representative image used for **B**. Values are means ± SD (n=3); *p* values are from unpaired student *t*-test. (*) *p*<0.05, (**), *p*<0.01, (***) *p*<0.001. **D**- TEM images showing (a) LDs of FFA-macrophages pushing the plasma membrane (PM) (arrows) or (b) close to PM rupture. **E** (**c, d**) Co-labelling of CD81 and PLIN2 in non-permeabilized THP1-macrophages. Arrows indicate the release of LDs from cells (x63). **F-** 3D reconstruction from Z-stack images of FFA-macrophages co-labelled with CD81 and PLIN2, processed with the Leica LAS X software 5.2.2. showing a corona of CD81 around the site of LD release. **G**- TEM images of FFA-macrophages co-labelled with CD81 and PLIN2 (immunogold labelling).

### LAM/TREM2^+^ macrophage-EVs affect skeletal muscle homeostasis and modulate macrophage phenotype

To determine whether LAM/TREM2^+^ macrophage EVs were functional, M0 macrophages and C2C12 skeletal muscle cells were treated for 24h with lEV-FFA and sEV-FFA and their respective controls. LEV-FFA and sEV-FFA induced the expression of the cell surface marker of M2 polarization CD206 (Fig. 4A), but not CD86 (Fig. S4B). lEV-FFA and sEV-FFA also induced the release of IL-1β (Fig. 4B) compared to lEV-CTR and sEV-CTR. lEV-FFA and lEV-CTR, but none of the sEV populations, induced the expression of TREM2 and PLIN2 (Fig. 4C-D, Fig. S4C). However, TREM2 levels were significantly higher in response to lEVs-FFA *vs* lEVs-CTR (Fig. 4C). Taken together, these data indicated that lEVs-FFA polarized M0 macrophages into LAM/TREM2^+^ compared to sEVs-FFA. Interestingly, PLIN2 was found to be exported into macrophage-derived EVs in response to FFA (Fig. 4E). This data suggested that in addition to the lipids they contain, macrophage EVs might also transfer proteins involved in LD formation to recipient macrophages via the EV route in response to FFA.

**Figure 4:**
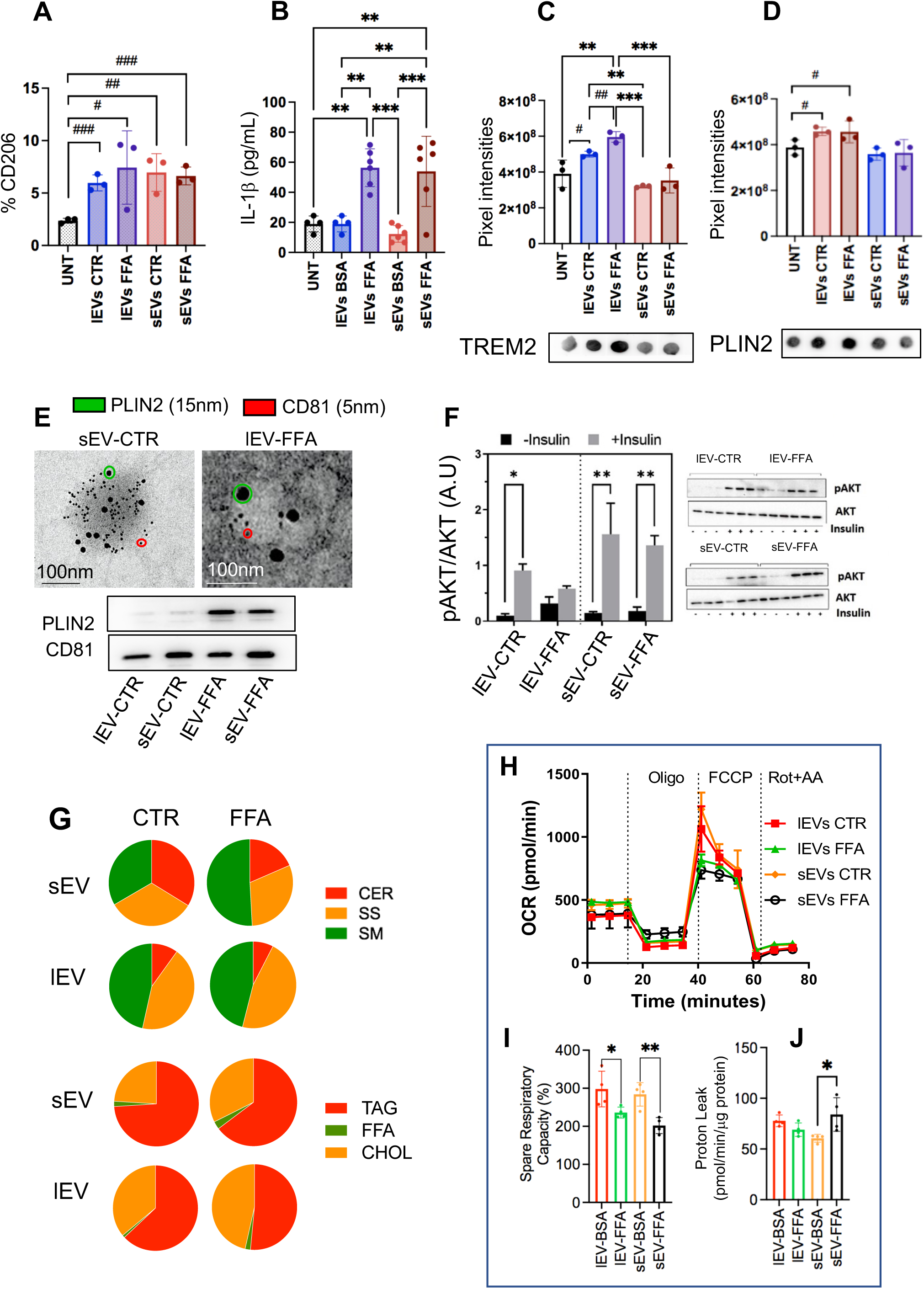
LAM/TREM2^+^ macrophage-released extracellular vesicles modulated macrophage phenotype and Skeletal Muscle metabolism. **A**- Percentage of macrophages expressing CD206 at the PM after treatment with EVs determined by FACS (the gating is provided in S4A). **B**- Quantification of IL-1β by ELISA assay in the supernatant of macrophages treated with EVs. **C-D-** Quantification of TREM2 (C) and PLIN2 (D) from dot-blot analyses. The quantification was performed by densitometry and is expressed as pixel intensity. Below representative blot used to generate the histograms (uncropped membranes are provided in S4C). (*) *p*<0.05, (**), *p*<0.01, (***), p<0.001 when *p* values are from one-way ANOVA and (#) *p*<0.05, (##), *p*<0.01, (##), *p*<0.001 when *p* values are from unpaired student *t*-test. **E**- Co-Immunogold-labelling of CD81 and PLIN2 at the surface of macrophage-EVs, visualized by TEM. Below, WB of EV protein extracts (10μg) to detect CD81 and PLIN2. **F**-Western blot quantification of [pAKT_Ser473_/AKT] ratios before and after insulin stimulation in recipient C2C12 in response to macrophage EVs. **G**-Lipid profiles of C2C12 muscle cells in response to macrophage EVs, analysed by TLC. Data are percentages of the total lipids (CER= ceramides, SS= sphingosine, SM: sphingomyelin, TAG= tryacylglycerols, FFA= free fatty acids, CHOL= cholesterol). **H-** Oxygen consumption rate (OCR) of C2C12 in response to macrophage EVs (Oligo = Oligomycin; Rot = Rotenone; AA = Antimycin A). **I**- Spare respiratory capacity (expressed as % of untreated C2C12). **J**- Detection of mitochondrial proton leak (difference in oxygen consumption before and after adding Oligomycin). Measurements were normalized by the protein concentration.

We then quantified the insulin-induced AKT phosphorylation in response to insulin in C2C12 as an indicator of muscle insulin-sensitivity. LEV-FFA exposure for 24h reduced p-AKT/AKT, *vs* lEV-CTR (Fig. 4F) and consequently reduced insulin-induced GLUT4 translocation at the plasma membrane in C2C12 (Fig. S5A). Instead, sEVs did not affect insulin-sensitivity (Fig. 4F). Recipient C2C12 lipid profiles showed that lEV-FFA induced cholesterol accumulation without affecting sphingomyelin levels (Fig. 4G, Fig S5B-C). Since the cholesterol/sphingomyelin ratio affects the PM fluidity, and consequently the activation of insulin receptors and GLUT4 membrane insertion (**20**), cholesterol enrichment could be one of the mechanisms to explain the altered insulin-sensitivity in myotubes in response to lEV-FFA. EVs from FFA-treated macrophages also affected the muscle cells bioenergetic efficiency (*i.e*.; decrease in OCR in response to FCCP Fig. 4H-I) *vs* CTR-EVs. Furthermore, sEV-FFA induced proton leak (Fig. 4I), reducing mitochondria efficiency to produce energy.

To explain these metabolic changes we performed RNAseq analyses of the recipient C2C12. A total of 824 and 790 genes were regulated in response to FFA-sEVs and FFA-lEVs, respectively (Fig. S5D). Interestingly, only 25 genes were in common demonstrating that FFA-sEVs and FFA-lEVs have distinct molecular signatures (Fig. S5E, Fig. 5A). To identify their unique functional roles in recipient muscle cells we performed gene ontology enrichment analyses on the regulated genes. FFA-lEVs regulated genes encoding extracellular matrix (ECM) proteins or proteins located in the nucleus (Fig.5B) whereas FFA-sEVs regulated genes significantly encoding proteins located in the mitochondria (Fig. 5C). The 824 and 790 regulated genes were then crossed with the 117 metabolic genes from the MetaCyc database to identify significantly enriched gene networks involved in metabolic regulations. No gene from the list of the genes modulated by FFA-lEVs was in common. Conversely 27 were included in the list of the genes down-regulated by sEV-FFA (Fig. 5D). They were included in 2 protein networks involved in fatty acid beta-oxidation and mitochondria respiratory chain complex. These data indicated that sEV-FFA-induced mitochondria dysfunction resulted from alterations at the transcriptional level. Similarly, we crossed the list of genes modulated by FFA-lEVs with the list of genes from the matrisome database (**21**) to identify significant components of ECM. There were 41 genes in common (Fig. 5E) that encoded for components of the extracellular matrix (including 9 collagens), significantly involved in muscle cell physiology and bone and joint formation (Fig. 5E).

**Figure 5:**
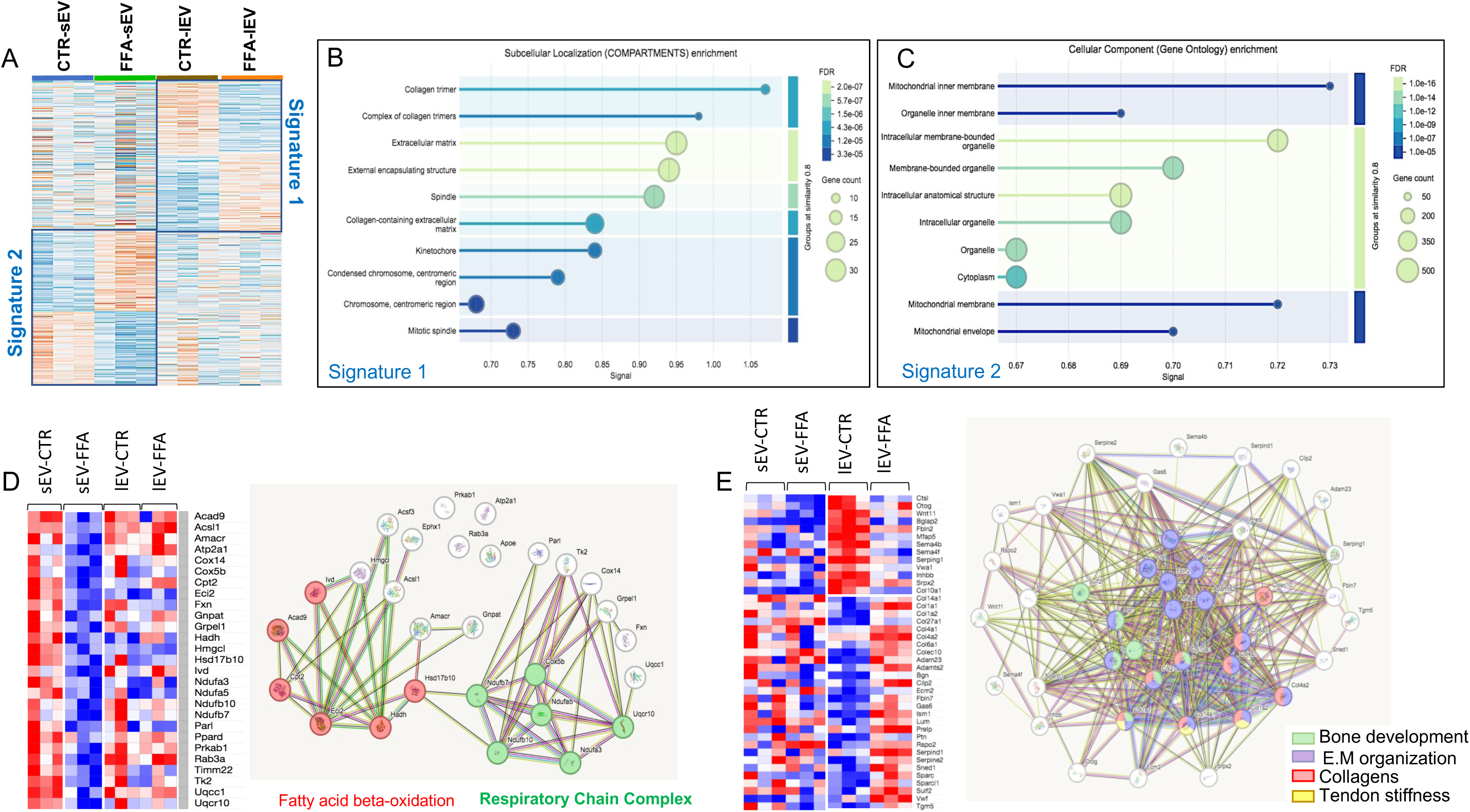
Cellular localization and Functional analyses of the genes regulated by macrophage EVs in recipient Skeletal muscle cells. **A**- Heatmap of the 1614 genes differentially regulated in response to macrophage EVs in recipient C2C12. **B-C** Cellular compartments significantly enriched in genes from signature 1 (**B**) or signature 2 (**C**). **D**- Heatmap and Functional analyses of the metabolic genes (MetaCyc Database, https://metacyc.org/) differentially regulated included in signatures 1 and 2. **E**- Heatmap and Functional analyses of the genes differentially regulated in response to macrophage EVs from signatures 1 and 2 contained in the matrisome database (https://matrisomedb.org/). All these analyses were performed with STRING 12.0 (https://string-db.org/).

## DISCUSSION

Recently, TREM2^+^-macrophages have emerged as key players in metabolic syndrome-related obesity, fatty liver, atherosclerosis and in tumors (22). Induced expression of the lipid sensor TREM2 is associated with phagocytic clearance of cellular debris, macrophage polarization and survival (23). Until now, the nutritional environment in the production and composition of EV-derived macrophages has not been extensively studied. Recently, we showed that under high glucose conditions, EVs released from M1 macrophage induced M2 polarization in recipient M0 macrophages (11). Here, we show that LAM/TREM2^+^ macrophage-derived EVs induced TREM2^+^ and Il-10 expression in M0 macrophages. This finding suggests that macrophage EVs play a critical role in mediating the metabolic reprogramming of macrophages, providing potential insights into how dietary factors and metabolic perturbations can shape immune cell behaviour in the context of metabolic diseases such as diabetes and obesity. An important finding of this study is that we show that FFA overload not only induces lipid storage in LAM/TREM2^+^ macrophages, but also alters their morphology and reduces the diversity and the number of the lipid-derived structures they release (*e.g*.; retractosomes, extracellular vesicles). This result suggests that LAM/TREM2^+^ macrophages reduce the amount of lipids they release into their environment, in line with their protective role in body fat accumulation at the early stage of obesity (**23**). However, analyses of energy production showed that FFA-overloaded LAM/TREM2^+^ macrophages are unable to utilize this excess of lipids and, surprisingly, we discover that they export and disseminate full LDs into their environment. The presence of TAG within EVs is not new, but has been the subject of much debate because the presence of contaminating lipoproteins in the EV preparations could not be ruled out, especially for EVs isolated from explants. In addition, TAG-containing EVs were usually < 300nm, probably due to the system used to measure EV size distribution in these studies, *i.e*.; the nanotracking analysis system (NTA) has a cut-off < 400nm (*e.g*. TAG-filled EVs from adipose tissues (**24**) or TAG-filled EVs isolated from hepathocytes (**25**)); or by the purification method (*i.e*.; size fractionation columns and filtration (**24**)). By using the dynamic light scattering method we had access to the full-size distribution of the LAM/TREM2^+^ macrophage EVs and were able to detect full LDs in the preparation of lEVs. This finding was confirmed by microscopy and PLIN2 labelling. Previously, SEM images showed that aggressive melanoma cells also release large LDs (called lipidosomes) that are taken up by neighbouring cells (**26**). However, lipidosomes cannot be compared to LDs released by LAM/TREM2^+^ macrophages because they are larger (2 to 6 µm) and also contain mitochondrial components. Therefore, our study introduces a new mechanism of TAG export from cells, distinct from the export within very low density lipoproteins (VLDL) released from the liver or within chylomicrons released from the intestine. This new mode of TAG export shows similarities to the release of milk fat globules by mammary epithelial cells, the only cells identified so far as releasing full LDs (**27, 28**). It cannot be excluded that other cell types capable of generating LDs in response to lipid overload, such as hepatocytes, pancreatic beta cells or enterocytes also export LDs. At present, we don’t know whether LAM/TREM2+ macrophage-derived LDs are destined to be exported after their synthesis and thus might represent a new form of local TAG exchange between cells, or whether this export is a form of rescue to protect LAM/TREM2+ macrophages from an excess of intracellular lipids that it can no longer handle. At the present time, we favour the first hypothesis for several reasons, *i.e*.; macrophage LDs accumulate mainly under the plasma membrane at the periphery of the cell, as the LDs from mammary epithelial cells; LDs are never in contact with mitochondria; the release of LDs occurs in specific regions on the surface of the macrophage, associated with the presence of the tetraspanin CD81, suggesting a specific mechanism linking LD synthesis and release. Interestingly, we found that CD81 is barely detected on cell surface, even in control macrophages, and is usually localized close to or associated with LDs or the endoplasmic reticulum, the site of LD formation. PLIN2 have been shown to interact with CD81 (**29**) positioning CD81 at the interface between lipid droplets and the ER. These interactions may play a role in regulating the dynamics of lipid droplet formation and metabolism, potentially linking CD81 to lipid homeostasis and cellular responses to metabolic stress. Consistently, CD81 is involved in the control of body fat (**30**), has a cholesterol-binding pocket that influences its conformation and activity (31), and can stimulate phosphatidylinositol-4 kinase activity at the plasma membrane (**32**), which is involved in vesicular trafficking and lipid transport (**33, 34**). Further studies are now needed to understand the role of CD81 in the release of LDs from macrophages.

The second important finding of this study is that LAM/TREM2^+^ macrophage-derived EVs modulate insulin-sensitivity (lEV-FFA), mitochondrial oxidative capacity (sEV-FFA), lipid profiles (sEV-FFA, lEV-FFA) and gene expressions encoding extracellular matrix components (lEV-FFA) in recipient muscle cells. Of note, TREM2 protein is not detected in myocytes (**35**), so if TREM2 is detected in SkM *in vivo*, it is mainly due to the presence of TREM2^+^ immune cells, including macrophages (36). Interestingly, our analysis of all previous skeletal muscle (SkM) transcriptome data recorded in the Gene Expression Omnibus (GEO) repository failed to find a correlation between SkM TREM2 expression and obesity or diabetes, in contrast to adipose tissue (**10**). Instead, TREM2 is strongly increased when SkM mass is affected (*e.g*.; atrophy-associated DMD (**37**), or in the presence of age-related atrophy (**36**), suggesting that TREM2 macrophages are important in skeletal muscle tissue remodeling and regeneration. Recently, it was described that TREM2 knock-out mice, with the TREM2-R47H variant, had reduced bone mass but induced SkM strength associated with fiber type-specific changes. The authors of this study suggested that it was unlikely that cytokines involved in bone–muscle crosstalk could explain these SkM phenotypic changes and postulated that the absence of TREM2-expressing macrophages in these animals could be at the origin of this phenotype (**38**). Although generated in a specific context of lipid overload, our data support a role for TREM2^+^ macrophages in the regulation of SkM homeostasis and highlight the TREM2^+^ macrophage-derived EVs as important actors in this process. In particular, they demonstrate the ability of TREM2^+^ macrophage-derived EVs to induce extracellular matrix (ECM) remodelling and collagen expression from muscle cells.

This study is the first to demonstrate that macrophage phenotypic plasticity induced by changes in microenvironmental FFA concentrations can consequently modify the biological activity of their derived EVs on muscle cells. Thus, macrophage-derived EVs represent a novel mechanism, independent of the cytokine pathway that explains how macrophages ‘talk to muscle’ and new therapeutic targets to prevent deleterious changes in muscle mass.

---

**Supplementary figure S1:**

Uncropped Western-blot used in this study.

**Supplementary Figure S2:**

**A**- Lipid profiles determined by TLC analyses of THP-1 macrophages treated with palmitate+oleate (FFA) supplemented with ^14^C-palmitate. **B**- Size of the lipid droplets contained in THP-1 macrophages treated for 24h with 500μM palmitate+oleate (1:2) determined manually from TEM images (below). **C**- Percentage of THP-1 cell viability determined with a MTT test. **F,G**- Quantification of IL-10 (**F**) and IL-1β (**G**) cytokine in the conditioned medium of THP-1 macrophages treated with 500μM palmitate+oleate (1:2) *vs* BSA (CTR). Polarized-THP-1 macrophages into M1 or M2 were used as positive control of cytokine production. (*) *p*<0.05, (**), *p*<0.01, (***), p<0.001; *p* values are from one-way ANOVA

**Supplementary Figure S3:**

**A**- CryoEM images of EV released from macrophages in the 20k and 100K pellet. **B**- Original TEM image used to build Fig. 1A-c. (**B**) and (**C**) show the formation of lipid droplets from the fusion of several small lipid droplets in THP-1 macrophages treated with FFA. **C**-Images showing the co-localization of lipid droplet release sites with CD81 that the plasma membrane. **D-E**- TEM images showing the localisation of CD81 in FFA- and CTR- THP-1 macrophages. CD81 gold particles = 25 nm. E.R = endoplasmic reticulum. LD = lipid droplet.

**Supplementary Figure S4:**

**A**- Gating strategy used for FACS analyses used to determine the percentage of CD206 (Fig. 4A) and CD86 (**B**) at the surface of THP-1 macrophages treated with EVs. **C**- Original Dot-blot showing the expression of TREM2 and PLIN2, used to draw the histograms Fig. 4C-D.

**Supplementary Figure 5:**

**A**- Immunodetection of GLUT4 and labelling with Phalloidin (F-actin) and DAPI (nucleus) in non-permeabilized C2C12 treated with macrophage-derived EVs, before (Basal) or after (+Insulin) 15min incubation with Insulin. Bar= 25μm. **B**-**C** Lipid profiles of C2C12 determined by TLC, in response to macrophage-derived EVs, used in Fig. 4G. TAG= Triacylglycerols; FFA= Free fatty acids; CHOL= Cholesterol; CER= ceramides; SS= sphingosine; SM= sphingomyelin. **D-** Ven Graph showing the gene number overlap between the list of genes regulated by small EV-FFA and the list of genes regulated by large EV-FFA, *vs* respective controls in C2C12 (lists in Table S2). **E**- Principal component analysis (PCA) based on the differentially regulated genes in C2C12 in response to macrophage EV-FFA and large EV-FFA, *vs* respective controls.

**Supplementary Figure 6:**

**A-** Heatmap of the differentially regulated genes in C2C12 in response to macrophage EV-FFA and large EV-FFA, *vs* respective controls. **B-C** Analyses of the genes included in signature 1 and 2, respectively by STRING 12.0 to determine their significant cellular compartments. **E**- Heatmap of the genes contained in the metabolic gene database and included in the list of genes regulated by macrophage EV-FFA or large EV-FFA. Right, significantly enriched functions related to protein networks retrieved from STRING 12.0. **F**- Heatmap of the genes contained in the matrisome database and included in the list of genes regulated by macrophage EV-FFA or large EV-FFA. Right, significantly enriched functions related to protein networks retrieved from STRING 12.0.

## Author Contributions

S.T., A.M.G., S.L.= fluorescence microscopy, cell culture, experiments with ^14^C; E.E-C, C.C.= sample preparations for TEM; F.B., A. E-J= FACS analyses, ELISA for cytokine quantification, bone marrow-derived human macrophages; C.C.D.S., C.B.= help for macrophage culture; M. Z., A.A.=EV preparations for cryoelectron microscopy and pictures; F.A., F.P.F.= H^1^NMR spectroscopy analysis; S.P.; Real-Time PCR and primer design; F.A., F.P.F.=H1NMR spectroscopy analysis; L.C.= help for Seahorse analysis and design of the experiment; B.G., S.H.= RNAseq; E.M.; data curation and RNAseq normalization; J.R., H.V.= data interpretation and helpfull discussion; S.T.= was involved in all the experiments of the study and drafted the manuscript; L.D.= study design, correction of the manuscript; S.R. = TEM and SEM pictures, bioinformatic analyses, supervision, study design, writing and editing. All authors have read, corrected the manuscript and agree with the submitted version. ST is the guarantors of this work and, as such, has full access to the data produced in Italy and France, and takes responsibility for the integrity of the data

Sample preparation for SEM and images were realized at the Centre Technologique des Microstructures (CTµ) from the Université Claude Bernard Lyon1, Lyon, FRANCE. We thank Armelle Penhoat for for showing us how to use the CarMeN laboratory’s DLS.

## Data Availability Statement

All data from this article are available upon request

## Conflicts of Interest

The authors declare no conflict of interest.

## Funding

This work is supported by the FRENCH NATIONAL FUNDING AGENCY (# ANR-21-CE14-0081; # ANR-20-CE18-0026).

## Abbreviations

CD86: cluster of differentiation 86
CD206: cluster of differentiation 206
EVs: extracellular vesicles
IL-8: Interleukin 8
IFNα: Interferon alpha
CM: conditioned medium
IL-1β: Interleukin-1 beta
IL-10: Interleukin-10
GAPDH: Glyceraldehyde-3-Phosphate Dehydrogenase
CD36: Fatty Acid Transporter
TEM: Transmission Electron Microscopy
SEM: Scanning Electron Microscopy
DAPI: 4′,6-diamidino-2-phenylindole
LDs: lipid droplets
LAM: lipid-associated macrophages
TREM2: Triggering receptor expressed on myeloid cells 2
FABP4: Fatty Acid Binding Protein 4
FAT/CD36: Fatty Acid Translocase
PLIN2: Perilipin 2
ABCA1: ATP Binding Cassette Subfamily A Member 1
RPMI: Roswell Park Memorial Institute medium
DMEM: Dulbecco’s Modified Eagle Medium
BSA: Bovine serum albumin
AKT: Protein Kinase B or PKB
GLUT4: Solute Carrier Family 2 Member 4
SEM: scanning electron microscopy
TEM: transmission electron microscopy
CryoEM: cryoelectronic microscopy
FACS: Fluorescence-Activated Cell Sorting
OCR: Oxygen consumption rate
DGAT1/2: Diacylglycerol O-acyltransferase 1 or 2
CD81: Tetraspanin-28
CD9: Tetraspanin-29
CD63: Lysosomal-Associated Membrane Protein 3
IL-10: Interleukin 10
IL-1β: Interleukin 1 beta
PM: Plasma membrane
TLC: Thin-layer chromatography
NMR: Nuclear Magnetic Resonance Spectroscopy
DAG: Diacylglycerols
TAG: Triacylglycerols
FFA: Free fatty acids
CHOL: cholesterol
CER: ceramides
SS: sphingosines
SM: sphingomyelins
SkM: Skeletal Muscle
sEV: small extracellular vesicles
lEV: large extracellular vesicles
MVB: Multivesicular bodies
DMSO: Dimethyl sulfoxide
MTT: 3-(4,5-Dimethylthiazol-2-yl)-2,5-Diphenyltetrazolium Bromide
PCA: Principal Component Analysis
BMD macrophages: Bone marrow-derived

## Notes

### Competing Interest Statement

The authors have declared no competing interest.

