## Supplementary figures and images for "LAM/TREM2 ^+^ macrophages release extracellular vesicles and extracellular lipid droplets which modulate the phenotype of recipient macrophages and homeostasis of skeletal muscle cells"

### SUPPLEMENTARY FIGURES

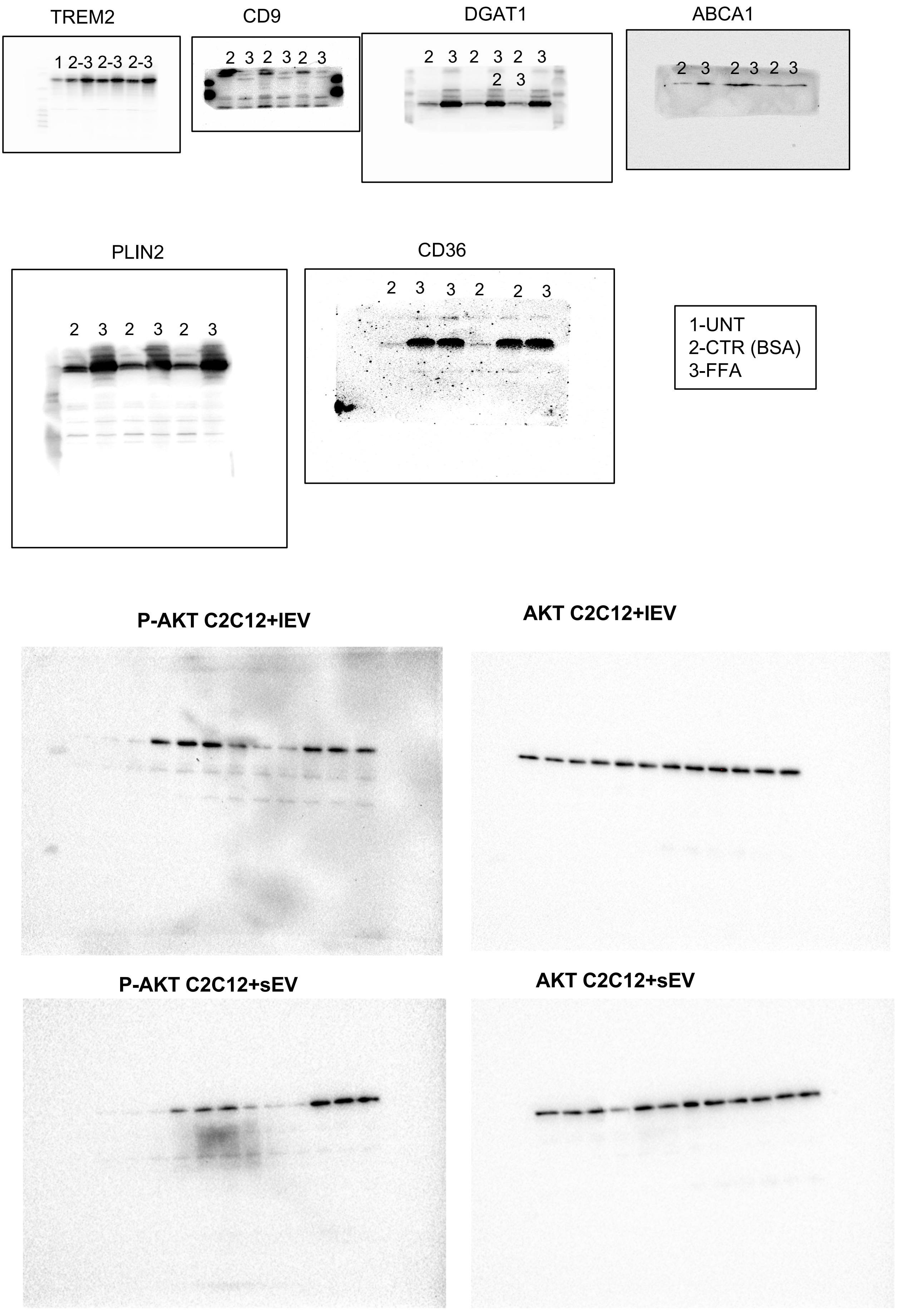

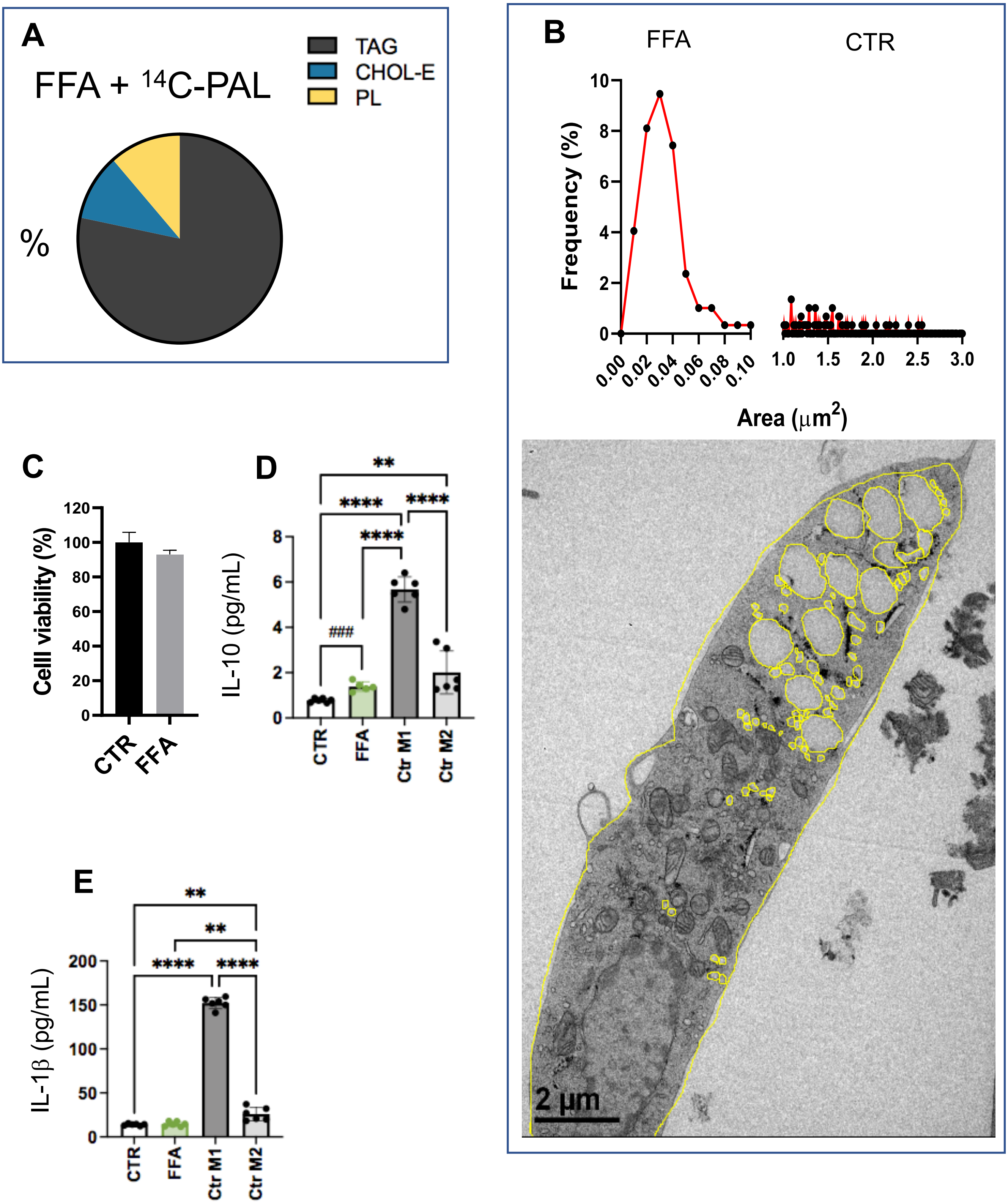

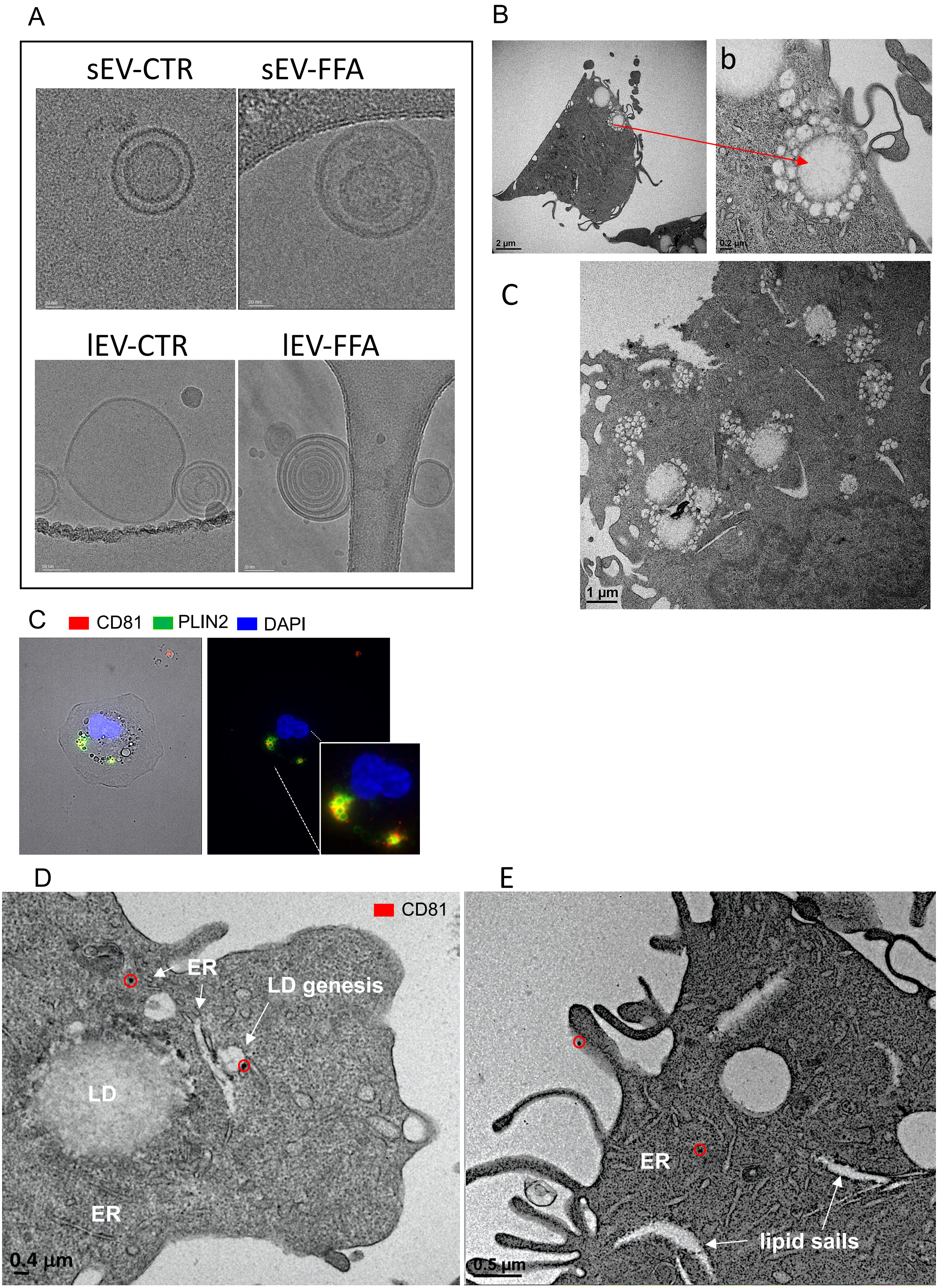

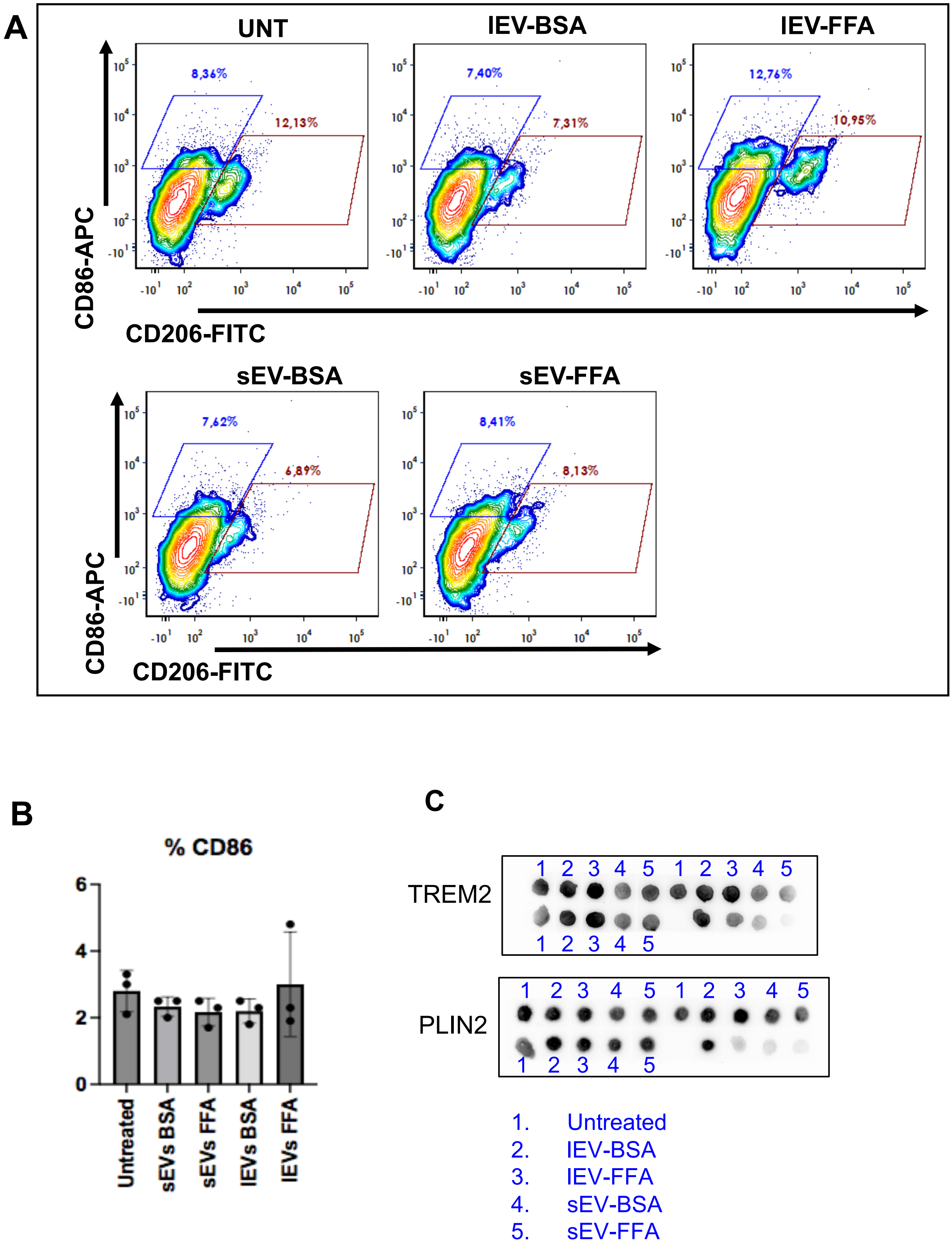

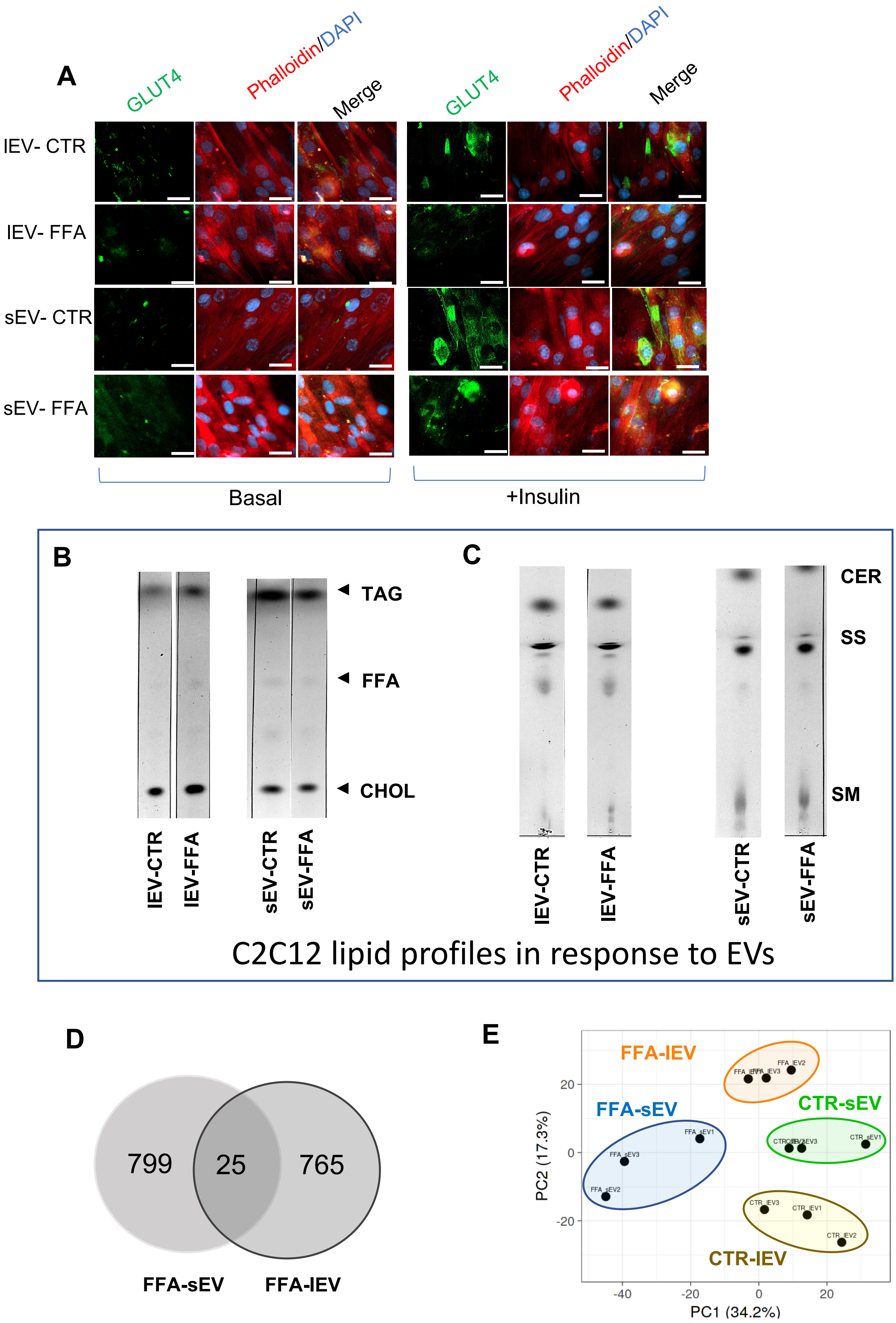
