## Supplementary material for "LAM/TREM2 ^+^ macrophages release extracellular vesicles and extracellular lipid droplets which modulate the phenotype of recipient macrophages and homeostasis of skeletal muscle cells": TABLE S1

**TABLE S1**: antibodies and primers used in this study

A-ANTIBODIES

| Protein Names | Symbol | Suppliers | REFERENCES | DILUTIONS |
|---|---|---|---|---|
| Triggering receptor expressed on myeloid cells 2 | TREM2 | Cell Signaling Technology | 55739S | 1/1000 |
| Perilipin-2 | PLIN2/ADRP | Proteintech | 15294-1-AP | 1/1000 |
| Cluster of differentiation 36 (fatty acid translocase) | CD36 | Abcam | ab252922 | 1/1000 |
| Arginase 1 | CD9 | Santa Cruz Biotechnology | sc-131118 | 1/500 |
| Fatty acid binding protein 4 | FABP4 | Cell Signaling Technology | 3544 | 1/500 |
| Diacylglycerol acyltransferase-1 | DGAT1 | Sigma Aldrich | SAB4301075 | 1/1000 |
| Phosphorylated protein kinase B | pAKT | Abcam | ab8932 | 1/1000 |
| Protein kinase B | AKt | Abcam | ab8805 | 1/1000 |
| Solute Carrier Family 2 Member 4 | GLUT4 | Abcam | ab33780 | 1/400 |
| Cluster of differentiation 63 | CD63 | Abcam | ab216130 | 1/1000 |
| Cluster of differentiation 81 | CD81 | Santa Cruz Biotechnology | sc-166028 | 1/1000 |
| Cluster of Differentiation 86 | CD86 | Miltenyi | 130-116-161 | 1/50 |
| Cluster of Differentiation 206 | CD206 | Miltenyi | 130-123-671 | 1/50 |
| APC anti-human IgG1 | APC anti-human IgG1 | Miltenyi | 130-113-446 | 1/50 |
| FITC anti-human IgG1 | FITC anti-human IgG1 | Miltenyi | 130-113-444 | 1/50 |
| Anti-rabbit IgG, HRP-linked | | Cell Signaling Technology | 7074 | 1/10000 |
| Anti-mouse IgG, HRP-linked | | Cell Signaling Technology | 7076 | 1/10000 |

B- PRIMERS

| Gene Symbols | Forward | Reverse |
|---|---|---|
| ABCA1 | 5'-CAGGAGGTGATGTTTCTGACCA-3' | 5'-TTGGCTGTTCTCCATGAAGGTC-3' |
| DGAT2 | 5'-AAGGGCTTTGTGAAACTGGC-3' | 5'-CCTCCTCGAAGATCACCTGC-3' |
