## Supplementary material for "LAM/TREM2 ^+^ macrophages release extracellular vesicles and extracellular lipid droplets which modulate the phenotype of recipient macrophages and homeostasis of skeletal muscle cells": TABLE S2

**TABLE S2**: Normalized gene counts. Only significantly regulated genes in C2C12 in response to macrophage EVs are listed
Data are available at GEO #GSE199222
Genes are listed in the same order they are represented in the Heatmap of Figure 5

| Treatments | CTR_sEV1 | CTR_sEV2 | CTR_sEV3 | FFA_sEV1 | FFA_sEV2 | FFA_sEV3 | CTR_lEV1 | CTR_lEV2 | CTR_lEV3 | FFA_lEV1 | FFA_lEV2 | FFA_lEV3 |
|---|---|---|---|---|---|---|---|---|---|---|---|---|
| ENSMUSG00000060572 | 136.51 | 117.81 | 121.52 | 99.24 | 96.12 | 90.81 | 103.41 | 97.90 | 78.53 | 130.52 | 114.05 | 117.74 |
| ENSMUSG00000070960 | 135.53 | 113.88 | 130.34 | 88.90 | 97.20 | 106.47 | 94.42 | 102.01 | 98.16 | 108.40 | 112.03 | 121.50 |
| ENSMUSG00000053675 | 169.66 | 141.37 | 157.78 | 128.18 | 115.56 | 134.66 | 131.29 | 116.00 | 121.72 | 134.95 | 154.42 | 145.05 |
| ENSMUSG00000019326 | 276.92 | 255.25 | 282.25 | 225.35 | 232.21 | 238.00 | 169.06 | 158.78 | 167.85 | 214.59 | 255.35 | 227.93 |
| ENSMUSG00000025105 | 10.73 | 14.73 | 12.74 | 16.54 | 18.36 | 15.66 | 17.09 | 16.45 | 19.63 | 12.17 | 7.06 | 11.30 |
| ENSMUSG00000070315 | 40.95 | 37.31 | 41.16 | 52.72 | 54.00 | 53.24 | 51.26 | 49.36 | 45.15 | 40.93 | 30.28 | 38.62 |
| ENSMUSG00000097061 | 55.58 | 53.01 | 61.74 | 77.53 | 82.08 | 82.46 | 62.95 | 60.88 | 53.99 | 51.99 | 50.46 | 45.21 |
| ENSMUSG00000083614 | 6.83 | 8.84 | 5.88 | 14.47 | 16.20 | 10.44 | 20.68 | 17.28 | 17.67 | 13.27 | 12.11 | 16.01 |
| ENSMUSG00000019763 | 600.63 | 625.37 | 636.03 | 714.29 | 668.54 | 686.85 | 633.07 | 603.85 | 605.63 | 660.35 | 635.85 | 660.26 |
| ENSMUSG00000097993 | 75.08 | 87.38 | 82.32 | 102.34 | 91.80 | 99.16 | 81.83 | 84.74 | 86.38 | 109.50 | 106.98 | 113.97 |
| ENSMUSG00000029648 | 22.43 | 25.53 | 25.48 | 41.35 | 33.48 | 37.58 | 18.88 | 18.92 | 25.52 | 35.40 | 40.37 | 29.20 |
| ENSMUSG00000053714 | 4.88 | 5.89 | 3.92 | 11.37 | 10.80 | 7.31 | 6.29 | 5.76 | 3.93 | 11.06 | 11.10 | 9.42 |
| ENSMUSG00000103984 | 7.80 | 9.82 | 7.84 | 18.61 | 28.08 | 19.83 | 11.69 | 12.34 | 11.78 | 13.27 | 14.13 | 13.19 |
| ENSMUSG00000086578 | 4.88 | 5.89 | 5.88 | 3.10 | 1.08 | 1.04 | 4.50 | 4.11 | 5.89 | 2.21 | 3.03 | 1.88 |
| ENSMUSG00000058932 | 8.78 | 7.85 | 5.88 | 4.13 | 3.24 | 3.13 | 8.09 | 10.69 | 10.80 | 5.53 | 6.06 | 4.71 |
| ENSMUSG00000048856 | 22.43 | 18.65 | 18.62 | 13.44 | 10.80 | 11.48 | 22.48 | 22.21 | 23.56 | 17.70 | 15.14 | 17.90 |
| ENSMUSG00000096981 | 20.48 | 21.60 | 16.66 | 11.37 | 12.96 | 14.61 | 15.29 | 16.45 | 16.69 | 14.38 | 12.11 | 14.13 |
| ENSMUSG00000015957 | 49.73 | 52.03 | 42.14 | 33.08 | 39.96 | 31.32 | 56.65 | 62.52 | 59.88 | 49.77 | 45.42 | 50.86 |
| ENSMUSG00000074886 | 1044.29 | 913.02 | 1022.16 | 867.27 | 754.94 | 778.70 | 979.28 | 1002.02 | 972.74 | 898.16 | 886.15 | 895.73 |
| ENSMUSG00000032118 | 100.43 | 91.30 | 105.84 | 85.80 | 73.44 | 86.64 | 142.98 | 141.50 | 128.59 | 109.50 | 112.03 | 87.59 |
| ENSMUSG00000107314 | 1583.49 | 1625.77 | 1533.73 | 1413.07 | 1486.12 | 1458.24 | 1586.27 | 1586.95 | 1490.03 | 1352.77 | 1422.08 | 1431.66 |
| ENSMUSG00000081999 | 2095.40 | 2100.93 | 2003.16 | 1905.11 | 1917.05 | 1873.69 | 2109.63 | 2209.72 | 2103.52 | 1927.95 | 1964.06 | 1957.23 |
| ENSMUSG00000032218 | 2006.67 | 2040.07 | 2062.94 | 1844.12 | 1905.17 | 1887.26 | 2132.11 | 2400.58 | 2222.29 | 1712.26 | 1866.16 | 1769.79 |
| ENSMUSG00000024640 | 2839.36 | 2810.74 | 2742.10 | 2569.78 | 2637.43 | 2719.20 | 2865.00 | 2880.20 | 2849.51 | 2725.46 | 2610.00 | 2706.96 |
| ENSMUSG00000086130 | 3.90 | 4.91 | 6.86 | 6.20 | 8.64 | 7.31 | 6.29 | 4.94 | 3.93 | 2.21 | 1.01 | 0.94 |
| ENSMUSG00000025658 | 6.83 | 6.87 | 5.88 | 4.13 | 6.48 | 2.09 | 8.99 | 9.87 | 5.89 | 3.32 | 1.01 | 3.77 |
| ENSMUSG00000037579 | 1.95 | 2.95 | 2.94 | 9.30 | 1.08 | 1.04 | 6.29 | 4.94 | 6.87 | 1.11 | 4.04 | 0.94 |
| ENSMUSG00000028972 | 9.75 | 6.87 | 12.74 | 6.20 | 9.72 | 7.31 | 33.27 | 23.04 | 31.41 | 9.95 | 11.10 | 10.36 |
| ENSMUSG00000028572 | 2.93 | 2.95 | 4.90 | 5.17 | 5.40 | 5.22 | 5.40 | 6.58 | 6.87 | 2.21 | 1.01 | 3.77 |

| | | | | | | | | | | | | |
|---|---|---|---|---|---|---|---|---|---|---|---|---|
| ENSMUSG00000107742 | 2.93 | 0.98 | 2.94 | 8.27 | 4.32 | 6.26 | 8.09 | 6.58 | 5.89 | 1.11 | 4.04 | 2.83 |
| ENSMUSG00000024459 | 4.88 | 6.87 | 6.86 | 7.24 | 2.16 | 8.35 | 6.29 | 8.23 | 8.83 | 1.11 | 5.05 | 3.77 |
| ENSMUSG00000103138 | 3.90 | 2.95 | 4.90 | 4.13 | 4.32 | 4.18 | 4.50 | 3.29 | 3.93 | 1.11 | 2.02 | 1.88 |
| ENSMUSG00000032940 | 1.95 | 0.98 | 0.98 | 2.07 | 4.32 | 2.09 | 2.70 | 2.47 | 1.96 | 1.11 | 1.01 | 0.94 |
| ENSMUSG00000020904 | 6.83 | 9.82 | 7.84 | 7.24 | 12.96 | 9.39 | 11.69 | 18.92 | 16.69 | 5.53 | 6.06 | 9.42 |
| ENSMUSG00000020787 | 3.90 | 3.93 | 0.98 | 5.17 | 1.08 | 1.04 | 6.29 | 7.40 | 6.87 | 3.32 | 3.03 | 2.83 |
| ENSMUSG00000029054 | 20.48 | 16.69 | 11.76 | 20.67 | 9.72 | 5.22 | 35.97 | 29.62 | 28.47 | 13.27 | 12.11 | 17.90 |
| ENSMUSG00000071392 | 1.95 | 5.89 | 3.92 | 4.13 | 3.24 | 3.13 | 5.40 | 7.40 | 6.87 | 2.21 | 4.04 | 2.83 |
| ENSMUSG00000030303 | 3.90 | 7.85 | 1.96 | 6.20 | 4.32 | 3.13 | 5.40 | 5.76 | 5.89 | 3.32 | 3.03 | 1.88 |
| ENSMUSG00000034387 | 12.68 | 17.67 | 10.78 | 9.30 | 12.96 | 12.53 | 22.48 | 27.15 | 19.63 | 11.06 | 12.11 | 10.36 |
| ENSMUSG00000097081 | 8.78 | 11.78 | 7.84 | 7.24 | 4.32 | 4.18 | 8.99 | 9.05 | 6.87 | 4.42 | 3.03 | 4.71 |
| ENSMUSG00000104186 | 13.65 | 14.73 | 9.80 | 7.24 | 16.20 | 8.35 | 13.49 | 13.99 | 17.67 | 6.64 | 8.07 | 7.54 |
| ENSMUSG00000108015 | 24.38 | 23.56 | 19.60 | 23.78 | 18.36 | 12.53 | 27.88 | 31.26 | 36.32 | 14.38 | 12.11 | 20.72 |
| ENSMUSG00000097468 | 21.45 | 15.71 | 15.68 | 15.51 | 16.20 | 11.48 | 16.19 | 13.99 | 14.72 | 9.95 | 6.06 | 6.59 |
| ENSMUSG00000025432 | 13.65 | 11.78 | 1.96 | 14.47 | 9.72 | 14.61 | 26.08 | 18.10 | 18.65 | 12.17 | 10.09 | 9.42 |
| ENSMUSG00000021091 | 35.10 | 22.58 | 19.60 | 27.91 | 27.00 | 12.53 | 44.06 | 46.07 | 49.08 | 23.23 | 28.26 | 18.84 |
| ENSMUSG00000055373 | 18.53 | 26.51 | 16.66 | 15.51 | 32.40 | 25.05 | 25.18 | 21.39 | 27.48 | 11.06 | 15.14 | 12.24 |
| ENSMUSG00000064036 | 8.78 | 3.93 | 4.90 | 5.17 | 2.16 | 1.04 | 4.50 | 3.29 | 3.93 | 2.21 | 2.02 | 1.88 |
| ENSMUSG00000103120 | 6.83 | 6.87 | 4.90 | 7.24 | 3.24 | 10.44 | 7.19 | 9.87 | 9.82 | 3.32 | 6.06 | 4.71 |
| ENSMUSG00000073073 | 4.88 | 1.96 | 2.94 | 2.07 | 8.64 | 3.13 | 3.60 | 4.11 | 3.93 | 2.21 | 2.02 | 1.88 |
| ENSMUSG00000047604 | 13.65 | 11.78 | 17.64 | 8.27 | 15.12 | 6.26 | 12.59 | 18.10 | 14.72 | 5.53 | 8.07 | 10.36 |
| ENSMUSG00000029053 | 8.78 | 9.82 | 7.84 | 10.34 | 4.32 | 11.48 | 14.39 | 12.34 | 9.82 | 6.64 | 8.07 | 4.71 |
| ENSMUSG00000039115 | 72.15 | 82.47 | 68.60 | 60.99 | 74.52 | 50.10 | 177.15 | 190.04 | 131.53 | 58.62 | 85.79 | 123.39 |
| ENSMUSG00000086873 | 7.80 | 7.85 | 8.82 | 10.34 | 12.96 | 16.70 | 8.99 | 11.52 | 9.82 | 5.53 | 6.06 | 4.71 |
| ENSMUSG00000106682 | 0.98 | 1.96 | 4.90 | 7.24 | 3.24 | 7.31 | 5.40 | 4.94 | 4.91 | 3.32 | 3.03 | 1.88 |
| ENSMUSG00000107868 | 17.55 | 17.67 | 18.62 | 23.78 | 17.28 | 24.01 | 23.38 | 19.74 | 27.48 | 12.17 | 10.09 | 16.01 |
| ENSMUSG00000039462 | 45.83 | 59.89 | 42.14 | 35.15 | 41.04 | 45.93 | 129.49 | 111.06 | 89.32 | 65.26 | 57.53 | 56.51 |
| ENSMUSG00000087578 | 6.83 | 10.80 | 11.76 | 6.20 | 8.64 | 8.35 | 11.69 | 10.69 | 13.74 | 5.53 | 7.06 | 7.54 |
| ENSMUSG00000041347 | 57.53 | 52.03 | 39.20 | 25.84 | 30.24 | 31.32 | 50.36 | 64.99 | 46.13 | 28.76 | 35.32 | 26.37 |
| ENSMUSG00000073234 | 13.65 | 11.78 | 12.74 | 7.24 | 19.44 | 14.61 | 17.09 | 18.92 | 12.76 | 8.85 | 11.10 | 7.54 |
| ENSMUSG00000027858 | 30.23 | 28.47 | 28.42 | 21.71 | 37.80 | 31.32 | 33.27 | 27.15 | 28.47 | 13.27 | 17.16 | 19.78 |
| ENSMUSG00000032492 | 83.85 | 86.39 | 68.60 | 97.17 | 59.40 | 65.76 | 163.66 | 157.13 | 124.66 | 88.49 | 81.75 | 91.36 |
| ENSMUSG00000073177 | 26.33 | 30.43 | 28.42 | 27.91 | 30.24 | 25.05 | 29.68 | 32.08 | 25.52 | 16.59 | 15.14 | 19.78 |
| ENSMUSG00000040751 | 30.23 | 44.18 | 30.38 | 34.11 | 32.40 | 30.27 | 53.06 | 57.59 | 53.99 | 30.97 | 32.30 | 34.85 |

| | | | | | | | | | | | | |
|---|---|---|---|---|---|---|---|---|---|---|---|---|
| ENSMUSG00000000627 | 24.38 | 40.25 | 25.48 | 37.21 | 30.24 | 14.61 | 32.37 | 37.84 | 30.43 | 24.33 | 22.20 | 14.13 |
| ENSMUSG00000085095 | 24.38 | 22.58 | 25.48 | 31.01 | 15.12 | 18.79 | 37.77 | 34.55 | 40.24 | 24.33 | 16.15 | 28.26 |
| ENSMUSG00000083414 | 2.93 | 7.85 | 3.92 | 5.17 | 7.56 | 3.13 | 6.29 | 8.23 | 6.87 | 4.42 | 3.03 | 5.65 |
| ENSMUSG00000085257 | 2.93 | 6.87 | 7.84 | 3.10 | 9.72 | 7.31 | 6.29 | 7.40 | 7.85 | 4.42 | 5.05 | 3.77 |
| ENSMUSG00000081169 | 37.05 | 38.29 | 49.98 | 45.48 | 39.96 | 20.88 | 51.26 | 60.06 | 51.04 | 40.93 | 31.29 | 28.26 |
| ENSMUSG00000049709 | 12.68 | 21.60 | 16.66 | 16.54 | 25.92 | 22.96 | 44.96 | 32.91 | 34.36 | 23.23 | 19.18 | 27.31 |
| ENSMUSG00000074486 | 92.63 | 81.48 | 78.40 | 77.53 | 73.44 | 75.16 | 141.18 | 153.02 | 134.48 | 76.32 | 109.00 | 84.77 |
| ENSMUSG00000020698 | 7.80 | 7.85 | 4.90 | 11.37 | 7.56 | 9.39 | 9.89 | 8.23 | 10.80 | 5.53 | 7.06 | 5.65 |
| ENSMUSG00000006462 | 12.68 | 12.76 | 13.72 | 18.61 | 9.72 | 12.53 | 16.19 | 17.28 | 17.67 | 9.95 | 13.12 | 9.42 |
| ENSMUSG00000049409 | 10.73 | 15.71 | 5.88 | 12.40 | 10.80 | 15.66 | 12.59 | 11.52 | 13.74 | 7.74 | 6.06 | 10.36 |
| ENSMUSG00000091019 | 15.60 | 18.65 | 11.76 | 22.74 | 20.52 | 12.53 | 19.78 | 22.21 | 23.56 | 17.70 | 12.11 | 12.24 |
| ENSMUSG00000052291 | 26.33 | 24.54 | 32.34 | 21.71 | 29.16 | 38.62 | 19.78 | 20.57 | 25.52 | 13.27 | 11.10 | 17.90 |
| ENSMUSG00000103226 | 5.85 | 2.95 | 3.92 | 3.10 | 9.72 | 3.13 | 7.19 | 7.40 | 5.89 | 4.42 | 4.04 | 4.71 |
| ENSMUSG00000058498 | 8.78 | 15.71 | 13.72 | 21.71 | 10.80 | 15.66 | 15.29 | 13.99 | 17.67 | 8.85 | 12.11 | 9.42 |
| ENSMUSG00000090659 | 22.43 | 25.53 | 29.40 | 29.98 | 41.04 | 36.53 | 33.27 | 24.68 | 31.41 | 22.12 | 21.19 | 15.07 |
| ENSMUSG00000032373 | 64.35 | 51.05 | 46.06 | 54.79 | 55.08 | 76.20 | 75.54 | 79.80 | 84.42 | 44.24 | 53.49 | 59.34 |
| ENSMUSG00000095079 | 26.33 | 28.47 | 31.36 | 34.11 | 27.00 | 16.70 | 51.26 | 55.94 | 45.15 | 27.65 | 36.33 | 36.73 |
| ENSMUSG00000076617 | 403.67 | 385.83 | 344.97 | 374.20 | 398.53 | 331.94 | 896.55 | 757.69 | 698.88 | 491.11 | 519.78 | 547.23 |
| ENSMUSG00000021572 | 27.30 | 25.53 | 29.40 | 25.84 | 44.28 | 35.49 | 44.06 | 41.96 | 46.13 | 26.55 | 33.31 | 28.26 |
| ENSMUSG00000002985 | 767.37 | 586.10 | 599.77 | 564.40 | 441.73 | 501.04 | 856.98 | 965.00 | 826.49 | 567.43 | 617.68 | 581.14 |
| ENSMUSG00000105107 | 2.93 | 4.91 | 0.98 | 3.10 | 3.24 | 4.18 | 7.19 | 5.76 | 6.87 | 5.53 | 4.04 | 3.77 |
| ENSMUSG00000053909 | 29.25 | 36.32 | 40.18 | 37.21 | 51.84 | 35.49 | 55.75 | 57.59 | 51.04 | 29.86 | 38.35 | 43.33 |
| ENSMUSG00000066175 | 7.80 | 2.95 | 6.86 | 8.27 | 9.72 | 13.57 | 6.29 | 6.58 | 7.85 | 4.42 | 4.04 | 5.65 |
| ENSMUSG00000053070 | 27.30 | 22.58 | 25.48 | 21.71 | 36.72 | 32.36 | 22.48 | 26.33 | 25.52 | 13.27 | 19.18 | 18.84 |
| ENSMUSG00000022157 | 181.36 | 163.95 | 137.20 | 143.68 | 159.84 | 170.15 | 294.05 | 314.26 | 254.23 | 170.34 | 199.84 | 225.11 |
| ENSMUSG00000083214 | 2.93 | 6.87 | 4.90 | 7.24 | 5.40 | 4.18 | 8.99 | 7.40 | 6.87 | 4.42 | 6.06 | 5.65 |
| ENSMUSG00000098702 | 30.23 | 40.25 | 40.18 | 27.91 | 37.80 | 33.40 | 35.07 | 35.38 | 28.47 | 22.12 | 20.19 | 26.37 |
| ENSMUSG00000026785 | 59.48 | 51.05 | 66.64 | 55.82 | 41.04 | 38.62 | 59.35 | 60.06 | 62.82 | 45.35 | 40.37 | 43.33 |
| ENSMUSG00000071537 | 53.63 | 51.05 | 52.92 | 42.38 | 33.48 | 48.02 | 60.25 | 57.59 | 53.01 | 42.03 | 36.33 | 43.33 |
| ENSMUSG00000038295 | 57.53 | 59.89 | 72.52 | 79.59 | 58.32 | 85.59 | 96.22 | 95.43 | 90.30 | 65.26 | 59.55 | 76.29 |
| ENSMUSG00000026628 | 62.40 | 57.92 | 59.78 | 64.09 | 62.64 | 52.19 | 89.92 | 90.49 | 77.54 | 59.73 | 72.67 | 51.80 |
| ENSMUSG00000109429 | 4.88 | 2.95 | 6.86 | 5.17 | 20.52 | 3.13 | 6.29 | 4.94 | 5.89 | 4.42 | 4.04 | 3.77 |
| ENSMUSG00000035678 | 226.21 | 232.67 | 172.48 | 196.40 | 171.72 | 191.02 | 250.89 | 272.31 | 247.36 | 173.66 | 179.65 | 198.74 |
| ENSMUSG00000101493 | 3.90 | 10.80 | 17.64 | 10.34 | 14.04 | 13.57 | 12.59 | 13.99 | 11.78 | 9.95 | 9.08 | 8.48 |

| | | | | | | | | | | | | |
|---|---|---|---|---|---|---|---|---|---|---|---|---|
| ENSMUSG00000079029 | 146.26 | 187.51 | 187.18 | 170.56 | 182.52 | 154.49 | 194.24 | 198.27 | 215.95 | 159.28 | 142.31 | 135.63 |
| ENSMUSG00000029153 | 102.38 | 92.28 | 99.96 | 111.64 | 147.96 | 105.43 | 111.51 | 113.53 | 129.57 | 95.13 | 84.78 | 75.35 |
| ENSMUSG00000048520 | 24.38 | 22.58 | 17.64 | 25.84 | 25.92 | 15.66 | 22.48 | 22.21 | 24.54 | 17.70 | 17.16 | 15.07 |
| ENSMUSG00000020566 | 107.26 | 113.88 | 86.24 | 85.80 | 115.56 | 72.02 | 165.46 | 149.73 | 144.29 | 102.87 | 106.98 | 122.44 |
| ENSMUSG00000079657 | 33.15 | 27.49 | 24.50 | 18.61 | 16.20 | 27.14 | 20.68 | 16.45 | 18.65 | 12.17 | 14.13 | 14.13 |
| ENSMUSG00000090176 | 22.43 | 17.67 | 18.62 | 32.04 | 22.68 | 36.53 | 33.27 | 28.79 | 35.34 | 24.33 | 19.18 | 27.31 |
| ENSMUSG00000016194 | 1142.77 | 1073.05 | 981.00 | 993.39 | 1078.95 | 978.08 | 1508.94 | 1381.28 | 1263.29 | 1019.83 | 1006.25 | 1009.70 |
| ENSMUSG00000023345 | 96.53 | 98.17 | 89.18 | 88.90 | 62.64 | 77.24 | 116.00 | 110.24 | 98.16 | 76.32 | 70.65 | 90.42 |
| ENSMUSG00000055313 | 11.70 | 13.74 | 15.68 | 10.34 | 19.44 | 10.44 | 11.69 | 13.16 | 12.76 | 9.95 | 10.09 | 7.54 |
| ENSMUSG00000026077 | 21.45 | 35.34 | 25.48 | 32.04 | 38.88 | 40.71 | 44.06 | 44.42 | 45.15 | 34.29 | 31.29 | 32.97 |
| ENSMUSG00000085487 | 12.68 | 14.73 | 10.78 | 8.27 | 19.44 | 4.18 | 9.89 | 8.23 | 8.83 | 5.53 | 7.06 | 7.54 |
| ENSMUSG00000053559 | 25.35 | 15.71 | 13.72 | 15.51 | 18.36 | 16.70 | 24.28 | 24.68 | 28.47 | 18.80 | 21.19 | 17.90 |
| ENSMUSG00000086644 | 29.25 | 31.42 | 42.14 | 32.04 | 34.56 | 31.32 | 44.96 | 50.18 | 45.15 | 33.18 | 39.36 | 32.97 |
| ENSMUSG00000032420 | 198.91 | 247.40 | 226.38 | 251.19 | 245.17 | 208.77 | 334.52 | 338.94 | 370.05 | 263.25 | 250.30 | 273.15 |
| ENSMUSG00000097296 | 168.68 | 194.39 | 219.52 | 218.11 | 258.13 | 206.68 | 196.04 | 216.36 | 198.28 | 164.81 | 131.21 | 164.83 |
| ENSMUSG00000043153 | 72.15 | 89.34 | 105.84 | 75.46 | 101.52 | 80.38 | 85.43 | 97.90 | 95.21 | 73.00 | 72.67 | 64.99 |
| ENSMUSG00000032942 | 10.73 | 6.87 | 9.80 | 2.07 | 5.40 | 8.35 | 16.19 | 14.81 | 14.72 | 12.17 | 12.11 | 10.36 |
| ENSMUSG00000079304 | 31.20 | 32.40 | 25.48 | 26.88 | 21.60 | 21.92 | 37.77 | 37.02 | 31.41 | 28.76 | 27.25 | 24.49 |
| ENSMUSG00000029136 | 35.10 | 32.40 | 38.22 | 25.84 | 32.40 | 27.14 | 35.07 | 38.67 | 44.17 | 28.76 | 33.31 | 27.31 |
| ENSMUSG00000040907 | 394.90 | 340.67 | 289.11 | 415.55 | 316.45 | 334.03 | 486.49 | 524.87 | 482.93 | 350.64 | 402.70 | 388.05 |
| ENSMUSG00000072905 | 123.83 | 176.71 | 159.74 | 150.92 | 169.56 | 135.70 | 161.86 | 182.63 | 184.54 | 133.84 | 132.22 | 138.46 |
| ENSMUSG00000098090 | 190.14 | 236.60 | 204.82 | 205.71 | 251.65 | 233.82 | 221.21 | 256.68 | 253.25 | 180.30 | 187.73 | 192.14 |
| ENSMUSG00000061411 | 76.05 | 55.96 | 66.64 | 72.36 | 69.12 | 78.29 | 71.04 | 79.80 | 88.34 | 57.52 | 61.57 | 64.99 |
| ENSMUSG00000022489 | 29.25 | 30.43 | 21.56 | 32.04 | 25.92 | 32.36 | 41.37 | 39.49 | 35.34 | 29.86 | 32.30 | 27.31 |
| ENSMUSG00000102224 | 268.14 | 208.13 | 240.11 | 218.11 | 231.13 | 198.33 | 363.30 | 331.54 | 323.92 | 235.60 | 287.65 | 267.49 |
| ENSMUSG00000060459 | 34.13 | 28.47 | 31.36 | 26.88 | 34.56 | 25.05 | 41.37 | 43.60 | 39.26 | 33.18 | 35.32 | 28.26 |
| ENSMUSG00000079559 | 152.11 | 133.52 | 120.54 | 120.94 | 146.88 | 148.23 | 176.25 | 176.88 | 154.11 | 148.22 | 118.09 | 129.04 |
| ENSMUSG00000087088 | 30.23 | 43.20 | 52.92 | 40.31 | 45.36 | 52.19 | 44.06 | 41.96 | 51.04 | 38.71 | 33.31 | 34.85 |
| ENSMUSG00000021947 | 94.58 | 85.41 | 103.88 | 96.13 | 83.16 | 85.59 | 97.12 | 106.95 | 98.16 | 74.11 | 78.72 | 83.83 |
| ENSMUSG00000022070 | 174.54 | 210.09 | 209.72 | 194.34 | 199.81 | 193.11 | 191.54 | 207.32 | 191.41 | 152.64 | 148.36 | 165.77 |
| ENSMUSG00000062694 | 1374.83 | 1269.40 | 1127.02 | 1187.72 | 978.51 | 932.15 | 1698.68 | 1832.93 | 1568.56 | 1300.79 | 1367.58 | 1370.44 |
| ENSMUSG00000021950 | 5805.49 | 5716.70 | 5140.21 | 6009.93 | 5376.38 | 5192.06 | 7777.58 | 8219.39 | 7748.55 | 5803.76 | 6367.56 | 6638.37 |
| ENSMUSG00000005951 | 97.51 | 111.92 | 102.90 | 127.15 | 83.16 | 90.81 | 142.98 | 129.16 | 124.66 | 90.70 | 113.04 | 111.14 |
| ENSMUSG00000024989 | 509.95 | 591.99 | 563.51 | 583.01 | 622.10 | 680.58 | 597.10 | 579.99 | 659.62 | 511.02 | 489.50 | 458.70 |

| | | | | | | | | | | | | |
|---|---|---|---|---|---|---|---|---|---|---|---|---|
| ENSMUSG00000082605 | 17.55 | 10.80 | 9.80 | 13.44 | 19.44 | 11.48 | 18.88 | 15.63 | 17.67 | 13.27 | 13.12 | 15.07 |
| ENSMUSG00000026708 | 210.61 | 218.93 | 261.67 | 209.84 | 255.97 | 268.27 | 215.82 | 225.41 | 234.60 | 175.87 | 185.71 | 176.13 |
| ENSMUSG00000099966 | 91.66 | 89.34 | 114.66 | 78.56 | 137.16 | 134.66 | 89.03 | 87.20 | 82.45 | 71.90 | 67.62 | 66.87 |
| ENSMUSG00000035459 | 70.20 | 72.65 | 59.78 | 64.09 | 58.32 | 58.46 | 101.61 | 93.79 | 101.10 | 78.53 | 69.64 | 88.54 |
| ENSMUSG00000017493 | 1006.26 | 749.07 | 867.32 | 826.96 | 760.34 | 845.51 | 1068.30 | 1052.21 | 1012.00 | 835.11 | 871.01 | 796.83 |
| ENSMUSG00000032783 | 355.90 | 385.83 | 349.87 | 394.87 | 308.89 | 367.43 | 402.86 | 431.08 | 392.63 | 321.88 | 351.23 | 307.05 |
| ENSMUSG00000105206 | 88.73 | 124.68 | 118.58 | 113.71 | 90.72 | 101.25 | 132.19 | 134.92 | 144.29 | 98.44 | 111.02 | 119.62 |
| ENSMUSG00000052131 | 28.28 | 16.69 | 19.60 | 18.61 | 20.52 | 22.96 | 28.78 | 24.68 | 27.48 | 21.02 | 20.19 | 23.55 |
| ENSMUSG00000047443 | 977.98 | 832.52 | 782.06 | 765.97 | 731.18 | 789.14 | 914.53 | 1043.98 | 1001.21 | 849.49 | 779.17 | 740.32 |
| ENSMUSG00000045382 | 691.31 | 681.33 | 683.07 | 659.50 | 753.86 | 721.29 | 824.61 | 870.39 | 968.81 | 711.23 | 672.18 | 748.79 |
| ENSMUSG00000034059 | 59.48 | 56.94 | 55.86 | 60.99 | 51.84 | 62.63 | 76.44 | 69.11 | 65.77 | 59.73 | 59.55 | 49.92 |
| ENSMUSG00000050069 | 22.43 | 21.60 | 18.62 | 18.61 | 22.68 | 17.75 | 27.88 | 27.15 | 24.54 | 21.02 | 21.19 | 21.66 |
| ENSMUSG00000084353 | 360.77 | 367.17 | 352.81 | 363.86 | 379.09 | 339.25 | 388.47 | 406.40 | 408.34 | 298.65 | 335.08 | 332.48 |
| ENSMUSG00000020876 | 412.45 | 350.48 | 395.93 | 368.00 | 311.05 | 281.84 | 407.36 | 431.91 | 365.15 | 311.92 | 342.15 | 313.65 |
| ENSMUSG00000024791 | 187.21 | 202.24 | 186.20 | 208.81 | 166.32 | 172.23 | 191.54 | 199.09 | 197.30 | 142.69 | 166.53 | 163.89 |
| ENSMUSG00000005699 | 133.58 | 103.08 | 100.94 | 103.37 | 95.04 | 88.73 | 104.31 | 103.66 | 93.25 | 81.85 | 82.76 | 78.18 |
| ENSMUSG00000023072 | 515.80 | 427.06 | 476.29 | 509.61 | 379.09 | 375.78 | 563.83 | 579.17 | 557.53 | 421.43 | 482.44 | 469.06 |
| ENSMUSG00000030008 | 172.58 | 132.54 | 128.38 | 135.41 | 118.80 | 117.95 | 145.68 | 160.42 | 145.27 | 110.61 | 134.23 | 121.50 |
| ENSMUSG00000031736 | 203.79 | 189.48 | 186.20 | 193.30 | 183.60 | 149.27 | 200.53 | 232.82 | 201.22 | 167.02 | 163.50 | 184.61 |
| ENSMUSG00000022322 | 540.18 | 672.50 | 672.29 | 542.69 | 727.94 | 699.37 | 598.90 | 640.04 | 615.45 | 508.81 | 492.53 | 509.56 |
| ENSMUSG00000062248 | 1329.00 | 1443.16 | 1351.45 | 1339.67 | 1363.00 | 1217.12 | 1434.30 | 1563.09 | 1589.17 | 1223.36 | 1285.83 | 1230.10 |
| ENSMUSG00000029651 | 34.13 | 22.58 | 33.32 | 36.18 | 18.36 | 29.23 | 35.97 | 37.84 | 40.24 | 27.65 | 34.32 | 31.08 |
| ENSMUSG00000025049 | 128.71 | 157.08 | 145.04 | 157.12 | 133.92 | 154.49 | 159.17 | 144.79 | 152.14 | 121.67 | 131.21 | 119.62 |
| ENSMUSG00000031200 | 50.70 | 62.83 | 82.32 | 69.26 | 78.84 | 99.16 | 62.95 | 60.06 | 67.73 | 57.52 | 47.44 | 50.86 |
| ENSMUSG00000022096 | 464.13 | 405.46 | 389.07 | 452.76 | 330.49 | 388.31 | 485.59 | 493.61 | 467.23 | 417.00 | 361.32 | 404.07 |
| ENSMUSG00000025486 | 151.13 | 120.75 | 131.32 | 130.25 | 114.48 | 104.38 | 150.17 | 148.90 | 129.57 | 112.82 | 120.10 | 117.74 |
| ENSMUSG00000033031 | 985.78 | 1147.66 | 1146.62 | 998.55 | 1404.04 | 1301.67 | 1151.04 | 1135.30 | 1163.17 | 949.04 | 906.34 | 967.31 |
| ENSMUSG00000091462 | 375.40 | 345.57 | 336.15 | 310.11 | 349.93 | 341.34 | 329.12 | 340.59 | 328.83 | 273.21 | 262.41 | 281.62 |
| ENSMUSG00000037544 | 866.83 | 956.22 | 898.68 | 978.91 | 873.74 | 995.82 | 1032.33 | 1105.68 | 1076.79 | 830.69 | 916.43 | 885.37 |
| ENSMUSG00000044576 | 18.53 | 10.80 | 11.76 | 12.40 | 10.80 | 14.61 | 12.59 | 11.52 | 11.78 | 9.95 | 9.08 | 10.36 |
| ENSMUSG00000079553 | 459.25 | 460.44 | 491.97 | 516.85 | 347.77 | 419.62 | 544.94 | 593.97 | 505.51 | 435.81 | 466.29 | 449.28 |
| ENSMUSG00000003779 | 3105.55 | 3288.85 | 3105.69 | 3169.32 | 3074.84 | 3287.05 | 3614.07 | 3776.10 | 3734.89 | 2894.69 | 3106.57 | 3144.00 |
| ENSMUSG00000092454 | 268.14 | 177.70 | 238.15 | 201.57 | 176.04 | 159.71 | 232.90 | 249.27 | 214.96 | 180.30 | 203.87 | 189.32 |
| ENSMUSG00000027331 | 1314.38 | 1426.48 | 1454.35 | 1389.29 | 1310.07 | 1481.21 | 1505.34 | 1565.56 | 1488.07 | 1211.19 | 1242.43 | 1301.68 |

| | | | | | | | | | | | | |
|---|---|---|---|---|---|---|---|---|---|---|---|---|
| ENSMUSG00000022584 | 2176.33 | 1812.30 | 1812.06 | 1841.02 | 1836.05 | 1654.49 | 2162.69 | 2340.52 | 2117.26 | 1929.06 | 1780.37 | 1744.36 |
| ENSMUSG00000063415 | 158.93 | 172.79 | 133.28 | 136.45 | 103.68 | 159.71 | 215.82 | 194.15 | 199.26 | 161.49 | 167.54 | 173.31 |
| ENSMUSG00000026622 | 739.09 | 737.29 | 743.84 | 717.39 | 666.38 | 765.13 | 820.11 | 898.37 | 886.36 | 689.11 | 735.77 | 723.36 |
| ENSMUSG00000037313 | 1446.01 | 1445.13 | 1511.19 | 1357.25 | 1555.24 | 1559.50 | 1550.30 | 1666.75 | 1526.35 | 1286.41 | 1356.47 | 1269.65 |
| ENSMUSG00000022033 | 877.55 | 924.80 | 931.02 | 813.52 | 1026.03 | 923.80 | 853.38 | 924.69 | 858.88 | 739.99 | 704.48 | 730.90 |
| ENSMUSG00000020309 | 325.67 | 313.18 | 351.83 | 265.66 | 516.25 | 374.74 | 299.45 | 348.82 | 313.12 | 288.69 | 247.27 | 258.08 |
| ENSMUSG00000056458 | 103.36 | 96.21 | 86.24 | 84.76 | 92.88 | 75.16 | 112.41 | 106.13 | 97.18 | 92.91 | 87.81 | 80.06 |
| ENSMUSG00000082693 | 130.66 | 130.57 | 151.90 | 131.28 | 136.08 | 125.26 | 140.28 | 139.86 | 133.49 | 107.29 | 112.03 | 122.44 |
| ENSMUSG00000069910 | 285.69 | 340.67 | 347.91 | 340.09 | 316.45 | 336.12 | 364.19 | 378.43 | 343.55 | 298.65 | 302.78 | 298.58 |
| ENSMUSG00000017716 | 1198.34 | 1245.83 | 1319.11 | 1076.08 | 1195.59 | 1200.42 | 1228.37 | 1313.00 | 1149.42 | 997.71 | 1062.77 | 1004.04 |
| ENSMUSG00000030867 | 899.00 | 963.09 | 837.92 | 940.67 | 733.34 | 844.47 | 1008.95 | 1142.70 | 1089.55 | 916.97 | 901.29 | 879.72 |
| ENSMUSG00000051457 | 493.38 | 467.31 | 494.91 | 541.66 | 421.21 | 494.78 | 526.96 | 589.86 | 593.85 | 506.60 | 453.17 | 464.35 |
| ENSMUSG00000031007 | 2260.18 | 2327.72 | 2277.57 | 2012.61 | 2985.20 | 2364.30 | 2562.85 | 2658.90 | 2552.10 | 2155.81 | 2108.39 | 2209.65 |
| ENSMUSG00000084404 | 119.93 | 116.83 | 118.58 | 114.74 | 154.44 | 122.13 | 145.68 | 148.08 | 135.46 | 117.25 | 116.07 | 124.33 |
| ENSMUSG00000078762 | 195.01 | 193.40 | 187.18 | 204.67 | 143.64 | 148.23 | 175.35 | 196.62 | 201.22 | 170.34 | 162.49 | 145.05 |
| ENSMUSG00000028295 | 312.02 | 290.60 | 331.25 | 261.53 | 355.33 | 307.93 | 260.78 | 245.98 | 249.32 | 211.27 | 226.08 | 193.09 |
| ENSMUSG00000016206 | 83.85 | 76.58 | 80.36 | 83.73 | 71.28 | 74.11 | 91.72 | 88.85 | 85.40 | 73.00 | 72.67 | 76.29 |
| ENSMUSG00000030654 | 1066.71 | 1148.64 | 1126.04 | 1031.63 | 1288.47 | 1165.97 | 1121.36 | 1230.73 | 1164.15 | 1061.87 | 918.45 | 955.07 |
| ENSMUSG00000048709 | 167.71 | 171.81 | 148.96 | 172.63 | 132.84 | 121.09 | 182.55 | 180.17 | 175.70 | 154.86 | 152.40 | 142.22 |
| ENSMUSG00000032400 | 340.29 | 441.79 | 390.05 | 343.19 | 415.81 | 461.38 | 348.01 | 369.38 | 400.48 | 324.09 | 294.71 | 315.53 |
| ENSMUSG00000082484 | 21.45 | 22.58 | 12.74 | 19.64 | 14.04 | 15.66 | 24.28 | 23.86 | 24.54 | 19.91 | 19.18 | 21.66 |
| ENSMUSG00000037902 | 498.25 | 458.47 | 432.19 | 484.81 | 405.01 | 491.65 | 596.20 | 547.90 | 582.07 | 508.81 | 457.20 | 478.48 |
| ENSMUSG00000043329 | 143.33 | 101.12 | 132.30 | 116.81 | 114.48 | 99.16 | 139.38 | 143.15 | 130.55 | 111.72 | 123.13 | 111.14 |
| ENSMUSG00000074280 | 198.91 | 238.56 | 202.86 | 649.16 | 190.08 | 230.69 | 231.11 | 260.79 | 231.65 | 202.42 | 194.79 | 209.10 |
| ENSMUSG00000036390 | 4145.94 | 3888.69 | 3824.04 | 3710.98 | 3672.10 | 3715.03 | 5196.74 | 5427.22 | 4744.93 | 4219.81 | 4264.22 | 4412.71 |
| ENSMUSG00000027496 | 681.56 | 619.48 | 660.53 | 652.26 | 564.85 | 661.79 | 633.97 | 732.18 | 677.29 | 540.89 | 557.12 | 617.87 |
| ENSMUSG00000019810 | 156.98 | 140.39 | 138.18 | 151.95 | 142.56 | 136.74 | 154.67 | 154.66 | 141.35 | 132.73 | 126.16 | 119.62 |
| ENSMUSG00000061451 | 289.59 | 213.04 | 254.81 | 234.65 | 172.80 | 188.94 | 304.84 | 319.20 | 292.51 | 236.71 | 258.38 | 275.03 |
| ENSMUSG00000048489 | 195.99 | 171.81 | 156.80 | 152.99 | 154.44 | 149.27 | 196.93 | 198.27 | 187.48 | 157.07 | 172.59 | 160.12 |
| ENSMUSG00000035365 | 270.09 | 336.74 | 320.47 | 321.48 | 372.61 | 421.71 | 329.12 | 306.86 | 347.48 | 276.53 | 281.59 | 269.38 |
| ENSMUSG00000036528 | 75.08 | 84.43 | 67.62 | 68.22 | 74.52 | 81.42 | 117.80 | 113.53 | 106.99 | 96.23 | 100.93 | 87.59 |
| ENSMUSG00000070371 | 61.43 | 41.23 | 49.98 | 44.45 | 39.96 | 38.62 | 47.66 | 41.96 | 45.15 | 36.50 | 38.35 | 38.62 |
| ENSMUSG00000027583 | 52.65 | 55.96 | 69.58 | 58.92 | 72.36 | 65.76 | 90.82 | 83.09 | 82.45 | 68.58 | 75.70 | 71.58 |
| ENSMUSG00000029521 | 151.13 | 159.04 | 150.92 | 158.16 | 181.44 | 172.23 | 174.45 | 155.49 | 156.07 | 143.79 | 129.19 | 136.57 |

| | | | | | | | | | | | | |
|---|---|---|---|---|---|---|---|---|---|---|---|---|
| ENSMUSG00000032925 | 51.68 | 58.90 | 88.20 | 83.73 | 74.52 | 76.20 | 80.93 | 75.69 | 84.42 | 64.15 | 69.64 | 69.70 |
| ENSMUSG00000062874 | 368.57 | 319.07 | 294.99 | 346.29 | 274.33 | 254.70 | 375.88 | 408.05 | 371.04 | 310.82 | 320.95 | 343.79 |
| ENSMUSG00000021391 | 244.74 | 264.09 | 256.77 | 219.14 | 291.61 | 247.39 | 205.03 | 215.54 | 212.02 | 174.77 | 189.75 | 170.48 |
| ENSMUSG00000058979 | 161.86 | 118.79 | 131.32 | 120.94 | 114.48 | 104.38 | 127.69 | 136.56 | 141.35 | 107.29 | 123.13 | 113.03 |
| ENSMUSG00000039452 | 115.06 | 115.85 | 111.72 | 114.74 | 97.20 | 104.38 | 100.72 | 97.08 | 109.94 | 89.59 | 80.74 | 90.42 |
| ENSMUSG00000026683 | 1297.80 | 1449.06 | 1407.31 | 1269.38 | 1622.20 | 1616.91 | 1368.65 | 1461.08 | 1453.71 | 1223.36 | 1240.41 | 1169.82 |
| ENSMUSG00000097006 | 53.63 | 64.80 | 65.66 | 56.85 | 46.44 | 38.62 | 58.45 | 57.59 | 60.86 | 48.67 | 51.47 | 49.92 |
| ENSMUSG00000021965 | 224.26 | 236.60 | 223.44 | 216.04 | 239.77 | 248.43 | 215.82 | 227.06 | 216.93 | 188.04 | 194.79 | 177.07 |
| ENSMUSG00000020101 | 527.51 | 444.73 | 431.21 | 467.23 | 344.53 | 432.15 | 545.84 | 491.14 | 488.82 | 450.19 | 422.89 | 421.96 |
| ENSMUSG00000048922 | 376.37 | 434.91 | 423.37 | 456.90 | 433.09 | 471.82 | 387.58 | 421.21 | 419.13 | 334.05 | 351.23 | 357.91 |
| ENSMUSG00000027569 | 474.85 | 456.51 | 452.77 | 493.07 | 523.81 | 458.25 | 455.92 | 518.29 | 494.71 | 434.70 | 394.63 | 421.96 |
| ENSMUSG00000026975 | 196.96 | 178.68 | 214.62 | 200.54 | 168.48 | 200.42 | 224.81 | 229.53 | 202.20 | 176.98 | 183.69 | 198.74 |
| ENSMUSG00000019942 | 1788.25 | 1969.38 | 1898.30 | 1815.18 | 1761.53 | 1843.42 | 1845.25 | 2037.78 | 1938.61 | 1635.94 | 1643.11 | 1698.21 |
| ENSMUSG00000101878 | 1346.55 | 1425.49 | 1253.45 | 1408.93 | 1181.55 | 1040.71 | 1514.33 | 1710.35 | 1610.76 | 1345.03 | 1351.43 | 1440.13 |
| ENSMUSG00000058258 | 1945.24 | 2119.59 | 2091.36 | 1953.69 | 2972.24 | 2394.57 | 2241.82 | 2134.85 | 2108.42 | 1829.51 | 1779.36 | 1944.04 |
| ENSMUSG00000027306 | 940.93 | 1096.61 | 1190.73 | 1121.56 | 1281.99 | 1305.84 | 1043.13 | 1094.16 | 1160.22 | 946.83 | 944.69 | 932.46 |
| ENSMUSG00000028678 | 824.90 | 829.57 | 827.14 | 919.99 | 700.94 | 905.01 | 914.53 | 1005.31 | 1007.10 | 763.22 | 862.94 | 881.60 |
| ENSMUSG00000040435 | 894.13 | 866.88 | 898.68 | 837.30 | 719.30 | 762.00 | 1062.01 | 1076.06 | 967.83 | 893.74 | 909.36 | 860.88 |
| ENSMUSG00000090188 | 39.00 | 38.29 | 31.36 | 50.65 | 38.88 | 26.10 | 57.55 | 55.12 | 53.99 | 47.56 | 46.43 | 48.98 |
| ENSMUSG00000030116 | 1235.40 | 1148.64 | 1017.26 | 1132.93 | 1215.03 | 1093.94 | 1377.65 | 1371.41 | 1281.94 | 1123.81 | 1171.78 | 1163.22 |
| ENSMUSG00000042549 | 67.28 | 58.90 | 49.00 | 64.09 | 56.16 | 62.63 | 79.13 | 77.33 | 70.67 | 66.37 | 66.61 | 62.16 |
| ENSMUSG00000031604 | 1795.08 | 1783.83 | 1725.82 | 1663.22 | 2416.02 | 1888.31 | 1958.56 | 1931.65 | 1795.30 | 1624.88 | 1565.40 | 1699.15 |
| ENSMUSG00000028555 | 30.23 | 36.32 | 39.20 | 40.31 | 34.56 | 42.80 | 42.26 | 40.31 | 42.21 | 34.29 | 36.33 | 36.73 |
| ENSMUSG00000029627 | 149.18 | 130.57 | 140.14 | 137.48 | 100.44 | 154.49 | 140.28 | 149.73 | 154.11 | 130.52 | 118.09 | 133.75 |
| ENSMUSG00000005481 | 2734.06 | 2648.75 | 2789.14 | 2651.44 | 2494.86 | 2393.53 | 2900.07 | 3200.22 | 2848.53 | 2619.27 | 2468.70 | 2619.37 |
| ENSMUSG00000004267 | 324.69 | 325.94 | 317.53 | 409.34 | 344.53 | 382.05 | 384.88 | 415.45 | 391.65 | 358.38 | 348.20 | 320.24 |
| ENSMUSG00000023015 | 3179.66 | 3114.09 | 3048.84 | 3077.32 | 2815.63 | 3108.56 | 3257.97 | 3500.50 | 3364.84 | 2907.96 | 2861.31 | 2973.52 |
| ENSMUSG00000022881 | 304.22 | 266.05 | 349.87 | 265.66 | 305.65 | 297.49 | 276.07 | 301.92 | 277.79 | 257.72 | 228.10 | 253.37 |
| ENSMUSG00000027715 | 2258.23 | 2348.33 | 2461.81 | 2164.57 | 2781.07 | 2693.11 | 2261.60 | 2410.45 | 2472.59 | 2145.85 | 1915.62 | 2118.29 |
| ENSMUSG00000033186 | 1175.92 | 1175.15 | 1226.01 | 959.27 | 1698.88 | 1283.92 | 1104.27 | 1076.89 | 1068.94 | 976.70 | 902.30 | 932.46 |
| ENSMUSG00000079507 | 227.19 | 190.46 | 204.82 | 189.17 | 100.44 | 129.44 | 189.74 | 197.44 | 181.59 | 159.28 | 177.63 | 156.35 |
| ENSMUSG00000038544 | 386.12 | 427.06 | 441.01 | 397.97 | 425.53 | 434.24 | 426.24 | 438.49 | 441.71 | 384.93 | 367.38 | 381.46 |
| ENSMUSG00000030677 | 638.66 | 624.39 | 638.97 | 621.25 | 529.21 | 633.61 | 648.36 | 705.86 | 672.38 | 557.48 | 602.54 | 599.98 |
| ENSMUSG00000048174 | 51.68 | 41.23 | 44.10 | 45.48 | 51.84 | 37.58 | 54.85 | 59.23 | 59.88 | 49.77 | 50.46 | 50.86 |

| | | | | | | | | | | | | |
|---|---|---|---|---|---|---|---|---|---|---|---|---|
| ENSMUSG00000023952 | 539.21 | 480.07 | 456.69 | 546.83 | 433.09 | 401.88 | 504.48 | 551.20 | 527.11 | 457.93 | 436.01 | 483.18 |
| ENSMUSG00000022177 | 527.51 | 480.07 | 415.53 | 448.63 | 371.53 | 406.05 | 471.21 | 506.77 | 465.27 | 420.32 | 440.05 | 395.59 |
| ENSMUSG00000040034 | 340.29 | 318.09 | 347.91 | 314.24 | 372.61 | 324.63 | 350.71 | 334.01 | 317.05 | 306.39 | 273.52 | 291.98 |
| ENSMUSG00000023224 | 555.78 | 472.22 | 455.71 | 471.37 | 432.01 | 545.93 | 582.71 | 584.93 | 574.22 | 520.98 | 522.81 | 472.82 |
| ENSMUSG00000002055 | 829.77 | 927.75 | 875.16 | 945.83 | 798.14 | 959.29 | 946.91 | 1020.95 | 1039.49 | 895.95 | 901.29 | 825.09 |
| ENSMUSG00000066152 | 275.94 | 250.34 | 274.41 | 253.26 | 264.61 | 236.95 | 276.97 | 273.13 | 268.95 | 226.75 | 248.28 | 241.12 |
| ENSMUSG00000021273 | 2936.87 | 2932.47 | 2631.36 | 2909.86 | 2868.55 | 2744.26 | 3686.01 | 3551.51 | 3345.21 | 2977.65 | 2988.48 | 3291.87 |
| ENSMUSG00000044337 | 544.08 | 564.50 | 512.55 | 639.86 | 567.01 | 580.38 | 624.98 | 684.47 | 653.73 | 585.13 | 569.24 | 563.24 |
| ENSMUSG00000048696 | 695.22 | 607.70 | 608.59 | 626.42 | 575.65 | 575.16 | 698.71 | 745.35 | 694.96 | 616.10 | 632.82 | 625.41 |
| ENSMUSG00000052364 | 72.15 | 66.76 | 66.64 | 62.02 | 75.60 | 52.19 | 50.36 | 53.47 | 48.10 | 46.46 | 44.41 | 42.38 |
| ENSMUSG00000033096 | 624.04 | 659.73 | 665.43 | 543.73 | 616.70 | 582.46 | 589.91 | 624.41 | 588.95 | 558.59 | 490.51 | 533.10 |
| ENSMUSG00000005233 | 744.94 | 894.37 | 924.16 | 806.29 | 1056.27 | 981.21 | 822.81 | 844.89 | 868.69 | 787.55 | 705.49 | 737.49 |
| ENSMUSG00000084113 | 129.68 | 136.46 | 136.22 | 139.55 | 181.44 | 144.05 | 163.66 | 179.34 | 162.94 | 150.43 | 151.39 | 143.17 |
| ENSMUSG00000042770 | 182.34 | 182.60 | 190.12 | 176.76 | 179.28 | 145.09 | 178.95 | 194.98 | 180.61 | 160.39 | 171.58 | 156.35 |
| ENSMUSG00000004896 | 235.96 | 229.73 | 245.99 | 253.26 | 284.05 | 296.45 | 250.89 | 270.66 | 244.41 | 227.86 | 215.99 | 230.76 |
| ENSMUSG00000029217 | 136.51 | 148.24 | 116.62 | 162.29 | 130.68 | 137.79 | 158.27 | 160.42 | 157.05 | 143.79 | 137.26 | 138.46 |
| ENSMUSG00000022722 | 450.48 | 438.84 | 427.29 | 357.66 | 513.01 | 461.38 | 394.77 | 417.10 | 394.59 | 380.50 | 344.17 | 340.02 |
| ENSMUSG00000025353 | 430.00 | 386.81 | 383.19 | 334.92 | 411.49 | 348.64 | 375.88 | 408.87 | 377.91 | 334.05 | 355.27 | 337.19 |
| ENSMUSG00000020846 | 2115.87 | 1982.14 | 1898.30 | 1927.85 | 2342.58 | 2131.52 | 2310.16 | 2177.63 | 2111.37 | 1918.00 | 1971.13 | 1937.45 |
| ENSMUSG00000038379 | 483.63 | 547.81 | 533.13 | 519.95 | 605.90 | 606.47 | 577.32 | 528.16 | 579.13 | 525.40 | 483.45 | 479.42 |
| ENSMUSG00000034311 | 902.90 | 1025.92 | 946.70 | 1121.56 | 1000.11 | 1127.35 | 1073.70 | 1054.67 | 1152.37 | 935.77 | 1017.36 | 947.53 |
| ENSMUSG00000027469 | 2490.29 | 2713.54 | 2578.43 | 2605.96 | 2857.75 | 2921.71 | 2829.93 | 2946.84 | 3004.60 | 2534.10 | 2556.51 | 2674.00 |
| ENSMUSG00000090266 | 393.92 | 379.94 | 364.57 | 318.38 | 410.41 | 342.38 | 361.50 | 371.85 | 352.39 | 338.47 | 302.78 | 319.30 |
| ENSMUSG00000001517 | 1345.58 | 1393.10 | 1316.17 | 1429.61 | 1151.31 | 1464.51 | 1390.23 | 1490.70 | 1491.01 | 1319.59 | 1253.53 | 1296.03 |
| ENSMUSG00000040084 | 1134.97 | 1204.60 | 1152.50 | 1192.89 | 1121.07 | 1237.99 | 1242.76 | 1269.39 | 1346.72 | 1142.61 | 1164.71 | 1111.42 |
| ENSMUSG00000037971 | 273.02 | 255.25 | 269.51 | 276.00 | 294.85 | 264.09 | 298.55 | 296.16 | 316.07 | 268.78 | 264.43 | 275.03 |
| ENSMUSG00000056220 | 424.15 | 494.80 | 512.55 | 526.15 | 635.06 | 536.53 | 483.79 | 462.35 | 506.49 | 403.73 | 438.03 | 447.39 |
| ENSMUSG00000042116 | 352.00 | 283.72 | 306.75 | 284.27 | 236.53 | 205.64 | 306.64 | 321.67 | 290.55 | 279.85 | 267.46 | 268.44 |
| ENSMUSG00000087385 | 50.70 | 39.27 | 44.10 | 41.35 | 37.80 | 42.80 | 42.26 | 40.31 | 44.17 | 37.61 | 36.33 | 38.62 |
| ENSMUSG00000083879 | 85.80 | 77.56 | 89.18 | 92.00 | 79.92 | 92.90 | 90.82 | 85.56 | 86.38 | 79.64 | 76.71 | 77.23 |
| ENSMUSG00000091952 | 786.87 | 815.83 | 771.28 | 790.78 | 710.66 | 741.13 | 923.53 | 965.83 | 898.14 | 832.90 | 811.46 | 835.45 |
| ENSMUSG00000024206 | 163.81 | 171.81 | 132.30 | 145.75 | 140.40 | 149.27 | 199.63 | 181.81 | 189.44 | 165.92 | 173.60 | 168.60 |
| ENSMUSG00000017314 | 217.44 | 203.22 | 186.20 | 200.54 | 147.96 | 206.68 | 208.63 | 221.30 | 221.84 | 189.14 | 184.70 | 206.27 |
| ENSMUSG00000012483 | 497.28 | 440.80 | 495.89 | 371.10 | 473.05 | 448.85 | 415.45 | 404.76 | 394.59 | 372.76 | 371.42 | 337.19 |

| | | | | | | | | | | | | |
|---|---|---|---|---|---|---|---|---|---|---|---|---|
| ENSMUSG00000022895 | 4041.61 | 3916.18 | 3869.12 | 4046.93 | 3984.22 | 4002.08 | 4261.53 | 4197.31 | 4464.20 | 3864.75 | 3773.71 | 3867.36 |
| ENSMUSG00000033965 | 318.84 | 322.01 | 290.09 | 321.48 | 305.65 | 343.42 | 383.08 | 388.30 | 361.22 | 329.62 | 338.11 | 340.96 |
| ENSMUSG00000029093 | 1966.69 | 1974.29 | 1670.94 | 2077.74 | 1588.72 | 1877.87 | 2439.65 | 2393.17 | 2405.84 | 2148.07 | 2063.98 | 2238.85 |
| ENSMUSG00000029659 | 3305.44 | 3232.89 | 3111.57 | 2953.28 | 3431.25 | 3065.76 | 3052.94 | 3331.03 | 3141.04 | 2714.39 | 2861.31 | 2914.18 |
| ENSMUSG00000030269 | 498.25 | 446.69 | 513.53 | 423.82 | 434.17 | 438.41 | 480.20 | 463.99 | 493.73 | 436.91 | 424.91 | 420.08 |
| ENSMUSG00000073700 | 1095.96 | 898.30 | 1032.94 | 925.16 | 806.78 | 910.23 | 1031.44 | 1052.21 | 970.78 | 869.40 | 935.60 | 920.22 |
| ENSMUSG00000054519 | 246.69 | 286.67 | 233.24 | 252.22 | 267.85 | 260.96 | 257.18 | 233.64 | 237.54 | 210.16 | 227.09 | 212.86 |
| ENSMUSG00000028020 | 460.23 | 466.33 | 411.61 | 418.65 | 655.58 | 549.06 | 459.51 | 420.39 | 450.54 | 414.79 | 377.47 | 396.53 |
| ENSMUSG00000018362 | 3087.03 | 3295.72 | 3172.33 | 3145.55 | 3629.97 | 3371.60 | 3381.17 | 3388.62 | 3485.57 | 2993.13 | 2993.53 | 3185.44 |
| ENSMUSG00000043004 | 832.70 | 954.26 | 809.50 | 714.29 | 1085.43 | 937.37 | 927.12 | 890.96 | 942.31 | 860.55 | 806.42 | 802.48 |
| ENSMUSG00000022109 | 612.34 | 594.94 | 598.79 | 560.27 | 572.41 | 625.26 | 606.09 | 649.09 | 600.72 | 536.46 | 582.36 | 541.58 |
| ENSMUSG00000062480 | 491.43 | 484.00 | 414.55 | 466.20 | 451.45 | 411.27 | 536.85 | 555.31 | 508.46 | 467.88 | 471.33 | 493.55 |
| ENSMUSG00000054520 | 627.94 | 650.90 | 545.87 | 615.05 | 507.61 | 554.28 | 739.18 | 723.96 | 703.79 | 628.27 | 652.00 | 662.14 |
| ENSMUSG00000037852 | 1709.27 | 1803.47 | 1767.96 | 1698.37 | 1949.45 | 1621.08 | 1970.25 | 1985.13 | 1998.49 | 1729.96 | 1811.66 | 1809.35 |
| ENSMUSG00000037035 | 1691.72 | 1558.03 | 1603.31 | 1719.04 | 1597.36 | 1812.11 | 1860.54 | 1905.33 | 1879.72 | 1715.58 | 1744.04 | 1616.27 |
| ENSMUSG00000051378 | 445.60 | 430.99 | 405.73 | 494.11 | 407.17 | 427.97 | 477.50 | 515.00 | 484.90 | 447.97 | 438.03 | 443.63 |
| ENSMUSG00000064080 | 10139.61 | 10016.74 | 8849.59 | 10371.11 | 7615.28 | 8444.67 | 11098.50 | 11483.78 | 11417.68 | 10012.51 | 10113.00 | 10496.31 |
| ENSMUSG00000023963 | 483.63 | 497.74 | 494.91 | 370.06 | 515.17 | 492.69 | 459.51 | 436.84 | 447.60 | 424.75 | 400.68 | 385.23 |
| ENSMUSG00000035683 | 437.80 | 506.58 | 509.61 | 454.83 | 507.61 | 524.01 | 453.22 | 463.17 | 492.75 | 407.05 | 440.05 | 422.90 |
| ENSMUSG00000036611 | 110.18 | 84.43 | 82.32 | 66.16 | 78.84 | 69.94 | 100.72 | 106.13 | 98.16 | 90.70 | 93.86 | 90.42 |
| ENSMUSG00000079555 | 582.11 | 662.68 | 639.95 | 640.89 | 855.38 | 777.66 | 589.01 | 621.95 | 632.13 | 566.33 | 547.03 | 549.12 |
| ENSMUSG00000023861 | 2096.37 | 1775.98 | 2004.14 | 1683.90 | 2324.22 | 1839.25 | 1899.21 | 1934.12 | 1783.52 | 1707.83 | 1712.75 | 1649.23 |
| ENSMUSG00000024246 | 118.96 | 100.14 | 144.06 | 114.74 | 139.32 | 122.13 | 129.49 | 124.22 | 120.73 | 112.82 | 114.05 | 111.14 |
| ENSMUSG00000056394 | 1126.19 | 1145.70 | 1153.48 | 1212.53 | 977.43 | 1133.61 | 1215.78 | 1263.64 | 1282.92 | 1103.90 | 1199.03 | 1094.46 |
| ENSMUSG00000082998 | 248.64 | 257.22 | 229.32 | 272.90 | 262.45 | 235.91 | 283.26 | 292.87 | 280.73 | 259.94 | 264.43 | 249.60 |
| ENSMUSG00000069083 | 1868.21 | 2105.84 | 1933.58 | 1990.91 | 2231.34 | 2029.23 | 2226.53 | 2245.91 | 2368.54 | 2060.68 | 1954.98 | 2163.50 |
| ENSMUSG00000073434 | 159.91 | 170.82 | 165.62 | 195.37 | 162.00 | 171.19 | 157.37 | 153.02 | 164.90 | 143.79 | 144.33 | 141.28 |
| ENSMUSG00000021113 | 214.51 | 249.36 | 232.26 | 222.25 | 278.65 | 259.92 | 249.99 | 259.97 | 243.43 | 225.65 | 233.14 | 222.28 |
| ENSMUSG00000017764 | 400.75 | 289.61 | 316.55 | 321.48 | 279.73 | 296.45 | 366.89 | 343.88 | 340.61 | 329.62 | 314.90 | 306.11 |
| ENSMUSG00000024512 | 1887.71 | 1955.64 | 1709.16 | 1911.31 | 2081.21 | 2033.40 | 2064.67 | 2227.82 | 2131.00 | 1933.48 | 1875.25 | 2000.55 |
| ENSMUSG00000037580 | 228.16 | 201.26 | 174.44 | 157.12 | 180.36 | 151.36 | 193.34 | 187.57 | 180.61 | 173.66 | 167.54 | 166.71 |
| ENSMUSG00000083220 | 268.14 | 339.68 | 297.93 | 333.88 | 343.45 | 348.64 | 325.53 | 320.84 | 338.64 | 314.14 | 284.62 | 292.92 |
| ENSMUSG00000029454 | 1169.09 | 1007.27 | 1002.56 | 1053.34 | 1021.71 | 1057.41 | 1125.86 | 1137.77 | 1050.29 | 1006.56 | 979.00 | 1014.40 |
| ENSMUSG00000011179 | 4688.07 | 4342.26 | 4392.45 | 4310.53 | 4134.35 | 4009.39 | 5080.74 | 5382.79 | 5020.76 | 4600.31 | 4620.49 | 4797.00 |

| | | | | | | | | | | | | |
|---|---|---|---|---|---|---|---|---|---|---|---|---|
| ENSMUSG00000027082 | 905.83 | 1075.99 | 1022.16 | 967.54 | 1271.19 | 1207.72 | 1020.64 | 1006.96 | 1023.78 | 923.60 | 899.27 | 940.94 |
| ENSMUSG00000026880 | 3821.25 | 3471.45 | 3594.72 | 3283.03 | 3663.46 | 3832.98 | 4161.71 | 4020.43 | 4036.24 | 3641.31 | 3757.56 | 3672.39 |
| ENSMUSG00000041117 | 741.04 | 679.37 | 705.61 | 660.53 | 500.05 | 670.15 | 758.96 | 782.37 | 739.13 | 674.73 | 694.39 | 698.88 |
| ENSMUSG00000031487 | 285.69 | 262.13 | 277.35 | 258.42 | 248.41 | 188.94 | 271.57 | 274.77 | 256.19 | 231.18 | 244.25 | 252.42 |
| ENSMUSG00000030539 | 561.63 | 495.78 | 475.31 | 509.61 | 504.37 | 528.18 | 571.92 | 586.57 | 589.93 | 537.57 | 535.93 | 513.32 |
| ENSMUSG00000026869 | 1587.39 | 1474.58 | 1557.25 | 1404.80 | 1657.84 | 1568.89 | 1513.43 | 1490.70 | 1435.06 | 1335.08 | 1319.13 | 1376.09 |
| ENSMUSG00000103436 | 658.16 | 543.89 | 601.73 | 560.27 | 505.45 | 479.12 | 672.64 | 703.39 | 659.62 | 603.94 | 625.75 | 618.82 |
| ENSMUSG00000013822 | 683.51 | 540.94 | 569.39 | 563.37 | 514.09 | 421.71 | 567.42 | 554.49 | 568.33 | 501.07 | 502.62 | 531.22 |
| ENSMUSG00000021930 | 342.24 | 382.88 | 416.51 | 385.57 | 519.49 | 408.14 | 408.26 | 400.64 | 402.45 | 380.50 | 362.33 | 357.91 |
| ENSMUSG00000105340 | 1099.86 | 961.13 | 960.42 | 1008.89 | 970.95 | 1010.44 | 1048.52 | 1090.87 | 1001.21 | 966.74 | 921.47 | 967.31 |
| ENSMUSG00000002733 | 855.12 | 870.81 | 910.44 | 864.17 | 1122.15 | 1018.79 | 943.31 | 1006.14 | 997.28 | 932.45 | 860.92 | 887.25 |
| ENSMUSG00000066441 | 851.22 | 810.92 | 877.12 | 810.42 | 945.02 | 949.89 | 868.67 | 840.78 | 864.77 | 754.37 | 789.26 | 799.66 |
| ENSMUSG00000053253 | 1613.72 | 1684.67 | 1626.83 | 1536.08 | 1994.81 | 1695.20 | 1758.03 | 1776.16 | 1697.14 | 1606.07 | 1549.25 | 1612.50 |
| ENSMUSG00000041471 | 167.71 | 184.57 | 176.40 | 175.73 | 191.16 | 193.11 | 156.47 | 157.95 | 164.90 | 144.90 | 141.30 | 150.70 |
| ENSMUSG00000086922 | 2351.84 | 2492.65 | 2449.07 | 2253.46 | 2740.03 | 2442.59 | 2409.08 | 2541.26 | 2533.45 | 2283.01 | 2229.50 | 2309.49 |
| ENSMUSG00000022131 | 751.77 | 669.55 | 680.13 | 658.47 | 648.02 | 626.30 | 709.51 | 682.00 | 687.10 | 638.23 | 637.87 | 618.82 |
| ENSMUSG00000028614 | 633.79 | 666.60 | 662.49 | 584.04 | 794.90 | 820.46 | 651.05 | 631.82 | 669.43 | 611.68 | 572.26 | 596.21 |
| ENSMUSG00000043923 | 136.51 | 144.32 | 136.22 | 154.02 | 162.00 | 128.39 | 135.79 | 129.98 | 133.49 | 119.46 | 125.15 | 119.62 |
| ENSMUSG00000030750 | 785.90 | 739.25 | 765.40 | 737.03 | 682.58 | 646.14 | 770.65 | 762.62 | 720.48 | 686.89 | 697.41 | 677.21 |
| ENSMUSG00000028583 | 6135.05 | 6047.55 | 5627.28 | 5585.08 | 6336.52 | 5775.57 | 6446.70 | 6809.32 | 6594.22 | 6236.25 | 5877.05 | 6044.04 |
| ENSMUSG00000027506 | 727.39 | 701.95 | 740.90 | 783.54 | 862.94 | 790.19 | 797.63 | 764.27 | 753.85 | 737.78 | 679.25 | 702.64 |
| ENSMUSG00000027811 | 195.99 | 212.06 | 203.84 | 180.90 | 261.37 | 193.11 | 202.33 | 217.19 | 213.00 | 184.72 | 194.79 | 199.68 |
| ENSMUSG00000031737 | 479.73 | 430.99 | 471.39 | 435.19 | 527.05 | 478.08 | 491.89 | 468.93 | 461.34 | 453.51 | 426.93 | 421.96 |
| ENSMUSG00000000916 | 340.29 | 331.83 | 345.95 | 331.82 | 282.97 | 307.93 | 338.12 | 357.04 | 333.74 | 318.56 | 305.81 | 320.24 |
| ENSMUSG00000026068 | 145.28 | 143.33 | 175.42 | 139.55 | 157.68 | 144.05 | 142.08 | 136.56 | 141.35 | 128.31 | 130.20 | 128.10 |
| ENSMUSG00000038975 | 1464.53 | 1532.50 | 1581.75 | 1424.44 | 1587.64 | 1436.32 | 1507.14 | 1611.63 | 1550.89 | 1440.16 | 1400.88 | 1459.91 |
| ENSMUSG00000098650 | 1283.17 | 1296.88 | 1348.51 | 1214.60 | 1398.64 | 1263.05 | 1224.77 | 1255.41 | 1165.13 | 1099.47 | 1112.23 | 1146.27 |
| ENSMUSG00000082691 | 807.35 | 807.98 | 735.02 | 860.04 | 750.62 | 707.72 | 963.99 | 909.88 | 975.69 | 876.04 | 852.84 | 898.55 |
| ENSMUSG00000019189 | 1594.22 | 1728.85 | 1607.23 | 1688.03 | 1424.56 | 1558.45 | 1798.49 | 1749.02 | 1730.52 | 1650.32 | 1598.70 | 1620.98 |
| ENSMUSG00000046417 | 399.77 | 307.29 | 346.93 | 333.88 | 299.17 | 326.72 | 362.40 | 341.41 | 359.26 | 313.03 | 333.06 | 335.31 |
| ENSMUSG00000038503 | 2205.58 | 2204.02 | 2223.67 | 2142.86 | 2133.06 | 2049.06 | 2386.60 | 2428.55 | 2317.50 | 2150.28 | 2194.18 | 2249.21 |
| ENSMUSG00000004394 | 1813.60 | 1705.29 | 1726.80 | 1647.72 | 1743.17 | 1686.85 | 1878.53 | 1896.28 | 1837.51 | 1760.93 | 1665.32 | 1768.85 |
| ENSMUSG00000048039 | 786.87 | 749.07 | 739.92 | 697.75 | 871.58 | 866.39 | 712.20 | 726.43 | 697.90 | 651.50 | 644.93 | 681.92 |
| ENSMUSG00000040007 | 521.66 | 464.37 | 479.23 | 525.12 | 382.33 | 424.84 | 510.77 | 505.95 | 486.86 | 475.63 | 470.33 | 447.39 |

| | | | | | | | | | | | | |
|---|---|---|---|---|---|---|---|---|---|---|---|---|
| ENSMUSG00000021981 | 1119.36 | 1057.34 | 1125.06 | 1021.30 | 1334.91 | 1272.44 | 1059.31 | 1026.70 | 1090.53 | 977.80 | 1004.24 | 966.37 |
| ENSMUSG00000095325 | 143.33 | 99.16 | 154.84 | 133.35 | 150.12 | 150.31 | 120.50 | 117.64 | 118.77 | 112.82 | 112.03 | 106.43 |
| ENSMUSG00000029234 | 1278.30 | 1181.04 | 1372.03 | 1128.80 | 1496.92 | 1381.00 | 1278.73 | 1237.31 | 1203.41 | 1191.28 | 1135.44 | 1128.37 |
| ENSMUSG00000060038 | 1346.55 | 1076.97 | 1161.32 | 1058.51 | 988.23 | 908.14 | 1144.74 | 1174.79 | 1148.44 | 1037.53 | 1102.14 | 1082.22 |
| ENSMUSG00000022364 | 549.93 | 643.04 | 592.91 | 643.99 | 673.94 | 739.04 | 690.62 | 684.47 | 714.59 | 658.14 | 635.85 | 649.90 |
| ENSMUSG00000058440 | 577.23 | 577.27 | 642.89 | 636.76 | 559.45 | 604.38 | 660.05 | 670.48 | 637.04 | 623.85 | 591.44 | 615.99 |
| ENSMUSG00000026317 | 693.27 | 615.55 | 588.01 | 621.25 | 609.14 | 612.73 | 699.61 | 679.53 | 677.29 | 634.91 | 625.75 | 653.66 |
| ENSMUSG00000026526 | 2389.86 | 2251.14 | 2331.47 | 2330.99 | 2312.34 | 2135.70 | 2468.43 | 2570.05 | 2479.46 | 2262.00 | 2342.54 | 2395.20 |
| ENSMUSG00000005687 | 1430.41 | 1591.41 | 1577.83 | 1485.43 | 1479.64 | 1513.57 | 1585.37 | 1651.12 | 1656.90 | 1495.46 | 1504.84 | 1558.81 |
| ENSMUSG00000025283 | 2097.35 | 2143.15 | 2194.27 | 2053.96 | 2168.70 | 2085.59 | 2180.67 | 2152.13 | 2165.35 | 2023.08 | 2042.79 | 1989.25 |
| ENSMUSG00000046152 | 273.02 | 259.18 | 254.81 | 232.58 | 241.93 | 277.66 | 261.68 | 264.90 | 253.25 | 243.34 | 239.20 | 245.83 |
| ENSMUSG00000037251 | 585.03 | 482.04 | 561.55 | 515.82 | 547.57 | 485.39 | 512.57 | 520.76 | 535.94 | 493.33 | 499.59 | 472.82 |
| ENSMUSG00000042524 | 3635.01 | 3265.28 | 3252.69 | 3423.61 | 2896.64 | 3099.16 | 3688.71 | 3824.64 | 3700.54 | 3454.38 | 3595.06 | 3426.56 |
| ENSMUSG00000024878 | 483.63 | 513.45 | 490.99 | 517.88 | 542.17 | 517.74 | 543.14 | 521.58 | 543.79 | 482.26 | 506.66 | 514.27 |
| ENSMUSG00000042472 | 579.18 | 588.07 | 575.27 | 547.86 | 604.82 | 543.84 | 558.43 | 566.00 | 556.55 | 530.93 | 510.70 | 529.34 |
| ENSMUSG00000035949 | 1184.69 | 1079.92 | 1246.59 | 1071.95 | 1297.11 | 1228.60 | 1216.68 | 1228.26 | 1190.65 | 1148.14 | 1098.10 | 1151.92 |
| ENSMUSG00000028271 | 928.25 | 880.63 | 920.24 | 821.79 | 1028.19 | 835.07 | 870.47 | 908.24 | 867.71 | 819.63 | 839.72 | 815.67 |
| ENSMUSG00000094518 | 2236.78 | 1893.79 | 2032.56 | 1738.68 | 2092.01 | 1893.53 | 2008.02 | 1956.33 | 1899.35 | 1793.00 | 1811.66 | 1880.93 |
| ENSMUSG00000071359 | 604.53 | 700.97 | 667.39 | 596.44 | 758.18 | 721.29 | 633.07 | 659.79 | 636.06 | 599.51 | 587.40 | 617.87 |
| ENSMUSG00000031253 | 4026.98 | 3884.76 | 3702.52 | 3776.10 | 3304.89 | 3591.85 | 4406.31 | 4269.71 | 4309.11 | 3923.37 | 4174.39 | 4058.56 |
| ENSMUSG00000030204 | 1379.70 | 1381.31 | 1303.43 | 1342.78 | 1278.75 | 1233.82 | 1375.85 | 1410.90 | 1353.59 | 1308.53 | 1290.87 | 1295.09 |
| ENSMUSG00000020868 | 547.98 | 490.87 | 516.47 | 477.57 | 395.29 | 487.47 | 501.78 | 513.35 | 489.81 | 467.88 | 485.46 | 462.46 |
| ENSMUSG00000054237 | 362.72 | 378.95 | 362.61 | 312.18 | 398.53 | 358.04 | 350.71 | 334.01 | 342.57 | 316.35 | 318.93 | 332.48 |
| ENSMUSG00000062352 | 789.80 | 731.40 | 715.42 | 753.57 | 699.86 | 651.36 | 789.54 | 756.04 | 771.52 | 723.40 | 717.60 | 744.09 |
| ENSMUSG00000025373 | 929.23 | 893.39 | 934.94 | 895.18 | 853.22 | 865.34 | 990.97 | 1007.78 | 1004.15 | 935.77 | 942.67 | 956.95 |
| ENSMUSG00000019970 | 934.10 | 834.48 | 956.50 | 906.55 | 1083.27 | 992.69 | 896.55 | 912.35 | 905.99 | 839.54 | 871.01 | 854.29 |
| ENSMUSG00000090213 | 1895.51 | 1495.20 | 1661.14 | 1577.43 | 1458.04 | 1562.63 | 1926.19 | 1987.59 | 1903.27 | 1822.87 | 1848.00 | 1829.13 |
| ENSMUSG00000037824 | 1349.48 | 1300.81 | 1294.61 | 1322.10 | 1248.51 | 1161.79 | 1516.13 | 1502.21 | 1455.68 | 1424.67 | 1370.60 | 1439.19 |
| ENSMUSG00000030291 | 1592.27 | 1588.46 | 1659.18 | 1566.05 | 1858.73 | 1573.07 | 1623.14 | 1647.00 | 1598.99 | 1515.37 | 1523.01 | 1572.94 |
| ENSMUSG00000040599 | 911.68 | 1015.12 | 1023.14 | 962.37 | 1128.63 | 1075.16 | 963.99 | 993.80 | 988.45 | 920.28 | 950.74 | 919.28 |
| ENSMUSG00000029131 | 3487.78 | 3626.57 | 3679.98 | 3454.62 | 3808.18 | 3698.33 | 3817.30 | 3748.95 | 3882.13 | 3640.21 | 3680.85 | 3546.18 |
| ENSMUSG00000038206 | 584.06 | 556.65 | 590.95 | 417.61 | 754.94 | 630.48 | 546.74 | 524.87 | 548.70 | 506.60 | 521.80 | 510.50 |
| ENSMUSG00000029063 | 2496.14 | 2377.79 | 2380.47 | 2365.10 | 2367.42 | 2372.65 | 2348.83 | 2407.16 | 2417.62 | 2245.40 | 2297.13 | 2270.87 |
| ENSMUSG00000007659 | 1262.70 | 1007.27 | 1124.08 | 1062.64 | 964.47 | 993.74 | 1137.55 | 1182.19 | 1147.46 | 1103.90 | 1095.07 | 1094.46 |

| | | | | | | | | | | | | |
|---|---|---|---|---|---|---|---|---|---|---|---|---|
| ENSMUSG00000025089 | 741.04 | 699.98 | 714.44 | 671.90 | 699.86 | 781.84 | 671.74 | 682.00 | 689.07 | 649.29 | 631.81 | 662.14 |
| ENSMUSG00000021665 | 1820.43 | 1618.90 | 1775.80 | 1534.01 | 1898.69 | 1818.37 | 1536.81 | 1523.60 | 1532.24 | 1461.17 | 1468.51 | 1439.19 |
| ENSMUSG00000022814 | 953.61 | 836.45 | 886.92 | 860.04 | 862.94 | 846.55 | 912.73 | 922.22 | 901.09 | 882.68 | 856.88 | 866.53 |
| ENSMUSG00000028057 | 1107.66 | 1029.85 | 1045.68 | 1033.70 | 916.94 | 905.01 | 1157.33 | 1158.33 | 1159.24 | 1100.58 | 1111.22 | 1102.00 |
| ENSMUSG00000032802 | 936.05 | 861.97 | 902.60 | 880.71 | 776.54 | 828.81 | 831.80 | 814.45 | 847.10 | 786.44 | 791.28 | 805.31 |
| ENSMUSG00000042901 | 2767.21 | 2958.00 | 3125.29 | 2884.02 | 3631.05 | 3211.90 | 2915.36 | 2830.02 | 2877.00 | 2791.82 | 2692.76 | 2772.90 |
| ENSMUSG00000030663 | 1613.72 | 1835.86 | 1676.82 | 1702.50 | 1698.88 | 1641.96 | 1829.07 | 1863.37 | 1815.91 | 1774.20 | 1743.03 | 1759.43 |
| ENSMUSG00000025228 | 4568.14 | 3979.99 | 4092.57 | 4094.48 | 3593.25 | 3638.83 | 4434.18 | 4421.08 | 4302.24 | 4191.05 | 4248.07 | 4245.05 |
| ENSMUSG00000055239 | 2590.72 | 2425.89 | 2322.65 | 2393.01 | 2382.54 | 2388.31 | 2751.69 | 2783.12 | 2717.00 | 2686.74 | 2664.50 | 2636.32 |
| ENSMUSG00000022051 | 2877.39 | 2824.48 | 2909.68 | 2803.39 | 3612.69 | 3339.24 | 2914.46 | 2995.38 | 2960.43 | 2840.49 | 2887.56 | 2858.61 |
| ENSMUSG00000030894 | 1824.33 | 1938.95 | 1835.58 | 1780.03 | 2000.21 | 2078.29 | 2070.06 | 2095.36 | 2093.70 | 2045.20 | 2011.50 | 2013.74 |
| ENSMUSG00000021901 | 1297.80 | 1236.02 | 1274.03 | 1292.12 | 993.63 | 1132.57 | 1319.19 | 1341.79 | 1310.40 | 1285.30 | 1266.65 | 1301.68 |
| ENSMUSG00000037058 | 3574.56 | 3874.95 | 3930.86 | 3671.70 | 4226.15 | 3921.71 | 4055.60 | 4051.70 | 4065.68 | 3991.95 | 3914.00 | 3916.34 |
| ENSMUSG00000029176 | 2148.05 | 2329.68 | 2319.71 | 2133.56 | 2481.90 | 2247.39 | 2209.45 | 2248.38 | 2210.51 | 2171.29 | 2165.92 | 2185.16 |
| ENSMUSG00000019087 | 4491.11 | 3974.10 | 4218.01 | 3805.05 | 3863.26 | 3859.08 | 4214.77 | 4190.73 | 4264.94 | 4102.56 | 4156.22 | 4144.27 |
| ENSMUSG00000031078 | 4688.07 | 4630.89 | 4712.92 | 4927.65 | 3668.86 | 4291.23 | 4809.17 | 4917.15 | 4829.35 | 4952.06 | 4975.76 | 4999.50 |
| ENSMUSG00000024059 | 566.51 | 611.63 | 599.77 | 569.57 | 716.06 | 657.62 | 659.15 | 667.19 | 655.69 | 675.83 | 672.18 | 688.51 |
| ENSMUSG00000102135 | 2445.44 | 2441.60 | 2201.13 | 2604.92 | 2443.02 | 1918.58 | 2685.15 | 2727.18 | 2737.61 | 2803.99 | 2797.73 | 2830.35 |
| ENSMUSG00000038991 | 7448.45 | 6644.45 | 6946.39 | 6618.78 | 5974.71 | 6176.40 | 6310.01 | 6313.24 | 6268.34 | 6549.28 | 6547.21 | 6502.74 |
| ENSMUSG00000082878 | 329.57 | 377.97 | 353.79 | 365.93 | 366.13 | 335.07 | 365.99 | 372.67 | 377.91 | 381.61 | 390.59 | 388.05 |
| ENSMUSG00000024726 | 761.52 | 725.51 | 844.78 | 668.80 | 1204.23 | 911.27 | 664.54 | 653.21 | 643.91 | 680.26 | 673.19 | 685.69 |
| ENSMUSG00000021253 | 5738.21 | 5220.92 | 5095.13 | 4954.52 | 4867.69 | 5036.53 | 5556.44 | 5631.24 | 5441.85 | 5704.21 | 5743.82 | 5847.19 |
| ENSMUSG00000031575 | 1550.34 | 1395.06 | 1551.37 | 1330.37 | 1499.08 | 1479.12 | 1383.94 | 1377.17 | 1359.48 | 1413.61 | 1435.20 | 1440.13 |
| ENSMUSG00000002948 | 1230.52 | 1206.56 | 1164.26 | 1341.74 | 1069.23 | 1236.95 | 1290.42 | 1264.46 | 1310.40 | 1342.82 | 1341.34 | 1350.66 |
| ENSMUSG00000025225 | 1586.42 | 1403.90 | 1385.75 | 1509.20 | 1108.11 | 1284.97 | 1351.57 | 1399.38 | 1397.76 | 1437.94 | 1455.38 | 1440.13 |
| ENSMUSG00000038335 | 1648.82 | 1669.95 | 1618.99 | 1553.65 | 1664.32 | 1607.51 | 1509.83 | 1539.23 | 1547.94 | 1590.59 | 1627.97 | 1583.30 |
| ENSMUSG00000024617 | 1439.18 | 1398.99 | 1351.45 | 1291.09 | 1213.95 | 1441.54 | 1718.46 | 1748.19 | 1734.44 | 1796.32 | 1830.84 | 1809.35 |
| ENSMUSG00000032185 | 2652.15 | 2294.34 | 2328.53 | 2334.09 | 2048.81 | 2323.59 | 2316.46 | 2357.80 | 2404.86 | 2434.55 | 2474.76 | 2495.98 |
| ENSMUSG00000035372 | 1459.66 | 1299.83 | 1333.81 | 1352.08 | 1118.91 | 1179.54 | 1413.62 | 1444.63 | 1426.23 | 1500.99 | 1501.81 | 1479.69 |
| ENSMUSG00000025130 | 31745.88 | 29066.52 | 30467.86 | 29550.37 | 26912.14 | 28624.19 | 29203.02 | 28424.40 | 28648.25 | 29738.88 | 30466.18 | 30114.73 |
| ENSMUSG00000065979 | 1385.55 | 1272.34 | 1315.19 | 1183.59 | 1421.32 | 1424.84 | 1220.28 | 1222.50 | 1224.02 | 1283.09 | 1293.90 | 1263.06 |
| ENSMUSG00000024271 | 2068.09 | 2105.84 | 2039.42 | 2159.40 | 2206.50 | 2103.34 | 2061.07 | 2116.75 | 2082.90 | 2156.91 | 2192.16 | 2212.48 |
| ENSMUSG00000038299 | 2212.40 | 2156.89 | 2244.25 | 2068.43 | 2391.18 | 2331.94 | 2034.09 | 2010.63 | 1998.49 | 2164.66 | 2097.29 | 2087.21 |
| ENSMUSG00000062937 | 2156.82 | 2235.43 | 2156.05 | 2277.24 | 2374.98 | 2251.56 | 2027.80 | 2032.02 | 2028.92 | 2118.20 | 2186.10 | 2108.87 |

| | | | | | | | | | | | | |
|---|---|---|---|---|---|---|---|---|---|---|---|---|
| ENSMUSG00000027187 | 2011.54 | 1980.18 | 2081.56 | 1867.89 | 2057.45 | 1991.65 | 1982.84 | 1986.77 | 1987.69 | 2069.53 | 2115.45 | 2095.68 |
| ENSMUSG00000016619 | 1702.45 | 1669.95 | 1601.35 | 1734.55 | 1599.52 | 1665.97 | 1738.24 | 1668.39 | 1672.60 | 1780.84 | 1797.53 | 1782.98 |
| ENSMUSG00000019066 | 561.63 | 508.54 | 555.67 | 568.53 | 490.33 | 489.56 | 483.79 | 481.27 | 491.77 | 516.55 | 518.77 | 502.96 |
| ENSMUSG00000036057 | 1679.05 | 1841.75 | 1669.96 | 1891.67 | 1392.16 | 1844.47 | 1758.03 | 1822.23 | 1831.62 | 1912.46 | 1946.90 | 1858.33 |
| ENSMUSG00000107951 | 21504.87 | 21340.19 | 21736.86 | 23371.94 | 22372.78 | 22060.52 | 24997.25 | 24151.40 | 24772.01 | 25363.11 | 26735.88 | 26089.14 |
| ENSMUSG00000051391 | 6222.81 | 6553.15 | 6344.65 | 6517.48 | 6190.72 | 6335.07 | 6846.86 | 6771.47 | 7032.00 | 7258.30 | 7334.45 | 7270.37 |
| ENSMUSG00000030816 | 1456.73 | 1256.63 | 1378.89 | 1355.18 | 1062.75 | 1305.84 | 1400.13 | 1346.73 | 1356.54 | 1449.00 | 1475.57 | 1425.06 |
| ENSMUSG00000038776 | 3583.33 | 2969.78 | 3319.33 | 2880.92 | 2879.35 | 2907.09 | 2779.57 | 2891.72 | 2876.02 | 3032.95 | 3003.62 | 3029.09 |
| ENSMUSG00000035914 | 1795.08 | 1630.68 | 1647.41 | 1504.03 | 1425.64 | 1649.27 | 1568.29 | 1567.20 | 1570.52 | 1693.45 | 1670.36 | 1631.34 |
| ENSMUSG00000026455 | 638.66 | 667.59 | 663.47 | 650.20 | 706.34 | 699.37 | 680.73 | 655.68 | 671.40 | 709.02 | 706.50 | 715.83 |
| ENSMUSG00000107713 | 1855.53 | 1999.81 | 1848.32 | 2086.01 | 2263.74 | 1617.95 | 2072.76 | 2072.33 | 2035.79 | 2185.67 | 2145.73 | 2233.20 |
| ENSMUSG00000001424 | 5024.46 | 4649.54 | 4350.31 | 4545.18 | 3948.58 | 4301.67 | 4665.29 | 4573.27 | 4574.14 | 4771.76 | 4937.41 | 4977.84 |
| ENSMUSG00000032419 | 1119.36 | 1183.98 | 1187.79 | 1052.31 | 1477.48 | 1289.14 | 981.08 | 986.39 | 997.28 | 1077.35 | 1048.64 | 1028.53 |
| ENSMUSG00000093798 | 19785.84 | 19487.63 | 19553.37 | 21590.88 | 19446.98 | 19157.60 | 23197.85 | 23217.66 | 23387.01 | 24086.66 | 25053.41 | 25146.32 |
| ENSMUSG00000022505 | 5578.30 | 5076.60 | 5165.69 | 5012.41 | 4618.20 | 5042.79 | 4749.82 | 4725.47 | 4884.32 | 5059.35 | 5151.37 | 5076.73 |
| ENSMUSG00000005973 | 5338.43 | 5508.57 | 5591.02 | 5328.72 | 5349.38 | 5373.69 | 5087.04 | 5034.80 | 4848.98 | 5294.95 | 5467.28 | 5179.40 |
| ENSMUSG00000083863 | 62801.42 | 58126.16 | 59412.78 | 60259.51 | 86372.00 | 70817.25 | 59984.22 | 57831.89 | 59639.52 | 64699.71 | 62907.53 | 61405.88 |
| ENSMUSG00000000826 | 2926.14 | 2855.90 | 2837.16 | 2873.68 | 2620.15 | 2849.68 | 2900.97 | 2746.93 | 2838.72 | 2926.77 | 3058.12 | 3055.46 |
| ENSMUSG00000074412 | 1476.24 | 1512.87 | 1361.25 | 1728.35 | 1651.36 | 1270.35 | 1636.63 | 1663.46 | 1578.37 | 1722.21 | 1716.79 | 1758.49 |
| ENSMUSG00000031148 | 958.48 | 986.65 | 1030.00 | 910.69 | 1101.63 | 1030.27 | 934.32 | 891.78 | 925.63 | 983.33 | 971.94 | 980.50 |
| ENSMUSG00000027833 | 1133.02 | 1147.66 | 1297.55 | 1110.19 | 1409.44 | 1374.74 | 1072.80 | 1048.09 | 1084.64 | 1151.46 | 1139.48 | 1131.20 |
| ENSMUSG00000039166 | 334.44 | 376.01 | 396.91 | 364.90 | 426.61 | 423.80 | 357.90 | 343.88 | 343.55 | 365.02 | 382.52 | 369.22 |
| ENSMUSG00000060594 | 893.15 | 854.12 | 867.32 | 861.07 | 817.58 | 889.35 | 839.00 | 793.89 | 800.97 | 888.21 | 856.88 | 855.23 |
| ENSMUSG00000032965 | 1065.74 | 1178.09 | 1116.24 | 1053.34 | 1145.91 | 1126.30 | 1138.45 | 1081.82 | 1095.44 | 1197.92 | 1205.08 | 1140.62 |
| ENSMUSG00000032468 | 862.92 | 1060.28 | 981.00 | 946.87 | 1069.23 | 1097.08 | 955.90 | 909.88 | 960.96 | 1015.41 | 1000.20 | 1008.75 |
| ENSMUSG00000062861 | 185.26 | 202.24 | 179.34 | 220.18 | 181.44 | 192.07 | 201.43 | 197.44 | 196.32 | 206.84 | 219.01 | 210.98 |
| ENSMUSG00000038290 | 3399.05 | 3160.24 | 3188.99 | 3248.92 | 2822.11 | 3218.16 | 3045.75 | 3022.52 | 3056.62 | 3248.65 | 3297.32 | 3218.40 |
| ENSMUSG00000033955 | 3577.48 | 3330.08 | 3298.75 | 3462.89 | 2599.63 | 3176.41 | 3249.88 | 3263.57 | 3268.65 | 3494.20 | 3546.62 | 3428.44 |
| ENSMUSG00000005103 | 5576.35 | 5587.11 | 5263.69 | 5372.14 | 5359.10 | 5493.73 | 5258.79 | 5614.79 | 5393.76 | 5881.19 | 5669.14 | 5862.26 |
| ENSMUSG00000051339 | 1478.19 | 1317.50 | 1309.31 | 1292.12 | 1294.95 | 1427.97 | 1401.03 | 1397.73 | 1424.27 | 1502.10 | 1545.21 | 1474.98 |
| ENSMUSG00000000194 | 1495.74 | 1534.47 | 1417.11 | 1521.61 | 1289.55 | 1441.54 | 1517.93 | 1514.55 | 1479.23 | 1582.84 | 1622.93 | 1626.63 |
| ENSMUSG00000069125 | 887.30 | 982.73 | 894.76 | 1138.10 | 1043.31 | 778.70 | 1077.30 | 1148.46 | 1124.89 | 1170.26 | 1188.93 | 1235.75 |
| ENSMUSG00000034863 | 707.89 | 631.26 | 636.03 | 651.23 | 527.05 | 604.38 | 677.13 | 662.26 | 660.60 | 731.14 | 714.57 | 700.76 |
| ENSMUSG00000031503 | 4124.49 | 3691.36 | 3515.33 | 3986.98 | 3312.45 | 3705.63 | 3802.01 | 3797.49 | 3765.32 | 4071.59 | 4028.04 | 4100.00 |

| | | | | | | | | | | | | |
|---|---|---|---|---|---|---|---|---|---|---|---|---|
| ENSMUSG00000083362 | 133.58 | 167.88 | 124.46 | 220.18 | 151.20 | 136.74 | 179.85 | 184.28 | 187.48 | 199.10 | 190.75 | 202.50 |
| ENSMUSG00000084166 | 201.84 | 271.94 | 199.92 | 285.30 | 250.57 | 243.21 | 282.36 | 273.13 | 286.62 | 309.71 | 292.69 | 302.34 |
| ENSMUSG00000081151 | 29.25 | 36.32 | 32.34 | 42.38 | 31.32 | 34.45 | 41.37 | 41.96 | 40.24 | 43.14 | 45.42 | 44.27 |
| ENSMUSG00000080763 | 195.01 | 177.70 | 136.22 | 203.64 | 203.05 | 146.14 | 201.43 | 213.90 | 211.04 | 223.43 | 226.08 | 224.17 |
| ENSMUSG00000033880 | 1035.51 | 927.75 | 937.88 | 932.40 | 777.62 | 941.54 | 869.57 | 901.66 | 866.73 | 976.70 | 942.67 | 920.22 |
| ENSMUSG00000040463 | 4163.49 | 3871.02 | 3707.42 | 4104.82 | 3062.96 | 3618.99 | 3685.11 | 3795.84 | 3820.29 | 4001.91 | 4157.23 | 4009.58 |
| ENSMUSG00000081642 | 352.97 | 340.67 | 325.37 | 395.91 | 483.85 | 384.13 | 414.55 | 428.62 | 434.84 | 468.99 | 454.18 | 453.04 |
| ENSMUSG00000022965 | 1120.34 | 919.89 | 1037.84 | 925.16 | 1038.99 | 977.03 | 865.97 | 847.36 | 850.04 | 924.71 | 931.57 | 907.97 |
| ENSMUSG00000026155 | 1707.32 | 1556.07 | 1643.49 | 1636.35 | 2001.29 | 1782.88 | 1502.64 | 1514.55 | 1563.65 | 1631.51 | 1688.53 | 1620.03 |
| ENSMUSG00000100862 | ####### | ######## | ######### | ######### | ######### | ######## | ######## | ######### | ######## | ######## | ######## | ######## |
| ENSMUSG00000058809 | 1457.71 | 1780.89 | 1699.36 | 1797.60 | 1764.77 | 1726.51 | 1673.50 | 1609.98 | 1681.44 | 1776.41 | 1819.74 | 1762.26 |
| ENSMUSG00000064357 | ######### | ######## | ######### | ######### | ######### | ######## | ######## | ######### | ######## | ######## | ######## | ######## |
| ENSMUSG00000081603 | 1003.33 | 1194.78 | 1035.88 | 1070.91 | 1317.63 | 1070.98 | 919.93 | 946.90 | 986.48 | 1019.83 | 1024.42 | 1038.89 |
| ENSMUSG00000048720 | 293.49 | 328.88 | 343.99 | 250.16 | 411.49 | 407.10 | 282.36 | 269.84 | 266.01 | 304.18 | 292.69 | 287.27 |
| ENSMUSG00000105854 | 621.11 | 732.38 | 563.51 | 786.65 | 799.22 | 584.55 | 762.56 | 735.48 | 780.35 | 815.20 | 813.48 | 833.56 |
| ENSMUSG00000004508 | 642.56 | 636.17 | 599.77 | 646.06 | 560.53 | 664.93 | 639.36 | 623.59 | 654.71 | 679.15 | 709.53 | 685.69 |
| ENSMUSG00000028433 | 3255.71 | 3285.90 | 3055.70 | 3347.12 | 2735.71 | 3490.60 | 3209.41 | 3302.24 | 3392.32 | 3642.42 | 3544.60 | 3539.59 |
| ENSMUSG00000038351 | 705.94 | 650.90 | 678.17 | 654.33 | 561.61 | 612.73 | 632.17 | 604.67 | 647.84 | 670.30 | 678.24 | 693.22 |
| ENSMUSG00000031502 | 4947.44 | 4662.31 | 4539.46 | 4972.09 | 3853.54 | 4504.17 | 4308.29 | 4361.85 | 4381.75 | 4685.48 | 4803.17 | 4653.83 |
| ENSMUSG00000025047 | 1046.24 | 1055.38 | 1054.50 | 1118.46 | 911.54 | 1201.46 | 1069.20 | 1013.54 | 1090.53 | 1127.13 | 1155.63 | 1163.22 |
| ENSMUSG00000039763 | 141.38 | 177.70 | 165.62 | 181.93 | 163.08 | 205.64 | 147.48 | 149.73 | 156.07 | 167.02 | 159.47 | 165.77 |
| ENSMUSG00000022594 | 2202.65 | 2027.30 | 1913.00 | 2039.49 | 1550.92 | 1742.17 | 1995.43 | 2099.48 | 2011.25 | 2232.13 | 2254.74 | 2144.66 |
| ENSMUSG00000100228 | 2300.16 | 2436.69 | 2199.17 | 2587.35 | 2812.39 | 2009.39 | 2496.31 | 2602.13 | 2506.94 | 2693.38 | 2720.01 | 2848.25 |
| ENSMUSG00000032898 | 650.36 | 675.44 | 638.97 | 715.32 | 618.86 | 693.11 | 702.31 | 657.32 | 688.08 | 723.40 | 767.05 | 734.67 |
| ENSMUSG00000030058 | 6292.04 | 6255.68 | 5839.94 | 6581.56 | 5448.74 | 5881.00 | 6385.55 | 6464.61 | 6896.55 | 6986.19 | 7310.23 | 7166.77 |
| ENSMUSG00000031375 | ######### | 94140.49 | 93956.55 | 92645.31 | 88027.68 | 97288.00 | 89954.29 | 89472.14 | 88379.06 | 97287.93 | 97050.53 | 96769.71 |
| ENSMUSG00000025867 | 587.96 | 500.69 | 538.03 | 640.89 | 632.90 | 650.31 | 552.14 | 580.81 | 578.15 | 605.04 | 641.90 | 613.16 |
| ENSMUSG00000036158 | 468.03 | 467.31 | 479.23 | 399.01 | 470.89 | 441.54 | 440.63 | 452.47 | 427.97 | 481.16 | 470.33 | 486.01 |
| ENSMUSG00000052488 | 1919.89 | 1920.29 | 1796.38 | 1866.86 | 1677.28 | 1970.77 | 1748.13 | 1847.74 | 1855.18 | 1954.50 | 1985.26 | 1995.84 |
| ENSMUSG00000038241 | 1175.92 | 1224.24 | 1223.07 | 1454.42 | 959.06 | 1276.62 | 1246.36 | 1262.81 | 1262.31 | 1421.35 | 1348.40 | 1341.24 |
| ENSMUSG00000069874 | 88.73 | 59.89 | 69.58 | 83.73 | 88.56 | 103.34 | 71.94 | 70.75 | 69.69 | 79.64 | 74.69 | 77.23 |
| ENSMUSG00000045248 | 266.19 | 272.93 | 236.18 | 253.26 | 211.69 | 247.39 | 241.00 | 246.80 | 255.21 | 280.95 | 265.44 | 263.73 |
| ENSMUSG00000031066 | 932.15 | 883.57 | 953.56 | 998.55 | 848.90 | 939.46 | 919.93 | 901.66 | 942.31 | 1034.21 | 1009.28 | 970.14 |
| ENSMUSG00000035392 | 513.85 | 576.28 | 571.35 | 557.16 | 460.09 | 627.35 | 530.56 | 513.35 | 522.20 | 575.18 | 583.36 | 549.12 |

| | | | | | | | | | | | | |
|---|---|---|---|---|---|---|---|---|---|---|---|---|
| ENSMUSG00000015016 | 135.53 | 145.30 | 137.20 | 128.18 | 106.92 | 105.43 | 116.90 | 120.93 | 125.64 | 137.16 | 132.22 | 127.15 |
| ENSMUSG00000046818 | 787.85 | 705.87 | 806.56 | 702.92 | 760.34 | 792.27 | 643.86 | 603.02 | 640.97 | 702.38 | 682.27 | 678.15 |
| ENSMUSG00000103823 | 3840.75 | 4208.74 | 3969.08 | 4794.30 | 4594.44 | 4014.61 | 5120.31 | 4685.16 | 4963.83 | 5255.13 | 5512.70 | 5376.25 |
| ENSMUSG00000037017 | 545.06 | 504.62 | 536.07 | 509.61 | 501.13 | 495.82 | 465.81 | 506.77 | 478.03 | 529.83 | 544.00 | 512.38 |
| ENSMUSG00000048170 | 1021.86 | 1098.57 | 1103.50 | 1065.74 | 1154.55 | 1203.55 | 1044.02 | 1115.55 | 1077.77 | 1206.77 | 1155.63 | 1178.29 |
| ENSMUSG00000034471 | 558.71 | 504.62 | 478.25 | 596.44 | 361.81 | 479.12 | 485.59 | 507.59 | 468.21 | 546.42 | 515.74 | 536.87 |
| ENSMUSG00000107639 | 184.29 | 214.02 | 203.84 | 229.48 | 227.89 | 198.33 | 225.71 | 221.30 | 211.04 | 247.77 | 232.13 | 240.18 |
| ENSMUSG00000041577 | 3226.46 | 2807.79 | 2658.80 | 3195.16 | 2741.11 | 2974.94 | 3126.68 | 2970.70 | 2855.40 | 3184.49 | 3311.45 | 3302.23 |
| ENSMUSG00000105263 | 825.87 | 747.11 | 800.68 | 690.51 | 658.82 | 811.06 | 578.22 | 599.73 | 595.82 | 663.67 | 654.01 | 625.41 |
| ENSMUSG00000018593 | ######## | ######## | ######## | ######## | ######## | 99204.49 | ######## | ######## | ######## | ######## | ######## | ######## |
| ENSMUSG00000037235 | 3696.44 | 3200.49 | 3613.34 | 3685.14 | 3168.80 | 3296.45 | 3004.38 | 3041.45 | 3195.03 | 3449.96 | 3435.60 | 3249.49 |
| ENSMUSG00000022842 | 566.51 | 459.46 | 481.19 | 497.21 | 413.65 | 456.16 | 461.31 | 436.84 | 434.84 | 492.22 | 492.53 | 478.48 |
| ENSMUSG00000000976 | 832.70 | 809.94 | 795.78 | 840.40 | 852.14 | 931.11 | 811.12 | 794.71 | 848.08 | 868.30 | 897.25 | 929.64 |
| ENSMUSG00000040124 | 603.56 | 632.24 | 717.38 | 670.87 | 758.18 | 621.08 | 561.13 | 519.93 | 552.63 | 597.30 | 608.60 | 591.50 |
| ENSMUSG00000029471 | 624.04 | 559.59 | 593.89 | 651.23 | 617.78 | 613.78 | 498.18 | 481.27 | 524.16 | 547.52 | 564.19 | 543.46 |
| ENSMUSG00000026043 | ######## | ######## | ######## | ######## | ######## | ######## | ######## | ######## | ######## | ######## | ######## | ######## |
| ENSMUSG00000081111 | 432.92 | 436.88 | 400.83 | 455.86 | 407.17 | 366.39 | 450.52 | 426.97 | 436.80 | 485.58 | 467.30 | 495.43 |
| ENSMUSG00000009733 | 461.20 | 450.62 | 462.57 | 458.96 | 509.77 | 485.39 | 430.74 | 406.40 | 433.86 | 485.58 | 450.14 | 465.29 |
| ENSMUSG00000102546 | 255.46 | 288.63 | 252.85 | 318.38 | 267.85 | 278.71 | 301.25 | 277.24 | 288.58 | 305.29 | 327.01 | 324.01 |
| ENSMUSG00000031380 | 387.10 | 334.77 | 396.91 | 332.85 | 359.65 | 365.34 | 294.05 | 283.00 | 301.34 | 332.94 | 330.04 | 306.11 |
| ENSMUSG00000037664 | 93.61 | 73.63 | 117.60 | 81.66 | 95.04 | 92.90 | 86.33 | 82.27 | 79.51 | 88.49 | 94.87 | 90.42 |
| ENSMUSG00000020456 | 4778.75 | 4359.93 | 4301.31 | 4544.14 | 3850.30 | 4518.78 | 4673.38 | 4643.20 | 4946.16 | 5095.85 | 5348.18 | 5303.73 |
| ENSMUSG00000002006 | 87.76 | 85.41 | 83.30 | 93.03 | 92.88 | 85.59 | 98.92 | 94.61 | 97.18 | 107.29 | 111.02 | 102.66 |
| ENSMUSG00000074364 | 7975.96 | 6829.02 | 7152.19 | 7473.65 | 6737.21 | 7639.87 | 6935.89 | 6836.47 | 6707.10 | 7653.18 | 7640.26 | 7320.29 |
| ENSMUSG00000101219 | 496.30 | 432.95 | 425.33 | 512.71 | 460.09 | 347.60 | 484.69 | 464.81 | 497.66 | 525.40 | 552.08 | 521.80 |
| ENSMUSG00000036545 | 3987.98 | 3680.56 | 3671.16 | 3754.40 | 3095.36 | 3288.10 | 3531.34 | 3268.51 | 3460.05 | 3629.15 | 3895.83 | 3814.61 |
| ENSMUSG00000002409 | 511.90 | 375.03 | 430.23 | 426.92 | 298.09 | 415.45 | 427.14 | 415.45 | 402.45 | 450.19 | 483.45 | 443.63 |
| ENSMUSG00000006906 | 618.19 | 709.80 | 692.87 | 648.13 | 690.14 | 630.48 | 604.29 | 553.66 | 580.11 | 638.23 | 631.81 | 652.72 |
| ENSMUSG00000028468 | 992.61 | 1048.50 | 920.24 | 1010.96 | 903.98 | 1013.57 | 919.03 | 914.82 | 959.98 | 1054.12 | 1010.29 | 1028.53 |
| ENSMUSG00000074182 | 1109.61 | 1172.20 | 1207.39 | 1120.53 | 1103.79 | 1113.78 | 1004.46 | 1034.11 | 1053.23 | 1154.78 | 1170.77 | 1099.17 |
| ENSMUSG00000022479 | 1219.80 | 1286.09 | 1270.11 | 1251.81 | 1196.67 | 1212.94 | 1067.41 | 1085.11 | 1069.92 | 1201.24 | 1201.05 | 1173.58 |
| ENSMUSG00000097789 | 1222.72 | 1170.24 | 1183.87 | 1187.72 | 1382.44 | 1346.55 | 1035.93 | 1060.43 | 1008.08 | 1171.37 | 1106.17 | 1174.52 |
| ENSMUSG00000022545 | 433.90 | 412.33 | 448.85 | 440.36 | 392.05 | 455.11 | 429.84 | 419.57 | 422.08 | 478.95 | 474.36 | 460.58 |
| ENSMUSG00000026932 | 1350.45 | 1282.16 | 1341.65 | 1363.45 | 1458.04 | 1411.27 | 1166.32 | 1246.36 | 1238.75 | 1354.99 | 1362.53 | 1343.12 |

| | | | | | | | | | | | | |
|---|---|---|---|---|---|---|---|---|---|---|---|---|
| ENSMUSG00000037287 | 667.91 | 606.72 | 663.47 | 604.71 | 739.82 | 722.34 | 569.22 | 585.75 | 593.85 | 676.94 | 640.89 | 627.29 |
| ENSMUSG00000039787 | 739.09 | 613.59 | 625.25 | 698.78 | 579.98 | 597.08 | 665.44 | 628.53 | 610.54 | 682.47 | 728.70 | 708.29 |
| ENSMUSG00000029661 | 37570.87 | 33930.08 | 32411.24 | 34459.40 | 33762.76 | 33972.82 | 31825.22 | 29564.63 | 29225.42 | 33376.88 | 34348.89 | 33205.04 |
| ENSMUSG00000022575 | 442.68 | 419.21 | 397.89 | 423.82 | 332.65 | 376.83 | 346.21 | 370.21 | 346.50 | 386.03 | 409.77 | 389.94 |
| ENSMUSG00000041757 | 198.91 | 229.73 | 197.96 | 250.16 | 163.08 | 230.69 | 223.91 | 231.17 | 222.82 | 243.34 | 259.39 | 255.25 |
| ENSMUSG00000031700 | 1604.94 | 1450.04 | 1546.47 | 1364.48 | 1461.28 | 1451.98 | 1374.05 | 1281.73 | 1358.50 | 1437.94 | 1558.33 | 1498.53 |
| ENSMUSG00000027185 | 841.47 | 794.23 | 776.18 | 806.29 | 725.78 | 794.36 | 715.80 | 739.59 | 724.40 | 804.14 | 793.30 | 843.92 |
| ENSMUSG00000079484 | 309.09 | 284.71 | 341.05 | 297.71 | 334.81 | 310.02 | 258.08 | 251.74 | 248.34 | 269.89 | 285.63 | 293.87 |
| ENSMUSG00000017376 | 352.97 | 288.63 | 320.47 | 346.29 | 478.45 | 422.76 | 315.64 | 335.65 | 310.18 | 345.11 | 371.42 | 362.62 |
| ENSMUSG00000058756 | 1590.32 | 1318.48 | 1484.73 | 1546.41 | 1217.19 | 1320.46 | 1433.40 | 1323.69 | 1434.08 | 1589.48 | 1639.07 | 1479.69 |
| ENSMUSG00000047045 | 490.45 | 451.60 | 457.67 | 476.54 | 503.29 | 489.56 | 410.06 | 386.66 | 392.63 | 438.02 | 440.05 | 458.70 |
| ENSMUSG00000031295 | 724.47 | 745.14 | 731.10 | 782.51 | 665.30 | 766.18 | 723.89 | 675.42 | 726.37 | 799.72 | 815.50 | 776.11 |
| ENSMUSG00000066151 | 1290.00 | 1247.80 | 1277.95 | 1315.90 | 1041.15 | 1210.85 | 1259.84 | 1160.80 | 1280.96 | 1358.30 | 1418.04 | 1391.16 |
| ENSMUSG00000046768 | 862.92 | 845.28 | 862.42 | 852.80 | 1018.47 | 897.70 | 783.24 | 767.56 | 775.44 | 849.49 | 854.86 | 916.45 |
| ENSMUSG00000037306 | 1806.78 | 1558.03 | 1660.16 | 1633.25 | 1569.28 | 1546.97 | 1522.42 | 1565.56 | 1568.56 | 1741.02 | 1754.13 | 1751.90 |
| ENSMUSG00000024501 | 9928.02 | 10223.89 | 10224.56 | 10474.48 | 11527.14 | 11588.71 | 9468.16 | 8965.56 | 9950.23 | 10822.18 | 10650.95 | 10511.38 |
| ENSMUSG00000001119 | 28315.63 | 24542.64 | 24672.02 | 25078.58 | 19486.95 | 21657.60 | 24061.13 | 21545.97 | 22876.59 | 25509.11 | 26484.57 | 25228.26 |
| ENSMUSG00000036825 | 910.70 | 966.04 | 1011.38 | 907.59 | 1006.59 | 1021.92 | 919.03 | 879.44 | 958.02 | 984.44 | 1039.56 | 1088.81 |
| ENSMUSG00000037750 | 606.48 | 627.33 | 568.41 | 638.83 | 562.69 | 663.88 | 594.40 | 566.00 | 608.58 | 693.53 | 662.09 | 644.25 |
| ENSMUSG00000037552 | 777.12 | 782.45 | 676.21 | 851.77 | 686.90 | 822.55 | 696.02 | 780.72 | 722.44 | 849.49 | 816.51 | 823.20 |
| ENSMUSG00000024999 | 547.01 | 618.50 | 641.91 | 653.30 | 747.38 | 803.76 | 562.03 | 537.21 | 559.50 | 643.76 | 624.75 | 609.40 |
| ENSMUSG00000097987 | 1224.67 | 1246.82 | 1087.82 | 1296.26 | 1735.61 | 1110.65 | 1253.55 | 1166.56 | 1247.58 | 1331.76 | 1409.97 | 1412.82 |
| ENSMUSG00000031778 | 1506.46 | 1244.85 | 1308.33 | 1248.71 | 1167.51 | 1223.38 | 1035.03 | 1132.01 | 1135.68 | 1193.49 | 1323.17 | 1225.39 |
| ENSMUSG00000039987 | 400.75 | 484.00 | 450.81 | 397.97 | 497.89 | 491.65 | 404.66 | 392.42 | 417.17 | 452.40 | 474.36 | 452.10 |
| ENSMUSG00000032735 | 2339.16 | 2226.60 | 2048.24 | 2199.71 | 1825.25 | 2085.59 | 2362.32 | 2193.26 | 2288.05 | 2445.61 | 2582.75 | 2744.64 |
| ENSMUSG00000081619 | 151.13 | 199.29 | 239.13 | 203.64 | 245.17 | 211.90 | 194.24 | 174.41 | 192.39 | 216.80 | 204.88 | 215.69 |
| ENSMUSG00000073062 | 251.56 | 253.29 | 277.35 | 226.38 | 331.57 | 346.55 | 247.29 | 229.53 | 245.39 | 280.95 | 266.45 | 273.15 |
| ENSMUSG00000043631 | 305.19 | 328.88 | 355.75 | 305.98 | 396.37 | 360.12 | 284.16 | 252.56 | 280.73 | 296.44 | 316.91 | 317.41 |
| ENSMUSG00000047583 | 159.91 | 209.11 | 190.12 | 193.30 | 223.57 | 221.29 | 156.47 | 165.36 | 161.96 | 181.40 | 188.74 | 180.84 |
| ENSMUSG00000096623 | 514.83 | 619.48 | 494.91 | 676.04 | 581.06 | 487.47 | 689.72 | 609.61 | 670.42 | 747.73 | 716.59 | 779.88 |
| ENSMUSG00000040836 | 450.48 | 484.00 | 440.03 | 462.06 | 429.85 | 518.79 | 403.76 | 377.61 | 415.21 | 461.25 | 435.00 | 467.17 |
| ENSMUSG00000036565 | 987.73 | 941.49 | 1041.76 | 1070.91 | 857.54 | 1035.49 | 877.66 | 870.39 | 944.28 | 1069.61 | 1011.30 | 988.97 |
| ENSMUSG00000026586 | 3570.66 | 3519.56 | 4120.99 | 3515.61 | 5131.21 | 4854.90 | 2944.13 | 2827.55 | 3192.08 | 3567.21 | 3342.74 | 3326.72 |
| ENSMUSG00000035504 | 134.56 | 153.15 | 154.84 | 138.52 | 110.16 | 160.75 | 160.07 | 162.89 | 153.13 | 179.19 | 172.59 | 192.14 |

| | | | | | | | | | | | | |
|---|---|---|---|---|---|---|---|---|---|---|---|---|
| ENSMUSG00000021728 | 1416.76 | 1564.90 | 1700.34 | 1325.20 | 1567.12 | 1421.71 | 1170.82 | 1090.87 | 1173.96 | 1268.71 | 1333.26 | 1323.34 |
| ENSMUSG00000078517 | 1056.96 | 1046.54 | 1027.06 | 1008.89 | 1144.83 | 1151.36 | 1080.89 | 1108.97 | 1162.18 | 1263.18 | 1259.58 | 1307.33 |
| ENSMUSG00000015134 | 108.23 | 102.10 | 103.88 | 93.03 | 97.20 | 112.73 | 89.92 | 98.72 | 91.29 | 111.72 | 104.97 | 103.61 |
| ENSMUSG00000034910 | 212.56 | 162.97 | 202.86 | 150.92 | 253.81 | 231.73 | 138.48 | 149.73 | 148.22 | 157.07 | 176.62 | 165.77 |
| ENSMUSG00000027230 | 5267.25 | 4655.43 | 4309.15 | 4616.50 | 4320.11 | 4728.60 | 4214.77 | 4260.66 | 4150.10 | 4889.01 | 5009.07 | 4586.01 |
| ENSMUSG00000039103 | 3553.11 | 3795.43 | 4058.27 | 3942.53 | 5770.59 | 5570.98 | 3983.66 | 3672.44 | 3995.01 | 4323.78 | 4767.85 | 4279.90 |
| ENSMUSG00000080775 | 714.72 | 832.52 | 780.10 | 791.81 | 819.74 | 738.00 | 764.36 | 691.05 | 772.50 | 847.28 | 808.43 | 901.38 |
| ENSMUSG00000014850 | 277.89 | 292.56 | 333.21 | 363.86 | 344.53 | 362.21 | 302.15 | 272.31 | 275.82 | 313.03 | 326.00 | 337.19 |
| ENSMUSG00000039158 | 137.48 | 123.70 | 110.74 | 135.41 | 103.68 | 152.40 | 112.41 | 125.05 | 124.66 | 131.63 | 139.28 | 145.05 |
| ENSMUSG00000000440 | 156.98 | 143.33 | 147.98 | 151.95 | 171.72 | 152.40 | 123.20 | 109.42 | 111.90 | 138.26 | 131.21 | 128.10 |
| ENSMUSG00000081568 | 155.03 | 244.45 | 158.76 | 282.20 | 210.61 | 172.23 | 228.41 | 241.87 | 251.28 | 273.21 | 269.48 | 290.10 |
| ENSMUSG00000091014 | 46.80 | 43.20 | 54.88 | 62.02 | 64.80 | 38.62 | 70.14 | 64.17 | 63.80 | 71.90 | 80.74 | 76.29 |
| ENSMUSG00000025608 | 79.95 | 62.83 | 64.68 | 72.36 | 45.36 | 79.33 | 62.95 | 57.59 | 64.78 | 74.11 | 71.66 | 68.76 |
| ENSMUSG00000001995 | 374.42 | 408.41 | 414.55 | 436.22 | 357.49 | 431.11 | 413.65 | 438.49 | 445.64 | 484.48 | 507.67 | 510.50 |
| ENSMUSG00000025150 | 1013.08 | 748.09 | 849.68 | 795.95 | 815.42 | 636.74 | 732.89 | 819.39 | 738.14 | 822.95 | 910.37 | 920.22 |
| ENSMUSG00000083627 | 312.99 | 278.82 | 277.35 | 311.14 | 227.89 | 265.14 | 248.19 | 255.03 | 276.80 | 295.33 | 308.84 | 300.46 |
| ENSMUSG00000109495 | 1546.44 | 1604.17 | 1386.73 | 1813.11 | 1570.36 | 1349.69 | 1802.09 | 1712.00 | 1875.79 | 2094.97 | 1993.33 | 2164.44 |
| ENSMUSG00000025507 | 70.20 | 47.12 | 62.72 | 58.92 | 52.92 | 61.59 | 44.96 | 49.36 | 50.06 | 58.62 | 52.48 | 56.51 |
| ENSMUSG00000045038 | 131.63 | 164.93 | 173.46 | 148.85 | 180.36 | 179.54 | 133.09 | 118.47 | 137.42 | 158.17 | 142.31 | 151.64 |
| ENSMUSG00000020747 | 621.11 | 612.61 | 555.67 | 685.34 | 457.93 | 629.44 | 583.61 | 538.03 | 601.71 | 683.58 | 667.14 | 654.61 |
| ENSMUSG00000037217 | 260.34 | 212.06 | 243.05 | 248.09 | 255.97 | 278.71 | 225.71 | 218.01 | 223.80 | 264.36 | 257.37 | 256.19 |
| ENSMUSG00000031523 | 850.25 | 911.06 | 925.14 | 954.10 | 901.82 | 973.90 | 819.21 | 720.67 | 753.85 | 912.54 | 857.89 | 903.26 |
| ENSMUSG00000050953 | 5246.78 | 5403.52 | 5588.08 | 4908.00 | 6502.85 | 5924.84 | 4612.23 | 4379.95 | 4966.77 | 5508.43 | 5523.80 | 5262.28 |
| ENSMUSG00000022836 | 888.28 | 999.42 | 948.66 | 967.54 | 896.42 | 1082.46 | 1057.51 | 972.41 | 1029.67 | 1158.10 | 1190.95 | 1223.50 |
| ENSMUSG00000083705 | 1015.03 | 1050.47 | 1017.26 | 1167.05 | 1706.44 | 1006.26 | 1100.68 | 987.22 | 1068.94 | 1185.75 | 1236.37 | 1264.95 |
| ENSMUSG00000073060 | 135.53 | 159.04 | 169.54 | 154.02 | 146.88 | 186.85 | 156.47 | 143.97 | 164.90 | 174.77 | 183.69 | 188.38 |
| ENSMUSG00000025140 | 221.34 | 194.39 | 190.12 | 189.17 | 167.40 | 180.58 | 178.95 | 159.60 | 173.74 | 195.78 | 190.75 | 216.63 |
| ENSMUSG00000031434 | 410.50 | 430.99 | 371.43 | 433.12 | 439.57 | 489.56 | 380.38 | 350.46 | 411.28 | 428.06 | 466.29 | 463.41 |
| ENSMUSG00000018334 | 360.77 | 416.26 | 381.23 | 423.82 | 278.65 | 382.05 | 313.84 | 299.46 | 331.77 | 365.02 | 364.35 | 394.65 |
| ENSMUSG00000108793 | 79.95 | 78.54 | 89.18 | 94.07 | 96.12 | 97.08 | 80.93 | 75.69 | 68.71 | 89.59 | 90.84 | 87.59 |
| ENSMUSG00000082634 | 73.13 | 70.69 | 59.78 | 92.00 | 116.64 | 57.41 | 87.23 | 74.86 | 85.40 | 95.13 | 96.89 | 102.66 |
| ENSMUSG00000063382 | 1419.68 | 1609.08 | 1509.23 | 1845.15 | 1411.60 | 1849.68 | 1524.22 | 1380.46 | 1581.32 | 1814.02 | 1835.88 | 1699.15 |
| ENSMUSG00000086670 | 84.83 | 138.43 | 85.26 | 141.62 | 110.16 | 92.90 | 128.59 | 123.40 | 131.53 | 151.54 | 144.33 | 162.00 |
| ENSMUSG00000055660 | 123.83 | 134.50 | 171.50 | 150.92 | 198.73 | 204.59 | 125.00 | 113.53 | 125.64 | 133.84 | 148.36 | 152.58 |

| | | | | | | | | | | | | |
|---|---|---|---|---|---|---|---|---|---|---|---|---|
| ENSMUSG00000026527 | 18.53 | 18.65 | 15.68 | 21.71 | 18.36 | 25.05 | 20.68 | 19.74 | 19.63 | 22.12 | 24.22 | 25.43 |
| ENSMUSG00000037568 | 477.78 | 508.54 | 468.45 | 484.81 | 509.77 | 568.89 | 396.57 | 430.26 | 442.69 | 523.19 | 506.66 | 489.78 |
| ENSMUSG00000031451 | 1468.43 | 1214.42 | 1372.03 | 1204.26 | 907.22 | 1075.16 | 1114.17 | 990.51 | 1085.62 | 1278.66 | 1364.55 | 1176.41 |
| ENSMUSG00000040483 | 235.96 | 207.15 | 212.66 | 206.74 | 234.37 | 203.55 | 178.05 | 193.33 | 188.46 | 228.96 | 222.04 | 220.40 |
| ENSMUSG00000044949 | 297.39 | 369.14 | 319.49 | 339.05 | 328.33 | 364.30 | 260.78 | 277.24 | 302.33 | 344.00 | 341.14 | 323.06 |
| ENSMUSG00000038740 | 568.46 | 568.43 | 584.09 | 595.41 | 407.17 | 602.30 | 484.69 | 460.70 | 513.36 | 579.60 | 605.57 | 568.90 |
| ENSMUSG00000042155 | 133.58 | 131.55 | 139.16 | 128.18 | 149.04 | 115.87 | 105.21 | 98.72 | 109.94 | 133.84 | 123.13 | 120.56 |
| ENSMUSG00000078484 | 315.92 | 281.76 | 264.61 | 320.45 | 236.53 | 286.01 | 276.07 | 265.73 | 235.58 | 319.67 | 322.97 | 292.92 |
| ENSMUSG00000028532 | 288.62 | 344.59 | 273.43 | 390.74 | 281.89 | 311.06 | 287.76 | 273.95 | 317.05 | 326.30 | 353.25 | 380.52 |
| ENSMUSG00000049551 | 22.43 | 21.60 | 30.38 | 24.81 | 21.60 | 18.79 | 25.18 | 24.68 | 26.50 | 29.86 | 32.30 | 30.14 |
| ENSMUSG00000083815 | 47.78 | 38.29 | 44.10 | 51.68 | 46.44 | 40.71 | 46.76 | 49.36 | 55.95 | 65.26 | 56.52 | 62.16 |
| ENSMUSG00000038855 | 275.94 | 320.05 | 269.51 | 266.69 | 216.01 | 256.78 | 210.42 | 219.66 | 233.61 | 277.63 | 271.50 | 254.31 |
| ENSMUSG00000038248 | 467.05 | 452.58 | 491.97 | 432.09 | 473.05 | 484.34 | 369.59 | 340.59 | 388.70 | 441.34 | 460.23 | 429.50 |
| ENSMUSG00000020275 | 224.26 | 247.40 | 280.29 | 246.02 | 340.21 | 376.83 | 201.43 | 201.56 | 211.04 | 268.78 | 232.13 | 243.95 |
| ENSMUSG00000025964 | 198.91 | 195.37 | 153.86 | 202.61 | 185.76 | 204.59 | 154.67 | 158.78 | 158.03 | 205.74 | 183.69 | 182.72 |
| ENSMUSG00000087591 | 891.20 | 714.71 | 753.64 | 751.50 | 717.14 | 777.66 | 777.85 | 734.65 | 740.11 | 873.83 | 979.00 | 893.84 |
| ENSMUSG00000032265 | 245.71 | 244.45 | 293.03 | 272.90 | 416.89 | 353.86 | 202.33 | 195.80 | 214.96 | 264.36 | 231.13 | 252.42 |
| ENSMUSG00000063455 | 251.56 | 284.71 | 266.57 | 292.54 | 232.21 | 330.90 | 234.70 | 222.12 | 265.03 | 280.95 | 306.82 | 293.87 |
| ENSMUSG00000105967 | 37.05 | 55.96 | 48.02 | 48.58 | 39.96 | 43.84 | 35.97 | 32.08 | 33.37 | 39.82 | 44.41 | 40.50 |
| ENSMUSG00000050014 | 317.87 | 372.08 | 332.23 | 308.04 | 363.97 | 363.26 | 255.39 | 265.73 | 232.63 | 332.94 | 286.64 | 309.88 |
| ENSMUSG00000107724 | 37.05 | 46.14 | 33.32 | 43.42 | 54.00 | 39.67 | 31.47 | 32.91 | 28.47 | 40.93 | 36.33 | 37.68 |
| ENSMUSG00000032177 | 156.98 | 197.33 | 205.80 | 198.47 | 174.96 | 206.68 | 161.86 | 170.29 | 164.90 | 207.95 | 188.74 | 219.46 |
| ENSMUSG00000053101 | 59.48 | 84.43 | 74.48 | 57.89 | 89.64 | 57.41 | 49.46 | 51.83 | 56.93 | 68.58 | 62.58 | 64.99 |
| ENSMUSG00000103715 | 12.68 | 12.76 | 9.80 | 11.37 | 14.04 | 11.48 | 10.79 | 9.87 | 8.83 | 12.17 | 13.12 | 11.30 |
| ENSMUSG00000104344 | 8.78 | 27.49 | 28.42 | 25.84 | 15.12 | 34.45 | 16.19 | 15.63 | 14.72 | 18.80 | 19.18 | 19.78 |
| ENSMUSG00000035954 | 327.62 | 452.58 | 416.51 | 426.92 | 416.89 | 512.53 | 341.71 | 296.99 | 361.22 | 426.96 | 398.67 | 420.08 |
| ENSMUSG00000028128 | 248.64 | 273.91 | 332.23 | 258.42 | 270.01 | 275.57 | 197.83 | 193.33 | 200.24 | 266.57 | 231.13 | 239.24 |
| ENSMUSG00000078651 | 25.35 | 21.60 | 27.44 | 28.94 | 22.68 | 30.27 | 22.48 | 24.68 | 25.52 | 33.18 | 28.26 | 29.20 |
| ENSMUSG00000063873 | 371.50 | 354.41 | 355.75 | 370.06 | 287.29 | 314.20 | 283.26 | 269.02 | 291.53 | 365.02 | 343.16 | 345.67 |
| ENSMUSG00000022371 | 376.37 | 471.24 | 447.87 | 395.91 | 396.37 | 479.12 | 318.33 | 259.14 | 325.88 | 352.85 | 378.48 | 397.47 |
| ENSMUSG00000066392 | 302.27 | 299.43 | 299.89 | 258.42 | 493.57 | 431.11 | 317.43 | 292.87 | 290.55 | 348.42 | 404.72 | 373.93 |
| ENSMUSG00000081173 | 28.28 | 18.65 | 28.42 | 21.71 | 44.28 | 24.01 | 17.09 | 18.92 | 15.71 | 21.02 | 20.19 | 23.55 |
| ENSMUSG00000001506 | 22641.78 | 22794.15 | 19996.34 | 23546.64 | 19869.28 | 22552.17 | 21708.70 | 18641.09 | 19918.12 | 23986.00 | 26045.53 | 25455.25 |
| ENSMUSG00000052632 | 309.09 | 373.06 | 349.87 | 324.58 | 345.61 | 386.22 | 281.46 | 280.53 | 328.83 | 398.20 | 356.28 | 363.57 |

| | | | | | | | | | | | | |
|---|---|---|---|---|---|---|---|---|---|---|---|---|
| ENSMUSG00000030074 | 1476.24 | 1452.98 | 1589.59 | 1420.30 | 2023.97 | 1845.51 | 1087.19 | 1076.89 | 1091.51 | 1360.52 | 1446.30 | 1287.55 |
| ENSMUSG00000050677 | 29.25 | 27.49 | 23.52 | 31.01 | 28.08 | 32.36 | 26.98 | 27.97 | 29.45 | 32.08 | 38.35 | 35.79 |
| ENSMUSG00000006800 | 1829.21 | 1713.14 | 1884.58 | 1861.69 | 1339.23 | 1702.50 | 1490.95 | 1309.71 | 1625.49 | 1808.49 | 1968.10 | 1801.82 |
| ENSMUSG00000082545 | 18.53 | 23.56 | 23.52 | 39.28 | 23.76 | 12.53 | 30.57 | 28.79 | 28.47 | 38.71 | 34.32 | 37.68 |
| ENSMUSG00000039824 | 2576.10 | 1756.34 | 1983.56 | 1668.39 | 1401.88 | 1539.66 | 1495.45 | 1697.19 | 1394.82 | 1839.46 | 2089.21 | 1860.21 |
| ENSMUSG00000034810 | 117.98 | 150.21 | 133.28 | 134.38 | 238.69 | 183.72 | 135.79 | 110.24 | 130.55 | 170.34 | 146.35 | 159.18 |
| ENSMUSG00000026249 | 873.65 | 859.03 | 1065.28 | 973.74 | 911.54 | 792.27 | 684.33 | 686.94 | 818.63 | 882.68 | 963.86 | 928.69 |
| ENSMUSG00000083743 | 69.23 | 76.58 | 67.62 | 94.07 | 111.24 | 65.76 | 81.83 | 75.69 | 75.58 | 99.55 | 97.90 | 97.96 |
| ENSMUSG00000036446 | 691.31 | 690.17 | 744.82 | 619.19 | 659.90 | 686.85 | 538.65 | 428.62 | 521.22 | 601.72 | 666.13 | 618.82 |
| ENSMUSG00000056219 | 9.75 | 8.84 | 6.86 | 7.24 | 3.24 | 6.26 | 7.19 | 6.58 | 7.85 | 8.85 | 10.09 | 8.48 |
| ENSMUSG00000027386 | 1612.74 | 1196.75 | 1679.76 | 1411.00 | 1188.03 | 1270.35 | 1116.86 | 1155.04 | 1173.96 | 1389.27 | 1564.39 | 1420.36 |
| ENSMUSG00000031698 | 96.53 | 87.38 | 100.94 | 103.37 | 104.76 | 85.59 | 86.33 | 83.09 | 78.53 | 103.97 | 114.05 | 97.01 |
| ENSMUSG00000049420 | 100.43 | 111.92 | 107.80 | 106.47 | 106.92 | 120.04 | 82.73 | 74.86 | 70.67 | 100.66 | 94.87 | 96.07 |
| ENSMUSG00000029838 | 1636.14 | 1714.13 | 2250.13 | 1569.16 | 1933.25 | 1689.98 | 1350.67 | 1303.12 | 1400.71 | 1872.64 | 1695.59 | 1613.44 |
| ENSMUSG00000083470 | 35.10 | 51.05 | 34.30 | 56.85 | 52.92 | 45.93 | 44.96 | 46.07 | 54.97 | 59.73 | 68.63 | 58.40 |
| ENSMUSG00000107369 | 233.04 | 202.24 | 204.82 | 204.67 | 159.84 | 202.51 | 184.35 | 194.98 | 179.63 | 238.92 | 245.26 | 234.53 |
| ENSMUSG00000109560 | 62.40 | 68.72 | 56.84 | 82.70 | 81.00 | 45.93 | 68.34 | 58.41 | 70.67 | 90.70 | 74.69 | 88.54 |
| ENSMUSG00000022037 | 33.15 | 22.58 | 28.42 | 18.61 | 24.84 | 24.01 | 17.98 | 22.21 | 22.58 | 26.55 | 26.24 | 28.26 |
| ENSMUSG00000082872 | 45.83 | 102.10 | 83.30 | 88.90 | 75.60 | 100.21 | 76.44 | 68.28 | 84.42 | 98.44 | 103.96 | 95.13 |
| ENSMUSG00000046952 | 34.13 | 56.94 | 36.26 | 47.55 | 44.28 | 52.19 | 32.37 | 33.73 | 39.26 | 46.46 | 43.40 | 48.04 |
| ENSMUSG00000086040 | 72.15 | 77.56 | 64.68 | 72.36 | 82.08 | 85.59 | 62.95 | 61.70 | 71.65 | 85.17 | 84.78 | 87.59 |
| ENSMUSG00000083890 | 45.83 | 35.34 | 32.34 | 27.91 | 31.32 | 32.36 | 30.57 | 27.97 | 31.41 | 40.93 | 40.37 | 36.73 |
| ENSMUSG00000084081 | 28.28 | 37.31 | 23.52 | 38.25 | 29.16 | 20.88 | 24.28 | 23.86 | 30.43 | 34.29 | 33.31 | 35.79 |
| ENSMUSG00000041112 | 720.57 | 822.70 | 777.16 | 734.96 | 748.46 | 815.24 | 606.99 | 520.76 | 662.56 | 798.61 | 758.98 | 810.02 |
| ENSMUSG00000033949 | 36.08 | 27.49 | 32.34 | 25.84 | 41.04 | 48.02 | 33.27 | 26.33 | 29.45 | 34.29 | 40.37 | 43.33 |
| ENSMUSG00000070526 | 14.63 | 20.62 | 26.46 | 29.98 | 24.84 | 18.79 | 19.78 | 17.28 | 15.71 | 23.23 | 23.21 | 23.55 |
| ENSMUSG00000067773 | 19.50 | 21.60 | 18.62 | 17.57 | 28.08 | 17.75 | 13.49 | 15.63 | 13.74 | 21.02 | 17.16 | 18.84 |
| ENSMUSG00000062488 | 31.20 | 43.20 | 66.64 | 43.42 | 33.48 | 44.89 | 35.07 | 33.73 | 41.23 | 47.56 | 51.47 | 48.04 |
| ENSMUSG00000064288 | 29.25 | 28.47 | 36.26 | 39.28 | 23.76 | 27.14 | 26.08 | 26.33 | 21.59 | 30.97 | 31.29 | 36.73 |
| ENSMUSG00000039145 | 44.85 | 66.76 | 81.34 | 81.66 | 85.32 | 102.30 | 53.95 | 55.94 | 59.88 | 76.32 | 74.69 | 77.23 |
| ENSMUSG00000031785 | 284.72 | 251.33 | 267.55 | 244.99 | 211.69 | 245.30 | 242.80 | 244.34 | 203.19 | 286.48 | 327.01 | 316.47 |
| ENSMUSG00000032596 | 101.41 | 91.30 | 99.96 | 111.64 | 69.12 | 82.46 | 70.14 | 84.74 | 68.71 | 102.87 | 97.90 | 100.78 |
| ENSMUSG00000006344 | 14.63 | 14.73 | 13.72 | 16.54 | 11.88 | 13.57 | 10.79 | 10.69 | 13.74 | 14.38 | 17.16 | 16.01 |
| ENSMUSG00000091426 | 260.34 | 329.87 | 215.60 | 362.83 | 305.65 | 232.78 | 315.64 | 247.63 | 322.94 | 441.34 | 356.28 | 399.36 |

| | | | | | | | | | | | | |
|---|---|---|---|---|---|---|---|---|---|---|---|---|
| ENSMUSG00000084124 | 129.68 | 166.90 | 103.88 | 164.36 | 159.84 | 103.34 | 140.28 | 123.40 | 139.38 | 196.89 | 159.47 | 188.38 |
| ENSMUSG00000056596 | 3.90 | 7.85 | 7.84 | 9.30 | 4.32 | 6.26 | 5.40 | 7.40 | 5.89 | 7.74 | 9.08 | 8.48 |
| ENSMUSG00000061540 | 4.88 | 9.82 | 8.82 | 9.30 | 3.24 | 6.26 | 8.09 | 6.58 | 7.85 | 11.06 | 9.08 | 10.36 |
| ENSMUSG00000061825 | 95.56 | 96.21 | 94.08 | 94.07 | 85.32 | 80.38 | 62.05 | 72.40 | 85.40 | 91.81 | 103.96 | 102.66 |
| ENSMUSG00000029309 | 221.34 | 191.44 | 283.23 | 215.01 | 190.08 | 240.08 | 186.14 | 161.25 | 147.24 | 228.96 | 247.27 | 199.68 |
| ENSMUSG00000058057 | 20.48 | 28.47 | 21.56 | 14.47 | 20.52 | 24.01 | 15.29 | 17.28 | 15.71 | 24.33 | 20.19 | 21.66 |
| ENSMUSG00000067455 | 34.13 | 30.43 | 37.24 | 40.31 | 27.00 | 28.18 | 24.28 | 31.26 | 23.56 | 32.08 | 38.35 | 38.62 |
| ENSMUSG00000069266 | 29.25 | 29.45 | 35.28 | 38.25 | 24.84 | 28.18 | 25.18 | 26.33 | 21.59 | 29.86 | 35.32 | 35.79 |
| ENSMUSG00000080844 | 32.18 | 67.74 | 43.12 | 52.72 | 62.64 | 51.15 | 55.75 | 54.30 | 62.82 | 88.49 | 68.63 | 82.89 |
| ENSMUSG00000060807 | 19.50 | 26.51 | 23.52 | 22.74 | 18.36 | 14.61 | 14.39 | 13.16 | 16.69 | 17.70 | 21.19 | 22.61 |
| ENSMUSG00000045672 | 69.23 | 72.65 | 93.10 | 96.13 | 87.48 | 105.43 | 53.06 | 55.12 | 49.08 | 78.53 | 68.63 | 71.58 |
| ENSMUSG00000042759 | 55.58 | 39.27 | 49.98 | 39.28 | 37.80 | 36.53 | 35.07 | 40.31 | 29.45 | 46.46 | 51.47 | 48.04 |
| ENSMUSG00000048478 | 20.48 | 25.53 | 25.48 | 19.64 | 18.36 | 18.79 | 24.28 | 22.21 | 26.50 | 37.61 | 30.28 | 33.91 |
| ENSMUSG00000081537 | 20.48 | 23.56 | 18.62 | 23.78 | 14.04 | 18.79 | 19.78 | 15.63 | 16.69 | 21.02 | 27.25 | 24.49 |
| ENSMUSG00000079450 | 18.53 | 20.62 | 24.50 | 17.57 | 28.08 | 25.05 | 15.29 | 18.10 | 18.65 | 21.02 | 26.24 | 25.43 |
| ENSMUSG00000101363 | 16.58 | 23.56 | 10.78 | 17.57 | 20.52 | 14.61 | 12.59 | 9.87 | 9.82 | 13.27 | 16.15 | 16.01 |
| ENSMUSG00000039321 | 32.18 | 32.40 | 23.52 | 33.08 | 33.48 | 29.23 | 29.68 | 23.86 | 21.59 | 35.40 | 39.36 | 31.08 |
| ENSMUSG00000082608 | 24.38 | 30.43 | 31.36 | 32.04 | 22.68 | 16.70 | 26.98 | 20.57 | 26.50 | 35.40 | 31.29 | 37.68 |
| ENSMUSG00000086657 | 5.85 | 8.84 | 7.84 | 10.34 | 5.40 | 10.44 | 8.09 | 6.58 | 6.87 | 11.06 | 9.08 | 10.36 |
| ENSMUSG00000023915 | 220.36 | 206.17 | 175.42 | 198.47 | 192.24 | 167.01 | 142.98 | 127.52 | 154.11 | 214.59 | 190.75 | 195.91 |
| ENSMUSG00000101609 | 90.68 | 113.88 | 153.86 | 162.29 | 535.69 | 509.39 | 87.23 | 118.47 | 114.84 | 150.43 | 153.41 | 153.53 |
| ENSMUSG00000022766 | 5.85 | 9.82 | 11.76 | 5.17 | 12.96 | 7.31 | 11.69 | 9.05 | 9.82 | 14.38 | 15.14 | 14.13 |
| ENSMUSG00000071573 | 72.15 | 61.85 | 53.90 | 53.75 | 74.52 | 60.54 | 38.67 | 46.07 | 45.15 | 57.52 | 70.65 | 57.45 |
| ENSMUSG00000038319 | 96.53 | 105.05 | 108.78 | 106.47 | 77.76 | 105.43 | 85.43 | 64.99 | 85.40 | 109.50 | 123.13 | 106.43 |
| ENSMUSG00000028637 | 15.60 | 17.67 | 13.72 | 16.54 | 17.28 | 25.05 | 12.59 | 9.87 | 10.80 | 17.70 | 14.13 | 16.01 |
| ENSMUSG00000038195 | 109.21 | 79.52 | 76.44 | 74.43 | 61.56 | 65.76 | 62.95 | 79.80 | 57.91 | 94.02 | 102.95 | 92.30 |
| ENSMUSG00000098120 | 7.80 | 10.80 | 7.84 | 14.47 | 3.24 | 12.53 | 9.89 | 7.40 | 8.83 | 13.27 | 13.12 | 11.30 |
| ENSMUSG00000048583 | 3752.02 | 2899.09 | 3111.57 | 2761.01 | 3038.12 | 3468.68 | 2534.98 | 2369.32 | 2873.07 | 3487.57 | 4151.18 | 3584.80 |
| ENSMUSG00000043811 | 49.73 | 33.38 | 40.18 | 35.15 | 22.68 | 33.40 | 21.58 | 29.62 | 21.59 | 30.97 | 39.36 | 34.85 |
| ENSMUSG00000087403 | 8.78 | 14.73 | 24.50 | 23.78 | 21.60 | 20.88 | 15.29 | 12.34 | 16.69 | 23.23 | 20.19 | 20.72 |
| ENSMUSG00000022206 | 401.72 | 436.88 | 488.05 | 381.44 | 416.89 | 447.81 | 257.18 | 215.54 | 257.17 | 352.85 | 356.28 | 349.44 |
| ENSMUSG00000029380 | 44.85 | 49.09 | 65.66 | 40.31 | 48.60 | 37.58 | 30.57 | 24.68 | 35.34 | 48.67 | 44.41 | 38.62 |
| ENSMUSG00000014813 | 292.52 | 269.00 | 310.67 | 257.39 | 302.41 | 347.60 | 216.72 | 179.34 | 205.15 | 313.03 | 290.67 | 272.20 |
| ENSMUSG00000097968 | 10.73 | 23.56 | 7.84 | 14.47 | 15.12 | 9.39 | 11.69 | 11.52 | 14.72 | 16.59 | 17.16 | 21.66 |

| | | | | | | | | | | | | |
|---|---|---|---|---|---|---|---|---|---|---|---|---|
| ENSMUSG00000063087 | 11.70 | 12.76 | 15.68 | 14.47 | 19.44 | 26.10 | 16.19 | 12.34 | 15.71 | 24.33 | 21.19 | 19.78 |
| ENSMUSG00000047793 | 265.22 | 315.14 | 328.31 | 383.50 | 257.05 | 354.91 | 291.36 | 214.72 | 265.03 | 420.32 | 393.62 | 326.83 |
| ENSMUSG00000030399 | 5429.11 | 4004.54 | 4765.84 | 3968.37 | 2742.19 | 3459.29 | 3485.48 | 3846.03 | 3691.70 | 5054.92 | 6089.00 | 5183.17 |
| ENSMUSG00000104876 | 79.95 | 75.59 | 78.40 | 67.19 | 69.12 | 89.77 | 62.05 | 51.01 | 56.93 | 80.75 | 97.90 | 73.47 |
| ENSMUSG00000001930 | 12.68 | 13.74 | 11.76 | 18.61 | 14.04 | 17.75 | 15.29 | 21.39 | 16.69 | 27.65 | 24.22 | 27.31 |
| ENSMUSG00000079055 | 9.75 | 13.74 | 10.78 | 15.51 | 7.56 | 11.48 | 12.59 | 11.52 | 7.85 | 14.38 | 16.15 | 16.95 |
| ENSMUSG00000051920 | 16.58 | 10.80 | 17.64 | 19.64 | 20.52 | 19.83 | 8.99 | 12.34 | 12.76 | 16.59 | 17.16 | 16.95 |
| ENSMUSG00000084118 | 10.73 | 4.91 | 12.74 | 5.17 | 6.48 | 10.44 | 6.29 | 5.76 | 4.91 | 7.74 | 8.07 | 9.42 |
| ENSMUSG00000037138 | 625.99 | 569.41 | 642.89 | 547.86 | 527.05 | 602.30 | 493.69 | 310.97 | 395.57 | 578.50 | 644.93 | 563.24 |
| ENSMUSG00000059352 | 5.85 | 10.80 | 8.82 | 12.40 | 11.88 | 6.26 | 9.89 | 9.05 | 7.85 | 15.49 | 13.12 | 11.30 |
| ENSMUSG00000103312 | 3.90 | 0.98 | 2.94 | 4.13 | 1.08 | 7.31 | 3.60 | 4.11 | 3.93 | 6.64 | 5.05 | 5.65 |
| ENSMUSG00000020486 | 13.65 | 16.69 | 8.82 | 12.40 | 12.96 | 14.61 | 7.19 | 11.52 | 10.80 | 15.49 | 13.12 | 16.01 |
| ENSMUSG00000022330 | 18.53 | 13.74 | 13.72 | 14.47 | 8.64 | 15.66 | 10.79 | 11.52 | 7.85 | 15.49 | 14.13 | 16.01 |
| ENSMUSG00000081258 | 4.88 | 6.87 | 8.82 | 15.51 | 16.20 | 6.26 | 13.49 | 9.05 | 11.78 | 18.80 | 17.16 | 16.01 |
| ENSMUSG00000004939 | 215.49 | 146.28 | 198.94 | 179.86 | 157.68 | 112.73 | 125.00 | 127.52 | 130.55 | 185.83 | 209.93 | 184.61 |
| ENSMUSG00000034570 | 105.31 | 77.56 | 109.76 | 92.00 | 56.16 | 80.38 | 92.62 | 71.57 | 68.71 | 106.19 | 131.21 | 118.68 |
| ENSMUSG00000108331 | 20.48 | 26.51 | 28.42 | 28.94 | 22.68 | 24.01 | 30.57 | 18.92 | 29.45 | 40.93 | 44.41 | 35.79 |
| ENSMUSG00000078891 | 12.68 | 12.76 | 19.60 | 14.47 | 17.28 | 22.96 | 9.89 | 9.87 | 11.78 | 15.49 | 16.15 | 16.95 |
| ENSMUSG00000043670 | 12.68 | 17.67 | 12.74 | 13.44 | 10.80 | 8.35 | 11.69 | 11.52 | 12.76 | 18.80 | 16.15 | 20.72 |
| ENSMUSG00000079998 | 37.05 | 29.45 | 33.32 | 41.35 | 60.48 | 34.45 | 28.78 | 27.15 | 40.24 | 44.24 | 48.45 | 56.51 |
| ENSMUSG00000084212 | 8.78 | 10.80 | 9.80 | 14.47 | 12.96 | 10.44 | 9.89 | 6.58 | 10.80 | 13.27 | 14.13 | 15.07 |
| ENSMUSG00000097312 | 0.98 | 4.91 | 1.96 | 8.27 | 21.60 | 21.92 | 2.70 | 2.47 | 1.96 | 3.32 | 4.04 | 3.77 |
| ENSMUSG00000104168 | 3.90 | 4.91 | 11.76 | 8.27 | 6.48 | 9.39 | 8.09 | 5.76 | 6.87 | 9.95 | 11.10 | 11.30 |
| ENSMUSG00000020788 | 21.45 | 28.47 | 23.52 | 27.91 | 17.28 | 21.92 | 25.18 | 18.92 | 23.56 | 36.50 | 34.32 | 34.85 |
| ENSMUSG00000044006 | 11.70 | 16.69 | 6.86 | 15.51 | 7.56 | 10.44 | 12.59 | 8.23 | 9.82 | 13.27 | 17.16 | 17.90 |
| ENSMUSG00000082054 | 11.70 | 6.87 | 1.96 | 5.17 | 8.64 | 6.26 | 8.09 | 7.40 | 6.87 | 9.95 | 13.12 | 12.24 |
| ENSMUSG00000102018 | 28.28 | 25.53 | 21.56 | 25.84 | 25.92 | 24.01 | 12.59 | 18.92 | 19.63 | 24.33 | 28.26 | 28.26 |
| ENSMUSG00000108638 | 6.83 | 4.91 | 2.94 | 5.17 | 10.80 | 6.26 | 5.40 | 4.94 | 4.91 | 7.74 | 7.06 | 9.42 |
| ENSMUSG00000073733 | 5.85 | 11.78 | 3.92 | 8.27 | 5.40 | 9.39 | 4.50 | 5.76 | 6.87 | 7.74 | 10.09 | 9.42 |
| ENSMUSG00000059571 | 3.90 | 11.78 | 10.78 | 9.30 | 11.88 | 4.18 | 8.09 | 7.40 | 10.80 | 15.49 | 15.14 | 11.30 |
| ENSMUSG00000014158 | 18.53 | 8.84 | 9.80 | 20.67 | 8.64 | 7.31 | 7.19 | 8.23 | 11.78 | 14.38 | 13.12 | 16.01 |
| ENSMUSG00000050022 | 10.73 | 7.85 | 9.80 | 5.17 | 8.64 | 5.22 | 5.40 | 6.58 | 8.83 | 13.27 | 10.09 | 10.36 |
| ENSMUSG00000028360 | 11.70 | 16.69 | 21.56 | 11.37 | 19.44 | 24.01 | 6.29 | 7.40 | 8.83 | 13.27 | 10.09 | 13.19 |
| ENSMUSG00000106551 | 16.58 | 13.74 | 10.78 | 16.54 | 17.28 | 15.66 | 7.19 | 9.87 | 12.76 | 16.59 | 14.13 | 17.90 |

| | | | | | | | | | | | | |
|---|---|---|---|---|---|---|---|---|---|---|---|---|
| ENSMUSG00000083387 | 11.70 | 19.63 | 15.68 | 14.47 | 21.60 | 11.48 | 11.69 | 11.52 | 11.78 | 17.70 | 17.16 | 22.61 |
| ENSMUSG00000109480 | 11.70 | 15.71 | 15.68 | 14.47 | 9.72 | 7.31 | 9.89 | 10.69 | 13.74 | 17.70 | 16.15 | 22.61 |
| ENSMUSG00000097893 | 2.93 | 2.95 | 1.96 | 5.17 | 1.08 | 4.18 | 3.60 | 3.29 | 2.94 | 5.53 | 5.05 | 5.65 |
| ENSMUSG00000096233 | 13.65 | 19.63 | 13.72 | 24.81 | 28.08 | 17.75 | 14.39 | 12.34 | 20.61 | 22.12 | 27.25 | 29.20 |
| ENSMUSG00000094306 | 6.83 | 14.73 | 17.64 | 10.34 | 24.84 | 5.22 | 4.50 | 7.40 | 6.87 | 12.17 | 11.10 | 8.48 |
| ENSMUSG00000037362 | 1209.07 | 1067.16 | 1249.53 | 951.00 | 1206.39 | 983.30 | 761.66 | 591.51 | 584.04 | 1037.53 | 1220.22 | 1021.00 |
| ENSMUSG00000106746 | 1.95 | 6.87 | 7.84 | 1.03 | 7.56 | 7.31 | 2.70 | 3.29 | 2.94 | 4.42 | 6.06 | 4.71 |
| ENSMUSG00000085433 | 11.70 | 16.69 | 12.74 | 18.61 | 11.88 | 10.44 | 11.69 | 8.23 | 8.83 | 15.49 | 15.14 | 18.84 |
| ENSMUSG00000034818 | 33.15 | 26.51 | 34.30 | 24.81 | 35.64 | 29.23 | 17.98 | 18.10 | 22.58 | 39.82 | 32.30 | 30.14 |
| ENSMUSG00000089684 | 6.83 | 7.85 | 6.86 | 9.30 | 6.48 | 5.22 | 8.09 | 8.23 | 3.93 | 9.95 | 13.12 | 12.24 |
| ENSMUSG00000098843 | 3.90 | 9.82 | 4.90 | 6.20 | 6.48 | 5.22 | 6.29 | 4.94 | 4.91 | 7.74 | 10.09 | 10.36 |
| ENSMUSG00000070780 | 10.73 | 6.87 | 10.78 | 9.30 | 6.48 | 10.44 | 9.89 | 8.23 | 11.78 | 21.02 | 16.15 | 15.07 |
| ENSMUSG00000051599 | 1.95 | 3.93 | 7.84 | 3.10 | 3.24 | 6.26 | 3.60 | 5.76 | 3.93 | 7.74 | 8.07 | 7.54 |
| ENSMUSG00000081693 | 7.80 | 24.54 | 9.80 | 22.74 | 17.28 | 14.61 | 12.59 | 17.28 | 17.67 | 33.18 | 23.21 | 27.31 |
| ENSMUSG00000009145 | 9.75 | 14.73 | 11.76 | 18.61 | 20.52 | 13.57 | 8.99 | 9.87 | 5.89 | 14.38 | 15.14 | 14.13 |
| ENSMUSG00000004035 | 31.20 | 23.56 | 24.50 | 34.11 | 17.28 | 18.79 | 17.09 | 19.74 | 15.71 | 36.50 | 28.26 | 28.26 |
| ENSMUSG00000081132 | 8.78 | 8.84 | 14.70 | 12.40 | 8.64 | 5.22 | 7.19 | 4.94 | 8.83 | 9.95 | 14.13 | 13.19 |
| ENSMUSG00000039457 | 186.24 | 150.21 | 231.28 | 192.27 | 129.60 | 178.50 | 133.09 | 82.27 | 115.83 | 181.40 | 222.04 | 188.38 |
| ENSMUSG00000057457 | 96.53 | 72.65 | 119.56 | 90.97 | 125.28 | 141.96 | 51.26 | 45.25 | 45.15 | 98.44 | 86.80 | 68.76 |
| ENSMUSG00000102496 | 6.83 | 7.85 | 21.56 | 7.24 | 38.88 | 20.88 | 6.29 | 4.94 | 3.93 | 8.85 | 9.08 | 9.42 |
| ENSMUSG00000054568 | 0.98 | 2.95 | 2.94 | 3.10 | 1.08 | 7.31 | 1.80 | 2.47 | 2.94 | 3.32 | 5.05 | 4.71 |
| ENSMUSG00000085289 | 18.53 | 23.56 | 17.64 | 27.91 | 7.56 | 19.83 | 17.09 | 9.87 | 11.78 | 21.02 | 25.23 | 24.49 |
| ENSMUSG00000083676 | 7.80 | 21.60 | 18.62 | 14.47 | 18.36 | 16.70 | 17.09 | 9.05 | 15.71 | 19.91 | 28.26 | 28.26 |
| ENSMUSG00000096206 | 7.80 | 15.71 | 19.60 | 10.34 | 27.00 | 6.26 | 6.29 | 7.40 | 6.87 | 13.27 | 14.13 | 10.36 |
| ENSMUSG00000096659 | 7.80 | 15.71 | 19.60 | 10.34 | 27.00 | 6.26 | 6.29 | 7.40 | 6.87 | 13.27 | 14.13 | 10.36 |
| ENSMUSG00000038591 | 156.98 | 154.13 | 146.02 | 192.27 | 218.17 | 174.32 | 98.02 | 84.74 | 109.94 | 182.51 | 190.75 | 166.71 |
| ENSMUSG00000095054 | 7.80 | 15.71 | 11.76 | 20.67 | 24.84 | 12.53 | 12.59 | 9.05 | 13.74 | 17.70 | 24.22 | 23.55 |
| ENSMUSG00000005705 | 8.78 | 4.91 | 4.90 | 12.40 | 4.32 | 7.31 | 7.19 | 4.11 | 4.91 | 7.74 | 12.11 | 10.36 |
| ENSMUSG00000034780 | 18.53 | 14.73 | 19.60 | 18.61 | 23.76 | 30.27 | 10.79 | 5.76 | 11.78 | 15.49 | 18.17 | 19.78 |
| ENSMUSG00000054409 | 3.90 | 0.98 | 0.98 | 3.10 | 3.24 | 1.04 | 1.80 | 1.65 | 2.94 | 3.32 | 4.04 | 4.71 |
| ENSMUSG00000105836 | 3.90 | 2.95 | 4.90 | 8.27 | 6.48 | 2.09 | 2.70 | 2.47 | 3.93 | 6.64 | 6.06 | 4.71 |
| ENSMUSG00000084157 | 4.88 | 10.80 | 10.78 | 9.30 | 8.64 | 10.44 | 3.60 | 4.94 | 7.85 | 11.06 | 9.08 | 11.30 |
| ENSMUSG00000056888 | 11.70 | 5.89 | 6.86 | 6.20 | 6.48 | 9.39 | 3.60 | 4.11 | 6.87 | 9.95 | 9.08 | 9.42 |
| ENSMUSG00000086266 | 8.78 | 11.78 | 13.72 | 12.40 | 8.64 | 11.48 | 3.60 | 6.58 | 6.87 | 11.06 | 10.09 | 12.24 |

| | | | | | | | | | | | | |
|---|---|---|---|---|---|---|---|---|---|---|---|---|
| ENSMUSG00000040624 | 14.63 | 12.76 | 19.60 | 15.51 | 17.28 | 13.57 | 14.39 | 7.40 | 10.80 | 22.12 | 22.20 | 19.78 |
| ENSMUSG00000021130 | 8.78 | 12.76 | 14.70 | 10.34 | 6.48 | 9.39 | 5.40 | 4.11 | 7.85 | 8.85 | 14.13 | 11.30 |
| ENSMUSG00000096205 | 8.78 | 15.71 | 18.62 | 10.34 | 27.00 | 6.26 | 5.40 | 7.40 | 6.87 | 14.38 | 14.13 | 10.36 |
| ENSMUSG00000094826 | 8.78 | 14.73 | 19.60 | 10.34 | 27.00 | 6.26 | 5.40 | 7.40 | 6.87 | 14.38 | 14.13 | 10.36 |
| ENSMUSG00000093815 | 8.78 | 15.71 | 19.60 | 10.34 | 27.00 | 6.26 | 5.40 | 7.40 | 6.87 | 14.38 | 14.13 | 10.36 |
| ENSMUSG00000095969 | 8.78 | 15.71 | 19.60 | 10.34 | 27.00 | 6.26 | 5.40 | 7.40 | 6.87 | 14.38 | 14.13 | 10.36 |
| ENSMUSG00000096214 | 8.78 | 15.71 | 19.60 | 10.34 | 27.00 | 6.26 | 5.40 | 7.40 | 6.87 | 14.38 | 14.13 | 10.36 |
| ENSMUSG00000005716 | 105.31 | 51.05 | 77.42 | 79.59 | 52.92 | 59.50 | 45.86 | 42.78 | 35.34 | 78.53 | 88.82 | 79.12 |
| ENSMUSG00000003477 | 59.48 | 18.65 | 76.44 | 35.15 | 24.84 | 31.32 | 19.78 | 13.99 | 13.74 | 26.55 | 37.34 | 31.08 |
| ENSMUSG00000104406 | 1.95 | 1.96 | 0.98 | 2.07 | 5.40 | 3.13 | 0.90 | 1.65 | 0.98 | 2.21 | 2.02 | 2.83 |
| ENSMUSG00000098749 | 3.90 | 2.95 | 9.80 | 1.03 | 11.88 | 6.26 | 2.70 | 2.47 | 3.93 | 7.74 | 5.05 | 5.65 |
| ENSMUSG00000093447 | 1.95 | 3.93 | 2.94 | 2.07 | 6.48 | 4.18 | 4.50 | 2.47 | 2.94 | 6.64 | 8.07 | 5.65 |
| ENSMUSG00000104125 | 0.98 | 2.95 | 3.92 | 2.07 | 1.08 | 1.04 | 1.80 | 1.65 | 1.96 | 3.32 | 4.04 | 3.77 |
| ENSMUSG00000068011 | 5.85 | 7.85 | 7.84 | 12.40 | 3.24 | 5.22 | 6.29 | 4.11 | 8.83 | 14.38 | 13.12 | 12.24 |
| ENSMUSG00000080764 | 2.93 | 5.89 | 4.90 | 9.30 | 5.40 | 7.31 | 1.80 | 4.94 | 2.94 | 7.74 | 5.05 | 7.54 |
| ENSMUSG00000081866 | 0.98 | 0.98 | 1.96 | 4.13 | 2.16 | 1.04 | 1.80 | 2.47 | 0.98 | 3.32 | 3.03 | 4.71 |
| ENSMUSG00000000544 | 6.83 | 6.87 | 4.90 | 9.30 | 9.72 | 3.13 | 3.60 | 4.11 | 3.93 | 7.74 | 7.06 | 10.36 |
| ENSMUSG00000085639 | 5.85 | 1.96 | 2.94 | 2.07 | 4.32 | 1.04 | 0.90 | 0.82 | 1.96 | 3.32 | 2.02 | 2.83 |
| ENSMUSG00000050751 | 3.90 | 1.96 | 3.92 | 2.07 | 2.16 | 1.04 | 1.80 | 2.47 | 2.94 | 4.42 | 6.06 | 5.65 |
| ENSMUSG00000098309 | 4.88 | 0.98 | 0.98 | 1.03 | 4.32 | 4.18 | 1.80 | 1.65 | 0.98 | 3.32 | 4.04 | 2.83 |
| ENSMUSG00000098330 | 4.88 | 0.98 | 0.98 | 1.03 | 4.32 | 4.18 | 1.80 | 1.65 | 0.98 | 3.32 | 4.04 | 2.83 |
| ENSMUSG00000098382 | 4.88 | 0.98 | 0.98 | 1.03 | 4.32 | 4.18 | 1.80 | 1.65 | 0.98 | 3.32 | 4.04 | 2.83 |
| ENSMUSG00000090273 | 7.80 | 1.96 | 2.94 | 2.07 | 4.32 | 7.31 | 1.80 | 2.47 | 1.96 | 5.53 | 5.05 | 3.77 |
| ENSMUSG00000074766 | 22.43 | 10.80 | 23.52 | 16.54 | 15.12 | 20.88 | 8.99 | 10.69 | 9.82 | 25.44 | 22.20 | 20.72 |
| ENSMUSG00000081747 | 5.85 | 2.95 | 3.92 | 5.17 | 6.48 | 4.18 | 3.60 | 1.65 | 2.94 | 6.64 | 6.06 | 6.59 |
| ENSMUSG00000082847 | 1.95 | 3.93 | 1.96 | 3.10 | 2.16 | 3.13 | 2.70 | 0.82 | 2.94 | 5.53 | 5.05 | 4.71 |
| ENSMUSG00000022512 | 2.93 | 7.85 | 7.84 | 7.24 | 5.40 | 3.13 | 2.70 | 3.29 | 2.94 | 7.74 | 6.06 | 7.54 |
| ENSMUSG00000091184 | 4.88 | 5.89 | 3.92 | 6.20 | 5.40 | 2.09 | 2.70 | 2.47 | 1.96 | 5.53 | 7.06 | 5.65 |
| ENSMUSG00000083026 | 4.88 | 8.84 | 5.88 | 7.24 | 3.24 | 4.18 | 4.50 | 1.65 | 2.94 | 8.85 | 6.06 | 8.48 |
| ENSMUSG00000049892 | 1.95 | 2.95 | 0.98 | 1.03 | 1.08 | 4.18 | 0.90 | 0.82 | 2.94 | 3.32 | 4.04 | 4.71 |
| ENSMUSG00000106051 | 8.78 | 3.93 | 11.76 | 13.44 | 17.28 | 6.26 | 6.29 | 4.94 | 4.91 | 15.49 | 12.11 | 14.13 |
| ENSMUSG00000081719 | 4.88 | 8.84 | 3.92 | 5.17 | 4.32 | 1.04 | 1.80 | 4.94 | 4.91 | 9.95 | 8.07 | 12.24 |
| ENSMUSG00000079015 | 5.85 | 0.98 | 2.94 | 1.03 | 6.48 | 3.13 | 1.80 | 1.65 | 1.96 | 5.53 | 4.04 | 4.71 |
| ENSMUSG00000036168 | 4.88 | 6.87 | 3.92 | 3.10 | 10.80 | 7.31 | 2.70 | 4.11 | 0.98 | 5.53 | 7.06 | 8.48 |

| | | | | | | | | | | | | |
|---|---|---|---|---|---|---|---|---|---|---|---|---|
| ENSMUSG00000101595 | 0.98 | 0.98 | 4.90 | 3.10 | 2.16 | 2.09 | 1.80 | 0.82 | 0.98 | 2.21 | 4.04 | 3.77 |
| ENSMUSG00000056492 | 15.60 | 17.67 | 13.72 | 28.94 | 20.52 | 20.88 | 8.99 | 7.40 | 11.78 | 19.91 | 34.32 | 28.26 |
| ENSMUSG00000031886 | 7.80 | 4.91 | 10.78 | 5.17 | 7.56 | 5.22 | 1.80 | 2.47 | 2.94 | 8.85 | 7.06 | 6.59 |
| ENSMUSG00000071178 | 4.88 | 0.98 | 2.94 | 1.03 | 5.40 | 3.13 | 1.80 | 1.65 | 0.98 | 5.53 | 4.04 | 4.71 |
| ENSMUSG00000066366 | 5.85 | 0.98 | 2.94 | 1.03 | 6.48 | 3.13 | 1.80 | 1.65 | 0.98 | 5.53 | 4.04 | 4.71 |
| ENSMUSG00000107735 | 1.95 | 2.95 | 3.92 | 7.24 | 2.16 | 4.18 | 1.80 | 0.82 | 0.98 | 4.42 | 3.03 | 4.71 |
| ENSMUSG00000089853 | 0.98 | 1.96 | 0.98 | 2.07 | 1.08 | 4.18 | 1.80 | 0.82 | 0.98 | 5.53 | 4.04 | 2.83 |
| ENSMUSG00000081831 | 4.88 | 6.87 | 6.86 | 6.20 | 2.16 | 3.13 | 2.70 | 2.47 | 2.94 | 12.17 | 9.08 | 7.54 |
| ENSMUSG00000086119 | 9.75 | 8.84 | 15.68 | 11.37 | 10.80 | 8.35 | 2.70 | 1.65 | 5.89 | 12.17 | 13.12 | 11.30 |
| ENSMUSG00000082495 | 0.98 | 4.91 | 1.96 | 5.17 | 3.24 | 3.13 | 1.80 | 0.82 | 0.98 | 3.32 | 4.04 | 5.65 |
| ENSMUSG00000082159 | 2.93 | 4.91 | 0.98 | 2.07 | 7.56 | 3.13 | 0.90 | 0.82 | 1.96 | 5.53 | 4.04 | 3.77 |
| ENSMUSG00000071553 | 2.93 | 0.98 | 0.98 | 5.17 | 1.08 | 1.04 | 1.80 | 0.82 | 0.98 | 4.42 | 4.04 | 5.65 |
| ENSMUSG00000106044 | 3.90 | 4.91 | 6.86 | 3.10 | 1.08 | 5.22 | 0.90 | 0.82 | 1.96 | 7.74 | 8.07 | 4.71 |
| ENSMUSG00000015120 | 4192.74 | 4227.39 | 4173.91 | 4270.21 | 4286.63 | 4307.93 | 4064.59 | 4077.20 | 3966.55 | 4018.50 | 3966.48 | 3899.38 |
| ENSMUSG00000024176 | 398.80 | 391.72 | 404.75 | 407.28 | 412.57 | 419.62 | 443.33 | 429.44 | 437.78 | 403.73 | 368.39 | 425.73 |
| ENSMUSG00000029552 | 1848.71 | 1845.68 | 1880.66 | 1972.30 | 1981.85 | 1908.14 | 1877.63 | 1859.26 | 1912.11 | 1794.11 | 1826.80 | 1876.23 |
| ENSMUSG00000027381 | 304.22 | 316.12 | 308.71 | 320.45 | 333.73 | 332.99 | 303.95 | 255.03 | 269.93 | 297.54 | 291.68 | 293.87 |
| ENSMUSG00000032477 | 373.45 | 356.37 | 366.53 | 381.44 | 395.29 | 393.53 | 317.43 | 387.48 | 361.22 | 366.12 | 335.08 | 305.17 |
| ENSMUSG00000020250 | 2792.56 | 2845.10 | 2749.94 | 2886.09 | 3067.28 | 3014.61 | 2814.64 | 2797.11 | 2914.30 | 2809.52 | 2811.86 | 2932.07 |
| ENSMUSG00000027531 | 808.32 | 818.78 | 835.96 | 877.61 | 882.38 | 874.74 | 838.10 | 843.25 | 815.69 | 807.46 | 735.77 | 854.29 |
| ENSMUSG00000001870 | 3177.71 | 3193.62 | 3190.95 | 3373.99 | 3390.21 | 3500.00 | 3632.06 | 3491.45 | 3669.13 | 3709.89 | 3761.59 | 3653.55 |
| ENSMUSG00000027639 | 1126.19 | 1125.08 | 1145.64 | 1211.50 | 1190.19 | 1249.48 | 1089.89 | 1038.22 | 1097.40 | 1162.52 | 1191.96 | 1074.69 |
| ENSMUSG00000074749 | 868.78 | 933.64 | 899.66 | 965.48 | 950.42 | 988.52 | 957.70 | 937.85 | 893.23 | 910.33 | 966.89 | 943.76 |
| ENSMUSG00000092376 | 0.98 | 0.98 | 0.98 | 1.03 | 1.08 | 1.04 | 0.90 | 0.82 | 0.98 | 1.11 | 1.01 | 3.77 |
| ENSMUSG00000061490 | 0.98 | 0.98 | 0.98 | 1.03 | 1.08 | 1.04 | 2.70 | 0.82 | 0.98 | 1.11 | 2.02 | 5.65 |
| ENSMUSG00000028602 | 2.93 | 2.95 | 2.94 | 3.10 | 3.24 | 3.13 | 1.80 | 4.94 | 3.93 | 3.32 | 2.02 | 3.77 |
| ENSMUSG00000042298 | 827.82 | 869.83 | 844.78 | 880.71 | 937.46 | 923.80 | 834.50 | 777.43 | 819.62 | 872.72 | 854.86 | 846.75 |
| ENSMUSG00000030347 | 448.53 | 476.15 | 476.29 | 510.65 | 504.37 | 496.87 | 465.81 | 455.76 | 486.86 | 464.57 | 478.40 | 466.23 |
| ENSMUSG00000004127 | 413.42 | 439.82 | 444.93 | 455.86 | 480.61 | 466.60 | 384.88 | 424.50 | 431.89 | 440.23 | 436.01 | 468.11 |
| ENSMUSG00000005370 | 948.73 | 1022.00 | 996.68 | 1049.20 | 1099.47 | 1075.16 | 934.32 | 939.50 | 1028.69 | 1010.98 | 934.60 | 994.63 |
| ENSMUSG00000053080 | 664.99 | 618.50 | 663.47 | 685.34 | 705.26 | 725.47 | 606.09 | 656.50 | 632.13 | 661.45 | 602.54 | 585.85 |
| ENSMUSG00000045969 | 903.88 | 936.58 | 918.28 | 1037.83 | 967.71 | 997.91 | 1051.22 | 1009.43 | 1051.27 | 1029.79 | 1056.72 | 984.26 |
| ENSMUSG00000030447 | 3135.78 | 3346.77 | 3280.13 | 3494.94 | 3485.25 | 3651.35 | 3346.10 | 3113.02 | 3575.88 | 3389.12 | 3418.44 | 3523.57 |
| ENSMUSG00000003778 | 1247.10 | 1309.65 | 1327.93 | 1360.35 | 1407.28 | 1473.90 | 1294.02 | 1277.62 | 1387.95 | 1347.24 | 1376.66 | 1301.68 |

| | | | | | | | | | | | | |
|---|---|---|---|---|---|---|---|---|---|---|---|---|
| ENSMUSG00000026867 | 2214.35 | 2273.72 | 2379.49 | 2422.99 | 2490.54 | 2614.82 | 2392.89 | 1980.19 | 2492.22 | 2402.47 | 2372.82 | 2353.76 |
| ENSMUSG00000058392 | 536.28 | 513.45 | 527.25 | 595.41 | 562.69 | 572.02 | 551.24 | 486.20 | 537.90 | 514.34 | 519.78 | 496.37 |
| ENSMUSG00000026430 | 317.87 | 299.43 | 295.97 | 343.19 | 333.73 | 326.72 | 342.61 | 301.10 | 342.57 | 321.88 | 330.04 | 310.82 |
| ENSMUSG00000024370 | 1301.70 | 1392.11 | 1368.11 | 1421.34 | 1494.76 | 1552.19 | 1385.74 | 1400.20 | 1479.23 | 1549.66 | 1342.34 | 1361.02 |
| ENSMUSG00000038418 | 2585.85 | 2753.79 | 2539.23 | 2878.85 | 2865.31 | 2951.98 | 2318.26 | 2461.46 | 2442.16 | 2842.70 | 2519.17 | 2823.76 |
| ENSMUSG00000031755 | 554.81 | 534.07 | 530.19 | 588.17 | 581.06 | 617.95 | 579.11 | 563.54 | 587.96 | 567.43 | 575.29 | 538.76 |
| ENSMUSG00000022765 | 738.12 | 799.14 | 812.44 | 866.24 | 839.18 | 889.35 | 776.95 | 730.54 | 879.49 | 801.93 | 859.91 | 886.31 |
| ENSMUSG00000020263 | 744.94 | 811.90 | 795.78 | 877.61 | 835.94 | 889.35 | 716.70 | 787.30 | 849.06 | 910.33 | 753.93 | 810.02 |
| ENSMUSG00000033450 | 204.76 | 199.29 | 219.52 | 236.72 | 233.29 | 220.25 | 229.31 | 225.41 | 241.47 | 220.12 | 190.75 | 243.95 |
| ENSMUSG00000024948 | 211.59 | 220.89 | 218.54 | 242.92 | 247.33 | 230.69 | 209.52 | 211.43 | 222.82 | 255.51 | 212.96 | 221.34 |
| ENSMUSG00000079215 | 1600.07 | 1753.40 | 1718.96 | 1805.87 | 1895.45 | 1918.58 | 1684.29 | 1585.30 | 1759.96 | 1668.01 | 1603.75 | 1782.04 |
| ENSMUSG00000024240 | 654.26 | 718.64 | 673.27 | 763.90 | 743.06 | 769.31 | 689.72 | 686.94 | 741.09 | 737.78 | 780.17 | 690.40 |
| ENSMUSG00000037997 | 201.84 | 211.08 | 220.50 | 234.65 | 230.05 | 240.08 | 196.04 | 199.91 | 227.73 | 212.37 | 182.68 | 210.98 |
| ENSMUSG00000045817 | 2479.57 | 2538.79 | 2657.82 | 2718.63 | 2977.64 | 2848.64 | 2380.30 | 2533.03 | 2509.89 | 2723.24 | 2407.14 | 2457.37 |
| ENSMUSG00000061533 | 361.75 | 366.19 | 383.19 | 402.11 | 403.93 | 431.11 | 428.04 | 401.47 | 473.12 | 367.23 | 378.48 | 395.59 |
| ENSMUSG00000033083 | 153.08 | 165.91 | 169.54 | 178.83 | 178.20 | 187.89 | 154.67 | 143.15 | 150.18 | 161.49 | 173.60 | 141.28 |
| ENSMUSG00000038543 | 135.53 | 131.55 | 139.16 | 152.99 | 146.88 | 155.53 | 121.40 | 145.61 | 130.55 | 136.05 | 135.24 | 131.86 |
| ENSMUSG00000030279 | 653.29 | 712.75 | 702.67 | 761.84 | 778.70 | 780.79 | 767.96 | 693.52 | 753.85 | 730.03 | 779.17 | 756.33 |
| ENSMUSG00000030515 | 373.45 | 404.48 | 403.77 | 419.68 | 444.97 | 461.38 | 391.17 | 361.16 | 372.02 | 388.24 | 445.09 | 387.11 |
| ENSMUSG00000040620 | 642.56 | 667.59 | 640.93 | 712.22 | 736.58 | 743.21 | 604.29 | 627.70 | 682.19 | 655.92 | 659.06 | 648.96 |
| ENSMUSG00000089764 | 3903.15 | 3992.76 | 3726.04 | 4431.47 | 4204.55 | 4446.76 | 3955.78 | 3864.95 | 4613.40 | 4546.11 | 4445.89 | 4645.35 |
| ENSMUSG00000040990 | 2485.42 | 2639.91 | 2567.65 | 2735.17 | 3003.56 | 2935.28 | 2602.42 | 2561.00 | 2659.09 | 2596.04 | 2687.72 | 2744.64 |
| ENSMUSG00000023284 | 351.02 | 369.14 | 379.27 | 389.70 | 430.93 | 419.62 | 347.11 | 354.57 | 355.33 | 411.47 | 396.65 | 340.96 |
| ENSMUSG00000022617 | 681.56 | 631.26 | 681.11 | 750.47 | 760.34 | 739.04 | 623.18 | 686.94 | 537.90 | 581.81 | 565.20 | 575.49 |
| ENSMUSG00000099689 | 175.51 | 176.71 | 194.04 | 209.84 | 207.37 | 199.37 | 202.33 | 171.12 | 189.44 | 186.93 | 186.72 | 160.12 |
| ENSMUSG00000048916 | 55.58 | 60.87 | 58.80 | 64.09 | 66.96 | 66.81 | 71.94 | 69.11 | 63.80 | 67.47 | 82.76 | 56.51 |
| ENSMUSG00000024045 | 1012.11 | 1063.23 | 1082.92 | 1156.71 | 1176.15 | 1233.82 | 1055.72 | 1108.15 | 1091.51 | 1150.35 | 1066.81 | 1123.66 |
| ENSMUSG00000032329 | 744.94 | 817.79 | 784.02 | 911.72 | 832.70 | 906.05 | 749.97 | 801.29 | 854.95 | 814.10 | 793.30 | 868.41 |
| ENSMUSG00000105156 | 249.61 | 253.29 | 266.57 | 283.23 | 290.53 | 297.49 | 285.06 | 312.62 | 329.81 | 293.12 | 317.92 | 310.82 |
| ENSMUSG00000058816 | 198.91 | 212.06 | 209.72 | 249.12 | 224.65 | 229.64 | 262.58 | 227.06 | 273.86 | 240.03 | 265.44 | 271.26 |
| ENSMUSG00000020160 | 348.10 | 345.57 | 346.93 | 376.27 | 410.41 | 395.62 | 340.81 | 365.27 | 366.13 | 344.00 | 378.48 | 368.28 |
| ENSMUSG00000021500 | 2059.32 | 2247.21 | 2246.21 | 2358.90 | 2474.34 | 2621.08 | 2241.82 | 2083.02 | 2367.56 | 2273.06 | 2246.66 | 2353.76 |
| ENSMUSG00000033565 | 5546.12 | 6029.88 | 5806.62 | 6164.98 | 6766.38 | 6872.64 | 5970.10 | 5993.22 | 6417.54 | 6445.30 | 6007.25 | 6355.81 |
| ENSMUSG00000045409 | 397.82 | 396.62 | 365.55 | 453.79 | 413.65 | 459.29 | 365.09 | 382.55 | 389.69 | 377.18 | 354.26 | 411.60 |

| | | | | | | | | | | | | |
|---|---|---|---|---|---|---|---|---|---|---|---|---|
| ENSMUSG00000022144 | 104.33 | 105.05 | 110.74 | 116.81 | 123.12 | 126.30 | 92.62 | 136.56 | 127.60 | 138.26 | 99.92 | 110.20 |
| ENSMUSG00000032221 | 117.98 | 133.52 | 128.38 | 137.48 | 152.28 | 145.09 | 121.40 | 144.79 | 158.03 | 136.05 | 141.30 | 119.62 |
| ENSMUSG00000004530 | 1929.64 | 1843.72 | 1819.90 | 2163.53 | 2125.50 | 2120.04 | 1972.95 | 2213.83 | 2219.34 | 2149.17 | 2243.63 | 2122.06 |
| ENSMUSG00000080715 | 307.14 | 348.52 | 330.27 | 361.79 | 400.69 | 367.43 | 334.52 | 299.46 | 284.66 | 296.44 | 326.00 | 306.11 |
| ENSMUSG00000021752 | 252.54 | 237.58 | 259.71 | 271.86 | 302.41 | 287.06 | 291.36 | 256.68 | 266.01 | 271.00 | 268.47 | 294.81 |
| ENSMUSG00000034333 | 768.34 | 770.67 | 786.96 | 887.95 | 892.10 | 892.48 | 846.19 | 788.13 | 888.33 | 838.43 | 845.78 | 903.26 |
| ENSMUSG00000056313 | 104.33 | 94.25 | 92.12 | 112.67 | 114.48 | 107.52 | 107.91 | 100.37 | 97.18 | 102.87 | 86.80 | 89.48 |
| ENSMUSG00000085385 | 256.44 | 292.56 | 269.51 | 303.91 | 321.85 | 317.33 | 208.63 | 274.77 | 235.58 | 268.78 | 201.86 | 245.83 |
| ENSMUSG00000047221 | 148.21 | 158.06 | 150.92 | 177.80 | 181.44 | 169.10 | 177.15 | 153.84 | 159.02 | 158.17 | 179.65 | 178.96 |
| ENSMUSG00000040102 | 447.55 | 493.82 | 488.05 | 513.75 | 587.54 | 555.32 | 451.42 | 430.26 | 457.41 | 478.95 | 469.32 | 461.52 |
| ENSMUSG00000038437 | 647.44 | 662.68 | 659.55 | 770.11 | 763.58 | 749.48 | 661.85 | 748.64 | 733.24 | 765.43 | 734.76 | 707.35 |
| ENSMUSG00000106870 | 42.90 | 48.11 | 45.08 | 49.62 | 52.92 | 55.32 | 32.37 | 31.26 | 42.21 | 40.93 | 27.25 | 35.79 |
| ENSMUSG00000027695 | 485.58 | 511.49 | 509.61 | 562.33 | 585.38 | 601.25 | 523.36 | 497.72 | 563.42 | 514.34 | 540.98 | 516.15 |
| ENSMUSG00000032580 | 2195.83 | 2542.72 | 2447.11 | 2751.71 | 2664.43 | 2936.32 | 2605.12 | 2288.69 | 2599.21 | 2723.24 | 2448.52 | 2603.36 |
| ENSMUSG00000017491 | 41.93 | 38.29 | 39.20 | 45.48 | 45.36 | 48.02 | 34.17 | 31.26 | 30.43 | 18.80 | 32.30 | 48.98 |
| ENSMUSG00000080989 | 309.09 | 338.70 | 316.55 | 356.63 | 382.33 | 384.13 | 392.07 | 332.36 | 376.92 | 327.41 | 404.72 | 366.39 |
| ENSMUSG00000072594 | 177.46 | 196.35 | 185.22 | 221.21 | 224.65 | 205.64 | 188.84 | 207.32 | 186.50 | 152.64 | 164.51 | 198.74 |
| ENSMUSG00000072812 | 1244.17 | 1394.08 | 1315.19 | 1585.69 | 1432.12 | 1596.03 | 1319.19 | 1267.75 | 1365.37 | 1360.52 | 1241.42 | 1337.47 |
| ENSMUSG00000001630 | 482.65 | 499.71 | 486.09 | 559.23 | 597.26 | 559.50 | 597.10 | 539.68 | 566.37 | 501.07 | 540.98 | 496.37 |
| ENSMUSG00000025316 | 626.96 | 567.45 | 599.77 | 680.17 | 735.50 | 684.76 | 614.19 | 622.77 | 605.63 | 648.18 | 623.74 | 632.00 |
| ENSMUSG00000062991 | 763.47 | 879.64 | 787.94 | 913.79 | 946.10 | 986.43 | 832.70 | 715.73 | 868.69 | 847.28 | 759.99 | 839.22 |
| ENSMUSG00000046179 | 748.84 | 866.88 | 784.02 | 931.36 | 883.46 | 997.91 | 840.80 | 816.10 | 948.20 | 874.93 | 829.63 | 897.61 |
| ENSMUSG00000021639 | 492.40 | 562.54 | 560.57 | 586.11 | 655.58 | 653.44 | 494.59 | 482.91 | 500.60 | 505.49 | 520.79 | 534.05 |
| ENSMUSG00000021431 | 496.30 | 496.76 | 540.97 | 580.94 | 638.30 | 580.38 | 494.59 | 505.95 | 480.97 | 474.52 | 483.45 | 463.41 |
| ENSMUSG00000040359 | 1038.43 | 1135.88 | 1108.40 | 1195.99 | 1381.36 | 1282.88 | 1153.73 | 949.37 | 1184.76 | 1119.38 | 1110.21 | 1115.19 |
| ENSMUSG00000025586 | 223.29 | 250.34 | 248.93 | 264.63 | 302.41 | 282.88 | 231.11 | 201.56 | 266.01 | 234.50 | 188.74 | 234.53 |
| ENSMUSG00000096435 | 294.47 | 288.63 | 282.25 | 325.62 | 330.49 | 362.21 | 293.15 | 309.33 | 333.74 | 300.86 | 330.04 | 315.53 |
| ENSMUSG00000096791 | 294.47 | 288.63 | 282.25 | 325.62 | 330.49 | 362.21 | 293.15 | 309.33 | 333.74 | 300.86 | 330.04 | 315.53 |
| ENSMUSG00000055065 | 4385.80 | 5059.91 | 5081.41 | 5422.79 | 5653.95 | 6027.13 | 4417.10 | 4543.66 | 4597.70 | 5252.92 | 4214.76 | 4828.08 |
| ENSMUSG00000004815 | 166.73 | 193.40 | 189.14 | 215.01 | 220.33 | 211.90 | 166.36 | 175.23 | 174.72 | 181.40 | 189.75 | 177.07 |
| ENSMUSG00000097772 | 83.85 | 82.47 | 84.28 | 92.00 | 99.36 | 104.38 | 88.13 | 95.43 | 76.56 | 76.32 | 82.76 | 62.16 |
| ENSMUSG00000090098 | 96.53 | 99.16 | 101.92 | 116.81 | 122.04 | 112.73 | 102.51 | 75.69 | 108.95 | 108.40 | 87.81 | 100.78 |
| ENSMUSG00000024384 | 1400.18 | 1500.11 | 1608.21 | 1712.84 | 1829.57 | 1783.92 | 1619.54 | 1554.86 | 1649.05 | 1724.43 | 1676.42 | 1697.27 |
| ENSMUSG00000026784 | 158.93 | 147.26 | 138.18 | 168.49 | 181.44 | 175.37 | 125.00 | 152.20 | 136.44 | 151.54 | 138.27 | 142.22 |

| | | | | | | | | | | | |
|---|---|---|---|---|---|---|---|---|---|---|---|
| ENSMUSG00000035455 | 386.12 | 435.89 | 397.89 | 444.49 | 495.73 | 502.09 | 399.27 | 382.55 | 419.13 | 384.93 | 308.84 | 356.97 |
| ENSMUSG00000053470 | 1384.58 | 1599.26 | 1509.23 | 1646.68 | 1805.81 | 1863.25 | 1504.44 | 1458.61 | 1581.32 | 1538.60 | 1541.17 | 1617.21 |
| ENSMUSG00000049349 | 45.83 | 51.05 | 48.02 | 58.92 | 56.16 | 56.37 | 58.45 | 37.84 | 35.34 | 57.52 | 37.34 | 61.22 |
| ENSMUSG00000108858 | 16.58 | 16.69 | 14.70 | 19.64 | 18.36 | 18.79 | 10.79 | 20.57 | 21.59 | 18.80 | 17.16 | 8.48 |
| ENSMUSG00000045795 | 179.41 | 161.99 | 171.50 | 212.94 | 190.08 | 204.59 | 151.07 | 179.34 | 175.70 | 155.96 | 177.63 | 175.19 |
| ENSMUSG00000048271 | 1103.76 | 1288.05 | 1209.35 | 1428.57 | 1329.51 | 1509.39 | 1330.88 | 1190.42 | 1386.97 | 1424.67 | 1245.45 | 1426.01 |
| ENSMUSG00000038894 | 333.47 | 299.43 | 323.41 | 359.73 | 362.89 | 411.27 | 319.23 | 292.87 | 410.30 | 335.15 | 328.02 | 350.38 |
| ENSMUSG00000039117 | 394.90 | 406.44 | 409.65 | 472.40 | 482.77 | 482.25 | 399.27 | 403.94 | 386.74 | 381.61 | 398.67 | 408.78 |
| ENSMUSG00000026643 | 1096.94 | 1227.18 | 1139.76 | 1256.98 | 1418.08 | 1440.50 | 1124.06 | 1171.50 | 1251.51 | 1142.61 | 1129.39 | 1189.59 |
| ENSMUSG00000083668 | 80.93 | 79.52 | 71.54 | 94.07 | 92.88 | 88.73 | 67.44 | 91.32 | 102.08 | 85.17 | 96.89 | 77.23 |
| ENSMUSG00000044617 | 299.34 | 316.12 | 265.59 | 335.95 | 362.89 | 348.64 | 301.25 | 300.28 | 350.42 | 336.26 | 304.80 | 349.44 |
| ENSMUSG00000015766 | 1851.63 | 2108.79 | 2214.85 | 2297.91 | 2544.55 | 2507.30 | 2115.93 | 2004.05 | 2335.17 | 2172.40 | 2058.94 | 2051.41 |
| ENSMUSG00000030126 | 939.95 | 1026.91 | 984.92 | 1071.95 | 1244.19 | 1198.33 | 1097.08 | 1015.19 | 1092.49 | 1129.34 | 1050.66 | 1077.51 |
| ENSMUSG00000027536 | 81.90 | 90.32 | 91.14 | 101.30 | 105.84 | 106.47 | 98.92 | 91.32 | 103.07 | 92.91 | 105.97 | 87.59 |
| ENSMUSG00000027667 | 604.53 | 659.73 | 642.89 | 707.05 | 798.14 | 766.18 | 669.94 | 662.26 | 639.99 | 685.79 | 620.71 | 668.73 |
| ENSMUSG00000020648 | 161.86 | 168.86 | 167.58 | 185.03 | 201.97 | 206.68 | 161.86 | 164.54 | 169.81 | 152.64 | 145.34 | 162.00 |
| ENSMUSG00000040282 | 166.73 | 153.15 | 174.44 | 179.86 | 208.45 | 201.46 | 169.06 | 187.57 | 190.43 | 163.70 | 173.60 | 161.06 |
| ENSMUSG00000037958 | 992.61 | 1110.35 | 1175.04 | 1280.75 | 1330.59 | 1311.06 | 1276.03 | 1260.34 | 1293.72 | 1226.68 | 1493.74 | 1217.85 |
| ENSMUSG00000042105 | 389.05 | 434.91 | 430.23 | 469.30 | 540.01 | 492.69 | 428.94 | 376.79 | 433.86 | 400.41 | 451.15 | 411.60 |
| ENSMUSG00000041096 | 695.22 | 767.72 | 736.00 | 923.09 | 814.34 | 899.79 | 859.68 | 750.28 | 830.41 | 819.63 | 842.75 | 858.05 |
| ENSMUSG00000027751 | 907.78 | 1018.07 | 946.70 | 1063.68 | 1213.95 | 1171.19 | 995.47 | 1036.58 | 1048.32 | 1090.62 | 968.91 | 1002.16 |
| ENSMUSG00000069094 | 577.23 | 689.18 | 648.77 | 714.29 | 795.98 | 789.14 | 704.11 | 682.00 | 746.98 | 723.40 | 609.61 | 692.28 |
| ENSMUSG00000037266 | 3261.56 | 3389.96 | 3526.11 | 3899.11 | 4212.11 | 4108.56 | 2944.13 | 3387.79 | 3135.15 | 3375.85 | 2802.78 | 2968.81 |
| ENSMUSG00000023951 | 2059.32 | 2417.06 | 2111.94 | 2717.60 | 2674.15 | 2528.18 | 2415.38 | 2670.42 | 2816.14 | 2538.52 | 2374.84 | 2709.79 |
| ENSMUSG00000038253 | 135.53 | 146.28 | 151.90 | 167.46 | 186.84 | 168.06 | 117.80 | 143.97 | 129.57 | 140.48 | 106.98 | 129.04 |
| ENSMUSG00000038002 | 499.23 | 557.63 | 530.19 | 586.11 | 652.34 | 673.28 | 518.87 | 499.37 | 592.87 | 553.06 | 532.90 | 549.12 |
| ENSMUSG00000094519 | 207.69 | 206.17 | 199.92 | 239.82 | 235.45 | 267.22 | 197.83 | 196.62 | 227.73 | 211.27 | 210.94 | 221.34 |
| ENSMUSG00000032235 | 302.27 | 288.63 | 313.61 | 353.53 | 379.09 | 362.21 | 305.74 | 248.45 | 286.62 | 304.18 | 296.73 | 278.80 |
| ENSMUSG00000096912 | 201.84 | 198.31 | 194.04 | 233.62 | 227.89 | 257.83 | 189.74 | 190.86 | 219.87 | 207.95 | 199.84 | 210.98 |
| ENSMUSG00000062519 | 353.95 | 359.32 | 346.93 | 413.48 | 455.77 | 414.40 | 327.33 | 301.10 | 355.33 | 382.71 | 343.16 | 378.64 |
| ENSMUSG00000097565 | 86.78 | 101.12 | 90.16 | 108.54 | 115.56 | 112.73 | 86.33 | 79.80 | 86.38 | 92.91 | 78.72 | 70.64 |
| ENSMUSG00000020185 | 347.12 | 399.57 | 374.37 | 443.46 | 438.49 | 479.12 | 342.61 | 371.03 | 446.62 | 438.02 | 313.89 | 409.72 |
| ENSMUSG00000020024 | 723.49 | 884.55 | 775.20 | 925.16 | 955.82 | 1012.53 | 792.24 | 720.67 | 822.56 | 869.40 | 813.48 | 742.20 |
| ENSMUSG00000042842 | 140.41 | 148.24 | 132.30 | 166.43 | 171.72 | 173.28 | 210.42 | 172.76 | 182.57 | 165.92 | 200.85 | 194.03 |

| | | | | | | | | | | | | |
|---|---|---|---|---|---|---|---|---|---|---|---|---|
| ENSMUSG00000039697 | 437.80 | 517.38 | 502.75 | 585.07 | 595.10 | 593.95 | 537.75 | 397.35 | 557.53 | 496.64 | 514.73 | 539.70 |
| ENSMUSG00000027635 | 243.76 | 261.14 | 267.55 | 299.77 | 298.09 | 342.38 | 246.39 | 238.58 | 291.53 | 296.44 | 253.33 | 259.96 |
| ENSMUSG00000034248 | 1724.87 | 2027.30 | 2037.46 | 2165.60 | 2422.50 | 2479.12 | 1887.52 | 1962.09 | 1961.19 | 2067.32 | 1635.04 | 1896.00 |
| ENSMUSG00000051890 | 193.06 | 224.82 | 202.86 | 265.66 | 246.25 | 246.35 | 227.51 | 231.17 | 204.17 | 207.95 | 213.97 | 222.28 |
| ENSMUSG00000046591 | 147.23 | 166.90 | 168.56 | 192.27 | 183.60 | 213.99 | 151.07 | 157.13 | 194.35 | 149.32 | 152.40 | 169.54 |
| ENSMUSG00000074384 | 170.63 | 186.53 | 188.16 | 207.77 | 230.05 | 228.60 | 166.36 | 174.41 | 185.52 | 182.51 | 168.55 | 178.96 |
| ENSMUSG00000026648 | 156.98 | 172.79 | 168.56 | 190.20 | 216.01 | 203.55 | 169.06 | 144.79 | 167.85 | 160.39 | 160.48 | 165.77 |
| ENSMUSG00000026196 | 185.26 | 184.57 | 184.24 | 207.77 | 232.21 | 238.00 | 166.36 | 208.96 | 212.02 | 220.12 | 190.75 | 179.90 |
| ENSMUSG00000033904 | 334.44 | 371.10 | 361.63 | 420.72 | 435.25 | 455.11 | 338.12 | 307.68 | 370.05 | 348.42 | 330.04 | 356.03 |
| ENSMUSG00000040195 | 254.49 | 235.62 | 240.11 | 278.07 | 294.85 | 324.63 | 253.59 | 258.32 | 267.97 | 245.56 | 223.05 | 240.18 |
| ENSMUSG00000031393 | 1096.94 | 1338.12 | 1261.29 | 1413.07 | 1555.24 | 1587.68 | 1236.46 | 1129.54 | 1279.97 | 1253.22 | 1254.54 | 1217.85 |
| ENSMUSG00000020070 | 375.40 | 450.62 | 409.65 | 460.00 | 531.37 | 532.36 | 417.25 | 371.03 | 451.52 | 450.19 | 370.41 | 433.26 |
| ENSMUSG00000033581 | 426.10 | 425.10 | 424.35 | 511.68 | 494.65 | 566.81 | 491.89 | 497.72 | 545.76 | 532.04 | 491.52 | 506.73 |
| ENSMUSG00000050064 | 342.24 | 379.94 | 347.91 | 422.78 | 438.49 | 459.29 | 392.97 | 386.66 | 396.56 | 419.22 | 395.64 | 402.18 |
| ENSMUSG00000042156 | 344.19 | 374.04 | 414.55 | 460.00 | 441.73 | 496.87 | 365.09 | 325.78 | 343.55 | 433.60 | 386.55 | 384.29 |
| ENSMUSG00000039191 | 1369.95 | 1496.18 | 1533.73 | 1661.16 | 1846.85 | 1936.32 | 1469.37 | 1272.69 | 1601.93 | 1523.11 | 1483.64 | 1578.59 |
| ENSMUSG00000041235 | 575.28 | 674.46 | 672.29 | 729.79 | 795.98 | 854.91 | 644.76 | 660.61 | 702.81 | 694.64 | 726.68 | 693.22 |
| ENSMUSG00000020899 | 346.14 | 329.87 | 341.05 | 410.38 | 421.21 | 430.06 | 349.81 | 401.47 | 416.19 | 410.37 | 345.17 | 403.12 |
| ENSMUSG00000024483 | 1600.07 | 1993.92 | 1765.02 | 2143.89 | 2106.05 | 2436.32 | 2007.12 | 1757.24 | 2321.43 | 2212.22 | 2197.21 | 2279.35 |
| ENSMUSG00000102550 | 1054.04 | 1237.00 | 1170.14 | 1330.37 | 1510.96 | 1492.69 | 1039.53 | 900.01 | 1142.55 | 1315.17 | 1237.38 | 1247.99 |
| ENSMUSG00000100798 | 38.03 | 36.32 | 33.32 | 40.31 | 48.60 | 45.93 | 35.07 | 53.47 | 34.36 | 45.35 | 41.38 | 32.02 |
| ENSMUSG00000069727 | 112.13 | 123.70 | 127.40 | 141.62 | 160.92 | 152.40 | 123.20 | 103.66 | 126.62 | 121.67 | 115.06 | 139.40 |
| ENSMUSG00000082743 | 117.01 | 111.92 | 117.60 | 136.45 | 158.76 | 138.83 | 151.97 | 142.32 | 183.55 | 175.87 | 168.55 | 199.68 |
| ENSMUSG00000019774 | 158.93 | 160.02 | 178.36 | 206.74 | 204.13 | 212.94 | 197.83 | 186.75 | 178.65 | 155.96 | 174.61 | 210.04 |
| ENSMUSG00000059645 | 139.43 | 148.24 | 148.96 | 182.96 | 192.24 | 173.28 | 139.38 | 149.73 | 142.33 | 164.81 | 131.21 | 116.79 |
| ENSMUSG00000027615 | 230.11 | 257.22 | 282.25 | 293.57 | 354.25 | 320.46 | 250.89 | 251.74 | 317.05 | 276.53 | 224.06 | 267.49 |
| ENSMUSG00000048521 | 65.33 | 62.83 | 66.64 | 79.59 | 82.08 | 83.51 | 145.68 | 136.56 | 92.27 | 80.75 | 107.99 | 95.13 |
| ENSMUSG00000004360 | 215.49 | 261.14 | 212.66 | 261.53 | 304.57 | 301.67 | 312.04 | 246.80 | 284.66 | 258.83 | 235.16 | 261.84 |
| ENSMUSG00000042097 | 101.41 | 117.81 | 115.64 | 128.18 | 149.04 | 145.09 | 129.49 | 132.45 | 149.20 | 123.88 | 117.08 | 142.22 |
| ENSMUSG00000034023 | 160.88 | 194.39 | 169.54 | 203.64 | 231.13 | 227.56 | 219.42 | 186.75 | 241.47 | 175.87 | 165.52 | 192.14 |
| ENSMUSG00000026743 | 623.06 | 666.60 | 649.75 | 732.89 | 855.38 | 862.21 | 666.34 | 610.43 | 690.05 | 696.85 | 637.87 | 671.56 |
| ENSMUSG00000068917 | 717.64 | 792.27 | 732.08 | 976.85 | 878.06 | 980.17 | 841.69 | 886.03 | 938.39 | 883.78 | 811.46 | 871.24 |
| ENSMUSG00000052125 | 249.61 | 299.43 | 277.35 | 330.78 | 374.77 | 342.38 | 291.36 | 344.70 | 322.94 | 334.05 | 268.47 | 258.08 |
| ENSMUSG00000021217 | 1020.88 | 1086.79 | 1116.24 | 1267.32 | 1359.76 | 1461.38 | 1091.68 | 970.76 | 1156.30 | 1187.96 | 1183.89 | 1179.23 |

| | | | | | | | | | | | | |
|---|---|---|---|---|---|---|---|---|---|---|---|---|
| ENSMUSG00000036036 | 205.74 | 229.73 | 227.36 | 302.87 | 286.21 | 251.57 | 248.19 | 292.05 | 271.90 | 263.25 | 228.10 | 255.25 |
| ENSMUSG00000037239 | 132.61 | 148.24 | 140.14 | 184.00 | 170.64 | 180.58 | 151.07 | 143.97 | 137.42 | 154.86 | 122.12 | 146.93 |
| ENSMUSG00000047735 | 489.48 | 604.75 | 538.03 | 623.32 | 757.10 | 695.20 | 499.98 | 478.80 | 612.50 | 529.83 | 498.59 | 538.76 |
| ENSMUSG00000030309 | 138.46 | 160.02 | 158.76 | 182.96 | 187.92 | 210.86 | 157.37 | 159.60 | 150.18 | 172.55 | 169.56 | 159.18 |
| ENSMUSG00000056919 | 483.63 | 601.81 | 583.11 | 653.30 | 684.74 | 784.97 | 536.85 | 524.87 | 684.16 | 657.03 | 596.49 | 578.31 |
| ENSMUSG00000020994 | 1464.53 | 1846.66 | 1757.18 | 1978.50 | 2298.30 | 2176.41 | 1788.60 | 1758.07 | 1956.28 | 1982.15 | 1874.24 | 1891.30 |
| ENSMUSG00000020601 | 160.88 | 192.42 | 170.52 | 198.47 | 231.13 | 238.00 | 208.63 | 181.81 | 186.50 | 199.10 | 217.00 | 190.26 |
| ENSMUSG00000039831 | 1772.65 | 2192.24 | 1901.24 | 2267.94 | 2526.19 | 2688.93 | 2183.37 | 1843.62 | 2198.73 | 2078.38 | 1965.07 | 2042.94 |
| ENSMUSG00000040459 | 1678.07 | 1834.88 | 1790.50 | 2076.70 | 2479.74 | 2212.94 | 1785.00 | 1776.16 | 1818.86 | 2009.80 | 1758.17 | 1754.72 |
| ENSMUSG00000036863 | 58.50 | 63.81 | 65.66 | 75.46 | 83.16 | 81.42 | 68.34 | 63.35 | 90.30 | 55.31 | 71.66 | 76.29 |
| ENSMUSG00000043336 | 3261.56 | 4111.55 | 3428.11 | 4356.01 | 4376.27 | 5106.47 | 4160.81 | 3512.02 | 4177.58 | 4101.46 | 4083.55 | 3882.43 |
| ENSMUSG00000033364 | 354.92 | 430.99 | 396.91 | 463.10 | 503.29 | 550.10 | 425.34 | 377.61 | 429.93 | 446.87 | 465.28 | 419.14 |
| ENSMUSG00000035545 | 724.47 | 889.46 | 727.18 | 1065.74 | 908.30 | 1028.18 | 898.35 | 923.87 | 938.39 | 956.79 | 764.03 | 872.18 |
| ENSMUSG00000055436 | 2229.95 | 2821.53 | 2796.98 | 3054.58 | 3572.73 | 3443.63 | 2840.72 | 2746.10 | 2809.27 | 2965.48 | 2765.43 | 2790.79 |
| ENSMUSG00000022723 | 163.81 | 156.10 | 149.94 | 189.17 | 191.16 | 223.38 | 164.56 | 157.95 | 165.89 | 174.77 | 178.64 | 166.71 |
| ENSMUSG00000089791 | 79.95 | 102.10 | 96.04 | 113.71 | 111.24 | 132.57 | 131.29 | 104.48 | 120.73 | 102.87 | 104.97 | 104.55 |
| ENSMUSG00000026098 | 152.11 | 195.37 | 184.24 | 209.84 | 243.01 | 231.73 | 175.35 | 169.47 | 157.05 | 178.08 | 176.62 | 166.71 |
| ENSMUSG00000027438 | 231.09 | 280.78 | 297.93 | 332.85 | 352.09 | 360.12 | 325.53 | 272.31 | 323.92 | 316.35 | 280.58 | 309.88 |
| ENSMUSG00000084808 | 10.73 | 9.82 | 8.82 | 11.37 | 12.96 | 13.57 | 8.09 | 11.52 | 7.85 | 11.06 | 9.08 | 7.54 |
| ENSMUSG00000078905 | 31.20 | 27.49 | 28.42 | 39.28 | 35.64 | 37.58 | 30.57 | 25.50 | 29.45 | 32.08 | 34.32 | 21.66 |
| ENSMUSG00000098506 | 164.78 | 186.53 | 198.94 | 247.05 | 224.65 | 240.08 | 231.11 | 209.78 | 242.45 | 210.16 | 245.26 | 234.53 |
| ENSMUSG00000108181 | 67.28 | 51.05 | 62.72 | 71.33 | 84.24 | 80.38 | 66.54 | 48.54 | 64.78 | 55.31 | 55.51 | 47.09 |
| ENSMUSG00000044703 | 13.65 | 15.71 | 11.76 | 18.61 | 16.20 | 18.79 | 12.59 | 7.40 | 7.85 | 5.53 | 13.12 | 11.30 |
| ENSMUSG00000053580 | 1827.26 | 2258.99 | 1881.64 | 2416.79 | 2473.26 | 2893.53 | 2289.48 | 1846.92 | 2412.71 | 2122.63 | 2198.22 | 2172.92 |
| ENSMUSG00000099750 | 15.60 | 12.76 | 12.74 | 17.57 | 19.44 | 16.70 | 10.79 | 16.45 | 12.76 | 18.80 | 11.10 | 16.95 |
| ENSMUSG00000103419 | 94.58 | 118.79 | 98.98 | 140.58 | 136.08 | 131.52 | 165.46 | 129.16 | 179.63 | 124.99 | 128.18 | 171.42 |
| ENSMUSG00000105013 | 52.65 | 54.98 | 46.06 | 67.19 | 66.96 | 66.81 | 49.46 | 60.06 | 70.67 | 70.79 | 45.42 | 62.16 |
| ENSMUSG00000090272 | 155.03 | 208.13 | 182.28 | 212.94 | 252.73 | 248.43 | 176.25 | 157.13 | 194.35 | 189.14 | 188.74 | 182.72 |
| ENSMUSG00000029068 | 1156.42 | 1392.11 | 1314.21 | 1569.16 | 1844.69 | 1648.22 | 1299.41 | 1327.80 | 1337.89 | 1400.34 | 1064.79 | 1213.14 |
| ENSMUSG00000097287 | 69.23 | 64.80 | 63.70 | 76.49 | 92.88 | 89.77 | 71.04 | 68.28 | 72.64 | 81.85 | 68.63 | 78.18 |
| ENSMUSG00000037355 | 426.10 | 531.12 | 476.29 | 567.50 | 671.78 | 641.96 | 471.21 | 399.00 | 474.10 | 536.46 | 418.85 | 522.74 |
| ENSMUSG00000097131 | 63.38 | 71.67 | 70.56 | 87.86 | 86.40 | 96.03 | 73.74 | 78.15 | 77.54 | 81.85 | 72.67 | 79.12 |
| ENSMUSG00000079165 | 94.58 | 87.38 | 85.26 | 130.25 | 111.24 | 110.65 | 102.51 | 127.52 | 102.08 | 100.66 | 81.75 | 106.43 |
| ENSMUSG00000050587 | 181.36 | 190.46 | 202.86 | 237.75 | 248.41 | 271.40 | 205.93 | 207.32 | 247.36 | 244.45 | 209.93 | 223.23 |

| | | | | | | | | | | | | |
|---|---|---|---|---|---|---|---|---|---|---|---|---|
| ENSMUSG00000046318 | 474.85 | 538.98 | 467.47 | 572.67 | 675.02 | 705.64 | 632.17 | 544.61 | 588.95 | 497.75 | 512.72 | 579.26 |
| ENSMUSG00000055228 | 103.36 | 116.83 | 118.58 | 134.38 | 163.08 | 149.27 | 89.92 | 77.33 | 112.88 | 119.46 | 77.71 | 91.36 |
| ENSMUSG00000036568 | 456.33 | 449.64 | 508.63 | 596.44 | 628.58 | 640.92 | 526.06 | 428.62 | 544.77 | 575.18 | 508.68 | 550.06 |
| ENSMUSG00000022822 | 1161.29 | 1401.93 | 1299.51 | 1723.18 | 1597.36 | 1791.23 | 1410.92 | 1209.34 | 1441.93 | 1463.38 | 1309.04 | 1481.58 |
| ENSMUSG00000040565 | 751.77 | 971.93 | 888.88 | 1025.43 | 1179.39 | 1262.00 | 914.53 | 738.77 | 1031.64 | 1006.56 | 874.04 | 981.44 |
| ENSMUSG00000040987 | 22.43 | 22.58 | 17.64 | 27.91 | 30.24 | 25.05 | 21.58 | 21.39 | 31.41 | 37.61 | 23.21 | 24.49 |
| ENSMUSG00000038331 | 211.59 | 247.40 | 241.09 | 274.96 | 324.01 | 330.90 | 244.59 | 233.64 | 293.49 | 283.16 | 255.35 | 260.90 |
| ENSMUSG00000089901 | 179.41 | 201.26 | 173.46 | 226.38 | 272.17 | 239.04 | 176.25 | 190.04 | 189.44 | 213.48 | 148.36 | 197.79 |
| ENSMUSG00000030424 | 36.08 | 39.27 | 46.06 | 56.85 | 49.68 | 55.32 | 49.46 | 43.60 | 43.19 | 43.14 | 41.38 | 52.75 |
| ENSMUSG00000040274 | 827.82 | 919.89 | 904.56 | 1033.70 | 1216.11 | 1287.05 | 809.32 | 890.14 | 929.55 | 1027.58 | 826.60 | 825.09 |
| ENSMUSG00000096449 | 1426.51 | 1779.90 | 1373.99 | 2252.43 | 2027.21 | 1848.64 | 2303.87 | 1739.97 | 2711.11 | 2780.76 | 2467.69 | 2626.90 |
| ENSMUSG00000003279 | 10.73 | 8.84 | 11.76 | 15.51 | 12.96 | 13.57 | 23.38 | 12.34 | 9.82 | 13.27 | 13.12 | 16.01 |
| ENSMUSG00000108456 | 38.03 | 36.32 | 40.18 | 46.52 | 54.00 | 53.24 | 35.07 | 36.20 | 50.06 | 37.61 | 35.32 | 41.44 |
| ENSMUSG00000039396 | 213.54 | 233.66 | 208.74 | 274.96 | 329.41 | 277.66 | 206.83 | 243.51 | 209.08 | 204.63 | 178.64 | 213.81 |
| ENSMUSG00000082852 | 6.83 | 6.87 | 5.88 | 8.27 | 9.72 | 8.35 | 9.89 | 6.58 | 7.85 | 9.95 | 14.13 | 11.30 |
| ENSMUSG00000026034 | 1918.91 | 2563.34 | 2370.67 | 2719.66 | 3389.13 | 3133.61 | 2344.33 | 2291.16 | 2435.29 | 2502.02 | 2102.33 | 2252.98 |
| ENSMUSG00000071723 | 90.68 | 115.85 | 95.06 | 140.58 | 130.68 | 135.70 | 101.61 | 102.83 | 104.05 | 96.23 | 85.79 | 86.65 |
| ENSMUSG00000059708 | 157.96 | 215.00 | 199.92 | 244.99 | 261.37 | 267.22 | 176.25 | 162.07 | 202.20 | 165.92 | 187.73 | 168.60 |
| ENSMUSG00000024524 | 12.68 | 11.78 | 13.72 | 15.51 | 18.36 | 17.75 | 21.58 | 18.92 | 26.50 | 13.27 | 18.17 | 15.07 |
| ENSMUSG00000027829 | 933.13 | 1281.18 | 1118.20 | 1367.58 | 1582.24 | 1558.45 | 1167.22 | 1148.46 | 1234.82 | 1183.54 | 1075.89 | 1106.71 |
| ENSMUSG00000032113 | 147.23 | 180.64 | 158.76 | 190.20 | 237.61 | 230.69 | 136.69 | 150.55 | 151.16 | 149.32 | 114.05 | 135.63 |
| ENSMUSG00000036377 | 80.93 | 80.50 | 81.34 | 107.50 | 103.68 | 117.95 | 79.13 | 76.51 | 85.40 | 92.91 | 75.70 | 111.14 |
| ENSMUSG00000037243 | 94.58 | 109.96 | 99.96 | 149.89 | 140.40 | 123.17 | 100.72 | 130.81 | 104.05 | 120.57 | 84.78 | 123.39 |
| ENSMUSG00000100178 | 29.25 | 21.60 | 22.54 | 32.04 | 35.64 | 32.36 | 25.18 | 25.50 | 34.36 | 27.65 | 25.23 | 31.08 |
| ENSMUSG00000037315 | 90.68 | 107.01 | 98.98 | 124.04 | 145.80 | 136.74 | 88.13 | 77.33 | 97.18 | 113.93 | 96.89 | 110.20 |
| ENSMUSG00000021732 | 66.30 | 53.01 | 65.66 | 83.73 | 85.32 | 84.55 | 63.85 | 58.41 | 79.51 | 60.84 | 74.69 | 69.70 |
| ENSMUSG00000034160 | 2654.10 | 3499.92 | 3255.63 | 3674.80 | 4552.32 | 4735.90 | 3558.32 | 2811.92 | 3394.29 | 3610.34 | 2999.59 | 3412.43 |
| ENSMUSG00000047227 | 55.58 | 47.12 | 49.98 | 65.12 | 73.44 | 72.02 | 53.95 | 46.89 | 53.99 | 57.52 | 41.38 | 44.27 |
| ENSMUSG00000082233 | 25.35 | 33.38 | 30.38 | 42.38 | 45.36 | 36.53 | 32.37 | 36.20 | 54.97 | 50.88 | 42.39 | 52.75 |
| ENSMUSG00000033022 | 6.83 | 5.89 | 6.86 | 9.30 | 8.64 | 9.39 | 7.19 | 10.69 | 7.85 | 14.38 | 8.07 | 10.36 |
| ENSMUSG00000024793 | 116.03 | 108.97 | 115.64 | 155.05 | 177.12 | 145.09 | 131.29 | 150.55 | 111.90 | 124.99 | 97.90 | 140.34 |
| ENSMUSG00000037818 | 161.86 | 215.98 | 172.48 | 235.68 | 287.29 | 254.70 | 169.06 | 152.20 | 176.68 | 184.72 | 178.64 | 186.49 |
| ENSMUSG00000090799 | 25.35 | 30.43 | 26.46 | 36.18 | 42.12 | 38.62 | 37.77 | 35.38 | 39.26 | 35.40 | 36.33 | 27.31 |
| ENSMUSG00000039529 | 48.75 | 68.72 | 56.84 | 71.33 | 92.88 | 84.55 | 60.25 | 47.72 | 76.56 | 69.68 | 60.56 | 62.16 |

| | | | | | | | | | | | | |
|---|---|---|---|---|---|---|---|---|---|---|---|---|
| ENSMUSG00000097041 | 21.45 | 18.65 | 19.60 | 28.94 | 28.08 | 28.18 | 37.77 | 22.21 | 33.37 | 28.76 | 37.34 | 32.97 |
| ENSMUSG00000089628 | 151.13 | 167.88 | 161.70 | 201.57 | 246.25 | 240.08 | 214.92 | 209.78 | 213.00 | 216.80 | 208.92 | 208.16 |
| ENSMUSG00000004187 | 30.23 | 38.29 | 31.36 | 47.55 | 49.68 | 45.93 | 25.18 | 34.55 | 35.34 | 27.65 | 25.23 | 29.20 |
| ENSMUSG00000052392 | 5.85 | 4.91 | 6.86 | 8.27 | 8.64 | 8.35 | 8.09 | 7.40 | 4.91 | 8.85 | 11.10 | 6.59 |
| ENSMUSG00000098099 | 12.68 | 10.80 | 13.72 | 19.64 | 18.36 | 15.66 | 21.58 | 22.21 | 24.54 | 22.12 | 26.24 | 29.20 |
| ENSMUSG00000098265 | 89.71 | 106.03 | 94.08 | 140.58 | 123.12 | 154.49 | 124.10 | 83.91 | 170.79 | 146.01 | 119.10 | 151.64 |
| ENSMUSG00000085547 | 16.58 | 21.60 | 24.50 | 25.84 | 32.40 | 32.36 | 25.18 | 20.57 | 39.26 | 33.18 | 27.25 | 40.50 |
| ENSMUSG00000040209 | 361.75 | 482.04 | 428.27 | 679.14 | 519.49 | 647.18 | 443.33 | 362.80 | 468.21 | 518.77 | 465.28 | 491.66 |
| ENSMUSG00000019971 | 400.75 | 524.25 | 495.89 | 594.38 | 713.90 | 757.83 | 549.44 | 410.52 | 556.55 | 535.36 | 520.79 | 497.31 |
| ENSMUSG00000081627 | 40.95 | 54.00 | 40.18 | 72.36 | 66.96 | 57.41 | 54.85 | 51.83 | 85.40 | 81.85 | 83.77 | 79.12 |
| ENSMUSG00000099102 | 19.50 | 13.74 | 18.62 | 26.88 | 27.00 | 21.92 | 20.68 | 20.57 | 26.50 | 29.86 | 28.26 | 22.61 |
| ENSMUSG00000030446 | 63.38 | 77.56 | 77.42 | 88.90 | 118.80 | 111.69 | 93.52 | 88.03 | 102.08 | 92.91 | 79.73 | 84.77 |
| ENSMUSG00000041247 | 17.55 | 21.60 | 20.58 | 26.88 | 33.48 | 27.14 | 14.39 | 24.68 | 32.39 | 25.44 | 27.25 | 27.31 |
| ENSMUSG00000085355 | 24.38 | 23.56 | 19.60 | 28.94 | 34.56 | 35.49 | 28.78 | 29.62 | 25.52 | 29.86 | 31.29 | 20.72 |
| ENSMUSG00000085396 | 293.49 | 407.42 | 377.31 | 451.73 | 546.49 | 583.51 | 348.01 | 361.16 | 356.31 | 405.94 | 257.37 | 359.80 |
| ENSMUSG00000048502 | 18.53 | 19.63 | 12.74 | 23.78 | 27.00 | 24.01 | 24.28 | 24.68 | 21.59 | 24.33 | 21.19 | 20.72 |
| ENSMUSG00000037033 | 42.90 | 33.38 | 44.10 | 57.89 | 63.72 | 56.37 | 53.06 | 46.89 | 53.99 | 53.09 | 51.47 | 52.75 |
| ENSMUSG00000084899 | 55.58 | 68.72 | 54.88 | 86.83 | 95.04 | 83.51 | 87.23 | 87.20 | 96.19 | 108.40 | 87.81 | 93.25 |
| ENSMUSG00000086370 | 154.06 | 179.66 | 178.36 | 211.91 | 268.93 | 280.79 | 187.94 | 187.57 | 191.41 | 188.04 | 148.36 | 145.99 |
| ENSMUSG00000038060 | 3.90 | 4.91 | 3.92 | 6.20 | 6.48 | 6.26 | 3.60 | 2.47 | 2.94 | 3.32 | 3.03 | 5.65 |
| ENSMUSG00000097571 | 31.20 | 36.32 | 34.30 | 52.72 | 54.00 | 44.89 | 44.96 | 46.89 | 38.28 | 35.40 | 24.22 | 39.56 |
| ENSMUSG00000072675 | 19.50 | 19.63 | 13.72 | 25.84 | 27.00 | 26.10 | 25.18 | 27.97 | 23.56 | 25.44 | 21.19 | 21.66 |
| ENSMUSG00000089659 | 56.55 | 53.01 | 37.24 | 77.53 | 66.96 | 75.16 | 67.44 | 83.91 | 77.54 | 75.22 | 75.70 | 65.93 |
| ENSMUSG00000058248 | 16.58 | 18.65 | 16.66 | 22.74 | 28.08 | 27.14 | 21.58 | 25.50 | 35.34 | 22.12 | 17.16 | 29.20 |
| ENSMUSG00000082957 | 15.60 | 13.74 | 14.70 | 19.64 | 21.60 | 25.05 | 17.09 | 14.81 | 22.58 | 29.86 | 14.13 | 15.07 |
| ENSMUSG00000099906 | 1320.23 | 1749.47 | 1616.05 | 2150.09 | 2734.63 | 2169.10 | 1908.20 | 1698.01 | 1860.08 | 2016.44 | 1669.35 | 2060.83 |
| ENSMUSG00000095550 | 22.43 | 17.67 | 15.68 | 28.94 | 24.84 | 30.27 | 18.88 | 19.74 | 18.65 | 19.91 | 16.15 | 13.19 |
| ENSMUSG00000104436 | 2.93 | 2.95 | 3.92 | 4.13 | 5.40 | 5.22 | 3.60 | 6.58 | 4.91 | 2.21 | 4.04 | 2.83 |
| ENSMUSG00000097787 | 10.73 | 12.76 | 15.68 | 17.57 | 22.68 | 18.79 | 8.99 | 15.63 | 10.80 | 11.06 | 9.08 | 15.07 |
| ENSMUSG00000062470 | 16.58 | 23.56 | 20.58 | 26.88 | 31.32 | 33.40 | 30.57 | 18.92 | 22.58 | 25.44 | 24.22 | 17.90 |
| ENSMUSG00000028687 | 23.40 | 33.38 | 28.42 | 37.21 | 47.52 | 43.84 | 25.18 | 39.49 | 22.58 | 42.03 | 32.30 | 39.56 |
| ENSMUSG00000109454 | 19.50 | 13.74 | 17.64 | 25.84 | 29.16 | 21.92 | 17.09 | 14.81 | 20.61 | 33.18 | 19.18 | 30.14 |
| ENSMUSG00000072672 | 19.50 | 19.63 | 13.72 | 25.84 | 28.08 | 26.10 | 25.18 | 27.97 | 23.56 | 25.44 | 21.19 | 22.61 |
| ENSMUSG00000102302 | 28.28 | 19.63 | 28.42 | 35.15 | 42.12 | 38.62 | 13.49 | 20.57 | 31.41 | 18.80 | 10.09 | 18.84 |

| | | | | | | | | | | | | |
|---|---|---|---|---|---|---|---|---|---|---|---|---|
| ENSMUSG00000095123 | 81.90 | 104.06 | 106.82 | 121.98 | 158.76 | 163.88 | 98.92 | 90.49 | 93.25 | 117.25 | 60.56 | 103.61 |
| ENSMUSG00000089726 | 15.60 | 22.58 | 20.58 | 27.91 | 33.48 | 28.18 | 14.39 | 26.33 | 17.67 | 23.23 | 24.22 | 17.90 |
| ENSMUSG00000060441 | 14.63 | 16.69 | 19.60 | 27.91 | 22.68 | 27.14 | 14.39 | 16.45 | 14.72 | 14.38 | 15.14 | 21.66 |
| ENSMUSG00000091985 | 15.60 | 21.60 | 18.62 | 27.91 | 28.08 | 29.23 | 18.88 | 26.33 | 24.54 | 21.02 | 23.21 | 24.49 |
| ENSMUSG00000029149 | 5.85 | 7.85 | 4.90 | 8.27 | 10.80 | 9.39 | 5.40 | 7.40 | 7.85 | 7.74 | 5.05 | 4.71 |
| ENSMUSG00000028248 | 1083.29 | 1547.23 | 1356.35 | 1860.66 | 2103.89 | 2138.83 | 1542.21 | 1425.70 | 1647.08 | 1697.88 | 1366.57 | 1461.80 |
| ENSMUSG00000004698 | 99.46 | 120.75 | 102.90 | 165.39 | 157.68 | 172.23 | 147.48 | 112.71 | 153.13 | 162.60 | 156.44 | 130.92 |
| ENSMUSG00000097586 | 12.68 | 15.71 | 18.62 | 23.78 | 27.00 | 21.92 | 19.78 | 11.52 | 15.71 | 28.76 | 18.17 | 19.78 |
| ENSMUSG00000072769 | 17.55 | 13.74 | 10.78 | 23.78 | 21.60 | 19.83 | 16.19 | 15.63 | 13.74 | 12.17 | 14.13 | 9.42 |
| ENSMUSG00000074415 | 803.45 | 1196.75 | 1016.28 | 1321.07 | 1720.48 | 1637.79 | 997.26 | 1049.74 | 1079.73 | 1243.27 | 852.84 | 1049.25 |
| ENSMUSG00000082511 | 21.45 | 17.67 | 22.54 | 35.15 | 34.56 | 26.10 | 29.68 | 27.97 | 44.17 | 28.76 | 31.29 | 36.73 |
| ENSMUSG00000105163 | 5.85 | 6.87 | 4.90 | 8.27 | 10.80 | 8.35 | 12.59 | 4.94 | 8.83 | 12.17 | 7.06 | 6.59 |
| ENSMUSG00000032226 | 4.88 | 5.89 | 3.92 | 8.27 | 7.56 | 7.31 | 8.09 | 4.94 | 2.94 | 3.32 | 6.06 | 3.77 |
| ENSMUSG00000080316 | 149.18 | 241.51 | 208.74 | 300.81 | 311.05 | 332.99 | 215.82 | 210.61 | 263.06 | 297.54 | 153.41 | 231.70 |
| ENSMUSG00000105540 | 5.85 | 6.87 | 7.84 | 9.30 | 11.88 | 11.48 | 17.09 | 8.23 | 7.85 | 7.74 | 10.09 | 5.65 |
| ENSMUSG00000097027 | 12.68 | 8.84 | 14.70 | 16.54 | 19.44 | 21.92 | 14.39 | 18.92 | 11.78 | 14.38 | 10.09 | 6.59 |
| ENSMUSG00000085173 | 17.55 | 17.67 | 14.70 | 23.78 | 28.08 | 28.18 | 17.98 | 17.28 | 26.50 | 26.55 | 15.14 | 15.07 |
| ENSMUSG00000044694 | 42.90 | 54.98 | 49.98 | 72.36 | 77.76 | 87.68 | 63.85 | 55.12 | 55.95 | 64.15 | 41.38 | 68.76 |
| ENSMUSG00000099618 | 7.80 | 5.89 | 5.88 | 10.34 | 10.80 | 10.44 | 7.19 | 5.76 | 6.87 | 3.32 | 4.04 | 10.36 |
| ENSMUSG00000083288 | 18.53 | 24.54 | 25.48 | 37.21 | 41.04 | 33.40 | 40.47 | 23.86 | 38.28 | 36.50 | 25.23 | 38.62 |
| ENSMUSG00000069892 | 197.94 | 275.87 | 203.84 | 300.81 | 383.41 | 419.62 | 298.55 | 291.23 | 373.00 | 289.80 | 264.43 | 308.94 |
| ENSMUSG00000070407 | 9.75 | 11.78 | 14.70 | 15.51 | 21.60 | 21.92 | 17.98 | 23.04 | 12.76 | 17.70 | 17.16 | 27.31 |
| ENSMUSG00000107215 | 20.48 | 30.43 | 32.34 | 43.42 | 50.76 | 41.75 | 32.37 | 13.99 | 27.48 | 27.65 | 14.13 | 36.73 |
| ENSMUSG00000034063 | 5.85 | 6.87 | 5.88 | 10.34 | 8.64 | 11.48 | 15.29 | 7.40 | 11.78 | 9.95 | 4.04 | 4.71 |
| ENSMUSG00000099241 | 7.80 | 14.73 | 12.74 | 16.54 | 19.44 | 21.92 | 11.69 | 18.92 | 17.67 | 9.95 | 15.14 | 23.55 |
| ENSMUSG00000083817 | 22.43 | 31.42 | 26.46 | 38.25 | 51.84 | 41.75 | 30.57 | 14.81 | 38.28 | 43.14 | 43.40 | 20.72 |
| ENSMUSG00000095474 | 3.90 | 5.89 | 6.86 | 8.27 | 9.72 | 9.39 | 3.60 | 3.29 | 7.85 | 3.32 | 8.07 | 4.71 |
| ENSMUSG00000098934 | 7.80 | 14.73 | 12.74 | 17.57 | 18.36 | 22.96 | 11.69 | 18.92 | 18.65 | 9.95 | 15.14 | 25.43 |
| ENSMUSG00000031849 | 2.93 | 2.95 | 2.94 | 4.13 | 5.40 | 5.22 | 5.40 | 4.11 | 3.93 | 5.53 | 5.05 | 6.59 |
| ENSMUSG00000080778 | 21.45 | 13.74 | 15.68 | 28.94 | 28.08 | 28.18 | 14.39 | 23.04 | 30.43 | 18.80 | 16.15 | 26.37 |
| ENSMUSG00000083303 | 12.68 | 6.87 | 11.76 | 16.54 | 19.44 | 16.70 | 23.38 | 15.63 | 26.50 | 14.38 | 17.16 | 16.95 |
| ENSMUSG00000093800 | 19.50 | 29.45 | 21.56 | 37.21 | 39.96 | 41.75 | 24.28 | 12.34 | 26.50 | 35.40 | 25.23 | 27.31 |
| ENSMUSG00000003934 | 8.78 | 6.87 | 7.84 | 14.47 | 11.88 | 13.57 | 7.19 | 6.58 | 13.74 | 5.53 | 6.06 | 8.48 |
| ENSMUSG00000049460 | 2.93 | 4.91 | 3.92 | 6.20 | 7.56 | 6.26 | 1.80 | 4.94 | 4.91 | 6.64 | 2.02 | 4.71 |

| | | | | | | | | | | | | |
|---|---|---|---|---|---|---|---|---|---|---|---|---|
| ENSMUSG00000083510 | 5.85 | 4.91 | 5.88 | 10.34 | 8.64 | 9.39 | 7.19 | 9.05 | 12.76 | 12.17 | 10.09 | 8.48 |
| ENSMUSG00000028417 | 9.75 | 11.78 | 6.86 | 14.47 | 18.36 | 15.66 | 11.69 | 11.52 | 14.72 | 11.06 | 9.08 | 11.30 |
| ENSMUSG00000078247 | 44.85 | 77.56 | 54.88 | 83.73 | 99.36 | 120.04 | 67.44 | 61.70 | 68.71 | 68.58 | 71.66 | 90.42 |
| ENSMUSG00000084378 | 8.78 | 13.74 | 13.72 | 19.64 | 17.28 | 25.05 | 20.68 | 18.10 | 17.67 | 21.02 | 21.19 | 27.31 |
| ENSMUSG00000090086 | 46.80 | 79.52 | 60.76 | 86.83 | 118.80 | 116.91 | 66.54 | 75.69 | 71.65 | 68.58 | 38.35 | 63.11 |
| ENSMUSG00000109489 | 37.05 | 42.22 | 41.16 | 66.16 | 76.68 | 65.76 | 46.76 | 55.12 | 65.77 | 53.09 | 48.45 | 54.63 |
| ENSMUSG00000105112 | 12.68 | 14.73 | 13.72 | 27.91 | 19.44 | 24.01 | 6.29 | 12.34 | 25.52 | 8.85 | 20.19 | 16.01 |
| ENSMUSG00000081427 | 6.83 | 12.76 | 8.82 | 18.61 | 16.20 | 14.61 | 8.09 | 8.23 | 11.78 | 19.91 | 10.09 | 17.90 |
| ENSMUSG00000021209 | 8.78 | 15.71 | 11.76 | 20.67 | 21.60 | 20.88 | 14.39 | 11.52 | 16.69 | 15.49 | 13.12 | 11.30 |
| ENSMUSG00000105084 | 11.70 | 15.71 | 11.76 | 25.84 | 21.60 | 20.88 | 12.59 | 19.74 | 21.59 | 23.23 | 21.19 | 18.84 |
| ENSMUSG00000033713 | 156.98 | 232.67 | 202.86 | 284.27 | 352.09 | 399.79 | 223.01 | 190.04 | 254.23 | 266.57 | 309.85 | 282.56 |
| ENSMUSG00000105382 | 2.93 | 3.93 | 3.92 | 7.24 | 5.40 | 6.26 | 4.50 | 4.94 | 2.94 | 3.32 | 3.03 | 2.83 |
| ENSMUSG00000054728 | 5.85 | 5.89 | 6.86 | 9.30 | 12.96 | 10.44 | 3.60 | 3.29 | 5.89 | 7.74 | 10.09 | 3.77 |
| ENSMUSG00000097892 | 10.73 | 10.80 | 10.78 | 18.61 | 20.52 | 17.75 | 14.39 | 9.05 | 9.82 | 13.27 | 7.06 | 5.65 |
| ENSMUSG00000081633 | 17.55 | 15.71 | 10.78 | 31.01 | 24.84 | 21.92 | 26.98 | 19.74 | 11.78 | 16.59 | 25.23 | 23.55 |
| ENSMUSG00000103928 | 21.45 | 21.60 | 35.28 | 44.45 | 49.68 | 44.89 | 30.57 | 39.49 | 45.15 | 45.35 | 26.24 | 28.26 |
| ENSMUSG00000085028 | 12.68 | 20.62 | 26.46 | 41.35 | 32.40 | 32.36 | 15.29 | 24.68 | 21.59 | 24.33 | 16.15 | 24.49 |
| ENSMUSG00000107205 | 8.78 | 4.91 | 8.82 | 11.37 | 16.20 | 12.53 | 3.60 | 8.23 | 9.82 | 6.64 | 11.10 | 3.77 |
| ENSMUSG00000097418 | 12.68 | 16.69 | 19.60 | 26.88 | 30.24 | 30.27 | 10.79 | 23.86 | 17.67 | 23.23 | 8.07 | 15.07 |
| ENSMUSG00000073164 | 4.88 | 5.89 | 6.86 | 11.37 | 11.88 | 8.35 | 7.19 | 9.05 | 7.85 | 6.64 | 10.09 | 6.59 |
| ENSMUSG00000081308 | 31.20 | 59.89 | 37.24 | 80.63 | 83.16 | 66.81 | 89.03 | 55.12 | 78.53 | 77.43 | 67.62 | 95.13 |
| ENSMUSG00000052631 | 4.88 | 2.95 | 3.92 | 6.20 | 8.64 | 6.26 | 1.80 | 2.47 | 5.89 | 4.42 | 2.02 | 3.77 |
| ENSMUSG00000101599 | 75.08 | 89.34 | 99.96 | 123.01 | 185.76 | 167.01 | 77.34 | 106.95 | 95.21 | 116.14 | 83.77 | 92.30 |
| ENSMUSG00000097760 | 2.93 | 3.93 | 5.88 | 9.30 | 6.48 | 7.31 | 3.60 | 4.11 | 3.93 | 6.64 | 4.04 | 3.77 |
| ENSMUSG00000087624 | 3.90 | 2.95 | 2.94 | 6.20 | 5.40 | 6.26 | 2.70 | 4.94 | 5.89 | 1.11 | 3.03 | 0.94 |
| ENSMUSG00000074622 | 4.88 | 6.87 | 4.90 | 11.37 | 9.72 | 9.39 | 12.59 | 5.76 | 5.89 | 7.74 | 8.07 | 10.36 |
| ENSMUSG00000065147 | 4.88 | 7.85 | 3.92 | 10.34 | 11.88 | 8.35 | 4.50 | 6.58 | 3.93 | 2.21 | 6.06 | 4.71 |
| ENSMUSG00000094374 | 8.78 | 6.87 | 7.84 | 11.37 | 15.12 | 16.70 | 11.69 | 11.52 | 17.67 | 16.59 | 14.13 | 13.19 |
| ENSMUSG00000065480 | 10.73 | 16.69 | 13.72 | 23.78 | 24.84 | 27.14 | 14.39 | 9.05 | 18.65 | 12.17 | 17.16 | 16.95 |
| ENSMUSG00000084146 | 7.80 | 5.89 | 3.92 | 10.34 | 10.80 | 11.48 | 12.59 | 8.23 | 7.85 | 14.38 | 11.10 | 11.30 |
| ENSMUSG00000097944 | 9.75 | 16.69 | 20.58 | 29.98 | 28.08 | 29.23 | 13.49 | 19.74 | 22.58 | 16.59 | 14.13 | 24.49 |
| ENSMUSG00000043243 | 20.48 | 21.60 | 19.60 | 43.42 | 33.48 | 38.62 | 30.57 | 18.10 | 19.63 | 35.40 | 16.15 | 16.01 |
| ENSMUSG00000103046 | 13.65 | 16.69 | 20.58 | 24.81 | 33.48 | 37.58 | 13.49 | 31.26 | 31.41 | 33.18 | 25.23 | 22.61 |
| ENSMUSG00000103477 | 6.83 | 7.85 | 4.90 | 10.34 | 14.04 | 12.53 | 6.29 | 6.58 | 10.80 | 6.64 | 10.09 | 8.48 |

| | | | | | | | | | | | | |
|---|---|---|---|---|---|---|---|---|---|---|---|---|
| ENSMUSG00000093629 | 12.68 | 15.71 | 8.82 | 22.74 | 22.68 | 25.05 | 17.09 | 11.52 | 7.85 | 13.27 | 13.12 | 15.07 |
| ENSMUSG00000081101 | 6.83 | 8.84 | 5.88 | 11.37 | 17.28 | 12.53 | 9.89 | 7.40 | 12.76 | 19.91 | 8.07 | 14.13 |
| ENSMUSG00000091442 | 4.88 | 4.91 | 4.90 | 9.30 | 10.80 | 8.35 | 6.29 | 4.11 | 9.82 | 5.53 | 10.09 | 5.65 |
| ENSMUSG00000104399 | 7.80 | 18.65 | 11.76 | 25.84 | 27.00 | 21.92 | 17.09 | 21.39 | 12.76 | 27.65 | 15.14 | 18.84 |
| ENSMUSG00000082793 | 6.83 | 6.87 | 2.94 | 13.44 | 9.72 | 9.39 | 12.59 | 6.58 | 9.82 | 13.27 | 11.10 | 15.07 |
| ENSMUSG00000100504 | 18.53 | 19.63 | 20.58 | 43.42 | 41.04 | 31.32 | 17.09 | 23.86 | 48.10 | 35.40 | 27.25 | 35.79 |
| ENSMUSG00000056155 | 3.90 | 4.91 | 2.94 | 8.27 | 8.64 | 6.26 | 2.70 | 4.11 | 3.93 | 1.11 | 5.05 | 1.88 |
| ENSMUSG00000104184 | 4.88 | 5.89 | 6.86 | 9.30 | 14.04 | 11.48 | 9.89 | 8.23 | 8.83 | 9.95 | 5.05 | 5.65 |
| ENSMUSG00000081050 | 2.93 | 3.93 | 5.88 | 8.27 | 9.72 | 7.31 | 7.19 | 8.23 | 10.80 | 6.64 | 2.02 | 7.54 |
| ENSMUSG00000087445 | 6.83 | 5.89 | 6.86 | 10.34 | 12.96 | 15.66 | 8.99 | 5.76 | 5.89 | 13.27 | 7.06 | 9.42 |
| ENSMUSG00000104501 | 1.95 | 2.95 | 1.96 | 4.13 | 4.32 | 5.22 | 0.90 | 4.94 | 3.93 | 1.11 | 2.02 | 3.77 |
| ENSMUSG00000100037 | 2.93 | 4.91 | 6.86 | 10.34 | 7.56 | 11.48 | 6.29 | 11.52 | 15.71 | 11.06 | 10.09 | 8.48 |
| ENSMUSG00000082534 | 5.85 | 4.91 | 3.92 | 9.30 | 8.64 | 11.48 | 1.80 | 2.47 | 11.78 | 1.11 | 16.15 | 10.36 |
| ENSMUSG00000105655 | 10.73 | 8.84 | 8.82 | 15.51 | 20.52 | 20.88 | 8.09 | 10.69 | 10.80 | 13.27 | 9.08 | 9.42 |
| ENSMUSG00000025375 | 4.88 | 3.93 | 5.88 | 8.27 | 10.80 | 10.44 | 7.19 | 10.69 | 12.76 | 7.74 | 10.09 | 4.71 |
| ENSMUSG00000102649 | 5.85 | 7.85 | 10.78 | 14.47 | 14.04 | 20.88 | 8.09 | 9.05 | 5.89 | 5.53 | 9.08 | 6.59 |
| ENSMUSG00000108465 | 16.58 | 20.62 | 26.46 | 34.11 | 43.20 | 51.15 | 17.98 | 19.74 | 36.32 | 37.61 | 23.21 | 25.43 |
| ENSMUSG00000102555 | 5.85 | 13.74 | 13.72 | 19.64 | 21.60 | 26.10 | 12.59 | 9.05 | 12.76 | 12.17 | 16.15 | 18.84 |
| ENSMUSG00000083700 | 14.63 | 18.65 | 18.62 | 38.25 | 38.88 | 28.18 | 28.78 | 29.62 | 26.50 | 32.08 | 30.28 | 31.08 |
| ENSMUSG00000099465 | 11.70 | 13.74 | 4.90 | 17.57 | 21.60 | 22.96 | 12.59 | 18.10 | 21.59 | 18.80 | 7.06 | 16.95 |
| ENSMUSG00000109390 | 9.75 | 3.93 | 6.86 | 12.40 | 17.28 | 12.53 | 8.09 | 8.23 | 6.87 | 3.32 | 7.06 | 9.42 |
| ENSMUSG00000023484 | 7.80 | 5.89 | 9.80 | 15.51 | 12.96 | 19.83 | 9.89 | 13.99 | 10.80 | 6.64 | 10.09 | 14.13 |
| ENSMUSG00000107465 | 3.90 | 2.95 | 3.92 | 6.20 | 8.64 | 7.31 | 2.70 | 2.47 | 3.93 | 4.42 | 5.05 | 7.54 |
| ENSMUSG00000036045 | 4.88 | 10.80 | 8.82 | 17.57 | 16.20 | 16.70 | 15.29 | 11.52 | 9.82 | 12.17 | 17.16 | 11.30 |
| ENSMUSG00000084377 | 6.83 | 4.91 | 5.88 | 14.47 | 12.96 | 9.39 | 5.40 | 9.87 | 8.83 | 16.59 | 6.06 | 7.54 |
| ENSMUSG00000033847 | 10.73 | 17.67 | 5.88 | 24.81 | 28.08 | 18.79 | 14.39 | 21.39 | 13.74 | 17.70 | 21.19 | 9.42 |
| ENSMUSG00000097445 | 0.98 | 2.95 | 2.94 | 5.17 | 4.32 | 5.22 | 4.50 | 6.58 | 3.93 | 3.32 | 7.06 | 3.77 |
| ENSMUSG00000063142 | 2.93 | 1.96 | 0.98 | 5.17 | 3.24 | 4.18 | 4.50 | 1.65 | 2.94 | 6.64 | 4.04 | 2.83 |
| ENSMUSG00000102425 | 1.95 | 4.91 | 5.88 | 8.27 | 8.64 | 10.44 | 0.90 | 6.58 | 4.91 | 9.95 | 10.09 | 4.71 |
| ENSMUSG00000086203 | 1.95 | 1.96 | 2.94 | 4.13 | 4.32 | 6.26 | 5.40 | 2.47 | 2.94 | 2.21 | 2.02 | 4.71 |
| ENSMUSG00000108764 | 6.83 | 8.84 | 2.94 | 14.47 | 15.12 | 10.44 | 12.59 | 5.76 | 11.78 | 8.85 | 17.16 | 8.48 |
| ENSMUSG00000037759 | 1.95 | 4.91 | 3.92 | 7.24 | 9.72 | 6.26 | 6.29 | 9.05 | 4.91 | 3.32 | 3.03 | 6.59 |
| ENSMUSG00000045008 | 2.93 | 6.87 | 3.92 | 8.27 | 11.88 | 10.44 | 2.70 | 5.76 | 4.91 | 4.42 | 7.06 | 6.59 |
| ENSMUSG00000031842 | 2.93 | 3.93 | 3.92 | 7.24 | 8.64 | 8.35 | 11.69 | 8.23 | 4.91 | 3.32 | 10.09 | 10.36 |

| | | | | | | | | | | | | |
|---|---|---|---|---|---|---|---|---|---|---|---|---|
| ENSMUSG00000029092 | 4.88 | 3.93 | 1.96 | 8.27 | 8.64 | 7.31 | 5.40 | 7.40 | 1.96 | 9.95 | 1.01 | 4.71 |
| ENSMUSG00000021604 | 2.93 | 2.95 | 0.98 | 5.17 | 4.32 | 6.26 | 1.80 | 2.47 | 5.89 | 4.42 | 4.04 | 1.88 |
| ENSMUSG00000082315 | 2.93 | 1.96 | 1.96 | 5.17 | 5.40 | 5.22 | 4.50 | 5.76 | 1.96 | 2.21 | 3.03 | 2.83 |
| ENSMUSG00000090942 | 2.93 | 9.82 | 4.90 | 15.51 | 10.80 | 14.61 | 4.50 | 2.47 | 6.87 | 9.95 | 8.07 | 7.54 |
| ENSMUSG00000047343 | 0.98 | 2.95 | 1.96 | 4.13 | 4.32 | 5.22 | 4.50 | 6.58 | 1.96 | 4.42 | 3.03 | 1.88 |
| ENSMUSG00000054910 | 0.98 | 1.96 | 1.96 | 4.13 | 3.24 | 4.18 | 3.60 | 1.65 | 1.96 | 3.32 | 3.03 | 3.77 |
| ENSMUSG00000034616 | 9.75 | 1.96 | 8.82 | 16.54 | 18.36 | 13.57 | 11.69 | 13.16 | 17.67 | 9.95 | 14.13 | 15.07 |
| ENSMUSG00000074817 | 5.85 | 5.89 | 7.84 | 14.47 | 19.44 | 12.53 | 7.19 | 7.40 | 8.83 | 7.74 | 7.06 | 12.24 |
| ENSMUSG00000039224 | 1.95 | 5.89 | 3.92 | 11.37 | 8.64 | 8.35 | 6.29 | 2.47 | 8.83 | 5.53 | 6.06 | 7.54 |
| ENSMUSG00000102691 | 3.90 | 4.91 | 6.86 | 11.37 | 16.20 | 10.44 | 7.19 | 9.05 | 6.87 | 11.06 | 10.09 | 7.54 |
| ENSMUSG00000040867 | 3.90 | 1.96 | 1.96 | 6.20 | 7.56 | 5.22 | 4.50 | 1.65 | 0.98 | 4.42 | 4.04 | 2.83 |
| ENSMUSG00000103082 | 4.88 | 3.93 | 1.96 | 9.30 | 8.64 | 8.35 | 3.60 | 5.76 | 2.94 | 5.53 | 2.02 | 1.88 |
| ENSMUSG00000042757 | 0.98 | 0.98 | 0.98 | 3.10 | 2.16 | 2.09 | 2.70 | 2.47 | 1.96 | 2.21 | 2.02 | 0.94 |
| ENSMUSG00000082531 | 0.98 | 0.98 | 0.98 | 3.10 | 2.16 | 2.09 | 1.80 | 0.82 | 1.96 | 1.11 | 2.02 | 1.88 |
| ENSMUSG00000107894 | 2.93 | 3.93 | 1.96 | 8.27 | 8.64 | 5.22 | 3.60 | 2.47 | 1.96 | 4.42 | 1.01 | 0.94 |
| ENSMUSG00000086916 | 6.83 | 2.95 | 3.92 | 8.27 | 14.04 | 12.53 | 3.60 | 7.40 | 8.83 | 5.53 | 7.06 | 12.24 |
| ENSMUSG00000083149 | 4.88 | 4.91 | 2.94 | 12.40 | 11.88 | 8.35 | 7.19 | 9.05 | 5.89 | 11.06 | 7.06 | 14.13 |
| ENSMUSG00000104503 | 3.90 | 1.96 | 3.92 | 8.27 | 7.56 | 9.39 | 2.70 | 1.65 | 2.94 | 1.11 | 7.06 | 1.88 |
| ENSMUSG00000029784 | 0.98 | 0.98 | 2.94 | 3.10 | 4.32 | 5.22 | 3.60 | 4.94 | 1.96 | 1.11 | 1.01 | 0.94 |
| ENSMUSG00000106354 | 4.88 | 2.95 | 1.96 | 7.24 | 8.64 | 9.39 | 4.50 | 0.82 | 5.89 | 1.11 | 2.02 | 0.94 |
| ENSMUSG00000021268 | 0.98 | 2.95 | 0.98 | 4.13 | 5.40 | 3.13 | 1.80 | 0.82 | 0.98 | 1.11 | 2.02 | 0.94 |
| ENSMUSG00000100005 | 9.75 | 6.87 | 8.82 | 20.67 | 23.76 | 21.92 | 9.89 | 9.87 | 15.71 | 12.17 | 11.10 | 11.30 |
| ENSMUSG00000109379 | 3.90 | 11.78 | 6.86 | 16.54 | 21.60 | 20.88 | 6.29 | 7.40 | 5.89 | 8.85 | 6.06 | 5.65 |
| ENSMUSG00000105556 | 1.95 | 6.87 | 6.86 | 10.34 | 18.36 | 12.53 | 1.80 | 6.58 | 7.85 | 4.42 | 8.07 | 4.71 |
| ENSMUSG00000045991 | 6.83 | 6.87 | 9.80 | 17.57 | 24.84 | 19.83 | 17.09 | 16.45 | 12.76 | 14.38 | 12.11 | 10.36 |
| ENSMUSG00000057182 | 3.90 | 3.93 | 3.92 | 10.34 | 9.72 | 11.48 | 11.69 | 7.40 | 10.80 | 6.64 | 7.06 | 11.30 |
| ENSMUSG00000085517 | 3.90 | 0.98 | 2.94 | 8.27 | 6.48 | 6.26 | 5.40 | 3.29 | 4.91 | 9.95 | 1.01 | 4.71 |
| ENSMUSG00000104626 | 1.95 | 5.89 | 12.74 | 13.44 | 21.60 | 20.88 | 3.60 | 2.47 | 6.87 | 5.53 | 11.10 | 3.77 |
| ENSMUSG00000085513 | 5.85 | 0.98 | 2.94 | 7.24 | 9.72 | 10.44 | 8.09 | 1.65 | 9.82 | 5.53 | 15.14 | 0.94 |
| ENSMUSG00000109198 | 2.93 | 1.96 | 2.94 | 7.24 | 8.64 | 6.26 | 8.99 | 4.94 | 6.87 | 5.53 | 6.06 | 1.88 |
| ENSMUSG00000104716 | 3.90 | 2.95 | 0.98 | 7.24 | 9.72 | 5.22 | 4.50 | 7.40 | 3.93 | 5.53 | 5.05 | 5.65 |
| ENSMUSG00000108221 | 2.93 | 0.98 | 1.96 | 6.20 | 6.48 | 4.18 | 6.29 | 2.47 | 5.89 | 1.11 | 6.06 | 4.71 |
| ENSMUSG00000102414 | 0.98 | 0.98 | 0.98 | 2.07 | 3.24 | 3.13 | 2.70 | 2.47 | 0.98 | 2.21 | 1.01 | 1.88 |
| ENSMUSG00000103976 | 5.85 | 4.91 | 4.90 | 16.54 | 16.20 | 12.53 | 8.09 | 6.58 | 9.82 | 5.53 | 4.04 | 9.42 |

| | | | | | | | | | | | | |
|---|---|---|---|---|---|---|---|---|---|---|---|---|
| ENSMUSG00000108394 | 2.93 | 0.98 | 5.88 | 7.24 | 11.88 | 9.39 | 0.90 | 1.65 | 2.94 | 1.11 | 5.05 | 2.83 |
| ENSMUSG00000040919 | 4.88 | 2.95 | 2.94 | 8.27 | 12.96 | 10.44 | 6.29 | 5.76 | 5.89 | 2.21 | 6.06 | 7.54 |
| ENSMUSG00000080892 | 3.90 | 2.95 | 0.98 | 8.27 | 5.40 | 9.39 | 3.60 | 7.40 | 8.83 | 5.53 | 3.03 | 6.59 |
| ENSMUSG00000032013 | 0.98 | 1.96 | 0.98 | 4.13 | 4.32 | 3.13 | 1.80 | 0.82 | 0.98 | 2.21 | 3.03 | 6.59 |
| ENSMUSG00000107227 | 2.93 | 4.91 | 8.82 | 14.47 | 17.28 | 17.75 | 6.29 | 4.11 | 6.87 | 3.32 | 4.04 | 6.59 |
| ENSMUSG00000106099 | 4.88 | 6.87 | 12.74 | 18.61 | 28.08 | 26.10 | 8.09 | 10.69 | 10.80 | 12.17 | 9.08 | 16.01 |
| ENSMUSG00000047894 | 0.98 | 1.96 | 1.96 | 6.20 | 4.32 | 4.18 | 3.60 | 4.94 | 0.98 | 4.42 | 3.03 | 9.42 |
| ENSMUSG00000017453 | 0.98 | 2.95 | 0.98 | 5.17 | 4.32 | 5.22 | 0.90 | 2.47 | 0.98 | 2.21 | 1.01 | 2.83 |
| ENSMUSG00000099018 | 2.93 | 0.98 | 0.98 | 7.24 | 4.32 | 4.18 | 2.70 | 3.29 | 3.93 | 1.11 | 6.06 | 5.65 |
| ENSMUSG00000068226 | 2.93 | 1.96 | 0.98 | 7.24 | 7.56 | 4.18 | 5.40 | 7.40 | 5.89 | 4.42 | 8.07 | 4.71 |
| ENSMUSG00000082794 | 1.95 | 7.85 | 6.86 | 13.44 | 21.60 | 22.96 | 7.19 | 2.47 | 0.98 | 3.32 | 4.04 | 4.71 |
| ENSMUSG00000021356 | 0.98 | 0.98 | 0.98 | 3.10 | 3.24 | 4.18 | 5.40 | 2.47 | 1.96 | 6.64 | 2.02 | 2.83 |
| ENSMUSG00000103065 | 1.95 | 0.98 | 1.96 | 7.24 | 4.32 | 6.26 | 0.90 | 5.76 | 0.98 | 1.11 | 1.01 | 1.88 |
| ENSMUSG00000103720 | 1.95 | 0.98 | 1.96 | 7.24 | 4.32 | 6.26 | 0.90 | 5.76 | 0.98 | 1.11 | 1.01 | 1.88 |
| ENSMUSG00000097054 | 0.98 | 2.95 | 0.98 | 5.17 | 7.56 | 5.22 | 2.70 | 2.47 | 7.85 | 4.42 | 8.07 | 1.88 |
| ENSMUSG00000097190 | 0.98 | 2.95 | 0.98 | 5.17 | 7.56 | 5.22 | 2.70 | 2.47 | 7.85 | 4.42 | 8.07 | 1.88 |
| ENSMUSG00000108954 | 1.95 | 1.96 | 1.96 | 6.20 | 8.64 | 7.31 | 4.50 | 3.29 | 1.96 | 4.42 | 4.04 | 7.54 |
| ENSMUSG00000107320 | 1.95 | 2.95 | 2.94 | 7.24 | 10.80 | 12.53 | 6.29 | 9.05 | 6.87 | 6.64 | 6.06 | 4.71 |
| ENSMUSG00000046958 | 2.93 | 0.98 | 0.98 | 6.20 | 5.40 | 8.35 | 2.70 | 2.47 | 3.93 | 1.11 | 3.03 | 4.71 |
| ENSMUSG00000109135 | 6.83 | 11.78 | 7.84 | 5.17 | 1.08 | 1.04 | 7.19 | 9.05 | 4.91 | 8.85 | 4.04 | 4.71 |
| ENSMUSG00000060560 | 3.90 | 4.91 | 5.88 | 2.07 | 1.08 | 2.09 | 0.90 | 4.11 | 1.96 | 1.11 | 5.05 | 3.77 |
| ENSMUSG00000027716 | 5.85 | 5.89 | 4.90 | 1.03 | 2.16 | 3.13 | 6.29 | 5.76 | 2.94 | 3.32 | 5.05 | 3.77 |
| ENSMUSG00000045761 | 3.90 | 3.93 | 5.88 | 1.03 | 3.24 | 1.04 | 5.40 | 2.47 | 5.89 | 7.74 | 5.05 | 6.59 |
| ENSMUSG00000104694 | 10.73 | 11.78 | 8.82 | 5.17 | 4.32 | 3.13 | 6.29 | 8.23 | 9.82 | 7.74 | 6.06 | 1.88 |
| ENSMUSG00000015533 | 2.93 | 1.96 | 2.94 | 1.03 | 1.08 | 1.04 | 0.90 | 1.65 | 2.94 | 1.11 | 2.02 | 2.83 |
| ENSMUSG00000029641 | 16.58 | 12.76 | 9.80 | 4.13 | 7.56 | 4.18 | 7.19 | 11.52 | 8.83 | 13.27 | 16.15 | 12.24 |
| ENSMUSG00000097839 | 3.90 | 4.91 | 3.92 | 2.07 | 1.08 | 2.09 | 3.60 | 2.47 | 0.98 | 2.21 | 3.03 | 1.88 |
| ENSMUSG00000098085 | 8.78 | 10.80 | 7.84 | 4.13 | 4.32 | 3.13 | 13.49 | 13.99 | 10.80 | 4.42 | 9.08 | 14.13 |
| ENSMUSG00000029410 | 6.83 | 4.91 | 7.84 | 2.07 | 3.24 | 3.13 | 1.80 | 4.11 | 5.89 | 4.42 | 4.04 | 2.83 |
| ENSMUSG00000040935 | 2.93 | 2.95 | 3.92 | 1.03 | 2.16 | 1.04 | 1.80 | 0.82 | 1.96 | 2.21 | 4.04 | 3.77 |
| ENSMUSG00000100632 | 9.75 | 8.84 | 7.84 | 5.17 | 2.16 | 4.18 | 2.70 | 8.23 | 7.85 | 2.21 | 9.08 | 4.71 |
| ENSMUSG00000091191 | 6.83 | 8.84 | 7.84 | 4.13 | 3.24 | 3.13 | 6.29 | 5.76 | 3.93 | 5.53 | 4.04 | 6.59 |
| ENSMUSG00000033316 | 6.83 | 6.87 | 4.90 | 4.13 | 2.16 | 2.09 | 5.40 | 6.58 | 5.89 | 5.53 | 5.05 | 6.59 |
| ENSMUSG00000033634 | 3.90 | 4.91 | 4.90 | 1.03 | 2.16 | 3.13 | 4.50 | 3.29 | 3.93 | 3.32 | 2.02 | 3.77 |

| | | | | | | | | | | | | |
|---|---|---|---|---|---|---|---|---|---|---|---|---|
| ENSMUSG00000045968 | 9.75 | 10.80 | 10.78 | 6.20 | 5.40 | 3.13 | 8.99 | 2.47 | 9.82 | 13.27 | 12.11 | 6.59 |
| ENSMUSG00000040891 | 2.93 | 2.95 | 2.94 | 2.07 | 1.08 | 1.04 | 4.50 | 0.82 | 4.91 | 1.11 | 3.03 | 0.94 |
| ENSMUSG00000083630 | 3.90 | 3.93 | 2.94 | 2.07 | 1.08 | 2.09 | 3.60 | 3.29 | 2.94 | 2.21 | 4.04 | 6.59 |
| ENSMUSG00000005493 | 3.90 | 3.93 | 2.94 | 2.07 | 2.16 | 1.04 | 0.90 | 1.65 | 0.98 | 1.11 | 3.03 | 1.88 |
| ENSMUSG00000098221 | 3.90 | 3.93 | 2.94 | 2.07 | 2.16 | 1.04 | 2.70 | 1.65 | 0.98 | 2.21 | 2.02 | 1.88 |
| ENSMUSG00000003526 | 20.48 | 18.65 | 18.62 | 13.44 | 8.64 | 6.26 | 9.89 | 12.34 | 13.74 | 7.74 | 11.10 | 16.01 |
| ENSMUSG00000107306 | 4.88 | 3.93 | 3.92 | 3.10 | 1.08 | 2.09 | 4.50 | 4.94 | 3.93 | 2.21 | 5.05 | 1.88 |
| ENSMUSG00000045231 | 3.90 | 3.93 | 4.90 | 2.07 | 1.08 | 3.13 | 2.70 | 0.82 | 8.83 | 4.42 | 4.04 | 3.77 |
| ENSMUSG00000106076 | 28.28 | 25.53 | 22.54 | 13.44 | 9.72 | 14.61 | 23.38 | 29.62 | 34.36 | 26.55 | 28.26 | 24.49 |
| ENSMUSG00000082621 | 4.88 | 4.91 | 4.90 | 2.07 | 2.16 | 3.13 | 5.40 | 3.29 | 2.94 | 2.21 | 5.05 | 7.54 |
| ENSMUSG00000048148 | 16.58 | 14.73 | 14.70 | 9.30 | 9.72 | 4.18 | 13.49 | 8.23 | 4.91 | 18.80 | 7.06 | 19.78 |
| ENSMUSG00000102890 | 7.80 | 4.91 | 5.88 | 3.10 | 2.16 | 4.18 | 8.09 | 4.94 | 4.91 | 15.49 | 6.06 | 10.36 |
| ENSMUSG00000071073 | 47.78 | 31.42 | 40.18 | 24.81 | 23.76 | 13.57 | 30.57 | 49.36 | 22.58 | 25.44 | 42.39 | 27.31 |
| ENSMUSG00000105361 | 113.11 | 74.61 | 95.06 | 60.99 | 51.84 | 34.45 | 45.86 | 45.25 | 51.04 | 32.08 | 81.75 | 41.44 |
| ENSMUSG00000106106 | 160.88 | 109.96 | 133.28 | 90.97 | 73.44 | 46.97 | 62.95 | 78.98 | 68.71 | 47.56 | 122.12 | 64.99 |
| ENSMUSG00000092202 | 31.20 | 26.51 | 26.46 | 14.47 | 14.04 | 15.66 | 19.78 | 16.45 | 16.69 | 17.70 | 23.21 | 24.49 |
| ENSMUSG00000048967 | 34.13 | 29.45 | 30.38 | 14.47 | 20.52 | 14.61 | 17.98 | 35.38 | 20.61 | 16.59 | 42.39 | 26.37 |
| ENSMUSG00000029452 | 25.35 | 31.42 | 23.52 | 17.57 | 12.96 | 12.53 | 29.68 | 26.33 | 22.58 | 25.44 | 25.23 | 19.78 |
| ENSMUSG00000021922 | 4.88 | 3.93 | 4.90 | 2.07 | 2.16 | 3.13 | 8.09 | 5.76 | 6.87 | 1.11 | 4.04 | 5.65 |
| ENSMUSG00000067642 | 22.43 | 17.67 | 16.66 | 11.37 | 9.72 | 9.39 | 13.49 | 16.45 | 17.67 | 15.49 | 16.15 | 21.66 |
| ENSMUSG00000095730 | 5.85 | 6.87 | 4.90 | 2.07 | 3.24 | 4.18 | 1.80 | 2.47 | 6.87 | 3.32 | 4.04 | 0.94 |
| ENSMUSG00000045555 | 28.28 | 21.60 | 22.54 | 12.40 | 14.04 | 12.53 | 20.68 | 23.86 | 28.47 | 27.65 | 27.25 | 25.43 |
| ENSMUSG00000020599 | 6.83 | 4.91 | 5.88 | 2.07 | 3.24 | 4.18 | 3.60 | 1.65 | 6.87 | 1.11 | 5.05 | 3.77 |
| ENSMUSG00000015083 | 11.70 | 9.82 | 12.74 | 8.27 | 4.32 | 6.26 | 1.80 | 5.76 | 7.85 | 9.95 | 5.05 | 3.77 |
| ENSMUSG00000056598 | 11.70 | 8.84 | 7.84 | 4.13 | 6.48 | 5.22 | 8.09 | 3.29 | 2.94 | 8.85 | 10.09 | 5.65 |
| ENSMUSG00000079963 | 5.85 | 7.85 | 8.82 | 4.13 | 3.24 | 5.22 | 8.09 | 2.47 | 9.82 | 9.95 | 9.08 | 8.48 |
| ENSMUSG00000021379 | 10.73 | 11.78 | 12.74 | 7.24 | 5.40 | 7.31 | 7.19 | 4.11 | 3.93 | 5.53 | 8.07 | 14.13 |
| ENSMUSG00000055413 | 35.10 | 31.42 | 26.46 | 21.71 | 12.96 | 18.79 | 25.18 | 25.50 | 36.32 | 25.44 | 33.31 | 40.50 |
| ENSMUSG00000003051 | 4.88 | 3.93 | 3.92 | 3.10 | 2.16 | 2.09 | 2.70 | 13.16 | 3.93 | 4.42 | 3.03 | 6.59 |
| ENSMUSG00000074973 | 3.90 | 4.91 | 3.92 | 2.07 | 2.16 | 3.13 | 3.60 | 3.29 | 1.96 | 1.11 | 3.03 | 4.71 |
| ENSMUSG00000026638 | 28.28 | 31.42 | 22.54 | 14.47 | 21.60 | 11.48 | 22.48 | 32.91 | 19.63 | 18.80 | 17.16 | 21.66 |
| ENSMUSG00000074577 | 35.10 | 26.51 | 30.38 | 21.71 | 11.88 | 19.83 | 27.88 | 22.21 | 23.56 | 15.49 | 27.25 | 24.49 |
| ENSMUSG00000084835 | 25.35 | 37.31 | 30.38 | 16.54 | 16.20 | 22.96 | 27.88 | 28.79 | 29.45 | 29.86 | 37.34 | 31.08 |
| ENSMUSG00000022219 | 31.20 | 25.53 | 27.44 | 16.54 | 20.52 | 13.57 | 20.68 | 15.63 | 17.67 | 17.70 | 32.30 | 32.02 |

| | | | | | | | | | | | | |
|---|---|---|---|---|---|---|---|---|---|---|---|---|
| ENSMUSG00000042357 | 42.90 | 43.20 | 42.14 | 32.04 | 25.92 | 20.88 | 30.57 | 34.55 | 18.65 | 23.23 | 33.31 | 35.79 |
| ENSMUSG00000097367 | 6.83 | 7.85 | 6.86 | 5.17 | 4.32 | 4.18 | 8.09 | 4.11 | 4.91 | 2.21 | 4.04 | 3.77 |
| ENSMUSG00000030518 | 23.40 | 18.65 | 21.56 | 16.54 | 10.80 | 13.57 | 18.88 | 15.63 | 17.67 | 11.06 | 14.13 | 19.78 |
| ENSMUSG00000042750 | 97.51 | 126.65 | 132.30 | 86.83 | 60.48 | 82.46 | 96.22 | 115.18 | 60.86 | 79.64 | 83.77 | 61.22 |
| ENSMUSG00000060550 | 234.99 | 211.08 | 196.98 | 176.76 | 110.16 | 128.39 | 190.64 | 201.56 | 185.52 | 184.72 | 207.91 | 167.65 |
| ENSMUSG00000004371 | 15.60 | 17.67 | 18.62 | 13.44 | 9.72 | 10.44 | 29.68 | 27.15 | 16.69 | 11.06 | 22.20 | 19.78 |
| ENSMUSG00000022225 | 15.60 | 13.74 | 15.68 | 10.34 | 7.56 | 11.48 | 20.68 | 15.63 | 11.78 | 14.38 | 11.10 | 13.19 |
| ENSMUSG00000059013 | 17.55 | 19.63 | 15.68 | 10.34 | 9.72 | 14.61 | 9.89 | 13.16 | 22.58 | 22.12 | 12.11 | 15.07 |
| ENSMUSG00000079355 | 37.05 | 38.29 | 43.12 | 25.84 | 28.08 | 25.05 | 29.68 | 28.79 | 31.41 | 26.55 | 29.27 | 29.20 |
| ENSMUSG00000106820 | 23.40 | 17.67 | 20.58 | 10.34 | 16.20 | 14.61 | 19.78 | 21.39 | 25.52 | 18.80 | 23.21 | 13.19 |
| ENSMUSG00000103309 | 93.61 | 80.50 | 74.48 | 52.72 | 52.92 | 60.54 | 71.94 | 86.38 | 62.82 | 58.62 | 74.69 | 64.05 |
| ENSMUSG00000074071 | 19.50 | 18.65 | 21.56 | 14.47 | 12.96 | 12.53 | 17.09 | 9.05 | 19.63 | 11.06 | 9.08 | 14.13 |
| ENSMUSG00000020475 | 1742.43 | 1371.50 | 1419.07 | 1189.79 | 929.90 | 927.97 | 1482.86 | 1602.58 | 1220.10 | 1318.48 | 1714.77 | 1600.25 |
| ENSMUSG00000002032 | 19.50 | 23.56 | 22.54 | 15.51 | 17.28 | 11.48 | 12.59 | 9.05 | 15.71 | 16.59 | 17.16 | 22.61 |
| ENSMUSG00000105808 | 42.90 | 40.25 | 48.02 | 35.15 | 22.68 | 31.32 | 28.78 | 38.67 | 35.34 | 26.55 | 30.28 | 27.31 |
| ENSMUSG00000035852 | 15.60 | 14.73 | 13.72 | 12.40 | 8.64 | 9.39 | 8.09 | 11.52 | 11.78 | 14.38 | 13.12 | 8.48 |
| ENSMUSG00000082417 | 541.16 | 406.44 | 458.65 | 376.27 | 290.53 | 305.85 | 401.06 | 530.63 | 361.22 | 386.03 | 445.09 | 385.23 |
| ENSMUSG00000085655 | 22.43 | 22.58 | 24.50 | 19.64 | 15.12 | 13.57 | 16.19 | 18.92 | 19.63 | 17.70 | 19.18 | 22.61 |
| ENSMUSG00000074179 | 40.95 | 42.22 | 48.02 | 31.01 | 29.16 | 31.32 | 44.06 | 31.26 | 36.32 | 47.56 | 48.45 | 28.26 |
| ENSMUSG00000081416 | 199.89 | 215.98 | 184.24 | 143.68 | 150.12 | 128.39 | 216.72 | 183.46 | 148.22 | 149.32 | 177.63 | 156.35 |
| ENSMUSG00000057003 | 16845.07 | 18216.27 | 14400.42 | 13008.07 | 10363.95 | 11593.93 | 26967.49 | 24177.72 | 22098.20 | 19273.97 | 23198.34 | 26177.67 |
| ENSMUSG00000071711 | 559.68 | 488.91 | 458.65 | 433.12 | 333.73 | 299.58 | 462.21 | 504.30 | 426.00 | 383.82 | 453.17 | 409.72 |
| ENSMUSG00000098530 | 37.05 | 34.36 | 38.22 | 31.01 | 21.60 | 25.05 | 29.68 | 35.38 | 31.41 | 24.33 | 34.32 | 29.20 |
| ENSMUSG00000045326 | 24.38 | 25.53 | 22.54 | 19.64 | 14.04 | 17.75 | 12.59 | 18.10 | 22.58 | 23.23 | 20.19 | 28.26 |
| ENSMUSG00000021751 | 33.15 | 35.34 | 42.14 | 31.01 | 24.84 | 22.96 | 31.47 | 28.79 | 45.15 | 34.29 | 46.43 | 37.68 |
| ENSMUSG00000023571 | 194.04 | 171.81 | 190.12 | 141.62 | 137.16 | 117.95 | 149.27 | 156.31 | 135.46 | 119.46 | 142.31 | 143.17 |
| ENSMUSG00000048981 | 30.23 | 26.51 | 25.48 | 21.71 | 17.28 | 19.83 | 34.17 | 26.33 | 19.63 | 29.86 | 48.45 | 27.31 |
| ENSMUSG00000108911 | 15.60 | 14.73 | 13.72 | 11.37 | 10.80 | 9.39 | 16.19 | 19.74 | 24.54 | 15.49 | 11.10 | 18.84 |
| ENSMUSG00000022602 | 40.95 | 40.25 | 44.10 | 31.01 | 30.24 | 29.23 | 31.47 | 32.08 | 41.23 | 40.93 | 24.22 | 35.79 |
| ENSMUSG00000062365 | 263.27 | 204.20 | 230.30 | 171.59 | 153.36 | 179.54 | 186.14 | 219.66 | 169.81 | 139.37 | 190.75 | 158.24 |
| ENSMUSG00000086784 | 78.00 | 88.36 | 94.08 | 75.46 | 59.40 | 54.28 | 86.33 | 96.25 | 69.69 | 87.38 | 80.74 | 72.52 |
| ENSMUSG00000030730 | 10236.14 | 8658.99 | 8714.34 | 7241.06 | 5852.67 | 7035.48 | 10166.88 | 9492.90 | 9461.40 | 10071.13 | 12814.85 | 11606.79 |
| ENSMUSG00000079654 | 82.88 | 68.72 | 73.50 | 64.09 | 52.92 | 48.02 | 66.54 | 79.80 | 62.82 | 59.73 | 71.66 | 66.87 |
| ENSMUSG00000003863 | 51.68 | 53.01 | 59.78 | 42.38 | 34.56 | 43.84 | 44.96 | 68.28 | 48.10 | 54.20 | 45.42 | 49.92 |

| | | | | | | | | | | | | |
|---|---|---|---|---|---|---|---|---|---|---|---|---|
| ENSMUSG00000021773 | 198.91 | 168.86 | 172.48 | 148.85 | 111.24 | 136.74 | 125.00 | 166.18 | 145.27 | 137.16 | 168.55 | 143.17 |
| ENSMUSG00000006651 | 250.59 | 207.15 | 219.52 | 175.73 | 149.04 | 173.28 | 209.52 | 238.58 | 194.35 | 201.31 | 194.79 | 191.20 |
| ENSMUSG00000039611 | 178.44 | 161.01 | 154.84 | 126.11 | 126.36 | 111.69 | 211.32 | 223.77 | 166.87 | 165.92 | 167.54 | 160.12 |
| ENSMUSG00000041351 | 120.91 | 109.96 | 104.86 | 71.33 | 83.16 | 92.90 | 149.27 | 127.52 | 136.44 | 109.50 | 124.14 | 97.96 |
| ENSMUSG00000025129 | 5.85 | 4.91 | 4.90 | 4.13 | 3.24 | 4.18 | 1.80 | 2.47 | 1.96 | 3.32 | 1.01 | 2.83 |
| ENSMUSG00000108634 | 212.56 | 179.66 | 198.94 | 146.79 | 163.08 | 127.35 | 170.86 | 219.66 | 151.16 | 153.75 | 163.50 | 198.74 |
| ENSMUSG00000035692 | 80.93 | 73.63 | 72.52 | 66.16 | 47.52 | 54.28 | 68.34 | 73.22 | 62.82 | 58.62 | 66.61 | 58.40 |
| ENSMUSG00000075702 | 1734.63 | 1374.44 | 1463.17 | 1303.50 | 1015.23 | 1064.72 | 1437.00 | 1543.35 | 1115.07 | 1246.59 | 1496.77 | 1241.40 |
| ENSMUSG00000046157 | 170.63 | 173.77 | 166.60 | 132.31 | 116.64 | 129.44 | 182.55 | 170.29 | 168.83 | 168.13 | 150.38 | 151.64 |
| ENSMUSG00000036186 | 278.87 | 239.55 | 268.53 | 217.08 | 184.68 | 181.63 | 196.04 | 226.24 | 213.98 | 223.43 | 220.02 | 200.62 |
| ENSMUSG00000105354 | 9.75 | 9.82 | 8.82 | 6.20 | 7.56 | 7.31 | 9.89 | 4.94 | 9.82 | 9.95 | 9.08 | 8.48 |
| ENSMUSG00000034108 | 296.42 | 246.42 | 258.73 | 216.04 | 200.89 | 178.50 | 226.61 | 294.52 | 231.65 | 264.36 | 237.18 | 215.69 |
| ENSMUSG00000024827 | 110.18 | 92.28 | 107.80 | 66.16 | 78.84 | 85.59 | 110.61 | 100.37 | 85.40 | 102.87 | 98.91 | 116.79 |
| ENSMUSG00000025213 | 80.93 | 70.69 | 87.22 | 68.22 | 57.24 | 52.19 | 55.75 | 64.99 | 49.08 | 58.62 | 67.62 | 65.93 |
| ENSMUSG00000029711 | 107.26 | 120.75 | 100.94 | 90.97 | 66.96 | 87.68 | 70.14 | 84.74 | 83.43 | 86.28 | 82.76 | 81.00 |
| ENSMUSG00000073755 | 131.63 | 137.44 | 158.76 | 117.84 | 98.28 | 106.47 | 117.80 | 116.00 | 142.33 | 150.43 | 112.03 | 119.62 |
| ENSMUSG00000051043 | 2229.95 | 1859.42 | 1919.86 | 1665.29 | 1361.92 | 1532.36 | 2188.77 | 2216.30 | 2076.03 | 2097.19 | 2329.42 | 2200.23 |
| ENSMUSG00000027316 | 275.94 | 247.40 | 225.40 | 186.07 | 198.73 | 185.80 | 252.69 | 290.41 | 199.26 | 225.65 | 231.13 | 226.99 |
| ENSMUSG00000039347 | 282.77 | 234.64 | 253.83 | 213.98 | 191.16 | 182.67 | 250.89 | 275.60 | 230.67 | 244.45 | 284.62 | 253.37 |
| ENSMUSG00000033313 | 174.54 | 150.21 | 170.52 | 132.31 | 123.12 | 123.17 | 154.67 | 147.26 | 131.53 | 119.46 | 159.47 | 128.10 |
| ENSMUSG00000003444 | 561.63 | 475.16 | 487.07 | 408.31 | 398.53 | 359.08 | 483.79 | 547.08 | 450.54 | 456.82 | 469.32 | 446.45 |
| ENSMUSG00000039670 | 90.68 | 81.48 | 76.44 | 73.39 | 60.48 | 56.37 | 65.64 | 75.69 | 63.80 | 69.68 | 73.68 | 69.70 |
| ENSMUSG00000060923 | 213.54 | 218.93 | 196.00 | 172.63 | 155.52 | 153.44 | 152.87 | 148.08 | 146.25 | 172.55 | 168.55 | 151.64 |
| ENSMUSG00000032648 | 2399.61 | 2107.81 | 2013.94 | 1725.24 | 1583.32 | 1705.63 | 2324.55 | 2329.83 | 2161.43 | 2388.09 | 2662.49 | 2605.24 |
| ENSMUSG00000020843 | 1089.14 | 911.06 | 978.06 | 836.26 | 780.86 | 677.45 | 928.92 | 1020.95 | 823.54 | 764.32 | 937.62 | 791.18 |
| ENSMUSG00000046718 | 127.73 | 106.03 | 112.70 | 93.03 | 89.64 | 84.55 | 100.72 | 107.77 | 96.19 | 80.75 | 103.96 | 112.08 |
| ENSMUSG00000025464 | 185.26 | 161.99 | 174.44 | 120.94 | 135.00 | 147.18 | 125.00 | 145.61 | 125.64 | 132.73 | 152.40 | 150.70 |
| ENSMUSG00000098021 | 53.63 | 54.00 | 49.98 | 45.48 | 34.56 | 41.75 | 58.45 | 65.81 | 61.84 | 70.79 | 66.61 | 48.98 |
| ENSMUSG00000049760 | 1116.44 | 993.53 | 986.88 | 873.48 | 799.22 | 721.29 | 970.29 | 1088.40 | 902.07 | 887.10 | 929.55 | 875.01 |
| ENSMUSG00000041241 | 508.00 | 416.26 | 480.21 | 404.18 | 314.29 | 367.43 | 437.93 | 487.03 | 419.13 | 413.69 | 493.54 | 444.57 |
| ENSMUSG00000038208 | 151.13 | 124.68 | 142.10 | 123.01 | 97.20 | 103.34 | 133.99 | 149.73 | 127.60 | 140.48 | 144.33 | 119.62 |
| ENSMUSG00000031312 | 1382.63 | 1270.38 | 1153.48 | 997.52 | 961.22 | 989.56 | 1203.19 | 1413.36 | 1086.60 | 1214.51 | 1236.37 | 1174.52 |
| ENSMUSG00000019659 | 625.99 | 531.12 | 550.77 | 498.24 | 407.17 | 418.58 | 477.50 | 721.49 | 483.92 | 522.08 | 486.47 | 505.79 |
| ENSMUSG00000025538 | 388.07 | 354.41 | 357.71 | 289.44 | 277.57 | 286.01 | 330.02 | 361.98 | 328.83 | 339.58 | 345.17 | 300.46 |

| | | | | | | | | | | | | |
|---|---|---|---|---|---|---|---|---|---|---|---|---|
| ENSMUSG00000020635 | 108.23 | 111.92 | 97.02 | 79.59 | 90.72 | 76.20 | 111.51 | 101.19 | 84.42 | 110.61 | 96.89 | 82.89 |
| ENSMUSG00000032860 | 380.27 | 352.45 | 332.23 | 290.47 | 249.49 | 290.19 | 373.19 | 389.95 | 393.61 | 363.91 | 358.30 | 391.82 |
| ENSMUSG00000033065 | 6402.22 | 5510.53 | 5512.62 | 5106.48 | 4037.14 | 4479.12 | 7283.89 | 7228.06 | 6250.67 | 6231.83 | 7457.58 | 6859.71 |
| ENSMUSG00000021750 | 396.85 | 357.36 | 388.09 | 332.85 | 285.13 | 276.62 | 381.28 | 334.83 | 340.61 | 331.83 | 383.53 | 428.56 |
| ENSMUSG00000031840 | 546.03 | 469.27 | 538.03 | 437.25 | 382.33 | 397.70 | 549.44 | 555.31 | 488.82 | 470.10 | 561.16 | 551.94 |
| ENSMUSG00000042978 | 619.16 | 504.62 | 567.43 | 466.20 | 389.89 | 469.73 | 495.48 | 565.18 | 500.60 | 492.22 | 547.03 | 527.45 |
| ENSMUSG00000029560 | 564.56 | 520.32 | 516.47 | 474.47 | 386.65 | 395.62 | 546.74 | 587.39 | 461.34 | 520.98 | 563.18 | 557.59 |
| ENSMUSG00000028672 | 444.63 | 401.53 | 392.01 | 338.02 | 290.53 | 343.42 | 345.31 | 351.28 | 324.90 | 339.58 | 317.92 | 309.88 |
| ENSMUSG00000074218 | 84.83 | 70.69 | 77.42 | 66.16 | 56.16 | 60.54 | 86.33 | 77.33 | 70.67 | 69.68 | 78.72 | 57.45 |
| ENSMUSG00000003355 | 482.65 | 431.97 | 460.61 | 383.50 | 357.49 | 340.29 | 384.88 | 412.16 | 327.85 | 383.82 | 445.09 | 314.59 |
| ENSMUSG00000051439 | 178.44 | 147.26 | 175.42 | 141.62 | 128.52 | 124.22 | 164.56 | 182.63 | 154.11 | 162.60 | 149.37 | 125.27 |
| ENSMUSG00000031750 | 132.61 | 112.90 | 122.50 | 108.54 | 85.32 | 96.03 | 103.41 | 97.08 | 140.37 | 112.82 | 107.99 | 104.55 |
| ENSMUSG00000054934 | 61.43 | 54.00 | 52.92 | 39.28 | 46.44 | 46.97 | 35.07 | 42.78 | 45.15 | 76.32 | 45.42 | 38.62 |
| ENSMUSG00000072115 | 148.21 | 157.08 | 156.80 | 128.18 | 110.16 | 126.30 | 137.58 | 116.82 | 118.77 | 136.05 | 121.11 | 121.50 |
| ENSMUSG00000030882 | 466.08 | 410.37 | 426.31 | 392.81 | 307.81 | 327.77 | 437.93 | 488.67 | 366.13 | 386.03 | 380.50 | 392.76 |
| ENSMUSG00000025466 | 158.93 | 146.28 | 139.16 | 132.31 | 108.00 | 110.65 | 98.92 | 134.10 | 114.84 | 98.44 | 105.97 | 103.61 |
| ENSMUSG00000034245 | 207.69 | 176.71 | 190.12 | 170.56 | 138.24 | 145.09 | 169.06 | 168.65 | 158.03 | 176.98 | 171.58 | 158.24 |
| ENSMUSG00000027628 | 806.37 | 699.98 | 692.87 | 579.91 | 598.34 | 560.54 | 649.26 | 631.82 | 632.13 | 548.63 | 668.14 | 615.99 |
| ENSMUSG00000022748 | 117.98 | 102.10 | 110.74 | 86.83 | 78.84 | 96.03 | 96.22 | 110.24 | 85.40 | 64.15 | 91.84 | 78.18 |
| ENSMUSG00000033938 | 3732.52 | 3111.15 | 3304.63 | 3015.30 | 2529.43 | 2493.73 | 3493.57 | 4039.36 | 3234.29 | 3096.00 | 3475.97 | 3241.95 |
| ENSMUSG00000029591 | 96.53 | 86.39 | 89.18 | 80.63 | 71.28 | 63.67 | 73.74 | 93.79 | 65.77 | 68.58 | 74.69 | 60.28 |
| ENSMUSG00000034259 | 832.70 | 680.35 | 750.70 | 649.16 | 570.25 | 574.11 | 759.86 | 827.62 | 629.19 | 644.86 | 749.90 | 644.25 |
| ENSMUSG00000040466 | 376.37 | 322.01 | 365.55 | 316.31 | 254.89 | 272.44 | 342.61 | 412.16 | 278.77 | 309.71 | 360.31 | 299.52 |
| ENSMUSG00000021235 | 338.34 | 290.60 | 298.91 | 261.53 | 230.05 | 244.26 | 297.65 | 319.20 | 233.61 | 258.83 | 269.48 | 275.03 |
| ENSMUSG00000090862 | 9424.89 | 7924.64 | 8227.27 | 7072.57 | 6839.82 | 6396.65 | 7035.70 | 8063.08 | 6489.19 | 6571.40 | 7328.40 | 6584.68 |
| ENSMUSG00000083689 | 768.34 | 674.46 | 726.20 | 602.65 | 565.93 | 554.28 | 598.00 | 528.16 | 370.05 | 478.95 | 547.03 | 466.23 |
| ENSMUSG00000079350 | 14.63 | 15.71 | 14.70 | 11.37 | 11.88 | 12.53 | 10.79 | 15.63 | 7.85 | 7.74 | 13.12 | 7.54 |
| ENSMUSG00000030996 | 584.06 | 534.07 | 505.69 | 478.60 | 414.73 | 397.70 | 630.37 | 699.28 | 515.33 | 526.51 | 670.16 | 607.51 |
| ENSMUSG00000079491 | 370.52 | 324.96 | 337.13 | 292.54 | 267.85 | 260.96 | 334.52 | 293.70 | 254.23 | 310.82 | 270.49 | 283.51 |
| ENSMUSG00000053093 | 2136.35 | 2100.93 | 1811.08 | 1786.23 | 1460.20 | 1580.37 | 2161.79 | 2761.73 | 2408.79 | 1999.85 | 2321.35 | 2922.65 |
| ENSMUSG00000005150 | 666.94 | 626.35 | 592.91 | 565.43 | 491.41 | 451.98 | 626.77 | 722.31 | 586.98 | 579.60 | 570.24 | 588.68 |
| ENSMUSG00000059534 | 1634.19 | 1578.65 | 1564.11 | 1416.17 | 1208.55 | 1201.46 | 1375.85 | 1727.63 | 1396.78 | 1415.82 | 1525.02 | 1410.94 |
| ENSMUSG00000092417 | 238.89 | 203.22 | 222.46 | 190.20 | 169.56 | 173.28 | 207.73 | 250.09 | 190.43 | 188.04 | 203.87 | 168.60 |
| ENSMUSG00000035824 | 240.84 | 234.64 | 248.93 | 202.61 | 198.73 | 180.58 | 219.42 | 251.74 | 198.28 | 194.68 | 228.10 | 215.69 |

| | | | | | | | | | | | | |
|---|---|---|---|---|---|---|---|---|---|---|---|---|
| ENSMUSG00000028464 | 48757.69 | 41675.07 | 42994.49 | 38669.66 | 32116.79 | 36448.81 | 46827.35 | 49707.93 | 42645.51 | 45555.15 | 53324.40 | 48263.82 |
| ENSMUSG00000027470 | 273.02 | 267.03 | 263.63 | 237.75 | 195.49 | 213.99 | 256.29 | 272.31 | 266.01 | 246.66 | 343.16 | 294.81 |
| ENSMUSG00000067288 | 12833.69 | 11052.48 | 11546.60 | 10352.50 | 9408.12 | 8800.62 | 8904.34 | 9956.07 | 8114.68 | 8158.67 | 10017.12 | 8647.40 |
| ENSMUSG00000029559 | 673.76 | 587.08 | 628.19 | 520.98 | 490.33 | 511.48 | 597.10 | 669.66 | 537.90 | 566.33 | 630.80 | 476.59 |
| ENSMUSG00000002014 | 3075.33 | 2793.06 | 2786.20 | 2388.88 | 2452.74 | 2135.70 | 2472.03 | 2672.06 | 2277.25 | 2343.85 | 2331.44 | 2363.18 |
| ENSMUSG00000062963 | 1665.40 | 1495.20 | 1590.57 | 1320.03 | 1279.83 | 1232.78 | 1365.95 | 1540.88 | 1230.90 | 1189.07 | 1400.88 | 1228.21 |
| ENSMUSG00000078681 | 343.22 | 310.23 | 306.75 | 265.66 | 272.17 | 236.95 | 295.85 | 317.55 | 294.47 | 303.07 | 315.91 | 299.52 |
| ENSMUSG00000033044 | 415.37 | 381.90 | 359.67 | 313.21 | 333.73 | 287.06 | 296.75 | 389.13 | 313.12 | 379.40 | 401.69 | 399.36 |
| ENSMUSG00000040952 | 11675.32 | 9888.13 | 10789.05 | 9455.25 | 8324.86 | 8344.46 | 9094.08 | 10324.63 | 7563.04 | 7574.64 | 8882.69 | 7487.95 |
| ENSMUSG00000032656 | 56.55 | 59.89 | 52.92 | 41.35 | 46.44 | 49.06 | 62.05 | 50.18 | 51.04 | 51.99 | 52.48 | 49.92 |
| ENSMUSG00000020743 | 296.42 | 278.82 | 281.27 | 254.29 | 223.57 | 216.07 | 264.38 | 264.90 | 251.28 | 271.00 | 256.36 | 243.95 |
| ENSMUSG00000071661 | 51.68 | 43.20 | 49.00 | 39.28 | 35.64 | 41.75 | 51.26 | 40.31 | 49.08 | 34.29 | 35.32 | 46.15 |
| ENSMUSG00000056116 | 976.03 | 870.81 | 896.72 | 790.78 | 712.82 | 722.34 | 875.87 | 819.39 | 767.59 | 777.60 | 789.26 | 865.59 |
| ENSMUSG00000022787 | 137.48 | 139.41 | 131.32 | 105.44 | 104.76 | 121.09 | 119.60 | 139.86 | 110.92 | 130.52 | 125.15 | 118.68 |
| ENSMUSG00000023755 | 90.68 | 79.52 | 86.24 | 71.33 | 69.12 | 67.85 | 70.14 | 92.14 | 59.88 | 56.41 | 66.61 | 58.40 |
| ENSMUSG00000021200 | 190.14 | 168.86 | 168.56 | 145.75 | 129.60 | 153.44 | 123.20 | 156.31 | 123.68 | 171.45 | 154.42 | 158.24 |
| ENSMUSG00000105175 | 233.04 | 264.09 | 269.51 | 222.25 | 189.00 | 211.90 | 211.32 | 261.61 | 255.21 | 266.57 | 218.00 | 245.83 |
| ENSMUSG00000109470 | 377.35 | 325.94 | 358.69 | 315.28 | 270.01 | 278.71 | 365.09 | 430.26 | 445.64 | 350.64 | 361.32 | 403.12 |
| ENSMUSG00000055200 | 215.49 | 185.55 | 216.58 | 155.05 | 165.24 | 182.67 | 182.55 | 170.29 | 155.09 | 189.14 | 168.55 | 198.74 |
| ENSMUSG00000034634 | 41.93 | 35.34 | 40.18 | 33.08 | 33.48 | 29.23 | 31.47 | 36.20 | 38.28 | 29.86 | 28.26 | 27.31 |
| ENSMUSG00000027332 | 1122.29 | 1007.27 | 1067.24 | 914.82 | 827.30 | 865.34 | 1255.35 | 1057.14 | 974.70 | 1058.55 | 1256.56 | 1108.59 |
| ENSMUSG00000023020 | 992.61 | 832.52 | 896.72 | 776.31 | 773.30 | 674.32 | 850.69 | 1020.95 | 785.26 | 814.10 | 868.99 | 853.34 |
| ENSMUSG00000034757 | 993.58 | 848.23 | 922.20 | 818.69 | 704.18 | 736.95 | 895.65 | 911.53 | 826.49 | 817.42 | 878.08 | 788.35 |
| ENSMUSG00000053560 | 1155.44 | 1059.30 | 1118.20 | 967.54 | 873.74 | 884.13 | 882.16 | 1160.80 | 905.01 | 1008.77 | 964.87 | 940.94 |
| ENSMUSG00000023034 | 182.34 | 176.71 | 158.76 | 145.75 | 131.76 | 146.14 | 155.57 | 168.65 | 165.89 | 161.49 | 154.42 | 178.02 |
| ENSMUSG00000013155 | 186.24 | 169.84 | 174.44 | 159.19 | 137.16 | 137.79 | 148.38 | 216.36 | 160.98 | 122.78 | 153.41 | 144.11 |
| ENSMUSG00000089847 | 722.52 | 621.44 | 679.15 | 604.71 | 522.73 | 528.18 | 649.26 | 730.54 | 541.83 | 575.18 | 621.72 | 592.44 |
| ENSMUSG00000025533 | 623.06 | 541.92 | 601.73 | 524.09 | 446.05 | 477.03 | 678.03 | 709.15 | 604.65 | 615.00 | 622.73 | 616.93 |
| ENSMUSG00000020549 | 1144.72 | 990.58 | 1008.44 | 905.52 | 807.86 | 866.39 | 1079.10 | 1065.37 | 1019.86 | 1004.35 | 1057.73 | 1072.80 |
| ENSMUSG00000059363 | 180.39 | 169.84 | 189.14 | 158.16 | 141.48 | 143.01 | 187.04 | 188.39 | 161.96 | 160.39 | 159.47 | 151.64 |
| ENSMUSG00000030609 | 992.61 | 851.17 | 948.66 | 834.20 | 689.06 | 769.31 | 922.63 | 1039.87 | 903.05 | 898.16 | 1009.28 | 1000.28 |
| ENSMUSG00000043333 | 75.08 | 65.78 | 70.56 | 52.72 | 59.40 | 61.59 | 90.82 | 87.20 | 82.45 | 70.79 | 85.79 | 67.82 |
| ENSMUSG00000067148 | 681.56 | 591.99 | 654.65 | 554.06 | 524.89 | 506.26 | 589.01 | 721.49 | 547.72 | 549.74 | 597.49 | 570.78 |
| ENSMUSG00000021361 | 1206.14 | 1120.17 | 1171.12 | 1027.50 | 939.62 | 909.18 | 1027.84 | 1211.81 | 931.51 | 945.72 | 1046.63 | 986.15 |

| | | | | | | | | | | | | |
|---|---|---|---|---|---|---|---|---|---|---|---|---|
| ENSMUSG00000027430 | 271.07 | 240.53 | 266.57 | 224.31 | 194.41 | 221.29 | 267.08 | 232.82 | 230.67 | 234.50 | 234.15 | 249.60 |
| ENSMUSG00000020340 | 927.28 | 984.69 | 917.30 | 775.27 | 730.10 | 826.72 | 1232.87 | 1063.72 | 1146.48 | 1037.53 | 1219.21 | 1208.43 |
| ENSMUSG00000032299 | 497.28 | 438.84 | 490.99 | 392.81 | 384.49 | 399.79 | 402.86 | 500.19 | 384.78 | 365.02 | 413.81 | 373.93 |
| ENSMUSG00000031762 | 5034.22 | 4500.32 | 4618.84 | 4146.17 | 3754.18 | 3781.83 | 3020.57 | 4343.75 | 3809.49 | 3484.25 | 3937.21 | 3536.76 |
| ENSMUSG00000020067 | 449.50 | 500.69 | 463.55 | 380.40 | 363.97 | 422.76 | 708.61 | 547.08 | 598.76 | 580.71 | 623.74 | 633.89 |
| ENSMUSG00000012848 | 13928.68 | 12423.00 | 12495.26 | 11803.81 | 10606.96 | 9696.23 | 12277.41 | 13431.06 | 11420.63 | 10644.10 | 12381.87 | 11467.39 |
| ENSMUSG00000057219 | 158.93 | 157.08 | 156.80 | 137.48 | 123.12 | 130.48 | 148.38 | 159.60 | 123.68 | 150.43 | 156.44 | 143.17 |
| ENSMUSG00000040048 | 3187.46 | 2775.39 | 2877.34 | 2692.79 | 2262.66 | 2364.30 | 3016.07 | 3361.47 | 2764.12 | 2851.55 | 2995.55 | 2847.30 |
| ENSMUSG00000036376 | 338.34 | 345.57 | 325.37 | 289.44 | 272.17 | 274.53 | 341.71 | 373.50 | 339.62 | 327.41 | 341.14 | 336.25 |
| ENSMUSG00000035203 | 1652.72 | 1412.73 | 1503.35 | 1373.79 | 1179.39 | 1233.82 | 1516.13 | 1591.06 | 1516.53 | 1529.75 | 1544.20 | 1498.53 |
| ENSMUSG00000033739 | 240.84 | 222.86 | 230.30 | 207.77 | 191.16 | 176.41 | 206.83 | 219.66 | 217.91 | 212.37 | 204.88 | 199.68 |
| ENSMUSG00000024902 | 757.62 | 646.97 | 682.09 | 561.30 | 617.78 | 553.24 | 602.49 | 678.71 | 462.32 | 525.40 | 609.61 | 521.80 |
| ENSMUSG00000073435 | 185.26 | 168.86 | 159.74 | 150.92 | 130.68 | 145.09 | 164.56 | 191.68 | 126.62 | 160.39 | 158.46 | 155.41 |
| ENSMUSG00000022856 | 212.56 | 199.29 | 181.30 | 151.95 | 166.32 | 174.32 | 161.86 | 193.33 | 153.13 | 149.32 | 170.57 | 194.97 |
| ENSMUSG00000061518 | 5197.05 | 4469.88 | 4628.64 | 4169.94 | 3786.58 | 3922.75 | 4489.04 | 5451.90 | 4148.14 | 4306.09 | 4598.29 | 4337.36 |
| ENSMUSG00000023089 | 1608.84 | 1703.33 | 1822.84 | 1453.38 | 1508.80 | 1306.89 | 1512.53 | 1712.00 | 1311.38 | 1360.52 | 1531.08 | 1413.76 |
| ENSMUSG00000032383 | 8547.34 | 7635.03 | 7899.95 | 7127.36 | 6528.77 | 6382.04 | 7169.69 | 8281.91 | 7046.73 | 7343.47 | 7195.17 | 7204.44 |
| ENSMUSG00000028648 | 2607.30 | 2562.35 | 2635.28 | 2304.12 | 2012.09 | 2178.49 | 2478.32 | 2844.00 | 2237.01 | 2262.00 | 2508.07 | 2153.14 |
| ENSMUSG00000098140 | 109.21 | 110.94 | 101.92 | 98.20 | 84.24 | 85.59 | 106.11 | 116.00 | 80.49 | 117.25 | 107.99 | 111.14 |
| ENSMUSG00000031388 | 1020.88 | 878.66 | 933.96 | 826.96 | 748.46 | 786.01 | 868.67 | 924.69 | 834.34 | 890.42 | 931.57 | 818.49 |
| ENSMUSG00000027597 | 2643.38 | 2372.88 | 2397.13 | 2272.07 | 1998.05 | 1914.40 | 2389.30 | 2642.45 | 2341.06 | 2106.03 | 2324.38 | 2171.98 |
| ENSMUSG00000025869 | 1001.38 | 943.46 | 926.12 | 784.58 | 805.70 | 805.84 | 862.38 | 995.44 | 817.65 | 817.42 | 848.81 | 828.85 |
| ENSMUSG00000024869 | 306.17 | 276.85 | 282.25 | 255.32 | 228.97 | 238.00 | 278.77 | 338.94 | 231.65 | 275.42 | 274.52 | 252.42 |
| ENSMUSG00000015289 | 388.07 | 345.57 | 356.73 | 320.45 | 316.45 | 273.49 | 340.81 | 374.32 | 300.36 | 305.29 | 292.69 | 290.10 |
| ENSMUSG00000029535 | 460.23 | 408.41 | 427.29 | 381.44 | 355.33 | 345.51 | 408.26 | 442.60 | 350.42 | 347.32 | 382.52 | 364.51 |
| ENSMUSG00000038312 | 755.67 | 663.66 | 682.09 | 640.89 | 544.33 | 569.94 | 652.85 | 728.07 | 639.01 | 662.56 | 671.17 | 603.75 |
| ENSMUSG00000033032 | 992.61 | 874.73 | 890.84 | 801.12 | 740.90 | 762.00 | 1220.28 | 1168.20 | 1034.58 | 884.89 | 1131.41 | 1020.06 |
| ENSMUSG00000010609 | 399.77 | 349.50 | 355.75 | 331.82 | 309.97 | 281.84 | 358.80 | 375.96 | 310.18 | 337.36 | 347.19 | 380.52 |
| ENSMUSG00000042320 | 50.70 | 54.00 | 58.80 | 47.55 | 45.36 | 43.84 | 61.15 | 48.54 | 53.01 | 58.62 | 56.52 | 52.75 |
| ENSMUSG00000085401 | 285.69 | 250.34 | 260.69 | 228.45 | 213.85 | 224.43 | 254.49 | 250.92 | 229.69 | 224.54 | 214.98 | 226.99 |
| ENSMUSG00000034371 | 368.57 | 333.79 | 336.15 | 320.45 | 273.25 | 275.57 | 327.33 | 324.96 | 293.49 | 309.71 | 356.28 | 288.22 |
| ENSMUSG00000074884 | 10616.41 | 9264.73 | 9825.69 | 8842.27 | 8100.21 | 7937.36 | 8768.55 | 9968.41 | 8404.25 | 8942.90 | 9639.65 | 9009.08 |
| ENSMUSG00000020471 | 740.07 | 666.60 | 644.85 | 619.19 | 533.53 | 565.76 | 611.49 | 686.94 | 664.53 | 548.63 | 648.97 | 620.70 |
| ENSMUSG00000009687 | 769.32 | 664.64 | 703.65 | 602.65 | 565.93 | 622.13 | 531.45 | 670.48 | 579.13 | 637.12 | 643.92 | 601.86 |

| | | | | | | | | | | | | |
|---|---|---|---|---|---|---|---|---|---|---|---|---|
| ENSMUSG00000013150 | 130.66 | 119.77 | 131.32 | 107.50 | 111.24 | 101.25 | 106.11 | 152.20 | 128.59 | 142.69 | 119.10 | 122.44 |
| ENSMUSG00000079610 | 174.54 | 153.15 | 157.78 | 141.62 | 138.24 | 127.35 | 120.50 | 144.79 | 120.73 | 147.11 | 153.41 | 137.51 |
| ENSMUSG00000020872 | 76.05 | 68.72 | 70.56 | 59.95 | 60.48 | 60.54 | 122.30 | 133.27 | 108.95 | 73.00 | 134.23 | 104.55 |
| ENSMUSG00000032044 | 226.21 | 229.73 | 215.60 | 199.50 | 180.36 | 184.76 | 201.43 | 222.12 | 189.44 | 197.99 | 188.74 | 214.75 |
| ENSMUSG00000029198 | 1576.67 | 1409.79 | 1424.95 | 1314.87 | 1252.83 | 1146.14 | 1428.00 | 1487.40 | 1299.61 | 1349.45 | 1404.92 | 1358.19 |
| ENSMUSG00000035642 | 821.97 | 828.59 | 864.38 | 728.76 | 722.54 | 667.01 | 857.88 | 914.82 | 750.90 | 799.72 | 838.71 | 792.12 |
| ENSMUSG00000009487 | 436.83 | 407.42 | 447.87 | 395.91 | 359.65 | 334.03 | 531.45 | 515.82 | 448.58 | 415.90 | 443.07 | 524.63 |
| ENSMUSG00000024217 | 2201.68 | 1972.33 | 2014.92 | 1807.94 | 1743.17 | 1669.10 | 1921.69 | 2101.12 | 1889.53 | 1839.46 | 1971.13 | 1829.13 |
| ENSMUSG00000020424 | 72.15 | 64.80 | 68.60 | 63.06 | 54.00 | 56.37 | 56.65 | 58.41 | 52.02 | 64.15 | 59.55 | 62.16 |
| ENSMUSG00000002205 | 378.32 | 355.39 | 336.15 | 305.98 | 303.49 | 293.32 | 300.35 | 325.78 | 294.47 | 325.20 | 314.90 | 291.98 |
| ENSMUSG00000101892 | 979.93 | 882.59 | 981.00 | 813.52 | 815.42 | 771.40 | 906.44 | 960.07 | 794.09 | 850.60 | 898.26 | 882.54 |
| ENSMUSG00000024866 | 117.01 | 104.06 | 103.88 | 96.13 | 90.72 | 87.68 | 104.31 | 110.24 | 96.19 | 98.44 | 103.96 | 131.86 |
| ENSMUSG00000062006 | 11567.09 | 10375.08 | 10774.35 | 10051.69 | 9156.48 | 8469.72 | 10283.78 | 10837.98 | 9282.76 | 8971.66 | 9931.33 | 9430.10 |
| ENSMUSG00000011114 | 2417.16 | 2152.97 | 2259.93 | 1988.84 | 1904.09 | 1887.26 | 2061.97 | 2329.83 | 1941.56 | 2049.62 | 2057.93 | 1993.96 |
| ENSMUSG00000060636 | 10774.37 | 9436.53 | 10348.04 | 8359.53 | 9265.56 | 8245.29 | 8642.66 | 9531.56 | 7547.33 | 7591.23 | 8022.78 | 7510.55 |
| ENSMUSG00000025260 | 1167.14 | 1096.61 | 1079.98 | 986.15 | 968.79 | 877.87 | 982.88 | 1166.56 | 935.44 | 955.68 | 992.12 | 998.39 |
| ENSMUSG00000018796 | 747.87 | 743.18 | 746.78 | 639.86 | 616.70 | 640.92 | 770.65 | 723.96 | 753.85 | 700.17 | 797.33 | 826.97 |
| ENSMUSG00000105518 | 15216.73 | 13608.95 | 15385.35 | 13546.63 | 12110.35 | 11863.24 | 14872.63 | 16823.79 | 14659.82 | 14577.43 | 14850.57 | 14082.99 |
| ENSMUSG00000026343 | 161.86 | 172.79 | 185.22 | 152.99 | 146.88 | 141.96 | 107.01 | 176.88 | 176.68 | 153.75 | 136.25 | 145.05 |
| ENSMUSG00000011254 | 412.45 | 370.12 | 412.59 | 362.83 | 318.61 | 335.07 | 357.00 | 433.55 | 371.04 | 383.82 | 341.14 | 330.60 |
| ENSMUSG00000034463 | 337.37 | 300.41 | 320.47 | 281.17 | 259.21 | 276.62 | 273.37 | 315.91 | 287.60 | 308.60 | 337.10 | 301.40 |
| ENSMUSG00000022235 | 488.50 | 496.76 | 509.61 | 442.42 | 411.49 | 421.71 | 419.95 | 436.02 | 405.39 | 464.57 | 581.35 | 518.98 |
| ENSMUSG00000020921 | 209.64 | 205.18 | 192.08 | 186.07 | 162.00 | 170.15 | 189.74 | 212.25 | 184.54 | 183.61 | 207.91 | 189.32 |
| ENSMUSG00000028573 | 268.14 | 246.42 | 248.93 | 229.48 | 209.53 | 212.94 | 287.76 | 282.18 | 234.60 | 245.56 | 299.76 | 255.25 |
| ENSMUSG00000015290 | 1235.40 | 1131.95 | 1118.20 | 1014.06 | 1022.79 | 940.50 | 1110.57 | 1163.27 | 990.41 | 1043.06 | 1065.80 | 1065.27 |
| ENSMUSG00000021131 | 3464.37 | 3390.95 | 3464.37 | 3020.47 | 2924.72 | 2870.56 | 3088.01 | 3688.07 | 2971.23 | 3021.89 | 2966.28 | 3059.23 |
| ENSMUSG00000026179 | 698.14 | 609.66 | 664.45 | 601.61 | 524.89 | 559.50 | 606.09 | 675.42 | 586.98 | 578.50 | 703.47 | 630.12 |
| ENSMUSG00000069920 | 487.53 | 444.73 | 431.21 | 411.41 | 390.97 | 363.26 | 380.38 | 417.10 | 372.02 | 350.64 | 420.87 | 405.01 |
| ENSMUSG00000021417 | 1522.06 | 1387.21 | 1422.99 | 1290.06 | 1216.11 | 1198.33 | 1252.65 | 1299.01 | 1146.48 | 1193.49 | 1306.01 | 1188.65 |
| ENSMUSG00000042616 | 206.71 | 221.87 | 211.68 | 169.53 | 197.65 | 180.58 | 205.03 | 228.70 | 197.30 | 204.63 | 186.72 | 173.31 |
| ENSMUSG00000062867 | 3916.80 | 3644.24 | 3708.40 | 3381.23 | 3013.28 | 3247.39 | 3399.15 | 3636.24 | 3392.32 | 3308.38 | 3451.74 | 3423.73 |
| ENSMUSG00000106392 | 54.60 | 54.98 | 53.90 | 46.52 | 49.68 | 43.84 | 52.16 | 55.12 | 39.26 | 40.93 | 42.39 | 34.85 |
| ENSMUSG00000041035 | 352.97 | 324.96 | 329.29 | 294.60 | 287.29 | 281.84 | 356.10 | 319.20 | 330.79 | 289.80 | 348.20 | 319.30 |
| ENSMUSG00000081895 | 4940.61 | 4367.78 | 4673.72 | 4101.72 | 4028.50 | 3876.82 | 4378.43 | 4570.81 | 3753.54 | 3875.81 | 4086.58 | 3943.65 |

| | | | | | | | | | | | | |
|---|---|---|---|---|---|---|---|---|---|---|---|---|
| ENSMUSG00000025512 | 862.92 | 806.99 | 755.60 | 738.06 | 666.38 | 678.50 | 812.92 | 855.59 | 809.80 | 684.68 | 816.51 | 785.53 |
| ENSMUSG00000028837 | 2295.28 | 2052.83 | 2111.94 | 1953.69 | 1866.29 | 1733.82 | 2046.68 | 2199.02 | 1892.48 | 1903.62 | 1926.72 | 1914.84 |
| ENSMUSG00000073616 | 2166.58 | 2071.48 | 2096.26 | 1856.52 | 1905.17 | 1685.80 | 1980.14 | 2102.77 | 1818.86 | 1806.28 | 1994.34 | 1799.93 |
| ENSMUSG00000034892 | 15558.00 | 15018.73 | 14637.59 | 13525.96 | 13453.91 | 11991.64 | 12184.79 | 13524.85 | 11523.69 | 11184.99 | 13920.02 | 12044.76 |
| ENSMUSG00000061613 | 2016.42 | 1979.20 | 2000.22 | 1822.41 | 1644.88 | 1703.55 | 1798.49 | 2154.60 | 1801.19 | 1948.97 | 1831.85 | 1917.67 |
| ENSMUSG00000105852 | 538.23 | 590.03 | 567.43 | 514.78 | 455.77 | 492.69 | 549.44 | 523.22 | 510.42 | 511.02 | 461.24 | 546.29 |
| ENSMUSG00000035674 | 1216.87 | 1147.66 | 1093.70 | 1027.50 | 1035.75 | 921.71 | 1031.44 | 1065.37 | 952.13 | 945.72 | 1122.32 | 935.29 |
| ENSMUSG00000050856 | 2491.27 | 2461.23 | 2338.33 | 2250.36 | 2044.49 | 1998.95 | 2217.54 | 2634.22 | 2008.30 | 2237.66 | 2437.42 | 2260.51 |
| ENSMUSG00000071041 | 1690.75 | 1589.44 | 1528.83 | 1460.62 | 1346.79 | 1346.55 | 1485.55 | 1592.71 | 1557.76 | 1509.84 | 1524.02 | 1569.17 |
| ENSMUSG00000060477 | 522.63 | 538.98 | 503.73 | 462.06 | 444.97 | 446.76 | 536.85 | 526.51 | 529.07 | 485.58 | 483.45 | 513.32 |
| ENSMUSG00000108204 | 727.39 | 670.53 | 671.31 | 595.41 | 575.65 | 623.17 | 562.93 | 606.31 | 490.79 | 515.45 | 593.46 | 484.13 |
| ENSMUSG00000020736 | 633.79 | 576.28 | 612.51 | 537.52 | 513.01 | 530.27 | 589.91 | 656.50 | 523.18 | 514.34 | 632.82 | 511.44 |
| ENSMUSG00000031389 | 537.26 | 482.04 | 492.95 | 454.83 | 411.49 | 445.72 | 475.70 | 432.73 | 432.87 | 511.02 | 512.72 | 452.10 |
| ENSMUSG00000039725 | 353.95 | 321.03 | 343.01 | 299.77 | 312.13 | 271.40 | 291.36 | 328.25 | 262.08 | 267.68 | 273.52 | 309.88 |
| ENSMUSG00000025428 | 13956.96 | 12549.64 | 13187.16 | 12000.22 | 11365.13 | 11089.76 | 12877.21 | 13221.28 | 12045.89 | 12384.01 | 12997.53 | 12635.32 |
| ENSMUSG00000028607 | 938.98 | 874.73 | 899.66 | 803.18 | 754.94 | 797.49 | 847.09 | 825.15 | 817.65 | 852.81 | 901.29 | 904.20 |
| ENSMUSG00000081176 | 2575.12 | 2371.90 | 2388.31 | 2158.36 | 2178.42 | 2032.36 | 2459.44 | 2301.03 | 2004.38 | 1985.47 | 2130.59 | 1934.62 |
| ENSMUSG00000057506 | 610.39 | 601.81 | 637.99 | 559.23 | 532.45 | 515.66 | 629.47 | 641.69 | 536.92 | 527.61 | 570.24 | 593.38 |
| ENSMUSG00000039911 | 833.67 | 816.81 | 782.06 | 740.13 | 703.10 | 670.15 | 843.49 | 807.05 | 812.74 | 734.46 | 803.39 | 786.47 |
| ENSMUSG00000033918 | 1055.99 | 949.35 | 969.24 | 829.03 | 874.82 | 882.04 | 928.02 | 1025.88 | 819.62 | 932.45 | 863.95 | 865.59 |
| ENSMUSG00000032508 | 437.80 | 399.57 | 397.89 | 361.79 | 353.17 | 360.12 | 365.09 | 371.85 | 342.57 | 330.73 | 410.78 | 381.46 |
| ENSMUSG00000005882 | 922.40 | 895.35 | 924.16 | 838.33 | 744.14 | 805.84 | 932.52 | 852.30 | 859.86 | 849.49 | 886.15 | 892.90 |
| ENSMUSG00000057278 | 1767.78 | 1834.88 | 1682.70 | 1543.31 | 1629.76 | 1431.10 | 1392.03 | 1669.22 | 1433.10 | 1362.73 | 1493.74 | 1379.85 |
| ENSMUSG00000025147 | 707.89 | 647.95 | 686.01 | 616.08 | 598.34 | 564.72 | 749.07 | 737.12 | 654.71 | 686.89 | 701.45 | 680.04 |
| ENSMUSG00000033207 | 1283.17 | 1281.18 | 1190.73 | 1053.34 | 1118.91 | 1100.21 | 1927.08 | 1591.06 | 1587.21 | 1348.35 | 1493.74 | 1486.29 |
| ENSMUSG00000024191 | 495.33 | 460.44 | 478.25 | 407.28 | 432.01 | 411.27 | 392.97 | 458.23 | 417.17 | 439.13 | 453.17 | 428.56 |
| ENSMUSG00000035215 | 537.26 | 492.84 | 508.63 | 457.93 | 425.53 | 459.29 | 487.39 | 588.22 | 440.73 | 431.38 | 486.47 | 410.66 |
| ENSMUSG00000026816 | 876.58 | 796.20 | 806.56 | 738.06 | 678.26 | 747.39 | 726.59 | 845.71 | 790.17 | 725.61 | 810.45 | 768.57 |
| ENSMUSG00000075312 | 494.35 | 479.09 | 482.17 | 427.95 | 411.49 | 431.11 | 544.04 | 554.49 | 499.62 | 558.59 | 512.72 | 529.34 |
| ENSMUSG00000090124 | 777.12 | 718.64 | 757.56 | 631.59 | 695.54 | 643.01 | 590.80 | 575.88 | 621.34 | 663.67 | 661.08 | 591.50 |
| ENSMUSG00000004270 | 1678.07 | 1522.69 | 1551.37 | 1442.01 | 1337.07 | 1381.00 | 1669.90 | 1715.29 | 1523.40 | 1515.37 | 1673.39 | 1620.98 |
| ENSMUSG00000040666 | 552.86 | 541.92 | 541.95 | 482.74 | 486.01 | 464.51 | 615.98 | 593.15 | 491.77 | 551.95 | 637.87 | 647.07 |
| ENSMUSG00000029513 | 821.00 | 744.16 | 774.22 | 727.72 | 676.10 | 645.09 | 752.67 | 811.98 | 733.24 | 759.90 | 738.79 | 767.63 |
| ENSMUSG00000025487 | 2796.46 | 2506.39 | 2715.64 | 2472.61 | 2248.62 | 2302.71 | 2731.01 | 2922.16 | 2397.01 | 2344.95 | 2463.66 | 2468.67 |

| | | | | | | | | | | | | |
|---|---|---|---|---|---|---|---|---|---|---|---|---|
| ENSMUSG00000057375 | 563.58 | 553.70 | 572.33 | 491.01 | 482.77 | 507.31 | 480.20 | 566.00 | 479.99 | 440.23 | 516.75 | 459.64 |
| ENSMUSG00000036678 | 741.04 | 681.33 | 739.92 | 653.30 | 600.50 | 641.96 | 705.01 | 774.14 | 723.42 | 662.56 | 685.30 | 665.91 |
| ENSMUSG00000054545 | 772.24 | 718.64 | 764.42 | 634.69 | 696.62 | 648.22 | 590.80 | 579.99 | 621.34 | 659.24 | 654.01 | 583.02 |
| ENSMUSG00000028884 | 501.18 | 454.55 | 463.55 | 435.19 | 387.73 | 424.84 | 388.47 | 494.43 | 428.95 | 429.17 | 441.06 | 407.83 |
| ENSMUSG00000025728 | 1334.85 | 1264.49 | 1222.09 | 1158.78 | 1040.07 | 1162.84 | 1195.10 | 1201.11 | 1318.26 | 1273.13 | 1200.04 | 1231.98 |
| ENSMUSG00000052934 | 422.20 | 402.52 | 387.11 | 378.33 | 356.41 | 331.94 | 406.46 | 412.99 | 363.18 | 376.08 | 455.19 | 448.33 |
| ENSMUSG00000022244 | 1501.59 | 1351.86 | 1480.81 | 1326.24 | 1275.51 | 1218.16 | 1459.48 | 1411.72 | 1364.39 | 1368.26 | 1501.81 | 1366.67 |
| ENSMUSG00000002741 | 2545.87 | 2317.90 | 2467.69 | 2274.14 | 2122.26 | 2066.80 | 2219.34 | 2419.50 | 2220.32 | 2204.48 | 2281.99 | 2295.36 |
| ENSMUSG00000039826 | 602.58 | 567.45 | 604.67 | 547.86 | 516.25 | 501.04 | 562.03 | 545.44 | 559.50 | 556.37 | 568.23 | 562.30 |
| ENSMUSG00000029363 | 536.28 | 530.14 | 541.95 | 502.38 | 465.49 | 453.03 | 513.47 | 575.88 | 484.90 | 444.66 | 438.03 | 468.11 |
| ENSMUSG00000040323 | 400.75 | 385.83 | 399.85 | 370.06 | 330.49 | 347.60 | 404.66 | 396.53 | 385.76 | 365.02 | 371.42 | 398.42 |
| ENSMUSG00000029388 | 1258.80 | 1200.67 | 1226.01 | 1105.02 | 1107.03 | 1044.88 | 1187.00 | 1206.05 | 1188.69 | 1139.29 | 1186.92 | 1164.16 |
| ENSMUSG00000024785 | 473.88 | 433.93 | 442.97 | 406.24 | 374.77 | 414.40 | 391.17 | 491.14 | 444.65 | 436.91 | 401.69 | 403.12 |
| ENSMUSG00000040557 | 176.49 | 170.82 | 166.60 | 162.29 | 144.72 | 148.23 | 187.94 | 184.28 | 157.05 | 173.66 | 186.72 | 164.83 |
| ENSMUSG00000051355 | 1693.67 | 1567.85 | 1666.04 | 1481.29 | 1495.84 | 1392.48 | 1478.36 | 1637.13 | 1293.72 | 1217.83 | 1444.28 | 1282.84 |
| ENSMUSG00000034432 | 1853.58 | 1696.46 | 1738.56 | 1646.68 | 1562.80 | 1481.21 | 1843.45 | 1809.89 | 1663.77 | 1691.24 | 1692.57 | 1648.29 |
| ENSMUSG00000025582 | 643.54 | 621.44 | 685.03 | 576.80 | 567.01 | 590.81 | 691.52 | 688.58 | 655.69 | 595.09 | 733.75 | 653.66 |
| ENSMUSG00000087260 | 1462.58 | 1339.10 | 1419.07 | 1238.37 | 1321.95 | 1195.20 | 1410.02 | 1489.87 | 1337.89 | 1304.10 | 1275.73 | 1325.23 |
| ENSMUSG00000030706 | 662.06 | 690.17 | 690.91 | 566.47 | 627.50 | 625.26 | 645.66 | 733.01 | 625.26 | 592.88 | 629.79 | 635.77 |
| ENSMUSG00000033554 | 271.07 | 285.69 | 269.51 | 228.45 | 252.73 | 254.70 | 269.77 | 315.09 | 263.06 | 279.85 | 264.43 | 259.96 |
| ENSMUSG00000078765 | 501.18 | 476.15 | 510.59 | 458.96 | 448.21 | 418.58 | 522.46 | 547.90 | 465.27 | 460.14 | 442.07 | 464.35 |
| ENSMUSG00000027487 | 358.82 | 334.77 | 369.47 | 316.31 | 330.49 | 300.63 | 303.05 | 334.01 | 315.09 | 336.26 | 318.93 | 341.90 |
| ENSMUSG00000105061 | 206.71 | 202.24 | 189.14 | 175.73 | 169.56 | 187.89 | 183.45 | 180.99 | 196.32 | 158.17 | 184.70 | 181.78 |
| ENSMUSG00000105194 | 206.71 | 202.24 | 189.14 | 175.73 | 169.56 | 187.89 | 183.45 | 180.99 | 196.32 | 158.17 | 184.70 | 181.78 |
| ENSMUSG00000081824 | 797.60 | 740.24 | 787.94 | 724.62 | 691.22 | 657.62 | 844.39 | 980.63 | 869.68 | 837.33 | 881.10 | 786.47 |
| ENSMUSG00000004070 | 1379.70 | 1327.32 | 1341.65 | 1239.41 | 1218.27 | 1158.66 | 1263.44 | 1351.66 | 1224.02 | 1226.68 | 1275.73 | 1253.64 |
| ENSMUSG00000084060 | 157.96 | 163.95 | 148.96 | 139.55 | 133.92 | 147.18 | 148.38 | 142.32 | 117.79 | 140.48 | 129.19 | 110.20 |
| ENSMUSG00000018449 | 181.36 | 176.71 | 193.06 | 162.29 | 167.40 | 162.84 | 147.48 | 157.95 | 160.98 | 140.48 | 136.25 | 153.53 |
| ENSMUSG00000039356 | 391.97 | 372.08 | 401.81 | 352.49 | 331.57 | 358.04 | 341.71 | 423.68 | 375.94 | 338.47 | 339.12 | 347.55 |
| ENSMUSG00000095991 | 292.52 | 317.10 | 310.67 | 263.59 | 273.25 | 286.01 | 333.62 | 274.77 | 263.06 | 280.95 | 336.09 | 277.85 |
| ENSMUSG00000004460 | 2732.11 | 2560.39 | 2554.91 | 2285.51 | 2370.66 | 2363.25 | 2554.76 | 2649.03 | 2444.12 | 2484.32 | 2639.27 | 2530.83 |
| ENSMUSG00000062997 | 19887.25 | 18477.42 | 18851.68 | 17090.15 | 17427.33 | 16745.28 | 16834.79 | 17639.07 | 14553.81 | 14834.04 | 17615.00 | 15250.92 |
| ENSMUSG00000027710 | 822.95 | 853.14 | 849.68 | 773.21 | 723.62 | 767.22 | 861.48 | 821.86 | 816.67 | 727.82 | 819.54 | 835.45 |
| ENSMUSG00000031543 | 2400.59 | 2287.47 | 2222.69 | 2141.83 | 2014.25 | 2038.62 | 2523.28 | 2669.59 | 2467.68 | 2511.98 | 2606.98 | 2647.63 |

| | | | | | | | | | | | | |
|---|---|---|---|---|---|---|---|---|---|---|---|---|
| ENSMUSG00000105053 | 388.07 | 368.15 | 383.19 | 352.49 | 328.33 | 341.34 | 362.40 | 364.45 | 354.35 | 321.88 | 358.30 | 342.84 |
| ENSMUSG00000029642 | 2098.32 | 1949.75 | 2059.02 | 1929.92 | 1772.33 | 1779.75 | 1954.06 | 2119.22 | 1747.20 | 1753.18 | 1886.35 | 1774.50 |
| ENSMUSG00000030801 | 397.82 | 396.62 | 387.11 | 370.06 | 338.05 | 352.82 | 430.74 | 390.77 | 396.56 | 383.82 | 469.32 | 425.73 |
| ENSMUSG00000040681 | 4474.53 | 4310.84 | 4474.77 | 3850.53 | 3991.78 | 4089.77 | 3955.78 | 4625.10 | 3773.17 | 3814.97 | 3743.43 | 3623.41 |
| ENSMUSG00000028882 | 893.15 | 890.44 | 834.98 | 793.88 | 818.66 | 744.26 | 850.69 | 883.56 | 812.74 | 840.64 | 885.14 | 850.52 |
| ENSMUSG00000095512 | 427.07 | 445.71 | 453.75 | 376.27 | 407.17 | 411.27 | 439.73 | 416.28 | 386.74 | 414.79 | 439.04 | 407.83 |
| ENSMUSG00000096579 | 426.10 | 443.75 | 452.77 | 376.27 | 405.01 | 411.27 | 437.93 | 414.63 | 385.76 | 412.58 | 438.03 | 406.89 |
| ENSMUSG00000029319 | 728.37 | 702.93 | 703.65 | 666.74 | 610.22 | 648.22 | 691.52 | 756.04 | 660.60 | 636.01 | 758.98 | 711.12 |
| ENSMUSG00000038213 | 288.62 | 267.03 | 279.31 | 253.26 | 247.33 | 252.61 | 265.28 | 284.65 | 281.71 | 248.87 | 277.55 | 277.85 |
| ENSMUSG00000022066 | 1293.90 | 1236.02 | 1237.77 | 1127.77 | 1129.71 | 1147.18 | 1245.46 | 1295.72 | 1252.49 | 1268.71 | 1331.24 | 1335.59 |
| ENSMUSG00000095463 | 1292.92 | 1232.09 | 1233.85 | 1123.63 | 1126.47 | 1147.18 | 1241.86 | 1290.78 | 1246.60 | 1263.18 | 1326.20 | 1336.53 |
| ENSMUSG00000026806 | 285.69 | 310.23 | 301.85 | 282.20 | 262.45 | 268.27 | 248.19 | 292.87 | 260.12 | 245.56 | 273.52 | 281.62 |
| ENSMUSG00000048440 | 336.39 | 322.01 | 309.69 | 280.13 | 299.17 | 297.49 | 261.68 | 334.83 | 314.10 | 286.48 | 290.67 | 295.75 |
| ENSMUSG00000071415 | 16531.10 | 15789.40 | 16446.71 | 14984.51 | 14876.31 | 14337.15 | 14414.02 | 17483.58 | 13582.06 | 14359.52 | 14936.36 | 14231.81 |
| ENSMUSG00000039737 | 550.91 | 518.36 | 530.19 | 474.47 | 491.41 | 485.39 | 549.44 | 552.84 | 465.27 | 465.67 | 521.80 | 469.06 |
| ENSMUSG00000056820 | 1673.20 | 1589.44 | 1636.63 | 1559.85 | 1428.88 | 1457.20 | 1650.12 | 1582.84 | 1551.87 | 1595.01 | 1537.14 | 1572.94 |
| ENSMUSG00000030615 | 649.39 | 656.79 | 679.15 | 581.97 | 625.34 | 594.99 | 580.91 | 594.80 | 500.60 | 577.39 | 511.71 | 546.29 |
| ENSMUSG00000031985 | 2081.75 | 1905.57 | 1982.58 | 1742.82 | 1806.89 | 1870.56 | 2068.27 | 1877.35 | 1904.26 | 1867.11 | 2004.43 | 1960.05 |
| ENSMUSG00000028582 | 823.92 | 755.94 | 796.76 | 748.40 | 703.10 | 707.72 | 693.32 | 767.56 | 697.90 | 763.22 | 755.95 | 730.90 |
| ENSMUSG00000091613 | 435.85 | 462.40 | 444.93 | 390.74 | 419.05 | 411.27 | 437.03 | 426.97 | 405.39 | 425.85 | 456.20 | 413.49 |
| ENSMUSG00000027984 | 1546.44 | 1501.09 | 1581.75 | 1365.52 | 1402.96 | 1440.50 | 1370.45 | 1442.16 | 1358.50 | 1517.58 | 1514.93 | 1409.05 |
| ENSMUSG00000025157 | 1209.07 | 1147.66 | 1144.66 | 1056.44 | 1091.91 | 1038.62 | 1129.45 | 1235.66 | 1051.27 | 1092.84 | 1155.63 | 1201.84 |
| ENSMUSG00000086926 | 65.33 | 64.80 | 68.60 | 59.95 | 58.32 | 62.63 | 71.04 | 81.45 | 92.27 | 89.59 | 104.97 | 74.41 |
| ENSMUSG00000030062 | 6077.53 | 5857.09 | 6044.77 | 5670.88 | 5216.54 | 5488.51 | 6135.56 | 5923.29 | 5787.37 | 5953.09 | 5946.69 | 5929.14 |
| ENSMUSG00000025290 | 23774.80 | 24495.51 | 24750.42 | 22515.01 | 22601.75 | 21455.09 | 22866.03 | 25981.04 | 21533.80 | 21524.91 | 23355.79 | 21605.79 |
| ENSMUSG00000031320 | 28582.80 | 27260.11 | 27332.78 | 25209.86 | 26418.57 | 24364.27 | 25292.20 | 26836.62 | 23616.70 | 23396.44 | 25762.93 | 24348.54 |
| ENSMUSG00000060519 | 2230.93 | 2064.61 | 2113.90 | 1992.97 | 1872.77 | 1990.60 | 2173.48 | 2300.21 | 2232.10 | 2170.19 | 2265.84 | 2267.11 |
| ENSMUSG00000020022 | 3414.65 | 3432.18 | 3256.61 | 3179.66 | 3066.20 | 2987.47 | 3572.71 | 3861.66 | 3318.71 | 3388.02 | 3395.22 | 3470.83 |
| ENSMUSG00000022031 | 2014.47 | 1917.35 | 1956.12 | 1845.15 | 1725.88 | 1811.06 | 1838.96 | 1814.83 | 1864.99 | 1914.68 | 1871.21 | 1891.30 |
| ENSMUSG00000015013 | 706.92 | 697.04 | 662.49 | 660.53 | 618.86 | 609.60 | 652.85 | 730.54 | 622.32 | 611.68 | 639.88 | 652.72 |
| ENSMUSG00000070044 | 710.82 | 710.78 | 742.86 | 645.03 | 667.46 | 668.06 | 696.92 | 692.70 | 652.75 | 754.37 | 624.75 | 692.28 |
| ENSMUSG00000034951 | 673.76 | 651.88 | 687.97 | 638.83 | 604.82 | 599.16 | 667.24 | 654.85 | 636.06 | 649.29 | 684.29 | 654.61 |
| ENSMUSG00000071646 | 1423.58 | 1368.55 | 1456.31 | 1333.47 | 1253.91 | 1301.67 | 1301.21 | 1313.00 | 1236.78 | 1276.45 | 1329.22 | 1295.09 |
| ENSMUSG00000021807 | 5570.50 | 5668.59 | 5558.68 | 5156.09 | 5299.70 | 4935.28 | 5605.00 | 5689.65 | 5033.52 | 5239.64 | 5171.56 | 5210.48 |

| | | | | | | | | | | | | |
|---|---|---|---|---|---|---|---|---|---|---|---|---|
| ENSMUSG00000020423 | 426.10 | 404.48 | 426.31 | 376.27 | 402.85 | 372.65 | 355.20 | 398.18 | 348.46 | 305.29 | 356.28 | 323.06 |
| ENSMUSG00000020180 | 1757.05 | 1798.56 | 1825.78 | 1724.21 | 1611.40 | 1606.47 | 1654.61 | 1967.03 | 1680.46 | 1795.22 | 1696.60 | 1713.28 |
| ENSMUSG00000037972 | 977.98 | 953.27 | 937.88 | 879.68 | 892.10 | 863.26 | 868.67 | 964.18 | 854.95 | 928.03 | 870.00 | 911.74 |
| ENSMUSG00000034192 | 849.27 | 904.19 | 873.20 | 791.81 | 822.98 | 799.58 | 850.69 | 814.45 | 776.43 | 728.93 | 860.92 | 775.17 |
| ENSMUSG00000021477 | 11570.02 | 11307.74 | 11609.33 | 10646.07 | 10670.68 | 10415.44 | 12972.53 | 12716.98 | 11723.93 | 10922.84 | 11571.42 | 11289.38 |
| ENSMUSG00000026000 | 906.80 | 881.61 | 940.82 | 847.63 | 861.86 | 802.71 | 962.19 | 1003.67 | 895.20 | 817.42 | 897.25 | 864.65 |
| ENSMUSG00000061024 | 547.01 | 567.45 | 537.05 | 520.98 | 501.13 | 498.96 | 526.96 | 607.96 | 494.71 | 567.43 | 523.82 | 515.21 |
| ENSMUSG00000025381 | 2475.67 | 2387.60 | 2530.41 | 2313.42 | 2309.10 | 2187.89 | 2469.33 | 2611.18 | 2389.15 | 2441.19 | 2467.69 | 2441.35 |
| ENSMUSG00000028729 | 1490.86 | 1480.47 | 1491.59 | 1352.08 | 1385.68 | 1375.78 | 1382.14 | 1482.47 | 1291.75 | 1340.61 | 1398.86 | 1276.25 |
| ENSMUSG00000009549 | 4659.79 | 4354.04 | 4561.02 | 4261.94 | 4198.07 | 4068.89 | 4704.86 | 4854.63 | 4257.09 | 4399.00 | 4617.46 | 4316.64 |
| ENSMUSG00000074813 | 517.75 | 489.89 | 503.73 | 466.20 | 461.17 | 467.64 | 557.53 | 598.09 | 478.03 | 548.63 | 490.51 | 534.05 |
| ENSMUSG00000101209 | 754.69 | 745.14 | 728.16 | 691.54 | 681.50 | 683.72 | 740.98 | 719.84 | 641.95 | 628.27 | 691.36 | 619.76 |
| ENSMUSG00000029777 | 5000.09 | 4693.72 | 4773.68 | 4465.58 | 4313.63 | 4581.41 | 4880.21 | 4863.68 | 4733.16 | 4542.79 | 5025.21 | 4818.66 |
| ENSMUSG00000038982 | 510.93 | 509.53 | 508.63 | 486.87 | 466.57 | 460.33 | 493.69 | 547.08 | 477.05 | 530.93 | 526.85 | 529.34 |
| ENSMUSG00000026470 | 868.78 | 840.37 | 840.86 | 790.78 | 761.42 | 807.93 | 847.09 | 890.96 | 849.06 | 868.30 | 853.85 | 870.30 |
| ENSMUSG00000038497 | 1022.83 | 1088.75 | 1061.36 | 1004.76 | 963.38 | 972.86 | 1173.52 | 1095.81 | 1164.15 | 1132.66 | 1192.97 | 1116.13 |
| ENSMUSG00000028029 | 2467.87 | 2468.11 | 2471.61 | 2329.96 | 2291.82 | 2249.48 | 2378.51 | 2268.95 | 2172.23 | 2093.87 | 2424.30 | 2172.92 |
| ENSMUSG00000095677 | 3905.10 | 3804.26 | 3711.34 | 3598.31 | 3614.85 | 3394.57 | 3788.52 | 3850.14 | 3485.57 | 3401.29 | 3570.84 | 3526.40 |
| ENSMUSG00000034263 | 790.77 | 793.25 | 777.16 | 723.59 | 714.98 | 757.83 | 736.48 | 791.42 | 769.55 | 732.24 | 757.97 | 739.38 |
| ENSMUSG00000021190 | 3019.75 | 2904.00 | 2974.36 | 2836.47 | 2722.75 | 2736.95 | 3263.36 | 3164.85 | 3306.93 | 3100.43 | 3286.22 | 3124.22 |
| ENSMUSG00000026926 | 2675.55 | 2689.98 | 2681.34 | 2537.73 | 2440.86 | 2525.05 | 2776.87 | 2812.74 | 2697.37 | 2677.89 | 2729.10 | 2815.28 |
| ENSMUSG00000025967 | 14251.42 | 13841.62 | 14491.57 | 13490.81 | 13528.43 | 12804.79 | 13215.32 | 14208.50 | 12579.87 | 13084.18 | 14366.12 | 13370.93 |
| ENSMUSG00000032575 | 3752.99 | 3636.38 | 3742.70 | 3538.35 | 3458.25 | 3420.66 | 3536.74 | 3598.40 | 3199.93 | 3487.57 | 3618.28 | 3403.95 |
| ENSMUSG00000032047 | 1510.36 | 1572.76 | 1553.33 | 1451.31 | 1483.96 | 1403.97 | 1376.75 | 1361.53 | 1301.57 | 1394.81 | 1435.20 | 1394.92 |
| ENSMUSG00000047084 | 576.26 | 549.78 | 571.35 | 546.83 | 518.41 | 529.23 | 555.73 | 602.20 | 470.17 | 486.69 | 523.82 | 533.10 |
| ENSMUSG00000025439 | 1261.72 | 1239.94 | 1206.41 | 1150.51 | 1178.31 | 1161.79 | 1214.88 | 1261.99 | 1175.93 | 1207.87 | 1144.53 | 1207.49 |
| ENSMUSG00000021248 | 8139.77 | 7801.93 | 7875.45 | 7282.41 | 7634.72 | 7552.18 | 7862.11 | 7743.06 | 7507.09 | 7476.20 | 7189.12 | 7397.53 |
| ENSMUSG00000033653 | 1010.16 | 1024.94 | 989.82 | 982.01 | 937.46 | 936.32 | 1028.74 | 987.22 | 993.35 | 1008.77 | 1108.19 | 1084.10 |
| ENSMUSG00000025785 | 326.64 | 315.14 | 322.43 | 303.91 | 304.57 | 307.93 | 324.63 | 363.62 | 294.47 | 344.00 | 306.82 | 317.41 |
| ENSMUSG00000025173 | 1954.99 | 1903.60 | 1929.66 | 1852.39 | 1832.81 | 1837.16 | 1926.19 | 1966.20 | 1848.31 | 1808.49 | 1769.27 | 1815.94 |
| ENSMUSG00000022538 | 1116.44 | 1134.90 | 1155.44 | 1081.25 | 1085.43 | 1091.86 | 1144.74 | 1192.06 | 1182.80 | 1092.84 | 1269.68 | 1196.19 |
| ENSMUSG00000022992 | 3892.42 | 3814.08 | 3898.52 | 3684.10 | 3744.46 | 3674.32 | 3637.45 | 3667.50 | 3508.15 | 3490.88 | 3713.15 | 3646.96 |
| ENSMUSG00000025236 | 647.44 | 641.08 | 659.55 | 620.22 | 633.98 | 610.65 | 631.27 | 658.14 | 598.76 | 600.62 | 523.82 | 599.98 |
